# CD155 regulates tumor growth and immune evasion in diffuse midline glioma

**DOI:** 10.1101/2025.09.09.674382

**Authors:** Theophilos Tzaridis, Ester Calvo Fernandez, Tanja Eisemann, Augusto Faria Andrade, Carlos A. O. de Biagi-Junior, Jennifer L. Hope, Oren J. Becher, Nada Jabado, Jon D. Larson, Suzanne J. Baker, Andrea Califano, Anindya Bagchi, Mariella G. Filbin, Linda M. Bradley, Peter D. Adams, Jovana Pavisic, Robert J. Wechsler-Reya

**Affiliations:** Cancer Genome and Epigenetics Program, NCI-Designated Cancer Center, Sanford Burnham Prebys Medical Discovery Institute, La Jolla, CA, USA; Departments of Systems Biology, Medicine, Biochemistry and Molecular Biophysics, and Biomedical Informatics, Columbia University Irving Medical Center, New York, NY 10032, USA; Department of Pathology and Cell Biology, Columbia University Irving Medical Center, New York 10032, USA; Departments of Human Genetics, Pediatrics and Medicine, McGill University, and Research Institute of the McGill University Health Centre, Montreal, QC, Canada; Cumming School of Medicine, Department of Biochemistry and Molecular Biology, University of Calgary & Charbonneau Cancer Institute, University of Calgary, AB, Canada; Department of Pediatric Oncology, Dana-Farber Boston Children’s Cancer and Blood Disorders Center, Boston, MA 02215, USA; Broad Institute of Harvard and MIT, Cambridge, MA; Immunity and Pathogenesis Program, NCI-Designated Cancer Center, Sanford Burnham Prebys Medical Discovery Institute, La Jolla, CA, USA; The Jack Martin Fund Division of Pediatric Hematology-Oncology, Mount Sinai Kravis Children’s Hospital, Icahn School of Medicine at Mount Sinai, NY, USA; Department of Developmental Neurobiology and Center of Excellence in Neuro-Oncology Sciences, St. Jude Children’s Research Hospital, 262 Danny Thomas Place, Memphis, TN, USA; Chan Zuckerberg Biohub New York, New York, NY 10032, USA; Department of Pediatrics, Memorial Sloan Kettering Cancer Center, New York, NY, USA; Department of Neurology and Herbert Irving Comprehensive Cancer Center, Columbia University Medical Center, New York, NY, USA

## Abstract

Diffuse midline glioma (DMG) is a devastating pediatric brain tumor with an unmet need for novel therapies. Immune checkpoint inhibitors have failed to prolong survival for DMG patients. In this study, we analyzed the expression of immune checkpoint molecules in human and murine DMG cells, as well as primary brain tumor samples, and identified CD155 as the most highly expressed. When murine DMG cells were co-cultured with CD8+ T cells, silencing of CD155 led to a marked increase in T cell-mediated killing. Strikingly, CD155-deficient DMG cells failed to grow in immunocompetent mice, and depletion of CD8+ T cells allowed these tumors to grow. CD155 also exerted cell-autonomous effects on tumor cells: silencing of CD155 led to induction of apoptosis of DMG cells and to delayed tumor growth in immunodeficient mice. Transcriptomic analyses identified FOXM1 as a key target of CD155. Notably, FOXM1 silencing also led to reduced proliferation of DMG cells *in vitro* and *in vivo*. Finally, treatment of DMG-bearing mice with Thiostrepton, a FOXM1-targeting agent, delayed tumor growth and prolonged survival. These studies demonstrate that CD155 regulates immune evasion and tumor growth in DMG, and suggest that targeting CD155 could be a valuable two-pronged therapeutic strategy for this disease.

**Conflict-of-interest statement:** The authors have declared that no conflict of interest exists.

## Introduction

Diffuse midline glioma (DMG) is an aggressive brain tumor that occurs mainly in children and young adults, with a median age at diagnosis of 6-7 years and a median overall survival of as low as 11 months(1). The current standard of care consisting of surgery (if feasible) and focal radiation, remains palliative offering primarily transient symptom relief and outcomes have not improved in decades. There have been several hundred clinical trials over the past 50 years and despite some encouraging results (2), none of these has identified a therapy that can prolong survival in this disease (3). An important molecular hallmark of DMG is a gain-of-function mutation in the gene encoding histone H3.1 (*H3.1K27M)* or histone H3.3 (*H3.3K27M*) (4). This mutation leads to global loss of tri-methylation of histone H3 lysine 27 (K27) and thereby an epigenetic dysregulation of tumor cells. In rare cases, the loss of this tri-methylation is not mediated by a mutation in a histone gene, but by overexpression of EZH inhibitory protein (EZHIP) (5). This has led to the definition of this entity as H3K27-altered DMG based on the most recent WHO classification (6).

Immunotherapy, particularly immune-checkpoint inhibition, is emerging as a powerful approach to treating human cancer. However, this approach has failed to prolong survival of patients with high-grade glioma (HGG), including DMG (7, 8). There are conflicting data on biomarkers of responsiveness to treatment with the programmed cell death 1 (PD1) inhibitor nivolumab, with one study showing that pediatric HGG patients with bi-allelic mismatch repair (MMR) deficiency (who have a high mutational burden), seem to benefit from this treatment (9). However, another study including a very large glioma cohort with more than 10,000 tumor samples demonstrated that MMR-deficient glioma patients do not benefit from PD-1 blockade (10). Notably, large clinical trials using inhibitors of PD-1 did not include screening of tumors and their microenvironment for expression of PD-1 or its ligand PD-L1 and intriguingly, a subset of long-term survivor adult glioblastoma patients treated with nivolumab had tumors with a PD-L1 score of >5% (8).

One potential explanation for the failure of immune checkpoint inhibition to improve survival of HGG and DMG patients is that the checkpoint molecules targeted are not expressed on tumor cells or on cells in the microenvironment. In this study, we investigated expression of immune checkpoint molecules on the surface of human and murine DMG cells, as well as primary pediatric brain tumors. Our studies revealed consistent expression of CD155 / Poliovirus receptor (PVR) in DMG cells and tissues and found compelling evidence for its role not only in immune evasion, but also in cell-autonomous regulation of DMG cell survival.

## Results

To screen for immune checkpoint molecules to target in DMG, we performed flow cytometric analysis of known checkpoints on patient-derived DMG cell lines. We chose checkpoint molecules previously reported to be expressed on the surface of tumor cells: B7-H3, CD155/PVR, PD-L1, PD-L2, CEACAM-1, CD252 (OX-40L), CD86 (B7-2), Galectin-9 and 4-1BB/CD137L. Notably, DMG cells consistently expressed high levels of B7-H3 and CD155/PVR but showed little or no expression of other checkpoint molecules, including PD-L1 and PD-L2, which are most commonly targeted in clinical trials (Figure 1a). To confirm the clinical relevance of these markers, we analyzed their expression on primary pediatric brain tumor samples (high-grade gliomas (HGG), DMGs, medulloblastomas (MB), ependymomas (EP), atypical teratoid rhabdoid tumors (ATRTs), a pleomorphic xanthoastrocytoma (PXA), a CNS high-grade neuroepithelial tumor (CNS-HGNET) and an epidermoid tumor; Figure 1b, Supplementary Table 1). These studies revealed that B7-H3 and CD155 were highly expressed on cells from almost all brain tumors we studied, both on the surface of tumor cells and also on infiltrating immune cells (Figure 1b, Supplementary Figure 1a,b and Supplementary Table 2). By contrast, we saw low expression of PD-L1 and variable, yet overall high levels of HLA-ABC (Figure 1b). Finally, we examined expression of CD155 and B7-H3 in murine models of DMG. To that end, we utilized three murine DMG models, all of which were engineered to express the H3.3K27M mutant protein and are p53-deficient. The first model (A-367 cells) additionally exhibits activation of PDGFRα because of a transgenic PDGFRA mutation (V544ins) (11). Cells from the second model (KAPP cells) overexpress wild-type PDGFRα and are Alpha-Thalassemia/mental retardation, X-linked (ATRX) deficient (12). Cells from the third model (PKC cells) have an overexpression of PDGF-B (13). Interestingly, all three express CD155, but not B7-H3 (Figure 1c-h). These studies identified CD155 as a ubiquitously expressed marker in both murine and human DMG.

**Figure 1.**
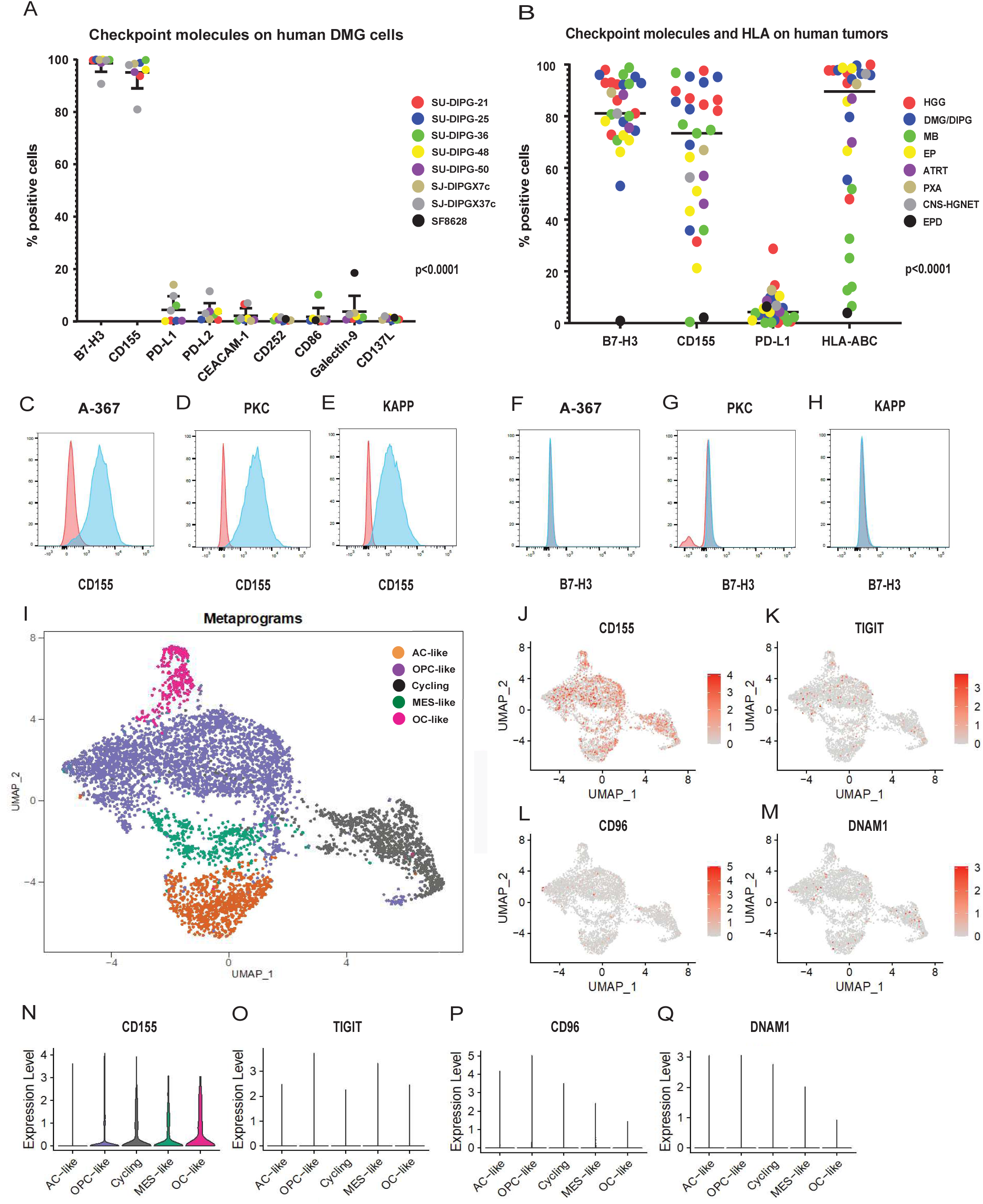
CD155 is highly expressed on human and murine DMG. A: Express ion of checkpoint molecules in human DMG cell lines, as assessed by flow cytometry (comparison between all subgroups; Friedman test); B: Expression of selected checkpoint molecules and HLA-ABC in primary human brain tumors (comparison between all subgroups; 1-way ANOVA test) C-H: Levels of CD155 (C-E) and B7-H3 (F-H) in murine DMG cells; I-M: scRNA sequencing data of human DMG tumors showing different cellular subpopulations (I) and expression of CD155 (J), as well as interaction partners TIGIT (K), CD96 (L) and DNAM1 (M) across tumors. Corresponding violin plots of these genes are shown in N-Q.

DMG is a highly heterogeneous tumor composed of diverse populations of tumor and non-tumor cells (14). To analyze the expression of CD155 and its interaction partners across the different cellular compartments of DMG, we interrogated single-cell RNA sequencing data from human H3-K27M mutant DMG tumors (14), and found high expression of CD155 across tumor samples. Among tumor cells, CD155 was expressed across all cell types including the oligodendrocyte precursor (OPC)-like cells, which are critical for tumor propagation (14) (Figure 1i-j, n). CD155 was also highly expressed in actively cycling (“cycling”), astrocyte-like (“AC-like”) and mesenchymal-like (“MES-like”) cells (Figure 1i-j, n). The immunosuppressive ligands of CD155, CD96 and T cell immunoreceptor with Ig and ITIM Domains (TIGIT) (15), which are expressed mainly on T cells and Natural Killer (NK) cells, were only expressed at low levels in human DMG tumors (Figure 1k-l,o-p and Supplementary Figure 1c-d). Interestingly, the immunostimulatory ligand, DNAX accessory-molecule 1 (DNAM1/CD226), was expressed in microglia and macrophages of human DMG tumors (Figure 1m,q and Supplementary Figure 1d). Notably, the expression levels of CD155 and its interaction partners were consistent across both pediatric and adult H3-K27M mutant DMG tumor samples (Supplementary Figure 1c). Additionally, we were able to confirm the low expression of CD96 and TIGIT, and the expression of DNAM1 in the microenvironment of our murine DMG models, (Supplementary Figure 1e-f). Together these studies suggest that CD155 is broadly expressed across different tumor cell populations in human and murine DMG.

To analyze the role of CD155 in immune evasion, we co-cultured murine DMG cells with CD8+ T cells (experimental outline shown in Figure 2a, gating strategy in Supplementary Figure 2b). To facilitate T cell recognition of tumor cells, we used the ovalbumin (OVA) system: DMG cells from the KAPP and PKC models were engineered to express OVA (DMG-OVA) and then infected with viruses encoding a control shRNA or an shRNA against CD155 (Figure 2b and Supplementary Figure 2a). DMG-OVA cells were co-cultured with CD8+ T cells from OT-I mice, which express T cell receptors that recognize OVA peptides in the context of class I MHC (H-2Kb). We observed dose-dependent CD8+ T-cell-mediated killing of DMG-OVA KAPP and PKC cells with different effector-to-target (E:T) ratios, and saw that lack of CD155 on DMG-OVA cells significantly increased T cell killing, as measured by the percentage of OVA-positive cells remaining (Figure 2c and Supplementary Figure 2c). The above-mentioned results are consistent with the possibility that CD155 acts as an immune checkpoint, and that its loss results in increased T cell activation and killing. However, the CD8+ T cells exposed to CD155-deficient tumor cells did not show elevated levels of interferon gamma compared to T cells cultured with CD155-expressing tumor cells (Figure 2d-e). Moreover, instead of an increase in expression of activation markers (CD25, CD44 and CD69) when incubated with CD155-deficient vs. CD155-expressing tumor cells, those T cells showed a small but significant decrease in CD25 and CD44 (Figure 2f-h and Supplementary Figure 2d-f). These findings suggest that loss of CD155 on tumor cells might not directly affect T cell activation, but rather, might make tumor cells more sensitive to T cell killing.

**Figure 2.**
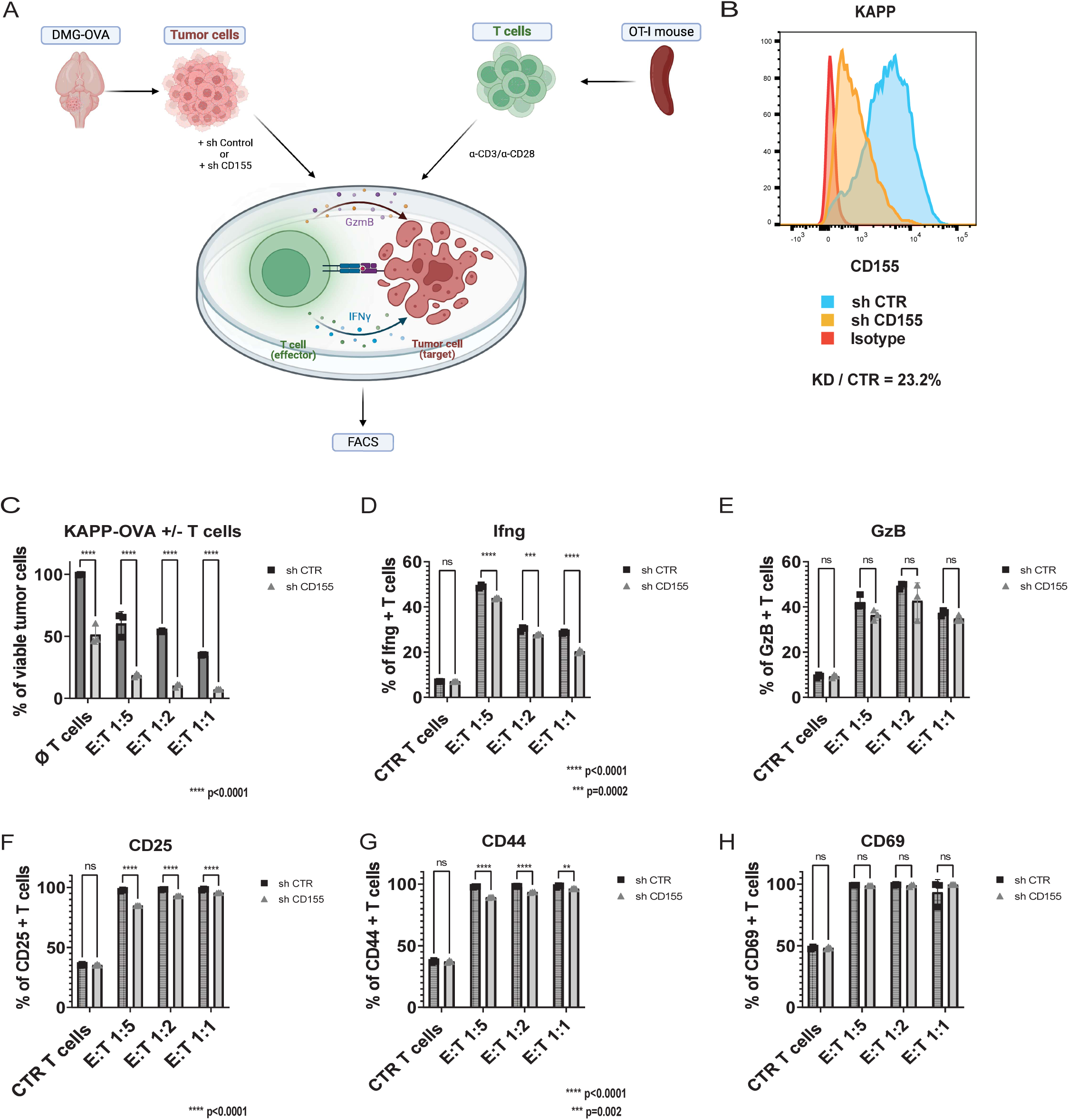
Loss of CD155 increases DMG cell sensitivity to T cell killing. A: Schematic of co-culture of OVA-expressing DMG cells with OVA-re-active (OT-I) T cells, DMG-OVA cells were transduced with either a control shRNA (sh Control) or an shRNA against CD155 (sh CD155) and co-cultured with OVA-reactive T cells isolated from the spleen of OT-I mice and activated with a-CD3/a-CD28 antibodies; after co-culture flow cytometric analysis was performed. B: CD155 levels on the surface of CD155-silenced KAPP cells (knockdown via shRNA1) versus control KAPP cells; C: Percentage of viable KAPP DMG tumor cells after culture without T cells (ø T cells) or with OT-I CD8+ T cells in different Effector (T cells) to Target (Tumor cells) (E:T) ratios; (2-way ANOVA test comparing sh CTR to sh CD155); D-H: Percentage of Interferon gamma (Ifng, D) positive or Granzyme B (GzB, E) positive, as well as CD25 (F), CD44 (G) and CD69 (H) positive CD8 T cells after co-culture with KAPP cells. Control T cells not co-cultured with tumor cells are labelled as CTR T cells (2-way ANOVA test comparing sh CTR to sh CD155).

To determine whether perturbation of CD155 also affects growth of DMG tumors *in vivo*, we transplanted control and CD155-deficient DMG cells (KAPP and A-367, representative plots shown in Figure 2b and Supplementary Figure 3a-c) into syngeneic, immunocompetent mice. As shown in Figure 3, CD155-deficient KAPP cells failed to grow at all (3a-b) and CD155-deficient A-367 cells grew far more slowly (3c) than CD155-expressing cells in immunocompetent mice. Notably, depletion of CD8+ T cells allowed these tumors to grow, highlighting a role for CD8+ T cells in rejection of CD155-deficient cells (Figure 3d-f). To further investigate the role of CD8+ T cells in targeting DMG cells, we injected KAPP-OVA cells expressing or lacking CD155 into immunodeficient mice, and performed adoptive transfer of OT-I CD8+ T cells. Our studies suggested that T cells were more efficient at clearing CD155-deficient than CD155-expressing DMG-OVA tumors, with a subset of CD155-deficient tumors being fully cleared (Figure 3g-i). Together these data suggested that loss of CD155 enhances sensitivity of tumor cells to T cell killing *in vitro* and *in vivo*.

**Figure 3:**
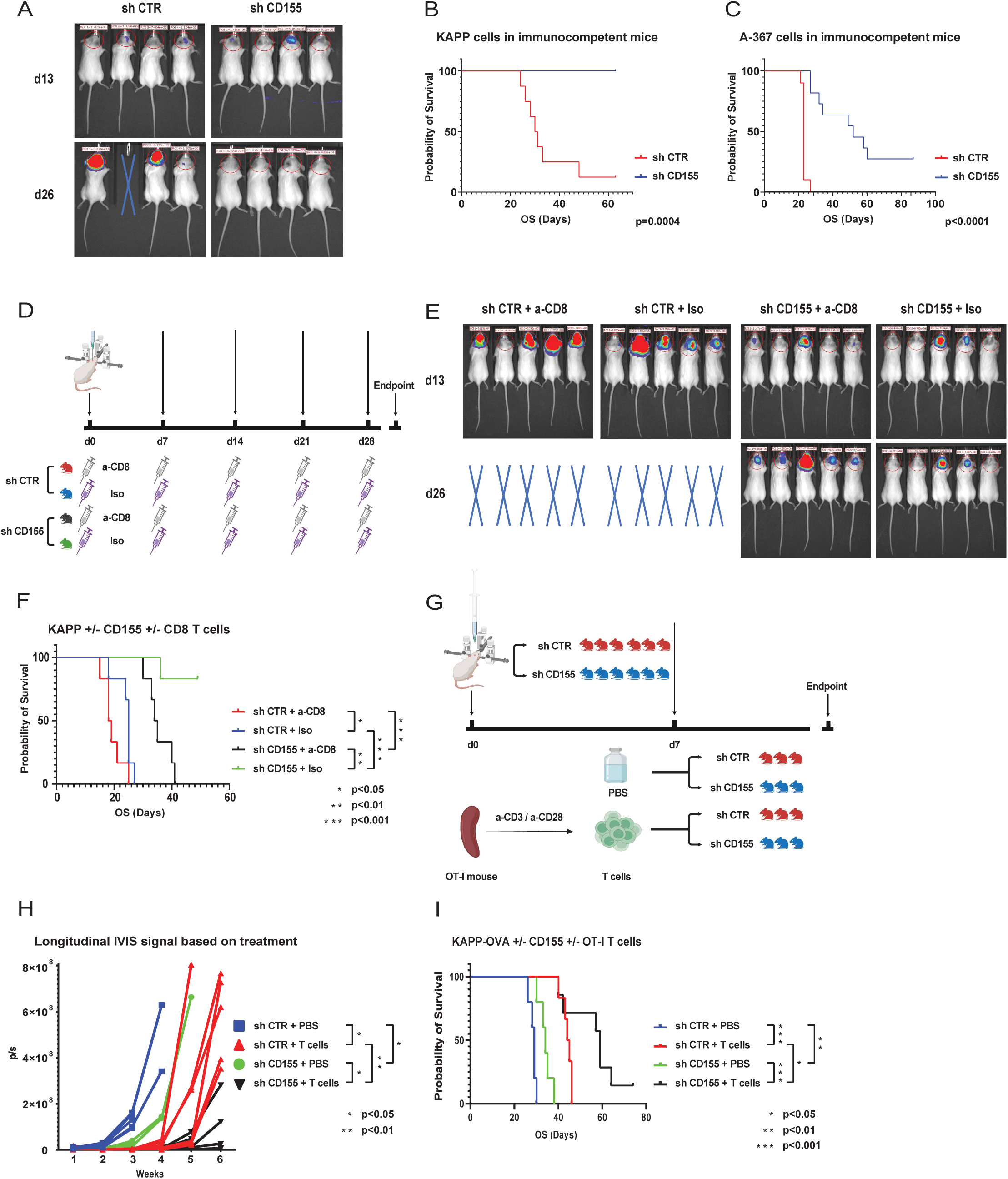
CD155-deficient DMG cells are cleared by CD8+ T cells. A: IVIS images of immunocompetent mice transplanted with control (sh CTR) or CD155-silenced (shCD155) DMG KAPP cells at day 13 (d13) and day 26 (d26) after transplantation; B-C: Survival of immunocompetent mice transplanted with DMG cells (b: KAPP, c: A-367) expressing an shRNA against CD155 (sh CD155) or a control-shRNA (sh CTR); D: Schematic of CD8 T cell depletion experiment: immunocompetent mice were either transplanted with control (sh CTR) or CD155-silenced (sh CD155) DMG cells, and treated weekly with either anti-CD8 antibody (a-CD8) or with isotype control antibody (Iso); E: corresponding IVIS images of subgroups listed in D at day 13 (d13) and day 26 (d26) after transplantation; F: corresponding survival of mice listed in D (Log-rank test with multiple comparisons; *p<0.05, **p<0.01, ***p<0.001); G: Schematic of adoptive T cell transfer in DMG-OVA-bearing mice: immunodeficient (NSG) mice were transplanted with either control or CD155-silenced DMG-OVA cells and were treated with either OT-I CD8 T cells or with PBS; H: corresponding IVIS images 1-6 weeks (x axis) after transplantation of mice listed in G (mixed effects model; *p<0.05, **p<0.01); I: corresponding survival of mice listed in G (Log-rank test with multiple comparisons; *p<0.05, **p<0.01, ***p<0.001).

While studying the effects of CD155 on sensitivity to T cell killing, we observed that the death of CD155-deficient cells was not fully dependent on T cells. For example, Figure 2c shows that even in the absence of T cells, CD155-deficient KAPP cells exhibit a 50% decrease in viability compared to CD155-expressing cells. When cultured in the absence of T cells, CD155-deficient DMG cells exhibited lower cell density (Supplementary Figure 3e-f) than CD155-expressing cells. CD155-deficient cells also showed much lower proliferation and viability compared to wild type DMG cells (Figure 4a-c, representative knockdown plots shown in Supp. Figure 3a-d), an effect that was not observed in CD155-negative murine medulloblastoma cells (Figure 4d). We observed similar effects of CD155 silencing in human DMG cells (Figure 4e-f, representative knockdown plots shown in Supplementary Figure 3g-h). To understand the basis for this reduced proliferation, we performed flow cytometric apoptosis assays and confirmed that CD155 silencing resulted in a marked increase in cell death (Figure 4g-h, Supplementary Figure 3i-p). Notably, CD155-deficient DMG cells (KAPP, A-367 and PKC) exhibited reduced growth in vivo, as monitored by IVIS and demonstrated in tissue sections (Supplementary Figure 4a-d), and were associated with longer mouse survival even in immunodeficient mice (Figure 4i-k). Even though CD155-negative tumors did grow in immunodeficient mice, they had a significantly lower number of tumor cells, as compared to control tumors at the endpoint of the *in vivo* experiment (Supplementary Figure 4e). Strikingly, human CD155-silenced DMG cells (SU-DIPG-6 and SU-DIPG-13) failed to grow in immunodeficient mice (Figure 4l-m and Supplementary Figure 4f-g). Together, these data suggested that in addition to regulating sensitivity to T cell killing, CD155 also exerts cell-intrinsic effects on tumor cell viability.

**Figure 4:**
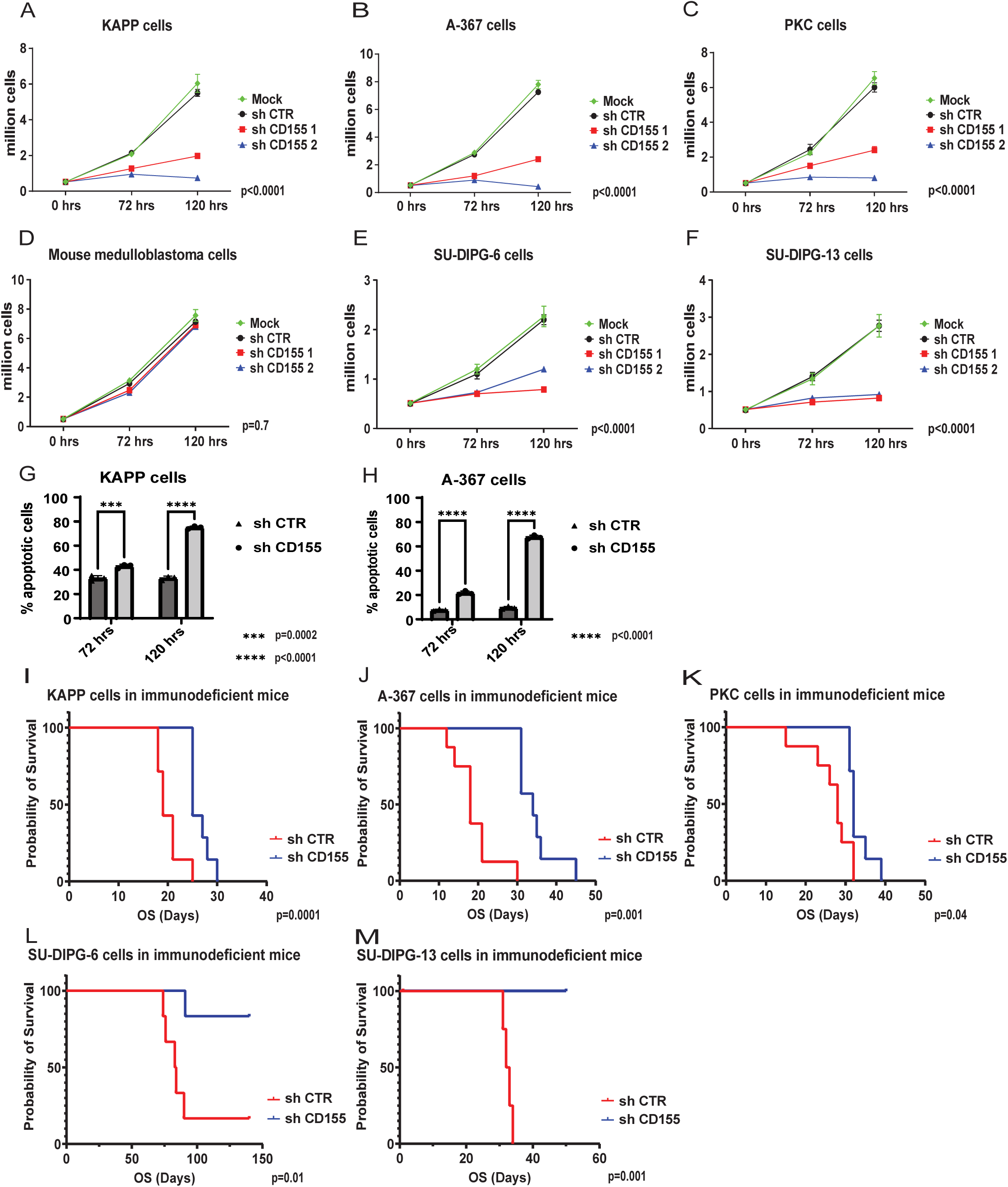
CD155 regulates DMG cell survival and tumor growth even in the absence of T cells. A-C, E-F: cell count of murine DMG cells (A: KAPP, B: A-367, C: PKC), human DMG cells (E: SU-DIPG-6, F: SU-DIPG-13) as well as mouse Group 3 medulloblastoma cells (D) either untreated (Mock), treated with control shRNA (sh CTR) or shRNA against CD155 (sh CD155-1 and sh CD155-2) at different time points: before treatment (0 hrs), after 72-hour exposure to shRNA viruses (72 hrs) and after 48 hours of puromycin selection (120 hrs), 2-way ANOVA test with multiple comparisons (p<0.0001: sh CD155-1 VS sh CTR; p<0.0001 sh CD155-2 VS sh CTR; n.s. Mock VS sh CTR). G-H: apoptotic assay with quantification of total apoptotic cells, i.e. Annexin V positive, 7-AAD positive and double positive DMG cells (G: KAPP, H: A-367) treated with control shRNA (sh CTR) or shRNA against CD155 (sh CD155) for 72 hours (72hrs) and after puromycin selection for 48 hours (120 hrs), (2-way ANOVA test); I-K: Survival of immunodeficient (NSG) mice transplanted with DMG cells (I: KAPP, J: A-367, K: PKC) expressing an shRNA against CD155 (sh CD155) or a control-shRNA (sh CTR). L-M: Survival of immunodeficient (NSG) mice transplanted with human DMG cells (L: SU-DIPG-6, M: SU-DIPG-13) expressing an shRNA against CD155 (sh CD155) or a control shRNA (sh CTR).

To gain deeper insight into the molecular mechanisms underlying the effects of CD155 on intrinsic tumor cell viability, we performed whole transcriptome analyses comparing CD155-expressing and CD155-deficient A-367 and KAPP DMG cells. We found a large number of differentially expressed genes (3,910 for A-367, of which 1,667 were up- and 2243 downregulated; 1,743 for KAPP, of which 915 were up- and 828 downregulated). Ingenuity pathway enrichment analysis (IPA, Qiagen, Hilden, Germany) of the differentially expressed genes revealed significant inactivation of pathways associated with the cell cycle (e.g. kinetochore metaphase pathway), oncogenic processes (e.g. Ephrin receptor signaling), and neuronal regulation (e.g. Netrin signaling). Notably, silencing of CD155 resulted in upregulation of only a few pathways, two of which were immune-related (hypercytokinemia in influenza and interferon signaling, Figure 5a). We validated the differential expression of several genes in CD155-deficient versus CD155-expressing cells, which were enriched in the differentially regulated pathways, by quantitative RT-PCR in A-367 and KAPP cells. As shown in Figure 5b-c, the genes encoding Rho associated coiled-coil containing protein kinase 2 (Rock2), epidermal growth factor receptor (Egfr), SH2 domain containing 3C (Sh2d3c), integrin alpha 4 (Itga4) and forkhead box M1 (FoxM1) were all significantly decreased in CD155-deficient cells compared to control cells. We were also able to validate some of these markers at the protein level (Figure 5d), showing decreased expression of Itga4 and FoxM1 in both cell lines, and decreased phospho-ERK (a target of EGFR signaling) in A-367 cells.

**Figure 5:**
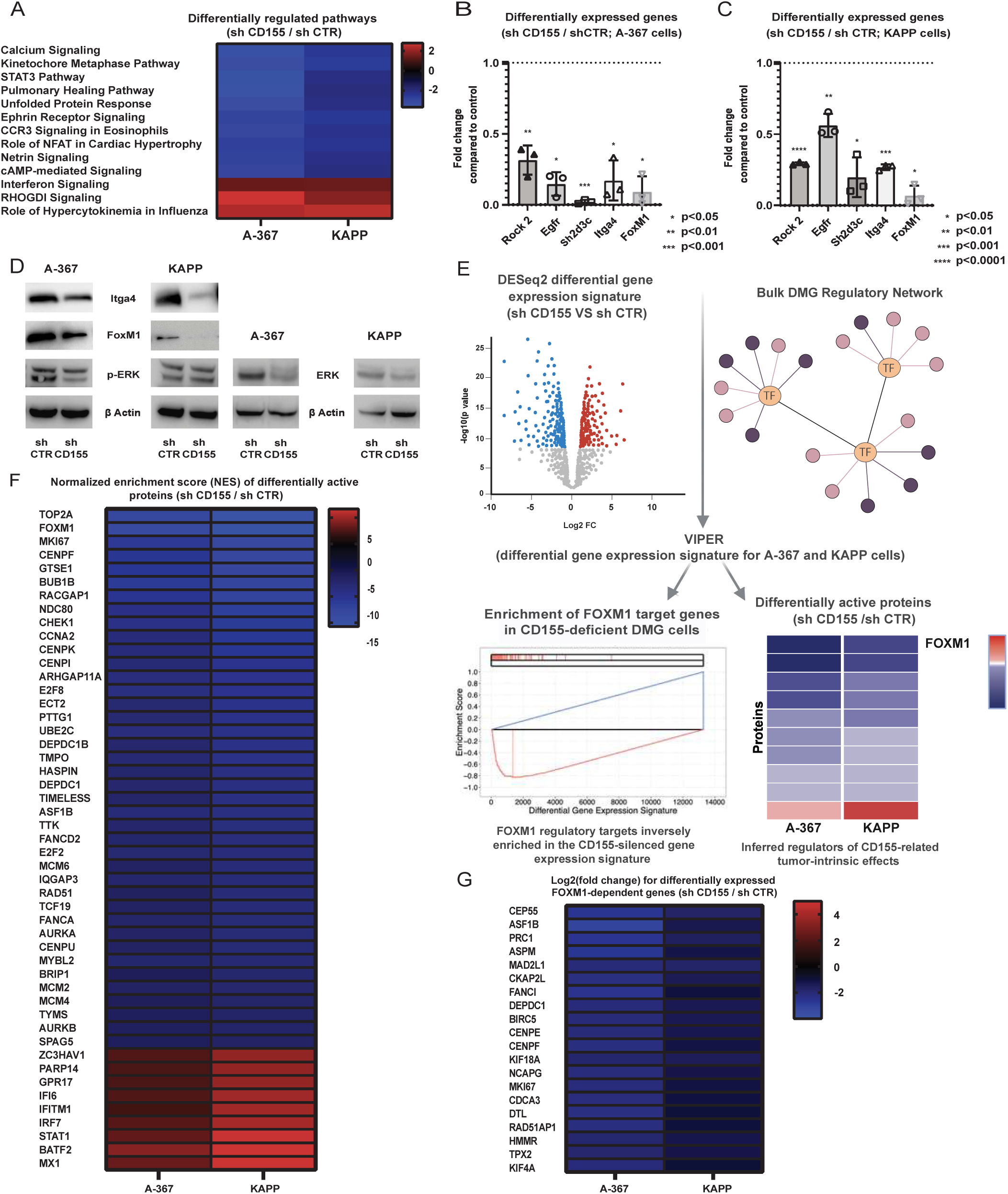
CD155 regulates oncogenic signaling pathways, including FOXM1. A: Ingenuity pathway analysis of genes differentially expressed between CD155-silenced and control DMG cells (A-367 and KAPP; sh CD155 / sh CTR) profiled by RNA seq. Depicted are the top 10 downregulated pathways with a Z-score lower than 2 (blue) as well as three commonly upregulated pathways (red); B-C: Differential expression of individual genes in CD155-silenced versus control A-367 (B) and KAPP (C) cells (unpaired t test); D: Protein levels of putative CD155 targets in CD155-silenced (sh CD155) versus control (sh CTR) A-367 and KAPP cells; E: schematic illustrating the regulatory network analysis of gene expression data: a previously published DMG regulatory network was interrogated with the differential gene expression signature computed by DESeq2 in CD155-deficient versus control A-367 and KAPP cells independently via VIPER to identify differential regulatory protein activity based on the enrichment (aREA) of regulatory protein target genes. The top significantly dysregulated proteins are master regulators driving the CD155-deficient transcriptional phenotype. These studies indicated that CD155 silencing leads to significant downregulation of FOXM1 transcriptional activity as inferred by the significant downregulation of FOXM1-target genes; F: heatmap depicting the VIPER-inferred normalized enrichment scores (NES) of the top differentially regulated proteins / master regulators in CD155-silenced versus control samples for A-367 and KAPP cells based on RNA seq data (inactive in blue, active in red); G: log2(foldchange) based on DESeq2 differential gene expression analysis in CD155-silenced versus control samples of the top 20 FOXM1 target genes in the DMG regulatory network for A-367 and KAPP cells (downregulated in blue, upregulated in red).

To better define the regulatory drivers of decreased proliferation and survival following CD155 knockdown, we performed a Virtual Inference of Protein Activity by Enriched Regulon (VIPER) (16) analysis of differential gene expression signatures between CD155-deficient and control A-367 and KAPP cells. Specifically, we interrogated a DMG gene network (17) to identify transcriptional regulators whose activity was significantly altered by silencing of CD155 based on the differential expression of their target genes (Figure 5e). The most aberrantly active VIPER-inferred transcriptional regulators, also known as Master Regulators (MR), are crucial for implementing and maintaining tumor-associated phenotypes, including tumorigenesis, metastatic progression, and drug resistance, as experimentally validated across various tumor types (18). The transcription factor FoxM1 emerged as one of the most significantly inactivated transcriptional regulators in this analysis, identifying it as a candidate MR of CD155-driven cell autonomous effects in DMG (Figure 5f). Indeed, FoxM1 has been demonstrated to play a major role in transcriptional activation of key oncogenes sustaining the biology and proliferation of DMG cells (16). It was identified to be among the most activated VIPER-inferred MRs in virtually every sample among a large cohort of DMG patient bulk RNA-seq profiles and was validated as a highly essential gene for DMG cell survival in a pooled CRISPR/Cas9-mediated knockout screen (16). Notably, we identified transcriptional and protein downregulation of FoxM1, as well as significantly reduced expression of the majority of FoxM1 positively regulated transcriptional targets, in the absence of CD155 (Figure 5g).

To study whether FoxM1 contributes to the effects of CD155 on cell viability, we used shRNA to silence FoxM1 in our murine DMG models. Our studies showed that FoxM1 knockdown resulted in reduced proliferation *in vitro* (Figure 6a-c). Moreover, silencing of FoxM1 led to delayed tumor growth in immunocompetent mice and to a lesser extent in immunodeficient mice (Figure 6d-e). These studies suggest that FoxM1 is an important target of CD155 and a key regulator of DMG cell survival.

**Figure 6:**
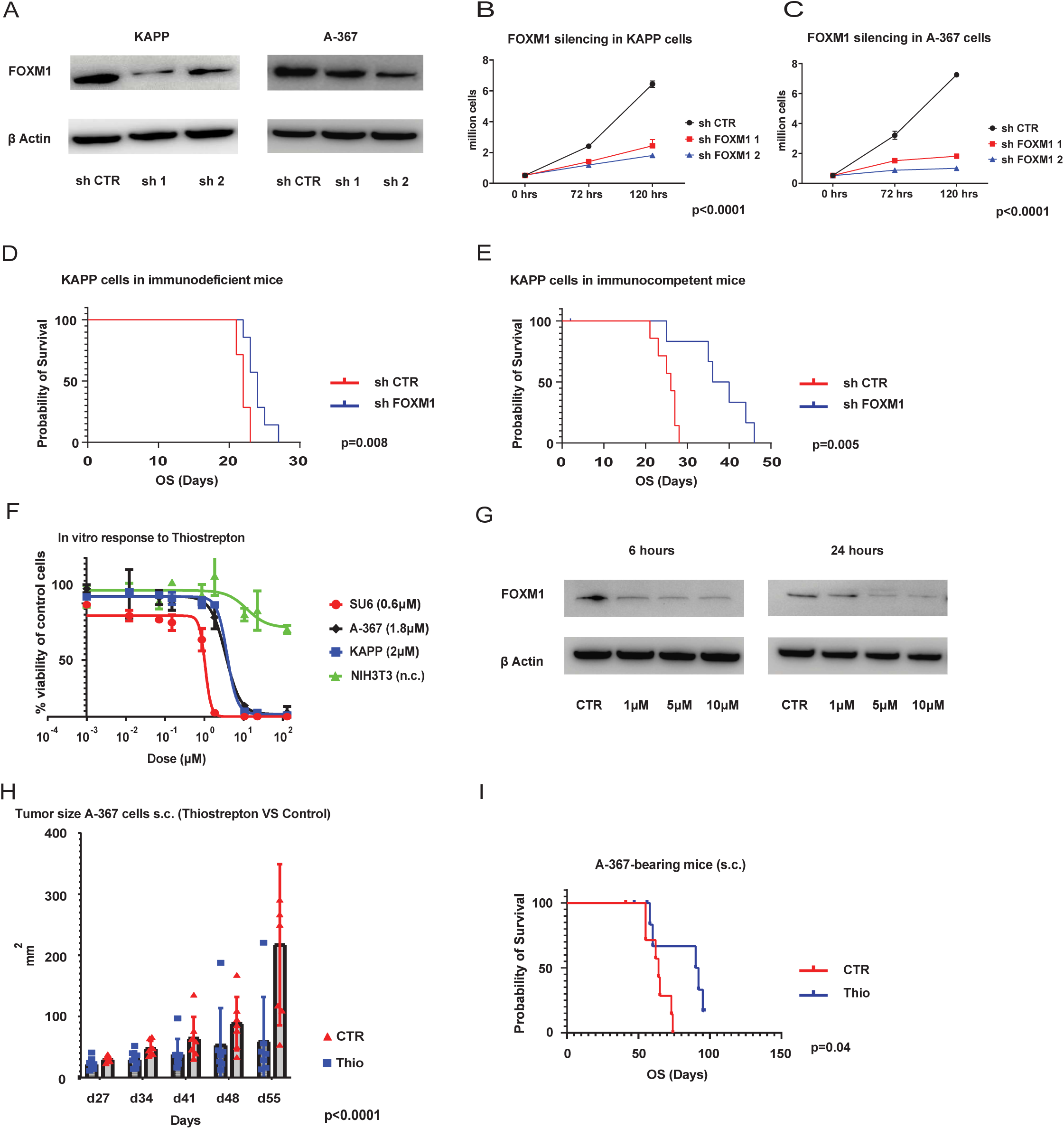
Silencing of FOXM1 and treatment with Thiostrepton lead to impaired tumor growth. A: FOXM1 knockdown via shRNA at the protein level for A-367 and KAPP cells; B,C: cell count of murine DMG cells (B: KAPP, C: A-367) either untreated (Mock), infected with control shRNA (sh CTR) or shRNA against FOXM1 (shFOXM1 1 and shFOXM1 2) at different time-points: before treatment (0 hrs), after 72-hour of exposure to shRNA viruses (72 hrs) and after 48 hours of puromycin selection (144 hrs), 2-way ANOVA test with multiple comparisons (p<0.0001: sh FOXM1-1 VS sh CTR; p<0.0001 sh FOXM1-2 VS sh CTR; n.s. Mock VS sh CTR); D-E: Survival of immunodeficient (NSG, D) or immunocompetent (E) mice transplanted with KAPP cells expressing an shRNA against FOXM1 (sh FOXM1) or a control shRNA (sh CTR); F: Cell viability assay of human (SU6), murine (A-367, KAPP) cells and control murine fibroblast cells (NIH-3T3) treated with different doses of Thiostrepton (Thio) normalized to vehicle control. Depicted are the IC50s for each cell line; G: dose-dependent reduction of FOXM1 levels in Thio treated A-367 cells for 6 and 24 hours; H: tumor size measured by caliper in Thio versus vehicle treated A-367-bearing NSG mice at different time points starting 27 days (d27) after transplantation of cells (2-way ANOVA test); I: Survival of A-367 bearing mice treated with either Thio or vehicle.

Previous studies have shown that the antibiotic Thiostrepton (Thio) can reduce FOXM1 expression as well as its transcriptional activity (19). Notably, our ingenuity pathway enrichment analysis of transcriptional changes in CD155-silenced DMG cells pinpointed Thio as a possible “Upstream Regulator”(20) in A-367 cells (z score 2.5) and to a lesser extent in KAPP cells (z score 0.8), indicating a high overlap between the transcriptional profile associated with CD155 silencing and that associated with Thio treatment (Supplementary Table 3). To test whether Thio could mimic the effects of CD155 and FoxM1 knockdown, we cultured human SU-DIPG-6 and murine KAPP and A-367 cells, as well as the murine fibroblast line NIH3T3, with vehicle or Thio. Thio potently reduced viability of DMG cells with an IC-50 in the low micromolar range, yet only at much higher concentrations for the NIH3T3 cells (Figure 6f). Moreover, Thio caused a dose-dependent reduction in FoxM1 expression in A-367 cells (Figure 6g).

To test whether Thio can also inhibit tumor growth in vivo, we treated tumor-bearing mice with vehicle or Thio. Given its complex molecular structure and a molecular weight of 1665 g/mol, we hypothesized that Thio might not be able to cross the blood-brain barrier and target DMG cells in the brain. Indeed, systemic administration of Thio via intraperitoneal (i.p.) injection failed to prolong survival of mice with orthotopically injected DMG tumor cells. Therefore, we injected mice with tumor cells subcutaneously (s.c.), and then treated them i.p. with Thio. Thio caused a striking delay in tumor growth and also a survival benefit for Thio treated mice (Figure 6h-i). These data indicate that Thio not only reduces FoxM1 expression and impairs viability of DMG cells *in vitro*, but also potently targets DMG tumors *in vivo*. Future experiments are warranted to identify the appropriate delivery strategy to the brain or alternative compounds with sufficient BBB permeability.

## Discussion

In this study, we identified CD155 as a novel regulator of tumor growth and immune evasion in DMG. We showed that CD155-deficient tumors are cleared by the immune system, and pinpointed CD8+ T cells as key players in this tumor cell rejection. Moreover, we showed that CD155 directly affects survival of DMG cells and that its silencing leads to downregulation of key oncogenic pathways, including FOXM1. Finally, we demonstrated that the FOXM1-targeting agent Thiostrepton efficiently targets DMG cells *in vitro* and *in vivo*. Our results suggest that targeting CD155 and thereby its transcriptional target FOXM1 could be a valuable therapeutic strategy for this devastating disease.

In this study, shRNA-mediated silencing of CD155 was found to sensitize tumor cells to CD8+ T cells. Notably, the known inhibitory interaction partners of CD155 on T cells (CD96 and TIGIT) showed very low expression in the microenvironment of human and murine DMG tumors. In contrast, we saw high levels of DNAM1 in all immune cell types of our murine model and also in microglia and macrophages of human tumors. Interestingly, although DNAM1 is generally considered a tumor-suppressive ligand of CD155 on T and NK cells (21), recent data suggest an opposing, potentially tumor-promoting function of DNAM1 (22). Further studies are needed to better understand the interactions between CD155 and its ligands, and to determine how these interactions contribute to immune evasion.

Although knockdown of CD155 increased tumor cell killing by CD8+ T cells, we did not observe an increase in activation markers on T cells exposed to CD155-negative tumor cells. These data suggest that CD155 may not act by inhibiting T cell activation, as a conventional immune checkpoint would. Rather, it appears to act within tumor cells themselves to enhance susceptibility to T cell killing. Our transcriptome studies revealed that silencing of CD155 led to upregulation of type I and type II Interferon in tumor cells, which could potentially recruit and activate CD8 T cells (23, 24). We also saw a downregulation of IL-4, a cytokine known to inhibit cytotoxic CD8 T cells (25). While these mechanisms may contribute to increased susceptibility of tumor cells to T cells, the tumor-cell intrinsic functions of CD155 may be more important drivers of immune evasion.

Indeed, silencing of CD155 led to apoptosis in the absence of T cells, and to delayed tumor growth even in immunodeficient mice, suggesting a key role for CD155 in regulating DMG cell survival. In our transcriptomic analyses of CD155-deficient cells, we found evidence for downregulation of key oncogenic pathways, including integrins and components of the mitogen-activated protein (MAP) kinase family, such as Rock2. Notably, both Itga4 and Rock2 are emerging as markers of biologically aggressive tumors and exciting therapeutic targets in cancer, and could contribute to the survival-promoting effects of CD155 (26–30). Future studies are warranted to understand the link between CD155 and these pathways and to evaluate the therapeutic potential of targeting these molecules in DMG.

Perhaps most excitingly, the transcriptomic profile of CD155-deficient DMG cells showed not only low expression of Forkhead box M1 (FoxM1), but also low expression of FoxM1 transcriptional targets, implicating FoxM1 as a master regulator of CD155-driven cell autonomous effects in DMG. Of note, silencing of FoxM1 in our murine DMG models also led to reduced cell viability and delayed tumor growth *in vivo* thereby underlining not only its importance as a target of CD155, but its autonomous role in DMG cell survival. FOXM1 has been identified as one of the most important transcriptional regulators of DMG biology and DMG cell proliferation (17). As a DMG master regulator, FOXM1 integrates upstream (epi)genetic signals to implement key transcriptional programs driving DMG cell survival independent of tumor genetic profiles, making it an attractive therapeutic target for the disease (18). This key homeostatic feature of FOXM1, implementing tight autoregulatory networks at a genomic and epigenetic level makes it a very attractive therapeutic target. Indeed, our data along with recent literature from other tumor entities (31) support the notion of targeting FOXM1 in DMG. Intriguingly, we found evidence for Thiostrepton downregulating FoxM1 expression and reducing viability of DMG cells. Thiostrepton has already been used in combination with proteasome inhibitors or chemotherapy and showed synergistic effects in breast carcinoma, leukemia and medulloblastoma, and its cytotoxic function has been mostly attributed to the targeting of FOXM1 (32–34). Considering its high molecular weight (1665g/mol) and complex structure, we hypothesized that Thiostrepton might not be capable of crossing the blood-brain barrier. Indeed, its failure to prolong survival of mice with orthotopically injected DMG tumor cells, and its marked effects on tumor size of subcutaneously transplanted DMG cells supports this notion, and future studies should focus on enhancing delivery of the agent to the brain.

In addition to targeting FOXM1, our studies suggest that targeting CD155 itself might have a potential benefit due to this double-pronged feature we have unraveled. Selecting a modality to therapeutically target CD155 poses a formidable challenge. A potential therapeutic approach could involve Antisense Oligonucleotides (ASOs) against CD155. Recent data have shown that ASOs against H3K27M mRNA led to delayed tumor growth in DMG-bearing mice (35). Based on our data showing pronounced effects on DMG cell proliferation and immune evasion following shRNA-mediated silencing of CD155, an ASO against CD155 could be a promising future therapeutic strategy. Another attractive approach might be a chimeric antigen receptor (CAR)-T cell. CAR-T therapies against disialoganglioside GD2 (36) or B7-H3 (37) have shown promising results for DMG patients. However, monotherapies are unlikely to substantially prolong survival of DMG patients; therefore multiantigen CAR-T targeting might be necessary to achieve durable responses (37). Based on our data, CD155 could be an effective target for CAR-T cells and could be combined with targeting of other antigens to enhance responses.

Future studies are necessary to further discern the mechanism of action of CD155 in inhibiting survival of tumor cells, on their own and in the presence of T cells. Such studies may help develop novel therapies against CD155 that exploit the dual function we have unraveled. DMG has an unmet need for novel therapeutic concepts and immunotherapy has so far failed to transfer the survival benefit it has brought about in other cancers to neuro-oncology. Our studies demonstrate that CD155 is expressed across human and murine DMG and functions not only as a modulator of T cell responses, but also as a regulator of tumor cell survival. Disrupting CD155 and its target FOXM1 represent very attractive therapeutic strategies, since they may weaken tumor cells and render them more susceptible to immunological attack, paving the way for therapies to improve outcomes for DMG patients.

## Materials and Methods

### Cell lines

Murine DMG cells (A-367, PKC and KAPP) and human DMG cells (SU-DIPG-13/21/25/36/48/50) were cultured in suspension flasks (Genesee Scientific, El Cajon, CA, USA) with Neurobasal-A medium supplemented with serum-free B-27 and N-2 Supplement, GlutaMAX, 100U/mL Penicillin-Streptomycin, 2μg/mL Heparin (all ThermoFisher Scientific, Waltham, MA, USA) and recombinant epidermal growth factor, as well as basic fibroblast growth factor (both 20ng/mL, Peprotech, Cranbury, NJ, USA). Human DMG cells SJ-DIPGX7c and SJ-DIPGX37c were cultured in adherent flasks with Neurobasal-A medium and the above-mentioned supplements, as well as KnockOut DMEM/F12, Sodium Pyruvate, Non-Essential Amino Acids, HEPES and recombinant Insulin (all ThermoFisher Scientific) and PDGF-AA/BB (Cell Guidance Systems, Cambridge, UK). Dissociation was performed enzymatically with Accutase (ThermoFisher) as indicated.

### Primary tumor cells and tissue dissociation

Human tumors were obtained from Rady Childrens’ Hospital after surgical tumor dissection. Both human and murine tumors were enzymatically dissociated via DNase (2500U, Worthington, Lakewood, NJ, USA) and Liberase (100μg/ml, Sigma-Aldrich, St. Louis, MO, USA). Enzymatic activity was inhibited using 1% FBS. Subsequently, the suspension was passed through a 70-micron filter (Corning, Corning, NY, USA) and remaining erythrocytes were removed using Red blood cell lysis buffer (Invitrogen, Waltham, MA, USA). At the end of the tumor dissociation, cell yield was counted using Trypan blue (Invitrogen) staining.

### Ethics Statement

The studies were approved by the institutional review boards at SBP and UCSD and were conducted in accordance with the tenets of the Declaration of Helsinki. Written informed consent was obtained by all donors.

### Flow cytometry

Cells were resuspended in FACS buffer (5% fetal bovine serum (FBS) in PBS) and stained with fluorophore-conjugated antibodies. For surface staining, 7-Aminocactinomycin D (7-AAD, Biolegend, San Diego, CA, USA) was used as a viability stain. For intracellular staining, fixation and permeabilization were performed using the eBioscience Foxp3/Transcription Factor staining buffer set based on the manufacturer’s guidelines (Invitrogen) and viability was tested using eFluor780 fixable dye (ThermoFisher Scientific, Waltham, MA, USA). A list of antibodies used can be found in Supplementary Table 6). Whenever human or murine material was analyzed ex vivo, an Fc receptor blocking step was included (Fc16/32, Biolegend, San Diego, CA, USA). Compensation was performed using UltraComp Beads (ThermoFisher Scientific). Apoptosis was measured by staining cells with Annexin V and 7-AAD (both Biolegend). For analysis, FlowJo Software Version 10.8 (FlowJo LLC, OR, USA) was used.

### Single-cell RNA sequencing

Single-cell RNA sequencing (scRNA-seq) analysis was conducted following the approach described by Liu et al. (38). The cohort included 18 patients, comprising 4 adults and 14 pediatric cases. Cell states were previously defined in the study by Ilon et al. In this study, we utilized the Seurat R package (v5.1.0) to generate Feature Plots depicting the expression of specific genes.

### Virus production and transduction of cells

293T cells were transfected for production of lentiviral production. 2^nd^ generation packaging plasmids psPAX2 and pMD2.G (both Addgene, Watertown, MA, USA) together with the plasmid of the gene of interest and polyethylenimine were mixed and incubated in OptiMEM (Invitrogen). Virus-containing supernatant was concentrated using the Optima-LX-80K ultracentrifuge (Beckman-Coulter, Carlsbad, CA, USA) and resuspended in Neurobasal medium (Invitrogen). Plasmids used in this study is provided in Supplementary Table 7. Lentiviruses were used to transduce tumor cells for 72 hours and subsequently, Puromycin selection was performed (for A-367, PKC, MP, SU-DIPG-6 and SU-DIPG-13 cells at a dose of 1.5μg/ml and for KAPP cells at a dose of 5μg/ml). For RNA experiments and Western blot studies on CD155, A-367 and KAPP cells were treated with shRNAs against CD155 or control shRNA for 48 hours.

### Isolation of CD8 T cells and co-culture experiments

To isolate CD8 T cells from either OT-I (C57BL/6-Tg(TcraTcrb)1100Mjb/J; 003831) or C57BL/6 (000664) mice (both Jackson Laboratory, Bar Harbor, ME, USA), we removed the spleen from a mouse of the corresponding strain, performed red blood cell lysis and subsequently used the negative selection MagniSort Mouse CD8 T cell Enrichment kit (ThermoFisher). After purification, OT-I CD8 T cells were activated in anti-CD3 and anti-CD28 coated plates for 48 hours and were thereafter cultured in RPMI media supplemented with IL-2 (Biolegend). DMG cells were lentivirally transduced with an ovalbumin-GFP construct and sorted for cells containing high levels of GFP. DMG-OVA cells were then transduced with shRNA against CD155 or a control shRNA for 48 hours and then co-cultured with OT-I CD8 T cells for 24 hours before analysis by flow cytometry. Tumor cell death was measured by percentage of OVA-positive cells and by positivity for 7-AAD, while CD8 T cells were analyzed for activation markers (CD25, CD44, CD69) using antibodies described above.

### In vivo experiments

#### a. Stereotactic injection of DMG cells

All animal experiments were approved by the Institutional Animal Care and Use Committee of Sanford Burnham Prebys Medical Discovery Institute, La Jolla. For experiments with immunocompetent mice and DMG KAPP cells, female 5 week-old B6(Cg)-Tyr^c-2J^/J (albino B6, 000058, Jackson Laboratories) were transplanted with KAPP tumor cells (300,000 cells per mouse). For experiments with immunocompetent mice and A-367 cells, 4-5 day old FVB-NJ pups (001800, Jackson Laboratories) were transplanted with A-367 tumor cells (25,000 cells per mouse). For experiments with immunodeficient mice, 8-10 week old NOD.Cg-Prkdcscid Il2rgtm1Wjl/SzJ (NSG, 005557, Jackson Laboratories) mice were used and each mouse was transplanted with 25,000 cells (applicable to PKC, A-367 and KAPP cells) or with 300,000 cells (applicable to SU-DIPG-6 and SU-DIPG-13). For all experiments with intracranial tumors, cells were stereotactically injected into the pons. For flank tumors, tumor cells were subcutaneously (s.c.) injected into the left flank of NSG mice. Tumor growth was monitored by bioluminescence imaging (BLI) using the in vivo imaging system (IVIS, PerkinElmer, Waltham, CA, USA) and flank tumors were additionally measured by digital caliper (ThermoFisher Scientific). Animals were kept alive until they displayed signs of morbidity or toxicity (>20% weight loss), whereupon they were euthanized.

#### b. Adoptive T cell transfer

For adoptive transfer experiments, CD155-deficient or CD155-wild type DMG-KAPP cells expressing OVA were transplanted into NSG mice. CD8+ T cells were purified from OT-I mice and activated, as previously described. Eight days after tumor transplantation, mice from each group (CD155-deficient and CD155-wild type) were randomized based on IVIS signal and 3 million CD8 T cells or PBS were intravenously injected into the NSG mice.

#### c. Immune cell depletion

For depletion of immune cells, tumors were transplanted into albino B6 mice. Immediately prior to tumor transplantation, 250 μg of either anti-CD8 (clone BE0004-1, BioXcell, Lebanon, NH, USA) or the corresponding isotype controls were injected i.p. into each mouse. Thereafter, antibodies were applied weekly at a dose of 200μg per mouse. Treatment was continued until the mice were euthanized per protocol, as mentioned above.

#### d. Treatment with Thiostrepton

NSG mice s.c. transplanted with DMG cells were randomized into two groups based on IVIS signal at day 7 after transplant. Mice were treated i.p. with either 100mg/kg Thiostrepton (Sellekchem) or with the corresponding DMSO control 3 times per week and was continued until the mice were euthanized per protocol, as mentioned above or if the tumors reached a surface area of 2cm^2^.

### H&E staining

Brains from A-367-bearing NSG mice were fixed in 4% neutral-buffered formalin, processed, and embedded in paraffin. Five-micrometer-thick sections were deparaffinized in xylene and rehydrated to water. Sections were stained with Harris Hematoxylin (Cancer Diagnostics, Durham, NC, USA) followed by washing steps with water and 70% Ethanol. Sections were then stained with Eosin Y (Cancer Diagnostics), dehydrated through a series of alcohol solutions and cleared with xylene. Subsequently, slides were covered with Cover-Seal-T (Cancer Diagnostics) and visualization was performed using the Aperio Leica AT2 microscope (Deer Park, IL, USA).

### Cell viability assay

Cells were seeded in 96-well plates (25,000 cells per well) and were treated for 72 hours with different concentrations of Thiostrepton (Sellekchem, Houston, TX, USA) or a DMSO control. After treatment, cell viability was determined using the CellTiter-Glo assay kit (Promega, Madison, WI, USA).

### RNA sequencing and quantitative reverse-transcription polymerase chain reaction (qRT-PCR)

RNA was extracted from cells using the RNeasy Mini Kit (Qiagen, Hilden, Germany). mRNA sequencing was performed by Novogene (Durham, NC, USA). Briefly, a library was constructed and after quality control (QC), sequencing was performed. Raw data stored in FASTQ format files were quality-checked by demonstrating i) an error distribution rate of less than 1% and ii) an equal GC distribution across reads. Low quality reads (s.a. adapter contamination) were excluded from downstream analysis. Alignment to the mus musculus reference genome GRCm38/mm10 was performed using HISAT2 (39). Gene expression levels were calculated using the Fragments Per Kilobase of transcript sequence per Millions base pairs sequenced (FPKM). For differential gene expression (DEG) analysis, DESeq was used as a normalization method (40) and a gene was considered differentially expressed if the absolute value of log2(foldchange) was greater than 1 and the adjusted p-value ≤ 0.05. Data were further analyzed using Ingenuity Pathway Analysis (IPA, Qiagen, Hilden, Germany) to analyze differentially regulated pathways, as well as differentially regulated “Upstream Regulators” (20). We applied the Virtual Inference of Protein Activity by Enriched Regulon Analysis (VIPER) algorithm to assess the differential activity of regulatory proteins following CD155 silencing, using a recently published DMG-specific gene regulatory network (16). First, we generated a differential gene expression signature of humanized genes (using the homologene package) with DESeq2, comparing CD155-deficient versus wild-type control samples independently for each cell line (A-367 and KAPP). Differential protein activity was then computed on this differential gene expression signature (Wald statistic) for each cell line using VIPER and the DMG gene regulatory network. Briefly, VIPER computes the activity of regulatory proteins (transcription factors, co-transcription factors, and signaling proteins) by assessing the enrichment of their inferred transcriptional targets in the network within the provided differential gene expression signature, using the analytic rank-based enrichment analysis (aREA) algorithm. This yielded normalized enrichment scores (NES; analogous to z-scores) for 6,138 regulatory proteins, representing their differential activity in CD155-deficient versus wild-type DMG cells for each of the two cell lines. A highly significant correlation was observed in differential gene expression and in protein activity between the two cell lines (Pearson correlation p-value < 2.2 x 10^-16^). We thus generated a combined CD155-deficient vs. wild-type protein activity signature by integrating NES scores across the two cell lines using Stouffer’s method. NES scores were transformed into p-values using a normal distribution with Bonferroni correction for multiple hypothesis testing.

To visualize the results, we generated a heatmap displaying the NES scores in each cell line for proteins with statistically significant differential protein activity in the integrated signature (Bonferroni-adjusted p-value < 0.01; Figure 5E). We also generated a heatmap displaying the NES scores for the top 100 FOXM1 target genes within the DMG regulatory network (Figure 5F).

### Validation of gene expression by quantitative RT-PCR

For qRT-PCR, cDNA was synthesized from extracted RNA using SuperScript Vilo kit (Invitrogen). Samples were then mixed with primers for individual genes and PowerUP SYBR Green Master Mix and real-time PCR was conducted in the QuantStudio 7 Pro machine (ThermoFisher Scientific). Beta-Actin and hypoxanthine-guanine phosphoribosyltransferase were used as reference genes for normalization. A list of primers used for qRT-PCR is listed in Supplementary Table 8.

### Western Blotting

For Western blot analyses, we used cells transduced with shRNAs against CD155, FOXM1 or control shRNAs, as well as tumor cells treated with Thiostrepton. Cells were harvested and lysis was performed using radioimmunoassay precipitation (RIPA) buffer. Protein concentration was measured using Bicinchonic acid (BCA) assay (ThermoFisher Scientific). 10μg of protein were loaded on a 4-20% Tris-Bis gradient gel (Invitrogen) and electrophoresis was performed at 120V for 90 minutes. A polyvinylidene fluoride (PVDF) membrane (ThermoFisher Scientific) was used for blotting, which was carried out in the Trans-Blot Turbo Transfer System (BioRad, Hercules, CA, USA). Primary antibody was applied overnight at 4°C with a list of antibodies found in the Supplementary Table 9. After exposure to the secondary antibody (HRP-linked mouse or rabbit, both Cell Signaling Technology, Danvers, MA, USA) for one hour at room temperature, chemiluminescent substrate SuperSignal West Femto (ThermoFisher Scientific) was applied and imaging was conducted in ChemiDoc Imaging Systems (BioRad).

### Statistical analysis

For statistical analysis, GraphPad Prism software version 9 (La Jolla, CA, USA) was used. For IC-50) determination, a non-linear regression analysis was applied. Survival studies were performed using Log-rank tests and survival was visualized via Kaplan Meier curves. Statistical tests used were non-parametric Friedman test, One-Way or Two-Way ANOVA, Wilcoxon rank test and Mixed-effects model (linear mixed model with restricted maximum likelihood).

## Data availability

Values for all data points in graphs are reported in the Supporting Data Values file. All data that support the findings of our study are available from the corresponding author upon request.

## Author contributions

TT designed the study, conducted experiments and acquired and analyzed data. ECF and JP conducted, and JP and AC supervised the bioinformatic part of the study. AFA, OJB, NJ, JDL and SJB provided reagents and helped with data analysis. JLH and LMB provided reagents and helped with immunology experiments. CAOBJ and MGF conducted the scRNA seq data analyses. TE and AB helped with data interpretation. RWR designed and supervised the study. TT and RWR wrote the manuscript with input from all co-authors.

## Acknowledgements

We gratefully acknowledge the Animal Facilities at UCSD and SBP for help with animal care and husbandry. We are highly appreciative of Yoav Altman and the flow cytometry core facility for their help with experimental planning and procedures. We gratefully thank Guillermina Garcia and Monica Sevilla from the histology core facility for their help with our experiments. We remain indebted to Dr. Meher Beigi Masihi for his valuable input and help with experiments and Bryan Hall for excellent technical assistance. We gratefully acknowledge Dr. Michelle Monje for her kind contribution of the patient-derived DMG lines. This work was supported by funding from the German Research Foundation (DFG) to TT (Reference number: TZ 102/1-1; project number: 454298163), the ChadTough Defeat DIPG Foundation to TT, the National Institute for Neurological Disorders and Stroke (R35 NS122339 to R.J. W-R) and the National Institute on Aging (P01 AG073084 to P.D.A.). Additional support was provided by the Swim Across America and Team Jack Foundations to JP. SBP’s Shared Resources are supported by SBP’s NCI Cancer Center Support Grant P30 CA030199. R.J.W-R’s laboratory was also funded by Ian’s Friends Foundation, Alex’s Lemonade Stand Foundation, the V Foundation, William’s Superhero Fund and the McDowell Charity Trust.

## References

1. Mackay A, Burford A, Carvalho D, Izquierdo E, Fazal-Salom J, Taylor KR, et al. Integrated Molecular Meta-Analysis of 1,000 Pediatric High-Grade and Diffuse Intrinsic Pontine Glioma. Cancer Cell. 2017;32(4):520–37 e5.

2. Venneti S, Kawakibi AR, Ji S, Waszak SM, Sweha SR, Mota M, et al. Clinical Efficacy of ONC201 in H3K27M-Mutant Diffuse Midline Gliomas Is Driven by Disruption of Integrated Metabolic and Epigenetic Pathways. Cancer Discov. 2023;13(11):2370–93.

3. Aziz-Bose R, and Monje M. Diffuse intrinsic pontine glioma: molecular landscape and emerging therapeutic targets. Curr Opin Oncol. 2019;31(6):522–30.

4. Schwartzentruber J, Korshunov A, Liu XY, Jones DT, Pfaff E, Jacob K, et al. Driver mutations in histone H3.3 and chromatin remodelling genes in paediatric glioblastoma. Nature. 2012;482(7384):226-31.

5. Sievers P, Sill M, Schrimpf D, Stichel D, Reuss DE, Sturm D, et al. A subset of pediatric-type thalamic gliomas share a distinct DNA methylation profile, H3K27me3 loss and frequent alteration of EGFR. Neuro Oncol. 2021;23(1):34–43.

6. Louis DN, Perry A, Wesseling P, Brat DJ, Cree IA, Figarella-Branger D, et al. The 2021 WHO Classification of Tumors of the Central Nervous System: a summary. Neuro Oncol. 2021;23(8):1231–51.

7. Omuro A, Brandes AA, Carpentier AF, Idbaih A, Reardon DA, Cloughesy T, et al. Radiotherapy combined with nivolumab or temozolomide for newly diagnosed glioblastoma with unmethylated MGMT promoter: An international randomized phase III trial. Neuro Oncol. 2023;25(1):123–34.

8. Lim M, Weller M, Idbaih A, Steinbach J, Finocchiaro G, Raval RR, et al. Phase III trial of chemoradiotherapy with temozolomide plus nivolumab or placebo for newly diagnosed glioblastoma with methylated MGMT promoter. Neuro Oncol. 2022;24(11):1935–49.

9. Bouffet E, Larouche V, Campbell BB, Merico D, de Borja R, Aronson M, et al. Immune Checkpoint Inhibition for Hypermutant Glioblastoma Multiforme Resulting From Germline Biallelic Mismatch Repair Deficiency. J Clin Oncol. 2016;34(19):2206–11.

10. Touat M, Li YY, Boynton AN, Spurr LF, Iorgulescu JB, Bohrson CL, et al. Mechanisms and therapeutic implications of hypermutation in gliomas. Nature. 2020;580(7804):517-23.

11. Larson JD, Kasper LH, Paugh BS, Jin H, Wu G, Kwon CH, et al. Histone H3.3 K27M Accelerates Spontaneous Brainstem Glioma and Drives Restricted Changes in Bivalent Gene Expression. Cancer Cell. 2019;35(1):140–55 e7.

12. Pathania M, De Jay N, Maestro N, Harutyunyan AS, Nitarska J, Pahlavan P, et al. H3.3(K27M) Cooperates with Trp53 Loss and PDGFRA Gain in Mouse Embryonic Neural Progenitor Cells to Induce Invasive High-Grade Gliomas. Cancer Cell. 2017;32(5):684–700 e9.

13. Lewis PW, Muller MM, Koletsky MS, Cordero F, Lin S, Banaszynski LA, et al. Inhibition of PRC2 activity by a gain-of-function H3 mutation found in pediatric glioblastoma. Science. 2013;340(6134):857-61.

14. Filbin MG, Tirosh I, Hovestadt V, Shaw ML, Escalante LE, Mathewson ND, et al. Developmental and oncogenic programs in H3K27M gliomas dissected by single-cell RNA-seq. Science. 2018;360(6386):331-5.

15. Li XY, Das I, Lepletier A, Addala V, Bald T, Stannard K, et al. CD155 loss enhances tumor suppression via combined host and tumor-intrinsic mechanisms. J Clin Invest. 2018;128(6):2613–25.

16. Alvarez MJ, Shen Y, Giorgi FM, Lachmann A, Ding BB, Ye BH, et al. Functional characterization of somatic mutations in cancer using network-based inference of protein activity. Nat Genet. 2016;48(8):838–47.

17. Fernández EC, Tomassoni L, Zhang X, Wang J, Obradovic A, Laise P, et al. Elucidation and Pharmacologic Targeting of Master Regulator Dependencies in Coexisting Diffuse Midline Glioma Subpopulations. bioRxiv. 2024:2024.03.17.585370.

18. Califano A, and Alvarez MJ. The recurrent architecture of tumour initiation, progression and drug sensitivity. Nat Rev Cancer. 2017;17(2):116–30.

19. Hegde NS, Sanders DA, Rodriguez R, and Balasubramanian S. The transcription factor FOXM1 is a cellular target of the natural product thiostrepton. Nat Chem. 2011;3(9):725–31.

20. Kramer A, Green J, Pollard J, Jr., and Tugendreich S. Causal analysis approaches in Ingenuity Pathway Analysis. Bioinformatics. 2014;30(4):523–30.

21. O’Donnell JS, Madore J, Li XY, and Smyth MJ. Tumor intrinsic and extrinsic immune functions of CD155. Semin Cancer Biol. 2020;65:189–96.

22. Nakamura-Shinya Y, Iguchi-Manaka A, Murata R, Sato K, Vo AV, Kanemaru K, et al. DNAM-1 promotes inflammation-driven tumor development via enhancing IFN-gamma production. Int Immunol. 2022;34(3):149–57.

23. Lu C, Klement JD, Ibrahim ML, Xiao W, Redd PS, Nayak-Kapoor A, et al. Type I interferon suppresses tumor growth through activating the STAT3-granzyme B pathway in tumor-infiltrating cytotoxic T lymphocytes. J Immunother Cancer. 2019;7(1):157.

24. Tau GZ, Cowan SN, Weisburg J, Braunstein NS, and Rothman PB. Regulation of IFN-gamma signaling is essential for the cytotoxic activity of CD8(+) T cells. J Immunol. 2001;167(10):5574–82.

25. Olver S, Groves P, Buttigieg K, Morris ES, Janas ML, Kelso A, et al. Tumor-derived interleukin-4 reduces tumor clearance and deviates the cytokine and granzyme profile of tumor-induced CD8+ T cells. Cancer Res. 2006;66(1):571–80.

26. Pulkka OP, Mpindi JP, Tynninen O, Nilsson B, Kallioniemi O, Sihto H, et al. Clinical relevance of integrin alpha 4 in gastrointestinal stromal tumours. J Cell Mol Med. 2018;22(4):2220–30.

27. Cobb DA, de Rossi J, Liu L, An E, and Lee DW. Targeting of the alpha(v) beta(3) integrin complex by CAR-T cells leads to rapid regression of diffuse intrinsic pontine glioma and glioblastoma. J Immunother Cancer. 2022;10(2).

28. Barcelo J, Samain R, and Sanz-Moreno V. Preclinical to clinical utility of ROCK inhibitors in cancer. Trends Cancer. 2023;9(3):250–63.

29. Zhang JG, Zhang DD, Liu Y, Hu JN, Zhang X, Li L, et al. RhoC/ROCK2 promotes vasculogenic mimicry formation primarily through ERK/MMPs in hepatocellular carcinoma. Biochim Biophys Acta Mol Basis Dis. 2019;1865(6):1113–25.

30. Zhang X, Li T, Yang M, Du Q, Wang R, Fu B, et al. Acquired temozolomide resistance in MGMT(low) gliomas is associated with regulation of homologous recombination repair by ROCK2. Cell Death Dis. 2022;13(2):138.

31. Katzenellenbogen BS, Guillen VS, and Katzenellenbogen JA. Targeting the oncogenic transcription factor FOXM1 to improve outcomes in all subtypes of breast cancer. Breast Cancer Res. 2023;25(1):76.

32. Pandit B, and Gartel AL. Thiazole antibiotic thiostrepton synergize with bortezomib to induce apoptosis in cancer cells. PLoS One. 2011;6(2):e17110.

33. Wang JY, Jia XH, Xing HY, Li YJ, Fan WW, Li N, et al. Inhibition of Forkhead box protein M1 by thiostrepton increases chemosensitivity to doxorubicin in T-cell acute lymphoblastic leukemia. Mol Med Rep. 2015;12(1):1457–64.

34. Lin J, Zheng Y, Chen K, Huang Z, Wu X, and Zhang N. Inhibition of FOXM1 by thiostrepton sensitizes medulloblastoma to the effects of chemotherapy. Oncol Rep. 2013;30(4):1739–44.

35. Zhang Q, Yang L, Liu YH, Wilkinson JE, and Krainer AR. Antisense oligonucleotide therapy for H3.3K27M diffuse midline glioma. Sci Transl Med. 2023;15(691):eadd8280.

36. Monje M, Mahdi J, Majzner R, Yeom KW, Schultz LM, Richards RM, et al. Intravenous and intracranial GD2-CAR T cells for H3K27M(+) diffuse midline gliomas. Nature. 2024.

37. Vitanza NA, Wilson AL, Huang W, Seidel K, Brown C, Gustafson JA, et al. Intraventricular B7-H3 CAR T Cells for Diffuse Intrinsic Pontine Glioma: Preliminary First-in-Human Bioactivity and Safety. Cancer Discov. 2023;13(1):114–31.

38. Liu I, Jiang L, Samuelsson ER, Marco Salas S, Beck A, Hack OA, et al. The landscape of tumor cell states and spatial organization in H3-K27M mutant diffuse midline glioma across age and location. Nat Genet. 2022;54(12):1881–94.

39. Mortazavi A, Williams BA, McCue K, Schaeffer L, and Wold B. Mapping and quantifying mammalian transcriptomes by RNA-Seq. Nat Methods. 2008;5(7):621–8.

40. Love MI, Huber W, and Anders S. Moderated estimation of fold change and dispersion for RNA-seq data with DESeq2. Genome Biol. 2014;15(12):550.

