## Supplementary Material for "CD155 regulates tumor growth and immune evasion in diffuse midline glioma"

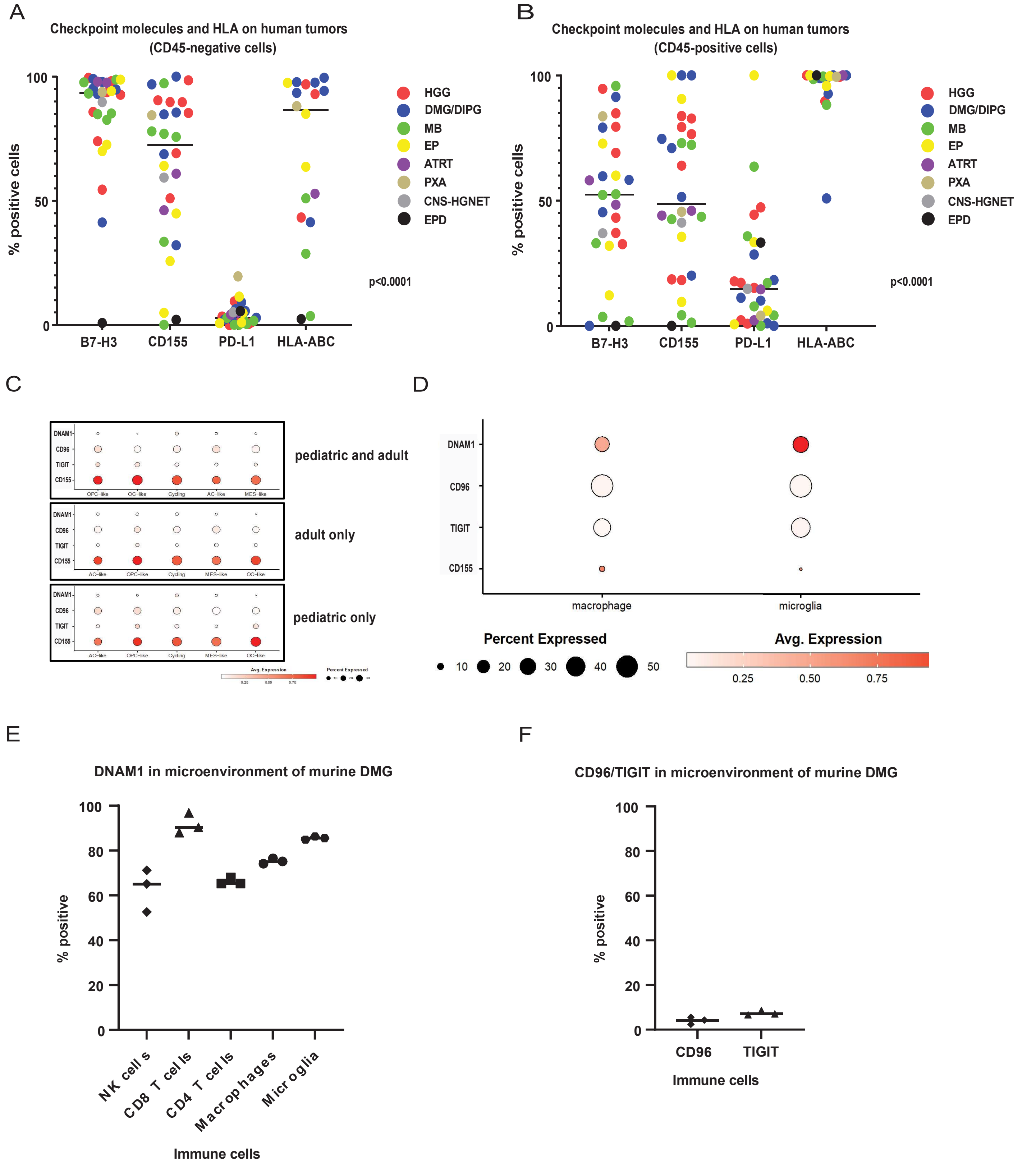

**Supplementary Figure 1. CD155 is expressed in human and murine DMG and in the microenvironment of these tumors.** A-B: Expression of selected checkpoint molecules and HLA-ABC in primary human brain tumors (1-way ANOVA test; A: CD45-negative, B: CD45-positive cells); C: scRNA sequencing data of human pediatric and adult DMG tumors showing expression levels of CD155, TIGIT, CD96 and DNAM1; D: scRNA sequencing data showing expression levels of CD155, TIGIT, CD96 and DNAM1 in the microenvironment of DMG tumors (macrophages and microglia); E: Expression of DNAM1 on the surface of immune cells in murine DMG tumors (KAPP); F: Expression of TIGIT and CD96 on the surface of CD45-positive cells.

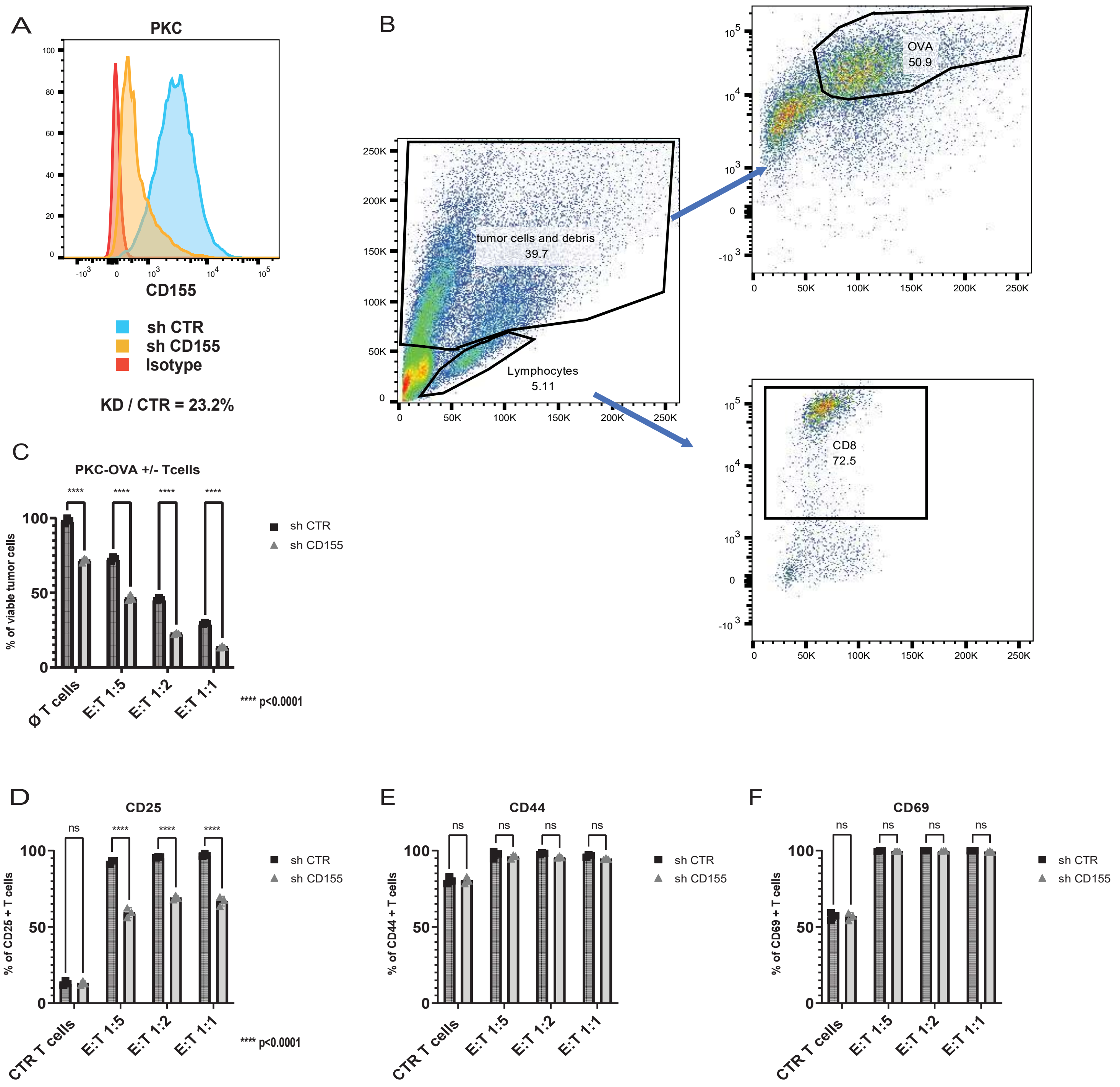

**Supplementary Figure 2. Loss of CD155 increases DMG cell sensitivity to T cell killing.** A: CD155 levels on the surface of CD155-silenced PKC cells (knockdown via shRNA1) versus control PKC cells; B: gating strategy of flow cytometric analysis of DMG-OVA and CD8 T cells; C: Percentage of viable PKC DMG tumor cells after co-culture with OT-I CD8 T cells in different Effector (T cells) to Target (Tumor) cells (E:T) and also without T cells (Ø T cells; 2-way ANOVA test comparing sh CTR to sh CD155); D-F: Percentage of CD25 (D), CD44 (E) and CD69 (F) positive CD8 T cells after co-culture with PKC-OVA cells. Control T cells not co-cultured with tumor cells are labelled as CTR T cells (2-way ANOVA test comparing sh CTR to sh CD155).

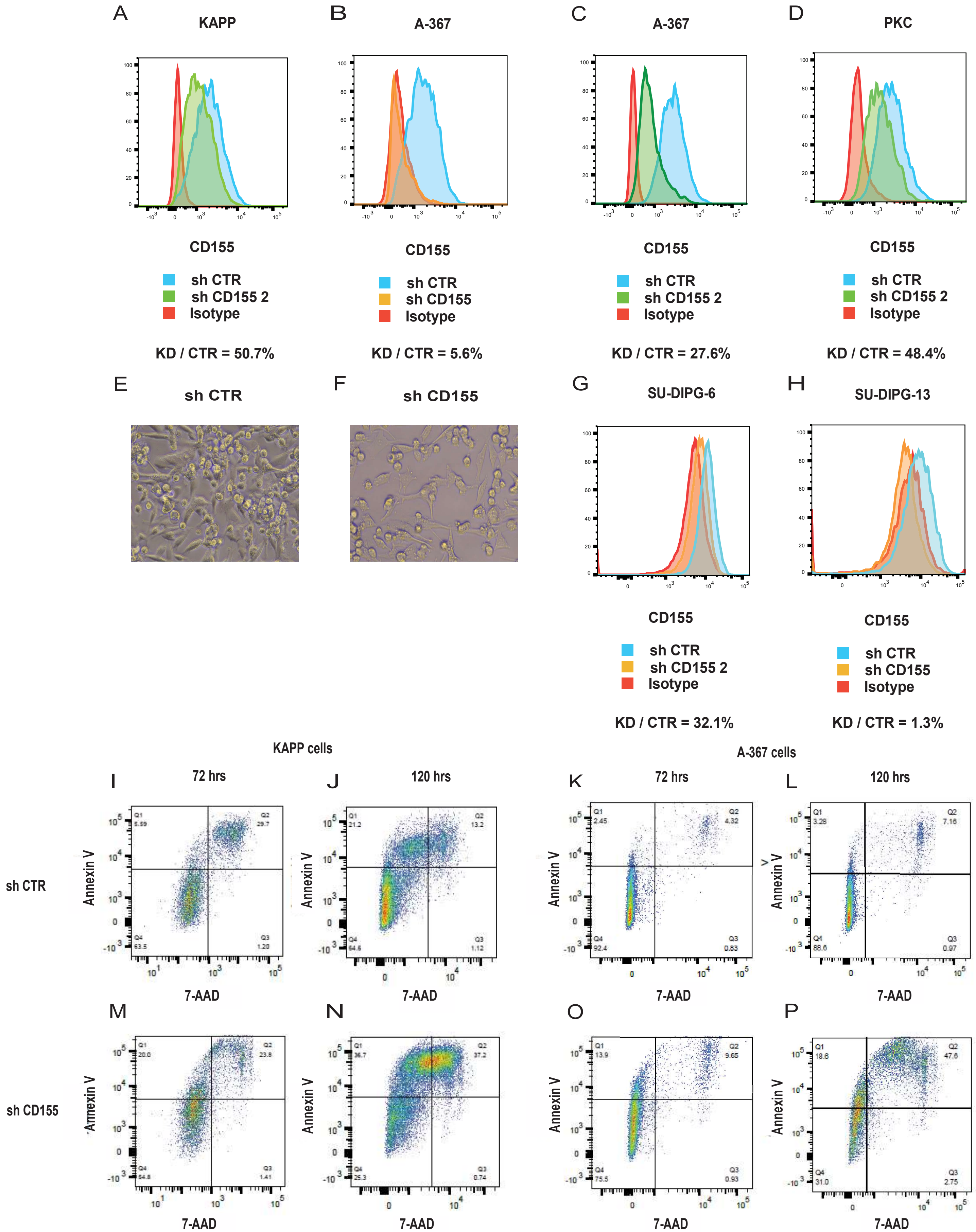

**Supplementary Figure 3. CD155 regulates DMG cell survival and tumor growth even in the absence of T cells.** A-D: CD155 levels on the surface of murine CD155-silenced versus control DMG cells (A: shRNA2 in KAPP, B: shRNA1 in A-367, C: shRNA2 in A-367 and D: shRNA2 in PKC cells); E-F: microscope pictures of control versus CD155-silenced KAPP cells; G-H: CD155 levels on the surface of human CD155-silenced versus control DMG cells (G: SU-DIPG-6, H: SU-DIPG-13); I-P: apoptotic assay with Annexin V and 7-AAD (I,J,M,N: KAPP, K,L,O,P: A-367) treated with control shRNA (sh CTR) or shRNA against CD155 (sh CD155) for 72 hours (72hrs) and after Puromycin selection for 48 hours (120 hrs), (2-way ANOVA test).

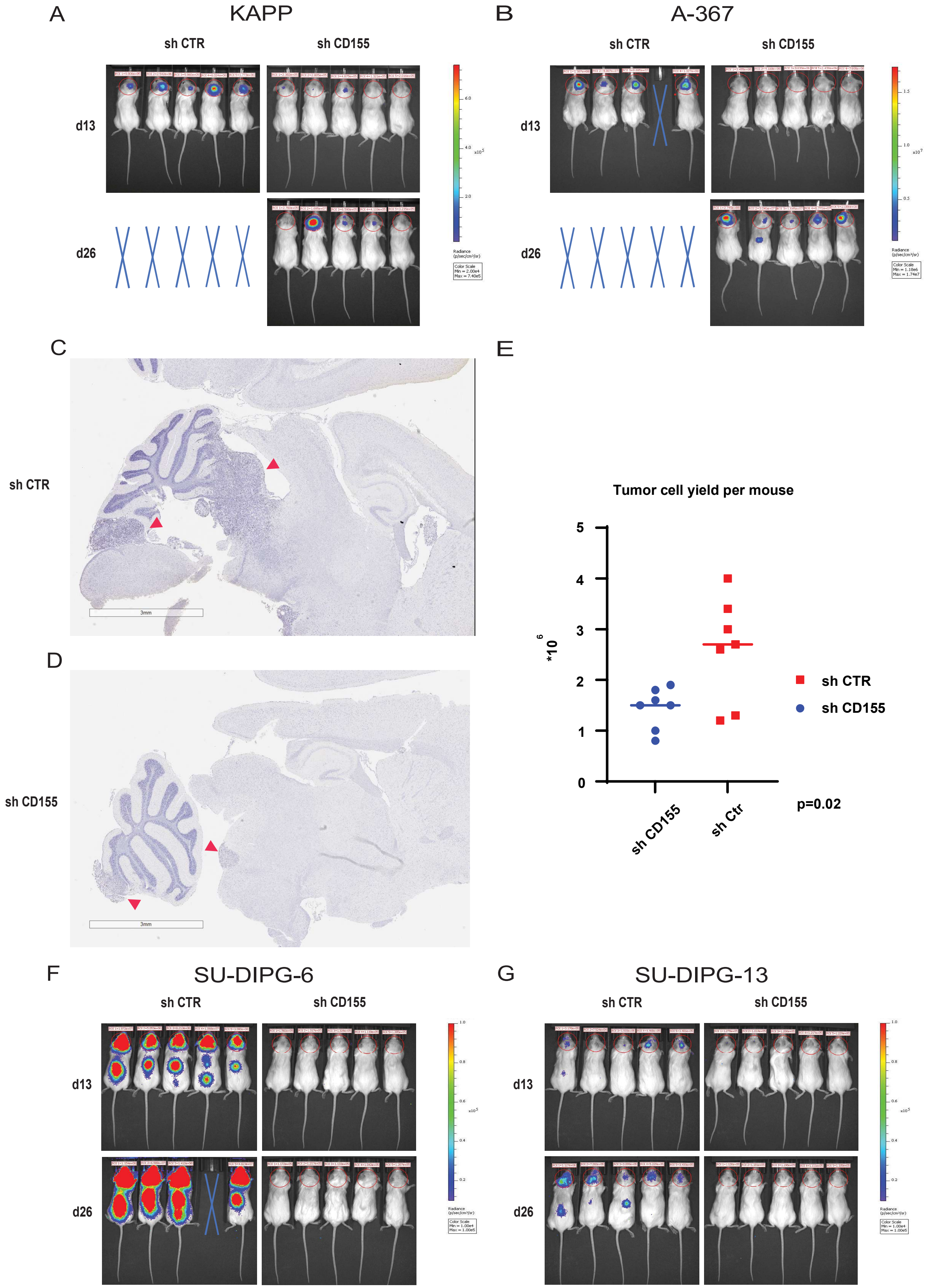

**Supplementary Figure 4. Loss of CD155 leads to delayed tumor growth and prolonged survival in immunodeficient mice.** A-B: representative IVIS images of KAPP (A) and A-367 (B) bearing NSG mice at d13 and d26 after transplant. C,D: representative H&E stained images of tumor sections from NSG mice with control DMG (sh CTR, C) or CD155-deficient (sh CD155, D) cells. Red arrows indicate the tumors. E: tumor cell yield of tumors from A-367 bearing NSG mice at endpoint of the experiment, as measured by Trypan blue staining (one-paired t test). F,G: representative IVIS images of SU-DIPG-6 (F) and SU-DIPG-13 (G) bearing NSG mice at d13 and d26 after transplant.

| **Table 1: Patient characteristics** | |
|---|---|
| n | 29 |
| Age (years) | 8.5 (0-18) |
| Sex | |
| Female | 14 (48.3%) |
| Male | 14 (48.3%) |
| n.k. | 1 (2.6%) |
| Disease status | |
| Primary | 21 (72.4%) |
| Recurrent | 8 (27.6%) |
| Entity | |
| High-grade glioma | 7 (24.1%) |
| DMG/DIPG | 7 (24.1%) |
| Medulloblastoma | 6 (20.7%) |
| Ependymoma | 4 (14%) |
| Atypical teratoid rhabdoid tumour | 2 (7%) |
| Pleomorphic xanthoastrocytoma | 1 (3.5%) |
| CNS HGNET | 1 (3.5%) |
| Epidermoid tumour | 1 (3.5%) |

| | HLA + | HLA/CD45- | HLA/CD45+ | | PDL1 + | CD45- | CD45+ | | B7H3 + | CD45- | CD45+ | | CD155 + | CD45- | CD45+ |
|---|---|---|---|---|---|---|---|---|---|---|---|---|---|---|---|
| HGG1 | 96.8% | 93.0% | 99.5% | | 28.7% | 0.8% | 47.3% | | 72.8% | 54.5% | 84.9% | | 31.5% | 51.0% | 18.6% |
| HGG2 | 97.7% | 96.9% | 100.0% | | 3.4% | 3.4% | 44.4% | | 86.1% | 92.7% | 69.1% | | 86.9% | 89.7% | 79.4% |
| HGG3 | 47.9% | 43.3% | 89.7% | | 10.3% | 9.5% | 17.2% | | 81.1% | 85.8% | 37.1% | | 89.7% | 90.5% | 82.8% |
| HGG4 | 99.9% | | | | 14.6% | 10.0% | 15.2% | | 92.3% | 74.0% | 94.7% | | 82.1% | 69.2% | 83.8% |
| HGG5 | | | | | 0.6% | 0.5% | 0.9% | | 93.0% | 98.2% | 79.5% | | 86.3% | 89.8% | 76.6% |
| HGG6 | 92.80% | | | | 1.3% | 1.0% | 17.8% | | 92.9% | 93.7% | 32.6% | | 84.5% | 85.4% | 18.3% |
| HGG7 | 97.70% | | | | 0.1% | 0.0% | 2.3% | | 97.9% | 99.5% | 43.2% | | 97.6% | 98.5% | 64.0% |
| DMG1 | 95.90% | | | | 1.12% | | | | 92.80% | | | | 95.20% | | |
| DMG2 | 55.4% | 41.4% | 98.8% | | 4.5% | 2.3% | 11.2% | | 53.0% | 41.3% | 91.4% | | 35.8% | 32.1% | 100.0% |
| DMG3 | 79.7% | 97.6% | 50.9% | | 5.8% | 2.9% | 10.1% | | 77.8% | 97.9% | 45.4% | | 85.6% | 85.6% | 74.7% |
| DMG4 | 96.4% | 93.5% | 98.9% | | 3.1% | 5.6% | 1.0% | | 74.4% | 93.0% | 58.3% | | 68.9% | 68.8% | 100.0% |
| DMG5 | 97.8% | 97.8% | 100.0% | | 9.1% | 9.1% | 0.0% | | 95.2% | 95.2% | 0.0% | | 95.3% | 100.0% | 20.1% |
| DMG6 | 99.6% | 99.6% | 100.0% | | 9.8% | 6.3% | 28.5% | | 96.0% | 99.1% | 79.2% | | 93.1% | 96.9% | 71.0% |
| DMG7 | 94.3% | 94.3% | 92.7% | | 3.2% | 2.2% | 18.3% | | 92.2% | 94.8% | 59.8% | | 82.7% | 84.8% | 51.5% |
| MB1 | 51.8% | 51.0% | 88.3% | | 0.6% | 0.4% | 7.8% | | 96.1% | 97.7% | 52.6% | | 74.7% | 75.7% | 43.6% |
| MB2 | 32.6% | 28.7% | 99.6% | | 3.8% | 2.9% | 17.1% | | 80.7% | 82.6% | 52.3% | | 35.9% | 33.5% | 72.3% |
| MB3 | 6.5% | 3.6% | 98.6% | | 0.1% | 0.1% | 0.0% | | 80.0% | 85.2% | 1.8% | | 73.4% | 78.0% | 4.3% |
| MB4 | 25.1% | | | | 2.3% | 1.8% | 35.8% | | 70.7% | 84.9% | 3.6% | | 0.4% | 0.1% | 1.3% |
| MB5 | 14.0% | | | | 2.9% | 1.7% | 63.6% | | 98.8% | 98.8% | 95.8% | | 76.8% | 77.0% | 73.0% |
| MB6 | 12.70% | | | | 0.2% | 0.1% | 4.2% | | 92.5% | 93.2% | 33.0% | | 96.8% | 97.3% | 42.6% |
| EP1 | 66.6% | 63.7% | 100.0% | | 1.0% | 1.0% | 0.6% | | 66.2% | 70.0% | 32.0% | | 64.2% | 64.1% | 100.0% |
| EP2 | 98.7% | 97.5% | 99.7% | | 5.6% | 4.9% | 6.1% | | 78.1% | 98.8% | 60.0% | | 51.0% | 4.9% | 90.6% |
| EP3 | 85.7% | 85.0% | 95.8% | | 4.2% | 0.7% | 33.4% | | 72.6% | 72.6% | 72.8% | | 43.3% | 44.9% | 35.6% |
| EP4 | 98.5% | | | | 10.4% | 11.4% | 100.0% | | 70.7% | 94.2% | 12.2% | | 21.2% | 25.7% | 9.6% |
| ATRT1 | 69.9% | 52.9% | 100.0% | | 8.5% | 4.2% | 14.6% | | 75.5% | 98.0% | 48.4% | | 46.1% | 46.2% | 46.0% |
| ATRT2 | 86.8% | | | | 4.4% | 5.0% | 2.2% | | 88.3% | 97.6% | 58.1% | | 56.9% | 60.9% | 44.1% |
| PXA | 92.4% | 88.1% | 99.5% | | 12.7% | 19.6% | 4.1% | | 89.2% | 93.8% | 83.7% | | 66.9% | 84.5% | 45.6% |
| CNS HGNET | 96.3% | | | | 6.7% | 5.1% | 14.8% | | 81.0% | 89.8% | 37.0% | | 56.3% | 59.4% | 41.2% |
| EPD | 3.9% | 2.4% | 100.0% | | 6.3% | 5.6% | 33.3% | | 0.8% | 0.8% | 0.0% | | 2.1% | 2.1% | 0.0% |

| Upstream Regulators | A-367 | KAPP |
|---|---|---|
| Eldr | -7.111 | -3.464 |
| CKAP2L | -6.403 | -3.317 |
| ATF4 | -6.345 | -4.681 |
| RNASEH2B | -6.258 | -4.918 |
| FOXM1 | -6.056 | -3.419 |
| aflatoxin B1 | -5.824 | -3.101 |
| CEBPB | -5.656 | -2.464 |
| AREG | -5.481 | -3.913 |
| Vegf | -5.405 | -4.049 |
| TREX1 | -5.216 | -5.297 |
| Ttc39aos1 | -5.21 | -5.359 |
| beta-estradiol | -5.202 | -1.514 |
| STAG2 | -5.189 | -5.274 |
| MITF | -5.189 | -0.48 |
| RABL6 | -5.145 | -3.317 |
| CSF2 | -5.111 | -2.01 |
| SN-011 | -5.096 | -4.943 |
| Insulin | -5.078 | -2.039 |
| HGF | -5.004 | -3.397 |
| PTGER4 | -4.985 | -4.352 |
| TRIM24 | -4.863 | -5.357 |
| IRGM | -4.852 | -5.297 |
| 8-bromo-cAMP | -4.837 | -0.836 |
| nelfinavir | -4.829 | -3.185 |
| PTGER2 | -4.76 | -3.158 |
| ESR1 | -4.665 | -1.723 |
| carbamazepine | -4.634 | -3.421 |
| PNPT1 | -4.624 | -5.174 |
| NKX2-3 | -4.58 | -5.189 |
| IL4 | -4.532 | -3.491 |
| ESR2 | -4.505 | -1.605 |
| MAPK1 | -4.482 | -4.079 |
| ERBB2 | -4.31 | -2.464 |
| MED1 | -4.268 | -1.673 |
| ACKR2 | -4.243 | -4.379 |
| dihydrotestosterone | -4.218 | -1.931 |
| SREBF1 | -4.179 | -1.013 |
| AR | -4.107 | -1.681 |
| tosedostat | -4.094 | -3.286 |
| TBX2 | -4.025 | -1.183 |
| NFAT5 | -4.023 | -3.176 |
| CD24 | -3.983 | -3.014 |
| SREBF2 | -3.935 | -1.875 |
| MAPK7 | -3.93 | -0.244 |
| forskolin | -3.875 | -1.987 |
| UCP1 | -3.867 | -4.364 |
| CREB1 | -3.847 | -2.085 |
| LIN9 | -3.818 | -1.172 |
| CITED2 | -3.681 | -4.3 |
| RORA | -3.637 | -0.419 |
| YAP1 | -3.622 | -1.746 |
| H2AZ1 | -3.615 | -1.913 |
| USP8 | -3.605 | -4.5 |

| | | |
|---|---|---|
| SP1 | -3.604 | -2.17 |
| estrogen | -3.592 | -1.887 |
| MIF | -3.53 | -2.967 |
| BRD4 | -3.505 | -1.153 |
| PGR | -3.503 | -3.041 |
| E2F1 | -3.488 | -1.359 |
| cephaloridine | -3.487 | -2.354 |
| clopidogrel | -3.477 | -3.035 |
| diclofenac | -3.471 | -2.601 |
| EIF2AK3 | -3.465 | -2.549 |
| thapsigargin | -3.463 | -3.561 |
| MYC | -3.459 | -2.097 |
| IGF2 | -3.457 | -1.516 |
| dimethyl itaconate | -3.422 | -2.138 |
| tunicamycin | -3.415 | -5.105 |
| CCND1 | -3.387 | -2.519 |
| Irgm1 | -3.364 | -5.224 |
| EWSR1 | -3.348 | -0.555 |
| diethylstilbestrol | -3.331 | -2.257 |
| Z36 | -3.302 | -2.63 |
| ERK1/2 | -3.29 | -3.433 |
| KDM1A | -3.279 | -0.755 |
| Mek | -3.276 | -2.686 |
| NGF | -3.262 | -2.653 |
| STAT6 | -3.261 | -4.151 |
| SYVN1 | -3.239 | -1.5 |
| PIK3CG | -3.222 | -4.663 |
| THEM6 | -3.183 | -2.704 |
| EIF2AK4 | -3.183 | -1.07 |
| PI3K (complex) | -3.181 | -1.073 |
| medroxyprogesterone acetate | -3.181 | 0.284 |
| PAX7 | -3.179 | -2.219 |
| CSF1 | -3.172 | -1.36 |
| EGFR | -3.135 | -1.846 |
| PG3-Oc | -3.13 | -2.39 |
| metribolone | -3.126 | -2.472 |
| EGR1 | -3.118 | -2.835 |
| EGF | -3.118 | -2.15 |
| CG | -3.116 | -1.828 |
| RACK1 | -3.111 | -1.709 |
| RARA | -3.097 | -1.758 |
| TGFB1 | -3.069 | -0.812 |
| NTRK2 | -3.066 | -1.528 |
| Lh | -3.058 | -2.715 |
| PIN1 | -3.037 | -2.749 |
| FSH | -3.036 | -2.515 |
| AKT1 | -3.035 | -1.887 |
| ASPSCR1-TFE3 | -3.02 | -0.839 |
| IOX2 | -3 | -1.342 |
| VH032 | -3 | -1.342 |
| E2F3 | -2.998 | -2.007 |
| TGFB3 | -2.982 | -1.668 |
| NCOR1 | -2.978 | -2.413 |
| DUSP11 | -2.976 | -3.719 |

| | | |
|---|---|---|
| KDM3A | -2.966 | -1.8 |
| Akt | -2.965 | -2.159 |
| JUN | -2.965 | -1.732 |
| NCOA3 | -2.898 | -1.738 |
| ATF6 | -2.895 | -2.299 |
| USP18 | -2.894 | -2.765 |
| lithium chloride | -2.864 | -0.779 |
| PRKD1 | -2.848 | -1.367 |
| BCR (complex) | -2.834 | -3.113 |
| CNB-001 | -2.828 | -2.449 |
| CD276 | -2.828 | -2.132 |
| CASR | -2.826 | -1.144 |
| SOCS1 | -2.82 | -3.279 |
| IL1A | -2.818 | -0.681 |
| Mapk | -2.808 | -1.527 |
| F2 | -2.793 | -1.535 |
| urethane | -2.792 | -0.587 |
| NFKB1 | -2.783 | -2.019 |
| Ap1 | -2.77 | -1.897 |
| NGLY1 | -2.769 | -2.632 |
| miR-182-5p (and other miRNAs w/seed UUGGCAA) | -2.76 | -2.853 |
| Z-LLL-CHO | -2.743 | -2.213 |
| KMT2D | -2.739 | -2.838 |
| VDR | -2.728 | -2.097 |
| IGF1 | -2.728 | -1.105 |
| NQO1 | -2.727 | -2.789 |
| SP3 | -2.726 | -2.364 |
| TREM1 | -2.703 | -3.332 |
| PRDM16 | -2.693 | -2.22 |
| Jnk | -2.686 | -2.483 |
| POU2F2 | -2.683 | -0.143 |
| methyl methanesulfonate | -2.663 | -2.432 |
| EIF4G1 | -2.646 | -1.342 |
| Pdgf (complex) | -2.636 | -2.022 |
| C5 | -2.632 | -2.004 |
| PRKCE | -2.624 | -2.202 |
| TAB1 | -2.621 | -1.715 |
| MYB | -2.615 | -1.597 |
| LG100268 | -2.608 | -0.816 |
| Hif1 | -2.601 | -1.98 |
| EIF4E | -2.599 | -0.354 |
| IL18 | -2.59 | -1.844 |
| RC3H1 | -2.584 | -3.051 |
| DNASE2 | -2.576 | -2.375 |
| CIRBP | -2.562 | -1.57 |
| IL1RN | -2.561 | -2.168 |
| ethanol | -2.561 | -2.151 |
| prostaglandin D2 | -2.561 | -1.131 |
| 48s | -2.556 | -1.109 |
| FN1 | -2.55 | -0.92 |
| KRAS | -2.549 | -1.612 |
| NLRX1 | -2.534 | -1.889 |
| MMP9 | -2.534 | -0.826 |
| MBD2 | -2.531 | -2.425 |

| | | |
|---|---|---|
| crocidolite asbestos | -2.53 | -1.987 |
| ASXL2 | -2.53 | -0.447 |
| memantine | -2.529 | -0.561 |
| IL1 | -2.527 | -2.244 |
| MAP2K1 | -2.514 | -0.676 |
| FGF2 | -2.508 | -2.441 |
| RAC1 | -2.506 | -2.044 |
| LIN28B | -2.503 | -2.714 |
| prostaglandin E2 | -2.503 | -2.109 |
| ELL2 | -2.496 | -2.121 |
| 1,4-bis[2-(3,5-dichloropyridyloxy)]benzene | -2.493 | -1.707 |
| ETV7 | -2.478 | -1.76 |
| MET | -2.46 | -1.838 |
| 2-deoxyglucose | -2.458 | -2.057 |
| GPAT4 | -2.449 | -2 |
| NOTCH3 | -2.444 | -1.152 |
| NRG1 | -2.437 | -1.396 |
| arsenite | -2.435 | -3.071 |
| 1-palmitoyl-2-oleoylphosphatidylserine | -2.433 | -1.195 |
| RPTOR | -2.42 | -0.91 |
| ouabain | -2.414 | -1.067 |
| filgrastim | -2.409 | -2.86 |
| adenosine triphosphate | -2.407 | -1.775 |
| YBX1 | -2.404 | -2.543 |
| IDO1 | -2.39 | -2.449 |
| mibolerone | -2.387 | -0.57 |
| PDGF BB | -2.383 | -1.31 |
| ERN1 | -2.375 | -3.486 |
| Pam3-Cys | -2.375 | -1.069 |
| CREB3 | -2.373 | -0.849 |
| ZC3H12C | -2.365 | -3.808 |
| bucladesine | -2.364 | -1.444 |
| TCF/LEF | -2.357 | -2.158 |
| 26s Proteasome | -2.351 | -0.858 |
| TARDBP | -2.348 | 0.323 |
| acetaminophen | -2.342 | -0.953 |
| CTNNB1 | -2.339 | -0.038 |
| vitamin K3 | -2.33 | -0.296 |
| PI3K (family) | -2.33 | 0.301 |
| VEGFA | -2.329 | -1.144 |
| KLF6 | -2.317 | -1.047 |
| ATF2 | -2.314 | -1.802 |
| CCAR2 | -2.309 | -1.89 |
| PLAT | -2.306 | -0.883 |
| kainic acid | -2.283 | -1.742 |
| RAS | -2.282 | -0.497 |
| HIF1A | -2.274 | -0.31 |
| NFkB (complex) | -2.273 | -0.212 |
| L-glutamic acid | -2.268 | -1.859 |
| TREM2 | -2.264 | -1.484 |
| diethylnitrosamine | -2.262 | -1.106 |
| azoxymethane | -2.262 | -0.4 |
| SMAD3 | -2.26 | -2.202 |
| SPZ1 | -2.244 | -2.176 |

| | | |
|---|---|---|
| TP63 | -2.244 | -2.124 |
| MASTL | -2.243 | -1.572 |
| ST3-Hel2A-2 | -2.236 | -2 |
| 6-aminopyrazolopyrimidine derivative compound II | -2.236 | -2 |
| NMU | -2.236 | -1.342 |
| TENM1 | -2.236 | -1 |
| NUP62 | -2.231 | -1.032 |
| SNHG11 | -2.224 | -0.927 |
| dithiothreitol | -2.222 | -1.4 |
| S1PR3 | -2.219 | -1.969 |
| USP37 | -2.219 | -1.414 |
| CCL2 | -2.216 | -2.1 |
| Tnf (family) | -2.214 | -1.167 |
| Cd2+ | -2.213 | -1.982 |
| NADPH oxidase | -2.213 | -0.555 |
| CLPP | -2.211 | -2.597 |
| PTK2 | -2.203 | -2.181 |
| ERK | -2.199 | -1.063 |
| NOX4 | -2.197 | -0.004 |
| histamine | -2.188 | -2.343 |
| COPS5 | -2.187 | -2.414 |
| HNF1A-AS1 | -2.184 | -1.213 |
| KMT2A | -2.172 | -2.376 |
| PIK3R1 | -2.158 | -0.467 |
| SLC2A3 | -2.157 | -1.98 |
| MMP2 | -2.155 | -0.329 |
| MAP3K1 | -2.152 | -2.942 |
| PGF | -2.15 | -2.169 |
| PTPN11 | -2.149 | -1.847 |
| RPSA | -2.143 | -2.481 |
| MRTFA | -2.131 | -1.519 |
| fluticasone propionate | -2.127 | -2.158 |
| 10E,12Z-octadecadienoic acid | -2.116 | -1.244 |
| CDK9 | -2.105 | -2.213 |
| valproic acid | -2.092 | -2.345 |
| PKM | -2.081 | -0.607 |
| Pka | -2.078 | -2.023 |
| GNRH | -2.074 | -1.412 |
| D-glucose | -2.073 | -0.883 |
| Gsk3 | -2.068 | -1.89 |
| STAR | -2.065 | -2.333 |
| G protein alpha | -2.065 | -1.265 |
| H1f1 | -2.065 | -1.265 |
| H1-6 | -2.065 | -1.265 |
| GATA6 | -2.061 | -1.775 |
| TCF7L2 | -2.06 | 0.465 |
| Ngf | -2.057 | -1.121 |
| Creb | -2.053 | -1.308 |
| GPER1 | -2.051 | -0.735 |
| MTORC1 | -2.05 | -0.142 |
| MMP1 | -2.048 | -2.26 |
| SOX4 | -2.047 | -1.705 |
| bicuculline | -2.044 | -1.44 |
| NFkB (family) | -2.039 | -0.677 |

| | | |
|---|---|---|
| cytokine | -2.037 | -1.252 |
| pilocarpine | -2.027 | -0.955 |
| NFE2L2 | -2.02 | -2.914 |
| CTCF | -2.014 | -1.048 |
| Ca2+ | -2.014 | -1.004 |
| ionomycin | -2.013 | -1.165 |
| RU.521 | -2 | -2.236 |
| H-151 | -2 | -2.158 |
| 1,2-dimethylhydrazine | -2 | -2.03 |
| ceruletide | -2 | -1.154 |
| pidnarulex | -2 | -0.711 |
| FURIN | -2 | -0.351 |
| MAP3K14 | -1.999 | -0.869 |
| KDM4D | -1.998 | -0.816 |
| MTOR | -1.998 | -0.548 |
| XBP1 | -1.991 | -3.485 |
| CpG oligonucleotide | -1.991 | -1.026 |
| BCR-ABL1 | -1.984 | -1.544 |
| glycochenodeoxycholate | -1.982 | -1.648 |
| caspase | -1.982 | -1.387 |
| INHBB | -1.982 | -0.555 |
| Ck2 alpha | -1.977 | -1.996 |
| IL-1R | -1.976 | -2.828 |
| IGFBP5 | -1.969 | -1.703 |
| premarin | -1.964 | -1.732 |
| PSMD10 | -1.964 | -1.467 |
| TGFA | -1.953 | -1.403 |
| histidinol | -1.951 | -2 |
| TRPC1 | -1.951 | -0.785 |
| cytarabine | -1.947 | -1.961 |
| M344 | -1.941 | -2.433 |
| sodium orthovanadate | -1.941 | -1.756 |
| TXN | -1.939 | -0.321 |
| RELB | -1.929 | -0.696 |
| 14,15-epoxyeicosatrienoic acid | -1.914 | -2 |
| 11,12-epoxyeicosatrienoic acid | -1.913 | -1 |
| SUB1 | -1.91 | -1.342 |
| MEX3A | -1.897 | -2 |
| FOXO1 | -1.895 | -1.421 |
| SLC15A4 | -1.894 | -0.21 |
| NAE1 | -1.89 | -1.732 |
| LINC00963 | -1.89 | -1 |
| F7 | -1.879 | -1.854 |
| GPR174 | -1.877 | -1.388 |
| NPM1 | -1.875 | -0.162 |
| STAT4 | -1.857 | 0.792 |
| CREM | -1.856 | -1.095 |
| IL2 | -1.849 | -1.71 |
| phenylephrine | -1.845 | -0.349 |
| bleomycin | -1.842 | -1.468 |
| calcimycin | -1.841 | -1.349 |
| FSHB | -1.838 | -0.625 |
| chenodeoxycholic acid | -1.834 | -1.704 |
| PDX1 | -1.822 | -1.235 |

| | | |
|---|---|---|
| MAP2K1/2 | -1.819 | -2.06 |
| norepinephrine | -1.813 | -0.81 |
| IGFBP2 | -1.809 | -0.862 |
| PITX2 | -1.806 | -1.535 |
| TBK1 | -1.801 | -1.199 |
| cigarette smoke | -1.801 | -0.024 |
| NFKB2 | -1.796 | -2.054 |
| NRAS | -1.788 | -3.357 |
| PRKCI | -1.786 | -1.265 |
| MAPK8 | -1.785 | -0.799 |
| IRF8 | -1.781 | -1.817 |
| WNT3A | -1.78 | -0.837 |
| IL10 | -1.776 | -1.668 |
| NR4A3 | -1.772 | 0.976 |
| PRKCB | -1.768 | -1.785 |
| CDK1 | -1.767 | -1.754 |
| NTRK1 | -1.765 | -0.82 |
| FABP5 | -1.764 | -2.449 |
| NfkB1-RelA | -1.761 | -1.555 |
| STAT3 | -1.76 | -2.379 |
| reactive oxygen species | -1.76 | -2.094 |
| ATF3 | -1.756 | -1.979 |
| okadaic acid | -1.75 | -1.765 |
| cobalt chloride | -1.75 | -0.334 |
| lactacystin | -1.743 | -1.821 |
| HRAS | -1.741 | -1.965 |
| CAMKK2 | -1.741 | -0.218 |
| PPP3R1 | -1.739 | -0.711 |
| deferoxamine | -1.732 | -0.993 |
| LLGL2 | -1.732 | -0.816 |
| JUNB | -1.727 | -0.56 |
| PTGES | -1.726 | -0.729 |
| sulindac sulfide | -1.721 | -1.587 |
| CTSB | -1.717 | -1.71 |
| SIRT1 | -1.715 | -3.187 |
| RGS4 | -1.706 | -1.414 |
| FLT1 | -1.701 | -1.89 |
| CCN2 | -1.697 | -0.447 |
| amphetamine | -1.696 | -2.58 |
| tributyrin | -1.691 | -2.554 |
| ZNF507 | -1.686 | -1.036 |
| cisplatin | -1.669 | -0.613 |
| BCL2L1 | -1.664 | -0.745 |
| TNFSF11 | -1.646 | -0.646 |
| SRC | -1.643 | -2.108 |
| mir-146 | -1.638 | -0.565 |
| ERG | -1.621 | -2.169 |
| lysophosphatidylcholine | -1.618 | -0.985 |
| TFEB | -1.606 | -2.204 |
| F2R | -1.606 | -1.771 |
| NCD-38 | -1.605 | -1.091 |
| CLIC4 | -1.604 | -2.345 |
| WNT1 | -1.596 | -0.729 |
| gentamicin | -1.595 | -1.616 |

| | | |
|---|---|---|
| CYP1B1 | -1.594 | -0.636 |
| mir-223 | -1.588 | -1.116 |
| SMARCA5 | -1.586 | -0.879 |
| GATA4 | -1.581 | -0.209 |
| CCL5 | -1.58 | -0.864 |
| estrogen receptor | -1.577 | -0.474 |
| DETA-NONOate | -1.573 | -1.637 |
| 6-hydroxydopamine | -1.572 | -1.172 |
| CAV1 | -1.546 | -1.207 |
| PRRX1 | -1.545 | -1.906 |
| homocysteine | -1.544 | -2.378 |
| IL10RA | -1.539 | -3.254 |
| nitric oxide | -1.536 | -1.961 |
| WWTR1 | -1.532 | -0.958 |
| sphingosine-1-phosphate | -1.529 | -1.239 |
| deoxynivalenol | -1.529 | -1.067 |
| trabectedin | -1.526 | 0.447 |
| NOTCH1 | -1.511 | -1.953 |
| FOS | -1.51 | -0.888 |
| GATA3 | -1.508 | -0.86 |
| CEBPA | -1.495 | -2.371 |
| Il3 | -1.488 | -0.835 |
| cadmium chloride | -1.486 | -0.46 |
| HBEGF | -1.472 | -2.041 |
| MAP2K4 | -1.47 | -2.408 |
| CD40 | -1.467 | -1.675 |
| mitomycin C | -1.465 | -1.645 |
| erlotinib | -1.463 | -1.948 |
| ziritaxestat | -1.46 | -1.886 |
| (1S,2R)-NCL-1 | -1.46 | -0.882 |
| SOCS3 | -1.454 | -0.895 |
| EDN1 | -1.446 | -0.425 |
| hydroxyurea | -1.443 | -0.473 |
| ASAH1 | -1.44 | -2.216 |
| cadmium | -1.427 | -2.013 |
| bufalin | -1.414 | -0.509 |
| N-acetylsphingosine | -1.409 | -0.875 |
| NR1I3 | -1.407 | -1.732 |
| EP300 | -1.406 | -1.574 |
| NF-Y | -1.406 | -0.772 |
| SRF | -1.396 | -1.032 |
| cyclic AMP | -1.39 | -0.244 |
| PDGF-AA | -1.377 | -0.879 |
| CCK | -1.375 | -0.57 |
| isobutylmethylxanthine | -1.374 | -1.675 |
| hydrogen peroxide | -1.373 | -1.931 |
| TRIM21 | -1.372 | -2.332 |
| choline | -1.372 | -0.557 |
| KAT5 | -1.357 | -1.951 |
| BMP7 | -1.354 | -1.656 |
| ALB | -1.328 | -1.276 |
| Pam3-Cys-Ser-Lys4 | -1.324 | -1.206 |
| N-Ac-Leu-Leu-norleucinal | -1.321 | -0.335 |
| sodium arsenite | -1.317 | -1.518 |

| | | |
|---|---|---|
| PLCG2 | -1.315 | -2.273 |
| VASP | -1.309 | -1.953 |
| SW016789 | -1.309 | -1.131 |
| CNR1 | -1.301 | -0.35 |
| GNAS | -1.292 | -0.385 |
| TNF | -1.292 | -0.278 |
| SERPINF1 | -1.289 | -0.866 |
| quinolinic acid | -1.285 | -1.352 |
| EHF | -1.279 | -1.106 |
| SP4 | -1.276 | -1.067 |
| farnesol | -1.272 | -1.857 |
| TRIM14 | -1.266 | -1.127 |
| CISH | -1.265 | -0.333 |
| POSTN | -1.248 | -2 |
| hyaluronic acid | -1.246 | -1.044 |
| Collagen type IV | -1.238 | -2 |
| TAC1 | -1.234 | -1.164 |
| BDNF | -1.233 | -0.533 |
| CD38 | -1.231 | -0.083 |
| PTHLH | -1.23 | -2.208 |
| SLC13A1 | -1.225 | -1.387 |
| ISG15 | -1.213 | -0.761 |
| GAST | -1.203 | -1.654 |
| carbonyl cyanide m-chlorophenyl hydrazone | -1.196 | -2.147 |
| SMARCD3 | -1.195 | -0.555 |
| PTPN1 | -1.189 | -1.937 |
| CFI-402257 | -1.188 | -2.425 |
| JAK inhibitor I | -1.188 | -0.243 |
| lovastatin | -1.184 | -0.217 |
| FGFR3 | -1.177 | -2.425 |
| ECSIT | -1.177 | -1.584 |
| cuprizone | -1.172 | -1.666 |
| CARM1 | -1.172 | -1.176 |
| APBB1 | -1.172 | -1.067 |
| methylprednisolone | -1.17 | -1.075 |
| INSR | -1.157 | -0.102 |
| PAK2 | -1.155 | -1.134 |
| TET3 | -1.145 | -0.765 |
| PTGS2 | -1.143 | -1.899 |
| NQO2 | -1.134 | -2 |
| FZD7 | -1.134 | -2 |
| tert-butyl-hydroquinone | -1.132 | -1.348 |
| SHH | -1.122 | -1.65 |
| FASN | -1.116 | -0.788 |
| FOXL2 | -1.113 | -1.713 |
| prexasertib | -1.095 | -1.807 |
| N-[(2Z)-3-(4,5-dihydro-1,3-thiazol-2-yl)-1,3-thiazolidin-2-yl idene]-5H dibenzo[a,d][7]annulen-5-amine | -1.095 | -1.584 |
| heparin | -1.069 | -2.736 |
| METTL3 | -1.067 | -1.076 |
| ERBB4 | -1.065 | -0.917 |
| SOD2 | -1.062 | -0.973 |
| prednisolone | -1.058 | -0.346 |
| MYOD1 | -1.057 | -0.479 |
| UBE2I | -1.046 | -1.982 |

| | | |
|---|---|---|
| halofuginone | -1.042 | -0.468 |
| cholecalciferol | -1.035 | -0.028 |
| Ige | -1.026 | -2.474 |
| DIO3 | -1.015 | -2.04 |
| IRS1 | -1.011 | -1.88 |
| LIN28A | -0.997 | -0.396 |
| arsenic | -0.993 | -1.961 |
| SPRY2 | -0.99 | -0.77 |
| MECP2 | -0.987 | -0.534 |
| TAFAZZIN | -0.983 | -0.945 |
| STAT5a/b | -0.982 | -1.409 |
| CD3 | -0.978 | -0.458 |
| HOXA7 | -0.971 | -0.816 |
| tetradecanoylphorbol acetate | -0.964 | -2.675 |
| CREBBP | -0.962 | -0.615 |
| HOXA9 | -0.961 | -0.547 |
| NPC1 | -0.935 | -2.035 |
| NDP | -0.933 | -0.626 |
| FGFR1 | -0.932 | -0.326 |
| IL5 | -0.931 | -0.812 |
| DUSP1 | -0.919 | -1.448 |
| PPRC1 | -0.905 | -2.2 |
| resiquimod | -0.904 | -0.983 |
| NUDT21 | -0.896 | -0.152 |
| cannabidiol | -0.894 | -0.933 |
| ARID1A | -0.887 | -0.418 |
| CDK4/6 | -0.879 | -1.982 |
| BMP6 | -0.869 | -0.778 |
| FGF1 | -0.864 | -0.587 |
| DCAF1 | -0.862 | -1.841 |
| NRIP1 | -0.859 | -1.4 |
| MRTFB | -0.857 | -0.396 |
| palmitic acid | -0.855 | -1.138 |
| RARRES2 | -0.854 | -0.035 |
| 25-hydroxycholesterol | -0.85 | -1.708 |
| ketamine | -0.85 | -0.64 |
| PRKACA | -0.849 | -1.215 |
| CCN1 | -0.84 | -0.492 |
| REL | -0.838 | -0.07 |
| SKIL | -0.835 | -0.559 |
| CYP1A2 | -0.832 | -0.816 |
| deoxycholate | -0.83 | -2.012 |
| soloxolone methyl | -0.816 | -1.633 |
| BIX 01294 | -0.816 | -1 |
| ALDH1A2 | -0.807 | -0.928 |
| taurine | -0.804 | -1.6 |
| gemcitabine | -0.804 | -0.511 |
| RXRA | -0.801 | -0.668 |
| Growth hormone | -0.801 | -0.318 |
| SB 216763 | -0.8 | -0.359 |
| GNAQ | -0.785 | -0.158 |
| SP110 | -0.781 | -1.606 |
| MAOA | -0.781 | -1.211 |
| ZBED6 | -0.777 | -0.017 |

| | | |
|---|---|---|
| TCF20 | -0.775 | -0.333 |
| INS | -0.77 | -1.318 |
| S100A9 | -0.768 | -0.51 |
| farnesyl pyrophosphate | -0.765 | -2.2 |
| NEDD9 | -0.762 | -1.265 |
| mir-142 | -0.76 | -1.587 |
| 2-aminopurine | -0.76 | -1.443 |
| NR0B2 | -0.76 | -0.184 |
| amitriptyline | -0.74 | -0.555 |
| KLF4 | -0.728 | -0.55 |
| paclitaxel | -0.724 | -0.395 |
| IRF4 | -0.715 | -0.18 |
| CLU | -0.711 | -0.853 |
| lysophosphatidic acid | -0.708 | -1.465 |
| RHOA | -0.702 | -0.916 |
| STAT5A | -0.699 | -0.321 |
| ETV6-RUNX1 | -0.693 | -2.779 |
| zymosan | -0.692 | -1.012 |
| bafilomycin A1 | -0.686 | -1.278 |
| Pdgf Ab | -0.686 | -0.555 |
| ROR1 | -0.679 | -0.816 |
| TAS4464 | -0.677 | -1.504 |
| IL15 | -0.677 | -0.499 |
| EIF4EBP1 | -0.676 | -0.752 |
| clofibrate | -0.675 | -1.729 |
| EWSR1-FLI1 | -0.675 | -0.761 |
| SN-38 | -0.673 | -2.425 |
| FGFR4 | -0.669 | -1.633 |
| FOXC2 | -0.668 | -0.786 |
| PTGS1 | -0.661 | -0.464 |
| 9,10-dimethyl-1,2-benzanthracene | -0.66 | -0.829 |
| PRDM1 | -0.656 | -1.671 |
| captopril | -0.656 | -1.219 |
| carbon tetrachloride | -0.656 | -0.734 |
| ADAM17 | -0.653 | -0.737 |
| GSTO1 | -0.638 | -0.44 |
| abemaciclib | -0.628 | -0.298 |
| mir-155 | -0.622 | -1.727 |
| S100A8 | -0.622 | -0.398 |
| CXCR4 | -0.618 | -1.23 |
| TCR | -0.61 | -1.2 |
| IL17A | -0.608 | -0.451 |
| entolimod | -0.606 | -0.739 |
| DCLK1 | -0.605 | -2.2 |
| TEAD | -0.594 | -1.469 |
| KL | -0.594 | -0.681 |
| methylnitronitrosoguanidine | -0.588 | -1.845 |
| mir-23 | -0.588 | -0.314 |
| CD40LG | -0.574 | -2.058 |
| LMNA | -0.574 | -1.729 |
| testosterone | -0.567 | -1.164 |
| AIM2 | -0.566 | -0.214 |
| nicotine | -0.562 | -1.203 |
| zinc | -0.537 | -0.357 |

| | | |
|---|---|---|
| ZEB1 | -0.516 | -0.204 |
| CEBPD | -0.514 | -1.713 |
| VTN | -0.49 | -0.341 |
| HDAC2 | -0.485 | -0.055 |
| NR3C2 | -0.474 | -0.362 |
| PLAUR | -0.467 | -1.664 |
| SLC16A2 | -0.466 | -1.463 |
| EBF1 | -0.464 | -0.448 |
| fluvastatin | -0.463 | -0.092 |
| ERVW-1 | -0.461 | -1.195 |
| KAT2B | -0.457 | -0.885 |
| raloxifene | -0.456 | -0.752 |
| 1-methyl-4-phenyl-1,2,3,6-tetrahydropyridine | -0.452 | -0.578 |
| methamphetamine | -0.451 | -1.311 |
| SMAD4 | -0.449 | -0.088 |
| TGFBR | -0.447 | -1.067 |
| BRAP | -0.447 | -1 |
| RNF216 | -0.447 | -1 |
| CYP1A1 | -0.434 | -0.036 |
| PDGFC | -0.425 | -0.106 |
| progesterone | -0.421 | -1.497 |
| Collagen Alpha1 | -0.42 | -1.934 |
| SNAI2 | -0.42 | -0.052 |
| RAF1 | -0.405 | -2.032 |
| telmisartan | -0.404 | -0.152 |
| LRPAP1 | -0.402 | -0.478 |
| AMBRA1 | -0.399 | -1.508 |
| TSH | -0.396 | -0.743 |
| cinnamaldehyde | -0.393 | -0.499 |
| ciprofloxacin | -0.381 | -1.236 |
| brefeldin A | -0.38 | -1.081 |
| SLC2A1 | -0.373 | -0.152 |
| FGF19 | -0.371 | -2.213 |
| Sb202190 | -0.369 | -0.278 |
| dimethylnitrosamine | -0.368 | -0.933 |
| GLIS1 | -0.368 | -0.64 |
| bisphenol A | -0.363 | -0.248 |
| 5-azacytidine | -0.359 | -1.076 |
| SAFB | -0.359 | -0.905 |
| DDIT3 | -0.357 | -0.136 |
| AIRE | -0.355 | -1.861 |
| CLOCK | -0.355 | -0.728 |
| SIX1 | -0.351 | -0.462 |
| advanced glycation end-products | -0.346 | -1.697 |
| PPP1R15A | -0.346 | -0.283 |
| SALL4 | -0.343 | -0.626 |
| leukotriene D4 | -0.338 | -0.913 |
| C5AR1 | -0.335 | -1.117 |
| miR-335-3p (miRNAs w/seed UUUUCAU) | -0.333 | -0.6 |
| CL 316243 | -0.328 | -0.555 |
| PTPN2 | -0.326 | -0.489 |
| TNFRSF1A | -0.315 | -1.858 |
| SMURF2 | -0.314 | -0.655 |
| nitrofurantoin | -0.312 | -0.662 |

| | | |
|---|---|---|
| PLX5622 | -0.31 | -1.784 |
| bortezomib | -0.309 | -0.819 |
| TP73 | -0.306 | -0.227 |
| mycophenolic acid | -0.286 | -1 |
| APOE | -0.282 | -0.07 |
| Calmodulin | -0.277 | -0.711 |
| ERBB3 | -0.275 | -0.894 |
| GFI1 | -0.268 | -1.365 |
| mevalonic acid | -0.261 | -1.344 |
| HYAL1 | -0.258 | -2.236 |
| STAU1 | -0.243 | -0.152 |
| aldosterone | -0.242 | -0.292 |
| enterotoxin B | -0.237 | -0.458 |
| 3-deazaneplanocin | -0.227 | -0.198 |
| oxaliplatin | -0.219 | -0.693 |
| TRIM37 | -0.213 | -0.816 |
| salmonella minnesota R595 lipopolysaccharides | -0.208 | -0.01 |
| 2,4,5,2',4',5'-hexachlorobiphenyl | -0.2 | -0.258 |
| OGT | -0.196 | -0.124 |
| DNMT3B | -0.194 | -0.51 |
| linsitinib | -0.192 | 0.652 |
| sulforafan | -0.19 | -2.193 |
| GDF15 | -0.189 | -0.659 |
| tetrachlorodibenzodioxin | -0.189 | -0.499 |
| IGFBP7 | -0.186 | -0.6 |
| TCF4 | -0.179 | -1.536 |
| miR-199a-5p (and other miRNAs w/seed CCAGUGU) | -0.161 | -0.905 |
| TGFBR2 | -0.153 | -1.574 |
| indomethacin | -0.153 | -0.298 |
| CDKN1B | -0.152 | -1.382 |
| naproxen | -0.152 | -0.152 |
| GDF2 | -0.15 | -0.139 |
| FOXO4 | -0.144 | -0.714 |
| RHO | -0.137 | -1.387 |
| S1PR2 | -0.128 | -0.298 |
| SSTR2 | -0.128 | -0.29 |
| paraquat | -0.12 | -0.946 |
| OTUB1 | -0.119 | -0.415 |
| anisomycin | -0.111 | -0.359 |
| RUNX1 | -0.105 | -0.803 |
| RGFP966 | -0.103 | -2.734 |
| PIK3CA | -0.103 | -0.287 |
| Raf | -0.097 | -1.06 |
| Smad2/3-Smad4 | -0.089 | -0.232 |
| WT1 | -0.082 | -1.681 |
| FZD8 | -0.082 | -0.625 |
| thioacetamide | -0.073 | -0.757 |
| PRKCD | -0.07 | -1.479 |
| VHL | -0.066 | -0.509 |
| BMP2 | -0.061 | -0.027 |
| thyroid hormone | -0.058 | -0.674 |
| cyclosporin A | -0.049 | -0.733 |
| morphine | -0.046 | -1.381 |
| luteolin | -0.045 | -0.013 |

| | | |
|---|---|---|
| glutathione | -0.036 | -0.164 |
| streptozocin | -0.032 | -0.564 |
| aphidicolin | -0.025 | -0.608 |
| PPARA | -0.019 | -0.07 |
| STK11 | 0.004 | 0.099 |
| YY1 | 0.008 | 0.607 |
| IRF2 | 0.04 | 0.644 |
| EZH2 | 0.068 | 0.233 |
| E. coli serotype 0127B8 lipopolysaccharide | 0.077 | 0.232 |
| BTG2 | 0.077 | 0.42 |
| KNG1 | 0.082 | 0.119 |
| vancomycin | 0.085 | 0.791 |
| CLEC4G | 0.087 | 1.32 |
| enalapril | 0.091 | 0.514 |
| 1,1-bis(3'-indolyl)-1-(4-hydroxyphenyl)methane | 0.109 | 0.903 |
| WWTR1-CAMTA1 | 0.113 | 0.788 |
| PELP1 | 0.116 | 0.734 |
| CHD4 | 0.117 | 0.632 |
| ITK | 0.119 | 0.38 |
| IKK (complex) | 0.124 | 0.257 |
| NSD2 | 0.135 | 2 |
| TNFSF13B | 0.148 | 0.12 |
| fenretinide | 0.157 | 0.491 |
| COLQ | 0.161 | 0.239 |
| CYP19A1 | 0.171 | 0.011 |
| 5-N-ethylcarboxamido adenosine | 0.173 | 0.137 |
| dimethyl sulfoxide | 0.174 | 2.725 |
| steroid | 0.186 | 0.447 |
| GATA1 | 0.186 | 2.299 |
| THPO | 0.189 | 0.964 |
| iron | 0.206 | 0.261 |
| AGN194204 | 0.207 | 1.456 |
| 15(S)-HETE | 0.208 | 0.469 |
| BMP10 | 0.209 | 0.087 |
| KIF1B | 0.218 | 0.254 |
| IRF9 | 0.223 | 3.845 |
| SYK | 0.224 | 0.207 |
| TAF4 | 0.242 | 0.873 |
| SNAI1 | 0.244 | 1.336 |
| WBP2 | 0.253 | 0.832 |
| ENG | 0.259 | 0.914 |
| FGF21 | 0.262 | 1.772 |
| F2RL1 | 0.264 | 0.633 |
| 4-hydroxytamoxifen | 0.265 | 0.467 |
| cyclopamine | 0.266 | 0.758 |
| corticosterone | 0.271 | 2.143 |
| PSMB11 | 0.275 | 0.84 |
| HIVEP1 | 0.277 | 1 |
| KLF11 | 0.278 | 1.554 |
| PLAU | 0.289 | 0.484 |
| MYRF | 0.306 | 0.356 |
| trametinib | 0.308 | 0.444 |
| PROCR | 0.321 | 0.43 |
| bevacizumab | 0.333 | 1.664 |

| | | |
|---|---|---|
| INHA | 0.336 | 1.576 |
| levodopa | 0.346 | 0.009 |
| cholesterol | 0.348 | 0.353 |
| OGA | 0.348 | 1.027 |
| OSM | 0.356 | 0.614 |
| 1,25-dihydroxyvitamin D | 0.358 | 0.456 |
| FAS | 0.362 | 0.006 |
| TFE3 | 0.378 | 0.378 |
| levothyroxine | 0.379 | 0.625 |
| NR4A1 | 0.379 | 0.945 |
| MYCN | 0.381 | 0.477 |
| carbamylcholine | 0.387 | 0.368 |
| zoledronic acid | 0.389 | 0.478 |
| LAMC1 | 0.391 | 0.068 |
| PARP1 | 0.4 | 0.96 |
| CpG ODN 2216 | 0.4 | 1.172 |
| EIF2AK2 | 0.411 | 1.544 |
| daunorubicin | 0.414 | 0.503 |
| citarinostat | 0.426 | 0.896 |
| NCOA2 | 0.428 | 2.552 |
| FOXO3 | 0.439 | 0.254 |
| LPAR1 | 0.44 | 0.928 |
| SB203580 | 0.445 | 0.224 |
| NELFB | 0.447 | 0.447 |
| Hdac | 0.453 | 0.728 |
| lonafarnib | 0.453 | 1.633 |
| MUC1 | 0.459 | 1.639 |
| ADCYAP1 | 0.473 | 1.49 |
| CX3CL1 | 0.477 | 1.291 |
| CDH2 | 0.477 | 1.969 |
| tetrodotoxin | 0.479 | 1.248 |
| PD184352 | 0.483 | 1.082 |
| MAP2K3 | 0.485 | 1.51 |
| Bvht | 0.491 | 1.114 |
| Smad | 0.497 | 0.444 |
| TEAD2 | 0.5 | 1.134 |
| TEAD3 | 0.5 | 1.414 |
| trans-cinnamaldehyde | 0.507 | 0.103 |
| gemfibrozil | 0.534 | 1.36 |
| TFAM | 0.549 | 1.982 |
| ACOX1 | 0.552 | 1.143 |
| doxorubicin | 0.552 | 2.029 |
| RIPK2 | 0.552 | 3.503 |
| CIC | 0.557 | 1.069 |
| IL12 (complex) | 0.564 | 1.949 |
| CDK8 | 0.568 | 0.223 |
| trovafloxacin | 0.571 | 0.18 |
| amiodarone | 0.577 | 0.333 |
| CGA | 0.583 | 0.507 |
| trans-hydroxytamoxifen | 0.59 | 1.456 |
| alitretinoin | 0.597 | 0.585 |
| HOXA4 | 0.61 | 0.126 |
| TFAP2A | 0.613 | 0.55 |
| carbon monoxide | 0.617 | 0.838 |

| | | |
|---|---|---|
| MYOCD | 0.629 | 1.042 |
| GSK3B | 0.635 | 0.145 |
| pimozide | 0.64 | 0.762 |
| EGOT | 0.64 | 1.118 |
| simvastatin | 0.642 | 1.288 |
| docetaxel | 0.654 | 1.975 |
| CEACAM1 | 0.664 | 0.478 |
| HDAC1 | 0.665 | 0.154 |
| TLR7 | 0.667 | 0.934 |
| LEPR | 0.692 | 0.686 |
| TCF7L1 | 0.696 | 0.777 |
| HAVCR1 | 0.702 | 2.128 |
| KCNJ10 | 0.707 | 1.342 |
| PLK2 | 0.707 | 2.333 |
| PLK4 | 0.707 | 2.333 |
| PTC-209 | 0.715 | 1.168 |
| MM-589 | 0.715 | 1.534 |
| plicamycin | 0.722 | 0.732 |
| BAX | 0.73 | 1.2 |
| NOS2 | 0.731 | 0.624 |
| SCD | 0.736 | 1.415 |
| SMARCA4 | 0.738 | 1.992 |
| Sn50 peptide | 0.746 | 1.191 |
| kaempferol | 0.747 | 0.89 |
| LDB1 | 0.75 | 0.365 |
| LMO2 | 0.75 | 0.539 |
| camptothecin | 0.759 | 1.132 |
| diphtheria toxin | 0.762 | 1 |
| ABCA1 | 0.764 | 1.118 |
| PTPRJ | 0.775 | 0.577 |
| SUMO2 | 0.777 | 0.213 |
| SOX9 | 0.801 | 1.474 |
| S1PR4 | 0.832 | 0.302 |
| MEF2C | 0.839 | 2.365 |
| TLR9 | 0.847 | 1.999 |
| leucine | 0.859 | 1.985 |
| TGFBR1 | 0.86 | 0.423 |
| SATB1 | 0.867 | 0.697 |
| imipramine | 0.873 | 0.254 |
| everolimus | 0.882 | 0.988 |
| ETV5 | 0.893 | 1.257 |
| LAMP2 | 0.896 | 0.093 |
| BAPTA-AM | 0.907 | 0.759 |
| cycloheximide | 0.909 | 2.304 |
| PPARGC1B | 0.919 | 2.559 |
| WNT5A | 0.927 | 0.431 |
| L-methionine | 0.936 | 0.793 |
| V-PYRRO/NO | 0.956 | 1.109 |
| TINCR | 0.995 | 1.432 |
| CDKN2B | 1 | 0.447 |
| ATXN3 | 1 | 0.816 |
| fluocinolone acetonide | 1 | 1.134 |
| NCOA5 | 1 | 2.186 |
| ISGF3 | 1 | 2.408 |

| | | |
|---|---|---|
| bardoxolone methyl | 1.003 | 1.159 |
| SB-431542 | 1.008 | 0.689 |
| 3M-002 | 1.016 | 1.463 |
| SHR2554 | 1.023 | 0.017 |
| SLC7A5 | 1.026 | 0.277 |
| conjugated linoleic acid | 1.035 | 0.128 |
| zinc protoporphyrin IX | 1.065 | 2.155 |
| MEF2A | 1.065 | 3.209 |
| STAT1/3/5 dimer | 1.066 | 1.387 |
| 3,3'-diindolylmethane | 1.067 | 1.735 |
| SMAD7 | 1.076 | 1.859 |
| ascorbic acid | 1.091 | 0.342 |
| mir-148 | 1.092 | 2.261 |
| 4-phenylbutyric acid | 1.099 | 1.977 |
| TP53 | 1.126 | 1.374 |
| TOB1 | 1.127 | 0.096 |
| pyrrolidine dithiocarbamate | 1.15 | 1.016 |
| pimagedine | 1.16 | 1.408 |
| PSEN1 | 1.164 | 0.674 |
| E2F4 | 1.175 | 0.13 |
| SOX10 | 1.19 | 0.936 |
| 2-hydroxy-1-naphthylaldehyde isonicotinoyl hydrazone | 1.192 | 0.132 |
| GJA1 | 1.192 | 0.728 |
| cercosporin | 1.192 | 1.131 |
| ZNF395 | 1.192 | 1.477 |
| SIN3A | 1.195 | 0.896 |
| TXNIP | 1.212 | 0.34 |
| SMAD1 | 1.224 | 0.842 |
| eicosapentenoic acid | 1.226 | 0.849 |
| BTK | 1.23 | 1.308 |
| triptolide | 1.233 | 1.731 |
| 15-deoxy-delta-12,14 -PGJ 2 | 1.239 | 0.083 |
| Map3k7 | 1.244 | 0.962 |
| xanthine derivative CB002 analog 4 | 1.265 | 0.447 |
| FUS-DDIT3 | 1.265 | 0.816 |
| PRL | 1.273 | 2.335 |
| sulindac | 1.276 | 1.015 |
| mir-133 | 1.291 | 1.27 |
| CHUK | 1.291 | 1.282 |
| Ifn gamma | 1.297 | 2.348 |
| N-[N-(3,5-difluorophenacetyl-L-Ala)]-S-phenylglycine t-butyl ester | 1.315 | 1.128 |
| N-nitro-L-arginine methyl ester | 1.317 | 0.29 |
| EZR | 1.317 | 1.889 |
| Immunoglobulin | 1.317 | 2.931 |
| PARP9 | 1.331 | 2.556 |
| isotretinoin | 1.335 | 1.212 |
| BTNL2 | 1.342 | 1.387 |
| MACROH2A1 | 1.343 | 1.554 |
| lipopolysaccharide | 1.37 | 2.348 |
| TBX5 | 1.376 | 1.618 |
| cetuximab | 1.377 | 0.445 |
| APP | 1.377 | 2.382 |
| SIRT6 | 1.379 | 0.611 |
| PAX5 | 1.387 | 0.707 |

| | | |
|---|---|---|
| CERS2 | 1.387 | 1.982 |
| embelin | 1.402 | 1.965 |
| ATP2A2 | 1.408 | 2.213 |
| MDL 73811 | 1.414 | 0.174 |
| helenalin | 1.414 | 1.969 |
| NKX2-1 | 1.421 | 1.871 |
| rottlerin | 1.431 | 1.742 |
| ALDH2 | 1.432 | 1.414 |
| AZ-1 | 1.437 | 1.32 |
| mir-30 | 1.446 | 0.505 |
| TWIST2 | 1.447 | 0.479 |
| dacinostat | 1.448 | 0.637 |
| PIK3R2 | 1.452 | 1.285 |
| DICER1 | 1.455 | 1.21 |
| NFATC2 | 1.467 | 2.145 |
| palbociclib | 1.469 | 0.964 |
| miR-30c-5p (and other miRNAs w/seed GUAAACA) | 1.471 | 2.259 |
| TLR4 | 1.474 | 1.227 |
| IGF2BP1 | 1.478 | 0.49 |
| TMPRSS2-ERG | 1.48 | 3.411 |
| Alpha catenin | 1.481 | 1.033 |
| IKBKG | 1.486 | 1.69 |
| TERT | 1.49 | 1.923 |
| noscapine | 1.501 | 2.216 |
| ZBTB16 | 1.504 | 1.478 |
| Interferon alpha | 1.505 | 3.577 |
| pembrolizumab | 1.508 | 1.463 |
| IL21 | 1.511 | 2.263 |
| uric acid | 1.518 | 0.079 |
| OPA1 | 1.519 | 0.831 |
| mir-455 | 1.524 | 1.96 |
| NGFR | 1.53 | 0.6 |
| STAT2 | 1.531 | 2.03 |
| mir-320 | 1.534 | 1.982 |
| TIMP3 | 1.546 | 0.707 |
| CCN5 | 1.55 | 0.113 |
| PD173074 | 1.551 | 1.456 |
| selumetinib | 1.554 | 1.347 |
| FADD | 1.556 | 1.676 |
| LHX1 | 1.589 | 1.732 |
| amino acids | 1.589 | 1.763 |
| BAP1 | 1.592 | 0.848 |
| curcumin | 1.597 | 1.826 |
| ARHGAP21 | 1.604 | 2.111 |
| Ro31-8220 | 1.607 | 1.929 |
| CNTF | 1.624 | 2.108 |
| morin | 1.633 | 1.982 |
| H-7 | 1.636 | 0.743 |
| staurosporine | 1.638 | 0.368 |
| FCGR2A | 1.64 | 1.562 |
| SGPL1 | 1.644 | 0.152 |
| salubrinal | 1.662 | 1.706 |
| TASL | 1.667 | 2.333 |
| miR-144-3p (miRNAs w/seed ACAGUAU) | 1.671 | 2.183 |

| | | |
|---|---|---|
| RORC | 1.673 | 1.118 |
| LIF | 1.683 | 1.575 |
| CBX7 | 1.695 | 2.416 |
| PAF1 | 1.698 | 1.387 |
| PP2/AG1879 tyrosine kinase inhibitor | 1.701 | 1.976 |
| picropodophyllin | 1.709 | 0.927 |
| FFAR3 | 1.723 | 0.507 |
| IFNE | 1.732 | 1.265 |
| JAK1/2 | 1.732 | 2.828 |
| SEL1L | 1.741 | 2.2 |
| IFRD1 | 1.744 | 1.387 |
| salinosporamide A | 1.757 | 2.2 |
| arsenic trioxide | 1.763 | 0.15 |
| CTF1 | 1.797 | 2.222 |
| SP600125 | 1.799 | 0.245 |
| ZFP36 | 1.813 | 2.021 |
| caffeic acid phenethyl ester | 1.819 | 1.988 |
| ATP2B2 | 1.828 | 2.523 |
| MXD1 | 1.838 | 0.389 |
| imiquimod | 1.84 | 2.402 |
| ELOVL5 | 1.846 | 1.519 |
| tretinoin | 1.849 | 4.175 |
| topotecan | 1.86 | 0.05 |
| concanavalin a | 1.86 | 1.207 |
| G protein alpha i | 1.881 | 3.096 |
| apigenin | 1.884 | 1.863 |
| KDM3B | 1.886 | 0.302 |
| evodiamine | 1.89 | 0.332 |
| NEIL2 | 1.89 | 0.692 |
| BMS-754807 | 1.89 | 1 |
| gossypin | 1.89 | 2.236 |
| CBL | 1.891 | 1.562 |
| GSK2816126 | 1.894 | 1.286 |
| HSPA5 | 1.894 | 1.375 |
| N-acetyl-L-cysteine | 1.9 | 3.028 |
| GPS2 | 1.905 | 0.597 |
| Sch-23390 | 1.919 | 2.335 |
| KLF2 | 1.927 | 1.847 |
| withaferin A | 1.929 | 0.305 |
| KLF3 | 1.934 | 1.671 |
| GW3965 | 1.937 | 2.313 |
| CLEC4A | 1.941 | 2.213 |
| IL27 | 1.944 | 2.934 |
| A-769662 | 1.951 | 1.987 |
| IFNA21 | 1.961 | 1.973 |
| IFNA16 | 1.961 | 1.976 |
| IFNA8 | 1.961 | 1.976 |
| IFNA6 | 1.961 | 1.976 |
| IFNA7 | 1.961 | 1.976 |
| IFNA5 | 1.961 | 1.976 |
| IFNA10 | 1.961 | 1.976 |
| DIABLO | 1.963 | 2.202 |
| decursin | 1.964 | 1.091 |
| HUWE1 | 1.967 | 0.075 |

| | | |
|---|---|---|
| dihydroartemisinin | 1.969 | 0.477 |
| piceatannol | 1.97 | 1.947 |
| U73122 | 1.976 | 0.436 |
| perifosine | 1.98 | 0.68 |
| L-cystine | 1.98 | 1.981 |
| PHB2 | 1.98 | 1.982 |
| setanaxib | 1.98 | 1.982 |
| DHCR24 | 1.982 | 1.342 |
| MIA-602 | 1.982 | 1.982 |
| sunitinib | 1.988 | 1.278 |
| xanthohumol | 1.994 | 0.447 |
| poly dA-dT | 1.994 | 1.848 |
| ADRB1 | 2 | 1.987 |
| GNE | 2 | 2 |
| GAS2L3 | 2 | 2.433 |
| DMH1 | 2 | 2.449 |
| wortmannin | 2.01 | 1.346 |
| thalidomide | 2.01 | 2.173 |
| ibrutinib | 2.011 | 1.221 |
| grape seed extract | 2.024 | 0.507 |
| PRKN | 2.031 | 1.408 |
| butyric acid | 2.037 | 0.574 |
| emodin | 2.038 | 0.458 |
| SLC6A4 | 2.04 | 1.134 |
| tazemetostat | 2.04 | 1.428 |
| RBM5 | 2.043 | 1.97 |
| propylthiouracil | 2.046 | 0.165 |
| NDRG1 | 2.049 | 1.794 |
| HIC1 | 2.052 | 0.207 |
| pristane | 2.073 | 2.573 |
| IFIH1 | 2.077 | 2.346 |
| mir-26 | 2.082 | 1.628 |
| HDAC3 | 2.092 | 1.773 |
| ITPR2 | 2.111 | 3.16 |
| ZFP91 | 2.121 | 1 |
| methylselenic acid | 2.138 | 1.387 |
| POLR2M | 2.138 | 1.698 |
| HNF4A | 2.162 | 1.302 |
| NFIX | 2.163 | 2.163 |
| triamcinolone acetonide | 2.17 | 1.809 |
| Bay 11-7082 | 2.17 | 2.024 |
| ursolic acid | 2.177 | 1.844 |
| gefitinib | 2.185 | 0.899 |
| SCH 58261 | 2.19 | 1.231 |
| imatinib | 2.2 | 0.491 |
| U18666A | 2.2 | 2.2 |
| DOCK8 | 2.2 | 2.711 |
| CREBZF | 2.211 | 0.398 |
| MEF2D | 2.211 | 1.744 |
| RPL22 | 2.213 | 2.219 |
| CLEC12A | 2.213 | 2.425 |
| NODAL | 2.219 | 0.478 |
| alvocidib | 2.232 | 2.621 |
| SP2509 | 2.233 | 1.91 |

| | | |
|---|---|---|
| L-histidine | 2.234 | 2.195 |
| bosutinib | 2.236 | 2.236 |
| IFN alpha/beta | 2.237 | 3.582 |
| GIP | 2.246 | 2.757 |
| COL18A1 | 2.261 | 1.618 |
| SASH1 | 2.268 | 2.858 |
| baicalein | 2.27 | 1.964 |
| FOXC1 | 2.286 | 3.024 |
| BCAP31 | 2.313 | 1.474 |
| entinostat | 2.323 | 0.922 |
| ELOVL3 | 2.324 | 0.577 |
| TNK1 | 2.333 | 3 |
| mir-802 | 2.343 | 1.46 |
| THZ2 | 2.345 | 0.447 |
| ACVRL1 | 2.345 | 1.633 |
| CDKN1A | 2.346 | 0.196 |
| DGCR8 | 2.369 | 2.022 |
| SU6656 | 2.384 | 1.913 |
| miR-503-5p (miRNAs w/seed AGCAGCG) | 2.393 | 0.817 |
| TICAM1 | 2.403 | 2.322 |
| FTI-277 | 2.407 | 1.103 |
| RARB | 2.413 | 2.552 |
| H89 | 2.414 | 0.78 |
| APC | 2.414 | 1.953 |
| FZD9 | 2.415 | 1.699 |
| CPT1B | 2.417 | 2.846 |
| TLR3 | 2.422 | 2.098 |
| epothilone B | 2.425 | 1.432 |
| 2'3'-cyclic guanosine monophosphate-adenosine monophosphate | 2.425 | 2.051 |
| MSR1 | 2.425 | 2.425 |
| TMEM120A | 2.425 | 2.606 |
| TRIB3 | 2.427 | 0.584 |
| CDN1163 | 2.449 | 1.432 |
| 1'-acetoxychavicol acetate | 2.449 | 2.236 |
| IFN Lambda | 2.449 | 2.236 |
| TNFSF10 | 2.449 | 2.737 |
| TGM2 | 2.457 | 4.093 |
| bisindolylmaleimide I | 2.459 | 1.007 |
| calphostin C | 2.477 | 2.32 |
| MAP2K6 | 2.519 | 2.797 |
| JAK | 2.525 | 2.121 |
| thiostrepton | 2.528 | 0.762 |
| REST | 2.536 | 2.963 |
| aldesleukin | 2.538 | 2.6 |
| CpG ODN 2006 | 2.538 | 3.177 |
| napabucasin | 2.546 | 2.646 |
| SAMSN1 | 2.556 | 2.887 |
| RB1 | 2.559 | 1.356 |
| Go 6976 | 2.575 | 1.159 |
| mir-27 | 2.578 | 0.271 |
| miR-143-3p (and other miRNAs w/seed GAGAUGA) | 2.584 | 1.941 |
| MAPKAP1 | 2.621 | 2.4 |
| Duxbl1 | 2.63 | 2.449 |
| dactolisib | 2.635 | 2.183 |

| | | |
|---|---|---|
| EGLN | 2.643 | 1.256 |
| tanespimycin | 2.645 | 1.848 |
| ETV6-NTRK3 | 2.646 | 2.646 |
| let-7a-5p (and other miRNAs w/seed GAGGUAG) | 2.675 | 2.467 |
| RBPJ | 2.695 | 3.266 |
| RUNX3 | 2.697 | 1.672 |
| TGS1 | 2.704 | 2.331 |
| miR-34a-5p (and other miRNAs w/seed GGCAGUG) | 2.725 | 0.986 |
| mir-34 | 2.73 | 1.794 |
| bromodeoxyuridine | 2.73 | 3.043 |
| IFNG | 2.748 | 4.033 |
| IFNA14 | 2.768 | 2.58 |
| PPP1R15B | 2.777 | 2.2 |
| mir-145 | 2.786 | 0.697 |
| miR-483-3p (miRNAs w/seed CACUCCU) | 2.787 | 1.826 |
| miR-291a-3p (and other miRNAs w/seed AAGUGCU) | 2.788 | 2.881 |
| docosahexaenoic acid | 2.791 | 2.877 |
| N-cor | 2.804 | 2.407 |
| IFNL3 | 2.804 | 2.951 |
| IFNL4 | 2.807 | 2.639 |
| oblimersen | 2.828 | 1.872 |
| KDM5B | 2.831 | 0.059 |
| IFNA4 | 2.858 | 3.594 |
| roscovitine | 2.9 | 1.62 |
| DACH1 | 2.913 | 2.36 |
| TCF3 | 2.914 | 1.324 |
| U0126 | 2.916 | 0.797 |
| ANGPTL4 | 2.923 | 1.92 |
| minocycline | 2.934 | 0.552 |
| pregna-4,17-diene-3,16-dione | 2.936 | 2.207 |
| mir-96 | 2.942 | 2.54 |
| IFNAR2 | 3 | 2.236 |
| TERF2IP | 3 | 2.333 |
| PTEN | 3.001 | 1.478 |
| adavosertib | 3.015 | 1.826 |
| HMOX1 | 3.019 | 1.368 |
| EIF3E | 3.057 | 1.369 |
| SPARC | 3.063 | 2.575 |
| ZBTB18 | 3.07 | 0.896 |
| miR-92a-3p (and other miRNAs w/seed AUUGCAC) | 3.102 | 2.399 |
| MPT0B291 | 3.116 | 2.42 |
| SENP3 | 3.154 | 4.146 |
| THZ531 | 3.157 | 2.324 |
| infliximab | 3.162 | 1.772 |
| SAHM1 | 3.162 | 2 |
| emactuzumab | 3.162 | 3 |
| BNIP3L | 3.165 | 2.132 |
| PAX6 | 3.18 | 2.401 |
| miR-17-5p (and other miRNAs w/seed AAAGUGC) | 3.203 | 2.884 |
| silibinin | 3.217 | 2.271 |
| IRF5 | 3.247 | 3.103 |
| CYB5R4 | 3.273 | 2.798 |
| sirolimus | 3.292 | 1.91 |
| ribavirin | 3.293 | 3.148 |

| | | |
|---|---|---|
| IFNAR1 | 3.295 | 5.468 |
| Rb | 3.297 | 0.447 |
| SKIV2L | 3.3 | 3 |
| diphenyleneiodonium | 3.307 | 2.793 |
| CGAS | 3.311 | 3.69 |
| IFNA1/IFNA13 | 3.318 | 2.91 |
| poly rI:rC-RNA | 3.329 | 4.671 |
| GRIN3A | 3.335 | 2.025 |
| IFN Beta | 3.383 | 4.048 |
| imipramine blue | 3.399 | 1.078 |
| SNCA | 3.407 | 2.429 |
| miR-145-5p (and other miRNAs w/seed UCCAGUU) | 3.43 | 2.59 |
| STING1 | 3.43 | 5.087 |
| IFN type 1 | 3.513 | 3.246 |
| stallimycin | 3.523 | 3.404 |
| MIR17HG | 3.53 | 1.163 |
| mir-122 | 3.612 | 2.259 |
| Ifnar | 3.628 | 4.538 |
| CDK19 | 3.693 | 2.242 |
| glutamine | 3.695 | 2.426 |
| miR-155-5p (miRNAs w/seed UAAUGCU) | 3.709 | 0.996 |
| Ifn | 3.74 | 3.826 |
| mir-8 | 3.742 | 3.118 |
| mir-183 | 3.804 | 3.431 |
| PML | 3.808 | 2.733 |
| PD98059 | 3.858 | 3.09 |
| miR-16-5p (and other miRNAs w/seed AGCAGCA) | 3.88 | 1.84 |
| vorinostat | 3.896 | 1.69 |
| TFRC | 3.9 | 1.872 |
| IRF1 | 3.925 | 3.363 |
| SMARCB1 | 3.931 | 2.447 |
| fulvestrant | 3.987 | 1.08 |
| TWNK | 3.988 | 2.449 |
| RNY3 | 4 | 3.742 |
| mir-15 | 4.049 | 2.866 |
| LY294002 | 4.102 | 2.784 |
| ZBTB10 | 4.107 | 4.968 |
| MAVS | 4.124 | 3.276 |
| HDL-cholesterol | 4.206 | 3.564 |
| mir-1 | 4.213 | 2.615 |
| torkinib | 4.214 | 0.561 |
| STAT1 | 4.218 | 5.35 |
| LONP1 | 4.231 | 1.342 |
| N-ethyl-N-nitrosourea | 4.257 | 3.162 |
| DDX58 | 4.591 | 3.278 |
| SPI1 | 4.682 | 3.436 |
| ARVib-7 | 4.811 | 1.89 |
| ARVib-31 | 4.811 | 1.89 |
| miR-1-3p (and other miRNAs w/seed GGAAUGU) | 4.874 | 1.412 |
| IFNL1 | 4.891 | 4.488 |
| IFNB1 | 4.949 | 4.825 |
| CDKN2A | 4.991 | 1.452 |
| miR-124-3p (and other miRNAs w/seed AAGGCAC) | 5.041 | 2.683 |
| NONO | 5.101 | 4.704 |

| | | |
|---|---|---|
| let-7 | 5.285 | 3.214 |
| 1,1-diarylethylene-diammonium derivative | 5.84 | 4.707 |
| IFNA2 | 5.975 | 6.294 |
| IRF3 | 5.977 | 5.428 |
| pyridostatin | 6.113 | 5.372 |
| l-asparaginase | 6.192 | 1.115 |
| IRF7 | 6.279 | 6.091 |

| | A-367 NES | A-367 adj. p | KAPP NES | KAPP adj. p |
|---|---|---|---|---|
| TOP2A | -12.944378 | 1.55E-34 | -9.38659 | 3.80E-17 |
| FOXM1 | -13.317702 | 1.12E-36 | -8.83547 | 6.12E-15 |
| MKI67 | -10.902203 | 6.91E-24 | -7.54216 | 2.84E-10 |
| CENPF | -10.673179 | 8.34E-23 | -7.54875 | 2.70E-10 |
| GTSE1 | -10.591648 | 2.00E-22 | -7.30431 | 1.71E-09 |
| BUB1B | -11.02891 | 1.70E-24 | -6.62288 | 2.16E-07 |
| RACGAP1 | -10.473474 | 7.02E-22 | -7.02992 | 1.27E-08 |
| NDC80 | -9.3572039 | 5.03E-17 | -7.02157 | 1.35E-08 |
| CHEK1 | -9.4709323 | 1.70E-17 | -6.40004 | 9.53E-07 |
| CCNA2 | -8.3914046 | 2.95E-13 | -6.88149 | 3.64E-08 |
| CENPK | -8.4088504 | 2.54E-13 | -6.60618 | 2.42E-07 |
| CENPI | -8.0341871 | 5.78E-12 | -6.37693 | 1.11E-06 |
| ARHGAP11A | -8.5519236 | 7.43E-14 | -5.75011 | 5.47E-05 |
| E2F8 | -8.8976115 | 3.50E-15 | -5.39282 | 0.0004257 |
| ECT2 | -7.6920311 | 8.89E-11 | -6.08381 | 7.20E-06 |
| PTTG1 | -7.7413119 | 6.04E-11 | -5.81757 | 3.67E-05 |
| UBE2C | -8.5869072 | 5.48E-14 | -4.88683 | 0.0062896 |
| DEPDC1B | -7.4442429 | 5.98E-10 | -5.89288 | 2.33E-05 |
| TMPO | -7.8419468 | 2.72E-11 | -5.49097 | 0.0002454 |
| HASPIN | -7.372392 | 1.03E-09 | -5.34782 | 0.0005464 |
| DEPDC1 | -7.4733631 | 4.80E-10 | -5.18624 | 0.0013171 |
| TIMELESS | -7.170224 | 4.60E-09 | -5.46362 | 0.0002864 |
| ASF1B | -8.344 | 4.41E-13 | -4.12389 | 0.2286595 |
| TTK | -7.3623825 | 1.11E-09 | -4.95819 | 0.0043673 |
| FANCD2 | -7.3836052 | 9.46E-10 | -4.82409 | 0.0086328 |
| E2F2 | -7.6890166 | 9.10E-11 | -4.28713 | 0.1110949 |
| MCM6 | -6.7252627 | 1.08E-07 | -5.03614 | 0.0029156 |
| IQGAP3 | -7.3127429 | 1.61E-09 | -4.36047 | 0.0796594 |
| RAD51 | -6.4982447 | 4.99E-07 | -5.09228 | 0.0021715 |
| TCF19 | -6.6951654 | 1.32E-07 | -4.84538 | 0.0077564 |
| FANCA | -7.2046678 | 3.57E-09 | -4.26176 | 0.1244905 |
| AURKA | -6.9389222 | 2.43E-08 | -4.48481 | 0.0447944 |
| CENPU | -6.6030999 | 2.47E-07 | -4.77308 | 0.0111364 |
| MYBL2 | -7.3721098 | 1.03E-09 | -3.79159 | 0.9187586 |
| BRIP1 | -5.8869202 | 2.42E-05 | -4.75048 | 0.0124563 |
| MCM2 | -6.5825164 | 2.84E-07 | -3.99832 | 0.3915596 |
| MCM4 | -6.0206255 | 1.07E-05 | -4.11385 | 0.2388423 |
| TYMS | -6.1042699 | 6.34E-06 | -3.87086 | 0.6656894 |
| AURKB | -6.2087016 | 3.28E-06 | -3.73088 | 1 |
| SPAG5 | -6.1062518 | 6.26E-06 | -3.59506 | 1 |
| E2F7 | -5.790282 | 4.31E-05 | -3.80004 | 0.8880165 |
| MCM8 | -6.3273226 | 1.53E-06 | -2.83884 | 1 |
| CDK2 | -5.4561047 | 0.000298732 | -3.68375 | 1 |
| MXD3 | -5.8633074 | 2.79E-05 | -3.22607 | 1 |
| CDCA7 | -5.4746186 | 0.000269124 | -3.55737 | 1 |

| | | | | |
|---|---|---|---|---|
| FOXN4 | -5.0045713 | 0.00343645 | -3.77332 | 0.9887696 |
| SUV39H2 | -5.4361462 | 0.000334185 | -3.20825 | 1 |
| ZNF367 | -5.2721301 | 0.000827706 | -2.89255 | 1 |
| ATAD2 | -4.5725981 | 0.029567572 | -3.56441 | 1 |
| NSD2 | -4.6259707 | 0.022885407 | -3.39886 | 1 |
| BRCA2 | -4.6651882 | 0.018925585 | -3.35106 | 1 |
| HELLS | -4.4790116 | 0.046028523 | -3.41423 | 1 |
| TONSL | -4.6729851 | 0.018220856 | -3.0702 | 1 |
| RTKN2 | -4.5157673 | 0.038723356 | -3.03306 | 1 |
| ZWINT | -5.3741221 | 0.00047236 | -2.15525 | 1 |
| ZNF90 | -4.5899964 | 0.027206904 | -2.82384 | 1 |
| ARHGAP11B | -4.0232451 | 0.352331717 | -3.36223 | 1 |
| TRIP13 | -4.7805872 | 0.01072824 | -2.58587 | 1 |
| ZNF695 | -4.7095805 | 0.015236186 | -2.5281 | 1 |
| DEK | -4.1754648 | 0.182530798 | -2.89858 | 1 |
| CDCA7L | -4.1785561 | 0.180067808 | -2.83965 | 1 |
| PCNA | -4.1381632 | 0.214889735 | -2.76914 | 1 |
| ZNF730 | -3.8948781 | 0.603046951 | -2.94252 | 1 |
| E2F1 | -3.5564863 | 1 | -3.27406 | 1 |
| TBC1D31 | -3.7111806 | 1 | -3.05924 | 1 |
| ZNF300 | -4.3438846 | 0.085922821 | -2.36502 | 1 |
| ZRANB3 | -3.5152012 | 1 | -3.05592 | 1 |
| CHAF1B | -3.8229083 | 0.809521517 | -2.73459 | 1 |
| SCML2 | -2.7796594 | 1 | -3.65862 | 1 |
| SUZ12 | -3.1847403 | 1 | -2.97968 | 1 |
| YEATS4 | -3.5838603 | 1 | -2.5153 | 1 |
| ZNF680 | -4.4384334 | 0.055620117 | -1.61313 | 1 |
| SMARCAD1 | -3.6571201 | 1 | -2.30582 | 1 |
| CHAF1A | -3.5676071 | 1 | -2.29464 | 1 |
| HMGCR | -4.2021733 | 0.16226703 | -1.62927 | 1 |
| SOD2 | -2.9369554 | 1 | -2.89388 | 1 |
| EZH2 | -3.5201406 | 1 | -2.2817 | 1 |
| NAMPT | -1.9194215 | 1 | -3.72047 | 1 |
| TRIM32 | -2.5145922 | 1 | -2.8193 | 1 |
| BHLHE40 | -2.5217172 | 1 | -2.78442 | 1 |
| ZNF726 | -2.1529188 | 1 | -3.15164 | 1 |
| PPP1R15A | -2.3836125 | 1 | -2.81523 | 1 |
| MED13 | -2.733335 | 1 | -2.44802 | 1 |
| TRAIP | -3.1736072 | 1 | -2.00126 | 1 |
| CEBPG | -2.3571961 | 1 | -2.81462 | 1 |
| CKS2 | -2.9129507 | 1 | -2.22522 | 1 |
| PIK3R2 | -2.8239376 | 1 | -2.30252 | 1 |
| ZNF678 | -2.8902956 | 1 | -2.20395 | 1 |
| MCM5 | -3.6296986 | 1 | -1.44215 | 1 |
| SRPK1 | -3.3750404 | 1 | -1.67999 | 1 |
| HMGB2 | -2.793656 | 1 | -2.22868 | 1 |

| | | | | |
|---|---|---|---|---|
| LIMD1 | -2.4959233 | 1 | -2.52106 | 1 |
| TGFBR1 | -3.1550199 | 1 | -1.78466 | 1 |
| BACH1 | -2.5735949 | 1 | -2.34353 | 1 |
| POLE | -2.9858029 | 1 | -1.91599 | 1 |
| ZIK1 | -2.421017 | 1 | -2.45278 | 1 |
| SHOC2 | -1.9352348 | 1 | -2.92625 | 1 |
| ARID5A | -2.101879 | 1 | -2.74278 | 1 |
| ZNF92 | -1.9492401 | 1 | -2.87712 | 1 |
| YARS1 | -2.1841677 | 1 | -2.62094 | 1 |
| ILF2 | -2.5601784 | 1 | -2.16987 | 1 |
| ADNP2 | -0.8665425 | 1 | -3.81931 | 0.821413 |
| PEG10 | -1.7017985 | 1 | -2.95578 | 1 |
| RFC4 | -2.6982008 | 1 | -1.95783 | 1 |
| MSH2 | -3.4318717 | 1 | -1.20097 | 1 |
| DNMT3B | -2.9239741 | 1 | -1.69919 | 1 |
| ZDHHC21 | -2.2148564 | 1 | -2.39383 | 1 |
| ZBTB38 | -2.0597844 | 1 | -2.50387 | 1 |
| OR5D16 | -2.2147057 | 1 | -2.34146 | 1 |
| ZHX1 | -1.744338 | 1 | -2.76069 | 1 |
| JRKL | -2.6199229 | 1 | -1.88244 | 1 |
| ZNF519 | -2.6150183 | 1 | -1.87668 | 1 |
| HCFC2 | -2.9957121 | 1 | -1.48831 | 1 |
| HSPA5 | -1.8349288 | 1 | -2.60156 | 1 |
| CHEK2 | -3.4698237 | 1 | -0.95841 | 1 |
| GPRC5A | -1.9190103 | 1 | -2.49711 | 1 |
| GPR85 | -2.1973543 | 1 | -2.19005 | 1 |
| CKS1B | -2.408581 | 1 | -1.96452 | 1 |
| SOX11 | -1.8819661 | 1 | -2.48361 | 1 |
| PBX3 | -1.7180486 | 1 | -2.61057 | 1 |
| FANCG | -2.9179792 | 1 | -1.40574 | 1 |
| MYB | -2.2207818 | 1 | -2.09431 | 1 |
| KSR2 | -2.272713 | 1 | -2.00737 | 1 |
| ARL5B | -2.6045538 | 1 | -1.67453 | 1 |
| AP3S1 | -1.4456763 | 1 | -2.81722 | 1 |
| PTPRJ | -1.8893873 | 1 | -2.36579 | 1 |
| LRP12 | -1.9573433 | 1 | -2.29602 | 1 |
| H2AX | -2.591534 | 1 | -1.64569 | 1 |
| MAP4K5 | -2.1612352 | 1 | -2.06153 | 1 |
| HACD1 | -2.9966621 | 1 | -1.22014 | 1 |
| HNRNPD | -2.6646245 | 1 | -1.53554 | 1 |
| CBX6 | -2.6346687 | 1 | -1.54413 | 1 |
| ZNF330 | -2.0885717 | 1 | -2.08638 | 1 |
| ALMS1 | -2.3771688 | 1 | -1.79205 | 1 |
| GUCY1A2 | -2.5707264 | 1 | -1.57431 | 1 |
| ADM2 | -1.717866 | 1 | -2.4249 | 1 |
| ZNF217 | -1.5733814 | 1 | -2.54246 | 1 |

| | | | | |
|---|---|---|---|---|
| ANP32A | -2.7879925 | 1 | -1.3168 | 1 |
| NAA15 | -1.7358142 | 1 | -2.36538 | 1 |
| CGGBP1 | -1.6769462 | 1 | -2.41911 | 1 |
| ZDHHC20 | -2.2288837 | 1 | -1.85885 | 1 |
| KIF5B | -2.0135137 | 1 | -2.06878 | 1 |
| HYCC1 | -1.6402556 | 1 | -2.43077 | 1 |
| KHDRBS1 | -2.6425472 | 1 | -1.36038 | 1 |
| ZMAT4 | -1.6648158 | 1 | -2.32893 | 1 |
| H2BC11 | -2.5049126 | 1 | -1.47151 | 1 |
| E2F3 | -2.3624839 | 1 | -1.60637 | 1 |
| GPRC6A | -1.5522833 | 1 | -2.41318 | 1 |
| SMAD5 | -2.1202191 | 1 | -1.84432 | 1 |
| BMPR2 | -1.3161264 | 1 | -2.64456 | 1 |
| TTL | -1.5352374 | 1 | -2.3919 | 1 |
| SP3 | -2.298397 | 1 | -1.626 | 1 |
| MXI1 | -2.0039727 | 1 | -1.91994 | 1 |
| RBPJ | -3.0454869 | 1 | -0.867 | 1 |
| PATZ1 | -2.1501932 | 1 | -1.75948 | 1 |
| NCOA6 | -2.3089179 | 1 | -1.58863 | 1 |
| FEN1 | -3.1954301 | 1 | -0.69928 | 1 |
| MCM3 | -2.2670945 | 1 | -1.62709 | 1 |
| AKAP12 | -2.0560603 | 1 | -1.82535 | 1 |
| LCOR | -2.3239803 | 1 | -1.55429 | 1 |
| HDGF | -2.8406421 | 1 | -1.0376 | 1 |
| ZNF138 | -1.836717 | 1 | -2.03428 | 1 |
| CSNK1A1 | -1.9963498 | 1 | -1.8635 | 1 |
| IGFBP2 | -2.2699813 | 1 | -1.5867 | 1 |
| ZNF670 | -2.985608 | 1 | -0.86935 | 1 |
| CEBPZ | -2.2233563 | 1 | -1.62358 | 1 |
| RBL1 | -3.0672409 | 1 | -0.75864 | 1 |
| ARHGAP33 | -2.2331273 | 1 | -1.57692 | 1 |
| FOXG1 | -1.602083 | 1 | -2.20579 | 1 |
| ZC4H2 | -2.5320251 | 1 | -1.26194 | 1 |
| ZNF564 | -2.0065053 | 1 | -1.78408 | 1 |
| CNTRL | -1.6619567 | 1 | -2.12727 | 1 |
| ADGRG7 | -2.2270767 | 1 | -1.56038 | 1 |
| KLF5 | -1.8983872 | 1 | -1.88715 | 1 |
| EGLN1 | -2.1684888 | 1 | -1.60431 | 1 |
| DPF3 | -2.0839455 | 1 | -1.6849 | 1 |
| ABL1 | -2.2858795 | 1 | -1.46108 | 1 |
| ASXL1 | -2.2137364 | 1 | -1.53023 | 1 |
| APAF1 | -1.6728516 | 1 | -2.0429 | 1 |
| EIF2AK3 | -1.539871 | 1 | -2.16202 | 1 |
| DUSP9 | -1.5536366 | 1 | -2.13978 | 1 |
| ENPP1 | -2.3752977 | 1 | -1.28403 | 1 |
| EPHB2 | -2.5762079 | 1 | -1.07891 | 1 |

| | | | | |
|---|---|---|---|---|
| RAB14 | -1.8948951 | 1 | -1.75967 | 1 |
| THAP12 | -2.6603391 | 1 | -0.99176 | 1 |
| ZNF280C | -1.7887889 | 1 | -1.85926 | 1 |
| MARCO | -1.9646641 | 1 | -1.6725 | 1 |
| CSNK2A2 | -2.8933972 | 1 | -0.74334 | 1 |
| ZNF268 | -1.9844168 | 1 | -1.64895 | 1 |
| H3.3A | -2.280542 | 1 | -1.34823 | 1 |
| WIZ | -0.9858009 | 1 | -2.63819 | 1 |
| RAP2A | -1.5601269 | 1 | -2.0512 | 1 |
| H4C1 | -2.2662564 | 1 | -1.33979 | 1 |
| SMARCE1 | -1.5786166 | 1 | -2.02276 | 1 |
| EWSR1 | -2.4474845 | 1 | -1.14429 | 1 |
| JAG1 | -2.8740397 | 1 | -0.68807 | 1 |
| RERE | -1.4427387 | 1 | -2.10922 | 1 |
| ZNF518B | -1.6678938 | 1 | -1.87297 | 1 |
| EPHA4 | -1.7737572 | 1 | -1.76674 | 1 |
| ABL2 | -1.9080597 | 1 | -1.63032 | 1 |
| SSRP1 | -2.6881641 | 1 | -0.84668 | 1 |
| RNF2 | -1.6554481 | 1 | -1.87447 | 1 |
| FGD1 | -1.9985497 | 1 | -1.5226 | 1 |
| BCL9 | -2.2429436 | 1 | -1.27699 | 1 |
| FOXJ3 | -1.0884083 | 1 | -2.42402 | 1 |
| SFN | -2.1111735 | 1 | -1.39871 | 1 |
| NFX1 | -2.1801351 | 1 | -1.32737 | 1 |
| CD47 | -1.9704379 | 1 | -1.53263 | 1 |
| RAB42 | -1.5643139 | 1 | -1.93709 | 1 |
| ZNF267 | -1.7047888 | 1 | -1.79547 | 1 |
| PDLIM1 | -2.0612078 | 1 | -1.43295 | 1 |
| IQGAP2 | -1.9231927 | 1 | -1.5678 | 1 |
| GPR173 | -2.7461783 | 1 | -0.74228 | 1 |
| RORB | -1.4390006 | 1 | -2.044 | 1 |
| GOLT1B | -1.5975166 | 1 | -1.84358 | 1 |
| TBC1D30 | -2.4968788 | 1 | -0.92323 | 1 |
| TNFAIP3 | -2.0988847 | 1 | -1.31866 | 1 |
| LMO4 | -2.080803 | 1 | -1.32286 | 1 |
| HIP1 | -2.3056289 | 1 | -1.09344 | 1 |
| NLGN1 | -1.7685401 | 1 | -1.62009 | 1 |
| DDX20 | -1.0297619 | 1 | -2.3518 | 1 |
| TCERG1 | -2.0279135 | 1 | -1.35298 | 1 |
| JUNB | -0.8011566 | 1 | -2.57809 | 1 |
| ARHGEF39 | -2.6427716 | 1 | -0.73315 | 1 |
| LIN28B | -1.7459446 | 1 | -1.6284 | 1 |
| ASIC2 | -1.3748463 | 1 | -1.99617 | 1 |
| YWHAH | -1.342668 | 1 | -2.02682 | 1 |
| CNGA3 | -1.4592974 | 1 | -1.90361 | 1 |
| ZNF462 | -2.082607 | 1 | -1.27653 | 1 |

| | | | | |
|---|---|---|---|---|
| XBP1 | -1.9800492 | 1 | -1.37825 | 1 |
| ZNF124 | -1.8779 | 1 | -1.47789 | 1 |
| H2BC17 | -2.8180172 | 1 | -0.5346 | 1 |
| FGFR1 | -2.6308329 | 1 | -0.72121 | 1 |
| PRKCI | -2.446854 | 1 | -0.90213 | 1 |
| DYRK2 | -1.4908526 | 1 | -1.85655 | 1 |
| ETV6 | -1.7754601 | 1 | -1.56929 | 1 |
| PRKAR2B | -2.5662629 | 1 | -0.76839 | 1 |
| RLIM | -1.9770669 | 1 | -1.35456 | 1 |
| ZNF215 | -1.658041 | 1 | -1.65551 | 1 |
| ADM | -1.6920489 | 1 | -1.61482 | 1 |
| HNRNPDL | -2.2672201 | 1 | -1.03701 | 1 |
| CITED1 | -1.5867713 | 1 | -1.7079 | 1 |
| OR51G2 | -0.918784 | 1 | -2.36913 | 1 |
| TBL1XR1 | -2.1777308 | 1 | -1.10776 | 1 |
| ZFAND5 | -0.1347416 | 1 | -3.14852 | 1 |
| HNRNPK | -2.4356312 | 1 | -0.84105 | 1 |
| MED1 | -2.1559764 | 1 | -1.11881 | 1 |
| RBFOX2 | -1.6216972 | 1 | -1.65174 | 1 |
| ATG10 | -2.6170774 | 1 | -0.65458 | 1 |
| RCOR2 | -1.8910138 | 1 | -1.37454 | 1 |
| NRBF2 | -1.03788 | 1 | -2.2253 | 1 |
| ZKSCAN7 | -1.7043871 | 1 | -1.55756 | 1 |
| HELZ | -1.5751164 | 1 | -1.67354 | 1 |
| OR2Z1 | -1.5564684 | 1 | -1.68832 | 1 |
| BRD7 | -1.3132098 | 1 | -1.92768 | 1 |
| TFCP2 | -1.2189206 | 1 | -2.01722 | 1 |
| BCL11A | -1.6675037 | 1 | -1.56786 | 1 |
| ATRX | -1.282949 | 1 | -1.93944 | 1 |
| SOCS3 | -1.1198714 | 1 | -2.10116 | 1 |
| OGFR | -2.7132235 | 1 | -0.50054 | 1 |
| TAS2R50 | -0.8229645 | 1 | -2.38465 | 1 |
| TBC1D15 | -1.1894002 | 1 | -2.01754 | 1 |
| L3MBTL3 | -0.8212719 | 1 | -2.38536 | 1 |
| ENO1 | -1.0424618 | 1 | -2.16165 | 1 |
| RAB6A | -0.8938075 | 1 | -2.30549 | 1 |
| SHANK2 | -2.0198122 | 1 | -1.17742 | 1 |
| FAT1 | -1.959909 | 1 | -1.23113 | 1 |
| ZNF460 | -0.5476024 | 1 | -2.64292 | 1 |
| XPO1 | -1.0892718 | 1 | -2.09208 | 1 |
| CDC42EP3 | -1.345513 | 1 | -1.83477 | 1 |
| FOSL2 | -1.3287052 | 1 | -1.84587 | 1 |
| RB1CC1 | -1.7645253 | 1 | -1.40847 | 1 |
| CDKN2C | -2.7427505 | 1 | -0.42989 | 1 |
| GRB10 | -1.0082406 | 1 | -2.15982 | 1 |
| POU2F1 | -2.7330897 | 1 | -0.43068 | 1 |

| | | | | |
|---|---|---|---|---|
| MRGPRX1 | -0.7778927 | 1 | -2.38557 | 1 |
| SUV39H1 | -1.6076103 | 1 | -1.54059 | 1 |
| TAF9B | -1.6614618 | 1 | -1.48615 | 1 |
| ZNF440 | -1.7830639 | 1 | -1.36425 | 1 |
| H2AC11 | -2.1320856 | 1 | -1.0102 | 1 |
| ZNF616 | -1.3501074 | 1 | -1.79203 | 1 |
| ZDBF2 | -1.3319043 | 1 | -1.80714 | 1 |
| SNX13 | -2.1369277 | 1 | -1.00204 | 1 |
| ACVR1B | -1.3249922 | 1 | -1.81062 | 1 |
| TAB2 | -1.8941733 | 1 | -1.23713 | 1 |
| KDM5B | -1.9445242 | 1 | -1.17373 | 1 |
| SH3RF1 | -0.9500046 | 1 | -2.16192 | 1 |
| PLAA | -1.747203 | 1 | -1.36248 | 1 |
| RHOBTB1 | -2.1605471 | 1 | -0.94353 | 1 |
| GPR3 | -0.7820835 | 1 | -2.32143 | 1 |
| ZNF624 | -1.2030885 | 1 | -1.89821 | 1 |
| DGKZ | -1.4811944 | 1 | -1.61965 | 1 |
| TRERF1 | -2.1465187 | 1 | -0.94963 | 1 |
| LZTS1 | -0.9400497 | 1 | -2.15322 | 1 |
| SUPT20H | -1.9771195 | 1 | -1.11603 | 1 |
| OTX1 | -2.2308652 | 1 | -0.86136 | 1 |
| KHDRBS3 | -1.8823748 | 1 | -1.20193 | 1 |
| DOCK1 | -2.3319148 | 1 | -0.75143 | 1 |
| RAB18 | -1.9196251 | 1 | -1.15204 | 1 |
| H2AC17 | -2.4269533 | 1 | -0.62944 | 1 |
| RAB30 | -2.0221063 | 1 | -1.03169 | 1 |
| BMPR1B | -0.693433 | 1 | -2.35833 | 1 |
| G3BP2 | -1.9085549 | 1 | -1.13526 | 1 |
| RAB39B | -1.7736701 | 1 | -1.26702 | 1 |
| DICER1 | -1.9302008 | 1 | -1.10738 | 1 |
| ITGA11 | -2.0952575 | 1 | -0.94225 | 1 |
| PIKFYVE | -1.8812301 | 1 | -1.15612 | 1 |
| POU5F2 | -1.4592259 | 1 | -1.57424 | 1 |
| LRRFIP1 | -1.368777 | 1 | -1.65929 | 1 |
| BCL11B | -1.6866513 | 1 | -1.33161 | 1 |
| BRAF | -2.1007417 | 1 | -0.90947 | 1 |
| RHOU | -1.9689304 | 1 | -1.03896 | 1 |
| EML4 | -1.1326987 | 1 | -1.87397 | 1 |
| ZC3H6 | -1.6305455 | 1 | -1.37172 | 1 |
| SAP130 | -1.9406775 | 1 | -1.05744 | 1 |
| DUSP6 | -1.6625635 | 1 | -1.32869 | 1 |
| TUSC3 | -1.4661085 | 1 | -1.5249 | 1 |
| CHAMP1 | -1.712755 | 1 | -1.27266 | 1 |
| MIB1 | -1.4502307 | 1 | -1.5345 | 1 |
| OR4D6 | -1.7379895 | 1 | -1.24324 | 1 |
| CSRNP2 | -1.0178922 | 1 | -1.95875 | 1 |

| | | | | |
|---|---|---|---|---|
| ZNF24 | -1.6415212 | 1 | -1.33183 | 1 |
| H1.3 | -2.3161757 | 1 | -0.65483 | 1 |
| NOCT | -1.0720587 | 1 | -1.88944 | 1 |
| FANCE | -1.6159961 | 1 | -1.34295 | 1 |
| TCF12 | -1.5157539 | 1 | -1.4368 | 1 |
| RYBP | -0.3422403 | 1 | -2.60877 | 1 |
| TAS1R2 | -2.0350767 | 1 | -0.91375 | 1 |
| SENP1 | -1.6534269 | 1 | -1.29439 | 1 |
| TBL1Y | -0.9335317 | 1 | -2.01123 | 1 |
| MSH6 | -1.0253241 | 1 | -1.91737 | 1 |
| IL6 | -0.9099793 | 1 | -2.03148 | 1 |
| MORF4L2 | -0.915578 | 1 | -2.02398 | 1 |
| ZNF507 | -1.6755721 | 1 | -1.25996 | 1 |
| ZNF107 | -1.6410027 | 1 | -1.29054 | 1 |
| RC3H2 | -1.8372949 | 1 | -1.09067 | 1 |
| SMARCC1 | -1.2663276 | 1 | -1.65966 | 1 |
| MCC | -1.8801824 | 1 | -1.04412 | 1 |
| LATS2 | -2.2463121 | 1 | -0.67191 | 1 |
| HMGXB4 | -1.9957396 | 1 | -0.92106 | 1 |
| MEF2A | -0.792767 | 1 | -2.12318 | 1 |
| TTF2 | -2.1586758 | 1 | -0.75554 | 1 |
| IL17RD | -1.2045131 | 1 | -1.70892 | 1 |
| IDE | -1.9581221 | 1 | -0.94934 | 1 |
| ZNF479 | -1.0529074 | 1 | -1.85366 | 1 |
| RNF6 | -0.9011485 | 1 | -2.0033 | 1 |
| CDC73 | -1.3712483 | 1 | -1.53058 | 1 |
| RPS6KA3 | -0.9526354 | 1 | -1.94869 | 1 |
| CASP3 | -2.251821 | 1 | -0.64616 | 1 |
| ITSN2 | -1.3498334 | 1 | -1.54656 | 1 |
| ERCC8 | -1.7468557 | 1 | -1.14645 | 1 |
| ITGA5 | -1.1656876 | 1 | -1.72513 | 1 |
| RNF138 | -1.4961013 | 1 | -1.38829 | 1 |
| IL20RB | -1.2580203 | 1 | -1.62521 | 1 |
| GZF1 | -1.1293072 | 1 | -1.7442 | 1 |
| CHRDL1 | -1.7152134 | 1 | -1.15575 | 1 |
| SOX4 | -2.0890298 | 1 | -0.77868 | 1 |
| SMARCD1 | -2.3431468 | 1 | -0.51489 | 1 |
| CDH2 | -1.756638 | 1 | -1.09999 | 1 |
| CYTH3 | -0.7779526 | 1 | -2.07528 | 1 |
| HSPD1 | -0.7163576 | 1 | -2.1341 | 1 |
| RBBP8 | -0.8933551 | 1 | -1.95532 | 1 |
| MAFG | -1.5911679 | 1 | -1.25413 | 1 |
| LRRN3 | -1.763907 | 1 | -1.08052 | 1 |
| MAP3K20 | -1.0718572 | 1 | -1.77058 | 1 |
| RALA | -1.2624306 | 1 | -1.57984 | 1 |
| ZNF43 | -1.546043 | 1 | -1.29508 | 1 |

| | | | | |
|---|---|---|---|---|
| TAF2 | -1.7845446 | 1 | -1.05493 | 1 |
| RUSC1 | -2.0177834 | 1 | -0.81977 | 1 |
| ADGRL3 | -2.2748587 | 1 | -0.5544 | 1 |
| NUP214 | -2.2263973 | 1 | -0.59095 | 1 |
| CNTLN | -1.6951499 | 1 | -1.11695 | 1 |
| ZFP36L2 | -1.4867465 | 1 | -1.32161 | 1 |
| RAB39A | -1.3649763 | 1 | -1.44296 | 1 |
| DGKB | -0.5460456 | 1 | -2.25891 | 1 |
| RAB9B | -1.682378 | 1 | -1.12128 | 1 |
| ASAH2 | -1.6748415 | 1 | -1.12613 | 1 |
| ATF6 | -0.8836544 | 1 | -1.91725 | 1 |
| OLIG3 | -1.1788329 | 1 | -1.61153 | 1 |
| ATXN1 | -2.4828676 | 1 | -0.30075 | 1 |
| F3 | -2.0668786 | 1 | -0.70612 | 1 |
| GCM1 | -0.831965 | 1 | -1.93939 | 1 |
| NONO | -1.9527962 | 1 | -0.81126 | 1 |
| GLI3 | -1.5994781 | 1 | -1.16128 | 1 |
| ZNF555 | -2.3345515 | 1 | -0.41357 | 1 |
| MAP3K2 | -0.5490936 | 1 | -2.19818 | 1 |
| SETBP1 | -1.6641891 | 1 | -1.08101 | 1 |
| ATXN3L | -1.0420006 | 1 | -1.70147 | 1 |
| CCNT2 | -0.9928689 | 1 | -1.7497 | 1 |
| ATF1 | -0.909016 | 1 | -1.83217 | 1 |
| PTEN | -1.5528205 | 1 | -1.18711 | 1 |
| ZNF66 | -1.7730105 | 1 | -0.95833 | 1 |
| FGD5 | -1.1369105 | 1 | -1.58953 | 1 |
| ATXN7 | -0.9633881 | 1 | -1.75229 | 1 |
| RICTOR | -1.8544068 | 1 | -0.85977 | 1 |
| ZNF286A | -2.0999344 | 1 | -0.61154 | 1 |
| NRXN1 | -1.9414603 | 1 | -0.76758 | 1 |
| ETV5 | -1.5243144 | 1 | -1.1838 | 1 |
| AHCTF1 | -1.431859 | 1 | -1.27175 | 1 |
| PHF6 | -0.6619559 | 1 | -2.03754 | 1 |
| GNG13 | -0.985303 | 1 | -1.71201 | 1 |
| SOX2 | -1.4204097 | 1 | -1.2761 | 1 |
| ZNF74 | -2.2601137 | 1 | -0.41956 | 1 |
| PIK3C2A | -1.3127005 | 1 | -1.3636 | 1 |
| SHC4 | -1.2842201 | 1 | -1.39077 | 1 |
| SRSF10 | -1.5430902 | 1 | -1.12767 | 1 |
| DNAJC27 | -1.85245 | 1 | -0.81767 | 1 |
| FPR2 | -0.611986 | 1 | -2.05562 | 1 |
| TSC22D2 | -0.7769397 | 1 | -1.88501 | 1 |
| TOPORS | -1.4097875 | 1 | -1.2521 | 1 |
| MAGI3 | -1.3650205 | 1 | -1.29156 | 1 |
| CNR1 | -2.2101669 | 1 | -0.44 | 1 |
| TMF1 | -0.6103974 | 1 | -2.03968 | 1 |

| | | | | |
|---|---|---|---|---|
| KISS1R | -0.2578431 | 1 | -2.39109 | 1 |
| CREM | -1.4597408 | 1 | -1.18703 | 1 |
| POLR3C | -1.7216426 | 1 | -0.92269 | 1 |
| ZNF451 | -1.3736817 | 1 | -1.26538 | 1 |
| HOXD9 | -1.4800336 | 1 | -1.15588 | 1 |
| ZIC2 | -0.4509625 | 1 | -2.18468 | 1 |
| OR8H2 | -1.6834232 | 1 | -0.94978 | 1 |
| POLR2A | -0.6699716 | 1 | -1.96261 | 1 |
| PRKDC | -1.2848578 | 1 | -1.34256 | 1 |
| LMO2 | -1.8509895 | 1 | -0.77559 | 1 |
| FER | -0.9872467 | 1 | -1.63321 | 1 |
| SRI | -1.4829231 | 1 | -1.13731 | 1 |
| ZNF257 | -1.3542036 | 1 | -1.26102 | 1 |
| ELAPOR2 | -0.988764 | 1 | -1.62381 | 1 |
| INSM1 | -0.8852843 | 1 | -1.72626 | 1 |
| EXT1 | -0.7191085 | 1 | -1.88677 | 1 |
| TNFRSF21 | -1.712987 | 1 | -0.88621 | 1 |
| RUVBL1 | -1.9708165 | 1 | -0.62238 | 1 |
| TRIM33 | -0.7489368 | 1 | -1.84283 | 1 |
| JMY | -1.4818848 | 1 | -1.10806 | 1 |
| ZNF654 | -0.6655006 | 1 | -1.92238 | 1 |
| BRCA1 | -1.5395048 | 1 | -1.04737 | 1 |
| KDM2B | -1.5402549 | 1 | -1.04471 | 1 |
| SMAD9 | -1.1398922 | 1 | -1.43143 | 1 |
| TULP3 | -1.4587617 | 1 | -1.11213 | 1 |
| ZBTB6 | -1.6556761 | 1 | -0.90951 | 1 |
| RAD17 | -1.8170345 | 1 | -0.74724 | 1 |
| FSTL1 | -0.192366 | 1 | -2.37125 | 1 |
| EAF1 | -1.0123547 | 1 | -1.55044 | 1 |
| TIPARP | -1.7963682 | 1 | -0.76205 | 1 |
| ANXA5 | -2.3429069 | 1 | -0.21465 | 1 |
| GRIK3 | -1.5137984 | 1 | -1.03992 | 1 |
| MTF2 | -1.7312321 | 1 | -0.8206 | 1 |
| ROR2 | -1.0010094 | 1 | -1.54364 | 1 |
| ZBTB10 | -1.0794609 | 1 | -1.46367 | 1 |
| MIER3 | -0.8499416 | 1 | -1.69072 | 1 |
| KRAS | -1.2840412 | 1 | -1.25626 | 1 |
| ZNF229 | -1.3653268 | 1 | -1.16522 | 1 |
| ALCAM | -2.0470746 | 1 | -0.48188 | 1 |
| GABBR2 | -1.428708 | 1 | -1.10001 | 1 |
| GSPT1 | -1.126156 | 1 | -1.39481 | 1 |
| RAB20 | -0.7129633 | 1 | -1.80602 | 1 |
| PRKAR1A | -1.0025553 | 1 | -1.51599 | 1 |
| ZNF728 | -1.9104689 | 1 | -0.60298 | 1 |
| PYGO1 | -0.436614 | 1 | -2.06028 | 1 |
| RAB10 | -1.7031949 | 1 | -0.79307 | 1 |

| | | | | |
|---|---|---|---|---|
| RCHY1 | -1.1983095 | 1 | -1.29639 | 1 |
| CAP1 | -1.3845237 | 1 | -1.10554 | 1 |
| PPIC | -1.0168874 | 1 | -1.46818 | 1 |
| GPR157 | -0.6509235 | 1 | -1.83249 | 1 |
| YAF2 | -1.413757 | 1 | -1.06787 | 1 |
| RALGAPB | -1.7093427 | 1 | -0.76876 | 1 |
| BPTF | -1.8837812 | 1 | -0.59123 | 1 |
| H3C2 | -2.0689976 | 1 | -0.40215 | 1 |
| SMURF2 | -1.9728594 | 1 | -0.49592 | 1 |
| RAB44 | -1.7013724 | 1 | -0.7637 | 1 |
| STAM | -0.6699159 | 1 | -1.7949 | 1 |
| AIFM1 | -1.6815217 | 1 | -0.78291 | 1 |
| CYCS | -1.5905579 | 1 | -0.86823 | 1 |
| ARL8B | -1.3036044 | 1 | -1.15309 | 1 |
| H2AC16 | -2.0133492 | 1 | -0.43855 | 1 |
| OR12D2 | -1.215629 | 1 | -1.2325 | 1 |
| GPR161 | -1.2233363 | 1 | -1.22006 | 1 |
| ZFP69B | -2.2128048 | 1 | -0.2277 | 1 |
| INSIG1 | -1.2116721 | 1 | -1.22748 | 1 |
| S1PR2 | -1.6453109 | 1 | -0.79376 | 1 |
| RAP1B | -1.2233658 | 1 | -1.21506 | 1 |
| ACVR2B | -0.2430369 | 1 | -2.19466 | 1 |
| RFX3 | -1.20271 | 1 | -1.22948 | 1 |
| AR | -0.5654803 | 1 | -1.86266 | 1 |
| ZBTB41 | -1.8408172 | 1 | -0.58514 | 1 |
| GRK5 | -0.2853695 | 1 | -2.13997 | 1 |
| PIK3CA | -1.5259353 | 1 | -0.89933 | 1 |
| YWHAG | -0.9263499 | 1 | -1.49637 | 1 |
| NTRK2 | -3.0102549 | 1 | 0.593212 | 1 |
| IFNAR2 | -1.9138411 | 1 | -0.50028 | 1 |
| OR11L1 | -0.4172227 | 1 | -1.9889 | 1 |
| DHCR24 | -2.3229176 | 1 | -0.08259 | 1 |
| ELP4 | -1.7372622 | 1 | -0.66801 | 1 |
| TNFSF18 | -0.919544 | 1 | -1.4856 | 1 |
| NKX2.2 | -1.298453 | 1 | -1.10423 | 1 |
| NFE2L2 | -0.7835356 | 1 | -1.61668 | 1 |
| CARM1 | -1.6534 | 1 | -0.74586 | 1 |
| ARHGAP25 | -0.54444 | 1 | -1.85337 | 1 |
| ZNF679 | -1.8513845 | 1 | -0.54412 | 1 |
| ZNF484 | -1.8566993 | 1 | -0.53637 | 1 |
| MAFB | -1.0851215 | 1 | -1.30716 | 1 |
| FGG | -0.0340388 | 1 | -2.35621 | 1 |
| RASSF9 | -1.8941397 | 1 | -0.49467 | 1 |
| TAF5L | -1.5997112 | 1 | -0.78851 | 1 |
| RRN3 | -0.4174291 | 1 | -1.96896 | 1 |
| VAV3 | -1.0702024 | 1 | -1.31501 | 1 |

| | | | | |
|---|---|---|---|---|
| RAB1A | -0.1833035 | 1 | -2.20118 | 1 |
| MS4A6E | -1.0636445 | 1 | -1.31808 | 1 |
| ZNF136 | -1.7430863 | 1 | -0.63561 | 1 |
| TAF9 | -1.9171881 | 1 | -0.46066 | 1 |
| KMT2A | -1.5444297 | 1 | -0.83201 | 1 |
| PDE3A | -0.1304771 | 1 | -2.24368 | 1 |
| ZKSCAN2 | -1.1091262 | 1 | -1.26398 | 1 |
| LMO3 | -0.3297902 | 1 | -2.03971 | 1 |
| ZNF765 | -0.6730976 | 1 | -1.69623 | 1 |
| KDM1A | -1.9966885 | 1 | -0.36465 | 1 |
| RB1 | -0.2094234 | 1 | -2.14679 | 1 |
| TCEAL1 | -1.7449904 | 1 | -0.61027 | 1 |
| MAF | -1.096401 | 1 | -1.25237 | 1 |
| PTPRK | -0.8753062 | 1 | -1.45809 | 1 |
| ARNT | -0.7393089 | 1 | -1.58182 | 1 |
| NRG1 | -1.1163462 | 1 | -1.20158 | 1 |
| STK38L | -1.391454 | 1 | -0.92506 | 1 |
| TICAM2 | -1.8213724 | 1 | -0.49502 | 1 |
| UBA3 | -0.8329357 | 1 | -1.47599 | 1 |
| MYOD1 | -0.1835727 | 1 | -2.1214 | 1 |
| GRB2 | -0.9245262 | 1 | -1.37866 | 1 |
| NRAS | -0.720671 | 1 | -1.5781 | 1 |
| ESF1 | -0.8826365 | 1 | -1.41486 | 1 |
| MOAP1 | -1.4866417 | 1 | -0.80975 | 1 |
| OR8J1 | -1.3684596 | 1 | -0.92606 | 1 |
| SLC12A2 | -0.922372 | 1 | -1.37052 | 1 |
| DCBLD2 | -0.9261576 | 1 | -1.3665 | 1 |
| PPP1R12A | -0.7389107 | 1 | -1.55257 | 1 |
| H4C14 | -1.7091441 | 1 | -0.57354 | 1 |
| RGS8 | -1.4866131 | 1 | -0.79409 | 1 |
| GSK3B | -1.5515574 | 1 | -0.72316 | 1 |
| SLIT2 | -1.1812846 | 1 | -1.09299 | 1 |
| CGB8 | -1.1391672 | 1 | -1.13115 | 1 |
| TEAD3 | -1.393692 | 1 | -0.87408 | 1 |
| GRK3 | -1.7639144 | 1 | -0.50031 | 1 |
| AKAP5 | -1.6566097 | 1 | -0.60673 | 1 |
| URI1 | -1.3452733 | 1 | -0.91582 | 1 |
| GDI2 | -1.5677256 | 1 | -0.69182 | 1 |
| TRIM24 | -1.6897861 | 1 | -0.56725 | 1 |
| PLCH1 | -1.3401147 | 1 | -0.91549 | 1 |
| ATF3 | -0.3827426 | 1 | -1.87102 | 1 |
| ICMT | -1.0319794 | 1 | -1.22039 | 1 |
| ZNF735 | -0.7372219 | 1 | -1.51511 | 1 |
| ZBTB5 | -0.6332621 | 1 | -1.61849 | 1 |
| TGIF1 | -1.4759812 | 1 | -0.7712 | 1 |
| GMPS | -1.3078437 | 1 | -0.93622 | 1 |

| | | | | |
|---|---|---|---|---|
| EPHA2 | -0.8206827 | 1 | -1.4212 | 1 |
| KLF17 | -0.150377 | 1 | -2.08909 | 1 |
| BCL3 | -1.1616219 | 1 | -1.07646 | 1 |
| FGFR2 | -1.3622376 | 1 | -0.8755 | 1 |
| SUPT16H | -1.5293318 | 1 | -0.70684 | 1 |
| COPS3 | -1.6241307 | 1 | -0.61167 | 1 |
| ZNF772 | -0.7698564 | 1 | -1.46593 | 1 |
| ALK | -0.9197319 | 1 | -1.30622 | 1 |
| ZNF569 | -1.336137 | 1 | -0.88848 | 1 |
| SELENOS | -1.5011144 | 1 | -0.72218 | 1 |
| IRAK4 | -1.9804598 | 1 | -0.23839 | 1 |
| FGF2 | -1.4270961 | 1 | -0.79163 | 1 |
| UBE2B | -1.2450156 | 1 | -0.96707 | 1 |
| ING2 | -1.0379475 | 1 | -1.17401 | 1 |
| KCNH1 | -0.5846061 | 1 | -1.62417 | 1 |
| LCORL | -1.3471259 | 1 | -0.86118 | 1 |
| ZNF256 | -0.9292517 | 1 | -1.27742 | 1 |
| ZNF639 | -1.2472332 | 1 | -0.95413 | 1 |
| MMP14 | -0.3680779 | 1 | -1.83271 | 1 |
| WDR77 | -1.5218976 | 1 | -0.67753 | 1 |
| RGS22 | -1.4517222 | 1 | -0.74692 | 1 |
| IL12RB2 | -1.1199556 | 1 | -1.07665 | 1 |
| APH1B | -0.6687024 | 1 | -1.52581 | 1 |
| TLE4 | -1.6789908 | 1 | -0.51319 | 1 |
| NR4A2 | -1.2975418 | 1 | -0.89085 | 1 |
| GAS1 | -1.1284875 | 1 | -1.05455 | 1 |
| ELP2 | -1.0183091 | 1 | -1.16251 | 1 |
| GRM1 | -1.3472801 | 1 | -0.83333 | 1 |
| EGF | -1.3028324 | 1 | -0.87689 | 1 |
| SH2D3C | -2.1387925 | 1 | -0.03913 | 1 |
| OR14I1 | -0.9445096 | 1 | -1.23162 | 1 |
| QKI | -2.0524226 | 1 | -0.12284 | 1 |
| AGO2 | -1.4060966 | 1 | -0.7648 | 1 |
| TET1 | -1.6425238 | 1 | -0.52409 | 1 |
| PTPRS | -1.2403939 | 1 | -0.92055 | 1 |
| ZNF441 | -1.6946634 | 1 | -0.46069 | 1 |
| CD2AP | -0.4257058 | 1 | -1.72667 | 1 |
| ZNF70 | -0.9672667 | 1 | -1.18472 | 1 |
| SMAD4 | -1.1759984 | 1 | -0.97491 | 1 |
| ASXL3 | -1.0187655 | 1 | -1.12877 | 1 |
| BMPR1A | -1.151994 | 1 | -0.99052 | 1 |
| OR1L3 | -1.6268916 | 1 | -0.51046 | 1 |
| ZNF727 | -1.1901714 | 1 | -0.9432 | 1 |
| CD58 | -0.939955 | 1 | -1.19058 | 1 |
| GCLC | -0.4244294 | 1 | -1.70288 | 1 |
| MAP3K15 | -1.851702 | 1 | -0.27423 | 1 |

| | | | | |
|---|---|---|---|---|
| FZD5 | -1.4473687 | 1 | -0.67395 | 1 |
| OR6P1 | -1.1537415 | 1 | -0.96639 | 1 |
| TGS1 | -1.2817531 | 1 | -0.83087 | 1 |
| FGF9 | -1.2678186 | 1 | -0.84076 | 1 |
| MED21 | -1.040318 | 1 | -1.06576 | 1 |
| ARL4C | -1.1834452 | 1 | -0.92064 | 1 |
| AGTR2 | -0.463767 | 1 | -1.63882 | 1 |
| CNOT7 | -1.0467785 | 1 | -1.05223 | 1 |
| OR2AJ1 | -1.2548077 | 1 | -0.84404 | 1 |
| ZFP30 | -1.23613 | 1 | -0.85883 | 1 |
| SIAH2 | -0.9861925 | 1 | -1.10687 | 1 |
| TRIP11 | -0.7046925 | 1 | -1.38808 | 1 |
| ZNF436 | -0.8646117 | 1 | -1.22737 | 1 |
| CHD9 | -0.6827897 | 1 | -1.40697 | 1 |
| PATJ | -0.5313814 | 1 | -1.55774 | 1 |
| TAF4B | -0.8502941 | 1 | -1.23872 | 1 |
| CLTC | -1.6494688 | 1 | -0.43938 | 1 |
| CRHR2 | -0.5616097 | 1 | -1.52631 | 1 |
| CAVIN4 | -1.4169383 | 1 | -0.67049 | 1 |
| HTR6 | -0.837808 | 1 | -1.24807 | 1 |
| PTN | -1.8506549 | 1 | -0.23 | 1 |
| PNOC | -1.323812 | 1 | -0.75198 | 1 |
| ZBED1 | -0.9041677 | 1 | -1.16773 | 1 |
| MET | -0.5786271 | 1 | -1.49137 | 1 |
| ARFGEF3 | -1.4833454 | 1 | -0.58408 | 1 |
| GNAT1 | -1.1244219 | 1 | -0.9421 | 1 |
| SUCNR1 | -0.1576216 | 1 | -1.90852 | 1 |
| CREB1 | -1.2509545 | 1 | -0.81449 | 1 |
| ITGAV | -1.6935761 | 1 | -0.36994 | 1 |
| ARID1A | -1.1495279 | 1 | -0.90547 | 1 |
| PLCB1 | -1.8153201 | 1 | -0.23848 | 1 |
| TSHZ2 | -0.8828322 | 1 | -1.17055 | 1 |
| ADGRE5 | -1.2644597 | 1 | -0.78891 | 1 |
| CARS1 | -1.7784092 | 1 | -0.27478 | 1 |
| OR52E8 | -1.0812818 | 1 | -0.97078 | 1 |
| GNAQ | -1.3375537 | 1 | -0.7118 | 1 |
| RCOR1 | -0.900434 | 1 | -1.14842 | 1 |
| ZMYM1 | -1.0939965 | 1 | -0.95384 | 1 |
| ASB7 | -0.4984105 | 1 | -1.54825 | 1 |
| STK39 | -0.6622756 | 1 | -1.38412 | 1 |
| OR9Q2 | -1.0561875 | 1 | -0.98834 | 1 |
| DDX3X | -1.125553 | 1 | -0.91549 | 1 |
| IPO7 | -1.3486419 | 1 | -0.68864 | 1 |
| KAT7 | -1.2384135 | 1 | -0.79724 | 1 |
| GABPB1 | -1.0453021 | 1 | -0.98822 | 1 |
| KRBOX5 | -0.5192317 | 1 | -1.51404 | 1 |

| | | | | |
|---|---|---|---|---|
| CASP8AP2 | -0.7718215 | 1 | -1.25818 | 1 |
| HMGN2 | -0.5644368 | 1 | -1.4647 | 1 |
| CSDE1 | -1.3495489 | 1 | -0.67898 | 1 |
| WLS | -1.8866907 | 1 | -0.13739 | 1 |
| DPF1 | -1.128314 | 1 | -0.89425 | 1 |
| SEMA6A | -1.3106438 | 1 | -0.71149 | 1 |
| JADE1 | -0.6030557 | 1 | -1.41875 | 1 |
| ZFP36 | -1.2593371 | 1 | -0.75853 | 1 |
| ZNF121 | -0.6230523 | 1 | -1.38814 | 1 |
| PAX8 | -0.4691789 | 1 | -1.54011 | 1 |
| KCNK2 | -1.8460273 | 1 | -0.16205 | 1 |
| TLR10 | -0.492708 | 1 | -1.51472 | 1 |
| STX2 | -0.1924927 | 1 | -1.81282 | 1 |
| CRY1 | -1.2259423 | 1 | -0.77909 | 1 |
| MIER1 | -1.0362189 | 1 | -0.96469 | 1 |
| DDIT4L | -1.5835133 | 1 | -0.41423 | 1 |
| PLXNA1 | -0.9750333 | 1 | -1.01747 | 1 |
| NFKBIZ | -0.7020448 | 1 | -1.28928 | 1 |
| NDFIP2 | -1.5402947 | 1 | -0.45041 | 1 |
| PKIA | -0.7627628 | 1 | -1.21637 | 1 |
| RHOQ | -0.7340905 | 1 | -1.24281 | 1 |
| AFF4 | -1.4337267 | 1 | -0.54144 | 1 |
| H4C6 | -0.8117617 | 1 | -1.16251 | 1 |
| FARP1 | -0.7915754 | 1 | -1.18216 | 1 |
| RUNX1 | 0.45772719 | 1 | -2.42874 | 1 |
| DDR1 | -1.4757005 | 1 | -0.48871 | 1 |
| GTF2A1 | -0.3526 | 1 | -1.60782 | 1 |
| CDKN2B | -1.6054084 | 1 | -0.34495 | 1 |
| PIK3R3 | -0.4810306 | 1 | -1.46834 | 1 |
| NEK6 | -1.4337489 | 1 | -0.51439 | 1 |
| ZNF597 | -0.4947787 | 1 | -1.45027 | 1 |
| NSD3 | -1.5382037 | 1 | -0.40664 | 1 |
| HSPB1 | -0.7456108 | 1 | -1.19852 | 1 |
| TTF1 | -1.310122 | 1 | -0.62467 | 1 |
| ACAP2 | -0.624781 | 1 | -1.30339 | 1 |
| ZSWIM6 | -0.9578204 | 1 | -0.96885 | 1 |
| FUBP3 | -0.4241312 | 1 | -1.50186 | 1 |
| TAF13 | -0.9763725 | 1 | -0.94717 | 1 |
| CEBPD | -0.7730403 | 1 | -1.14646 | 1 |
| HMG20A | -1.1645594 | 1 | -0.75487 | 1 |
| SYDE2 | -0.6061468 | 1 | -1.31278 | 1 |
| P2RY1 | -0.5549718 | 1 | -1.35917 | 1 |
| COLEC12 | -0.6666474 | 1 | -1.2466 | 1 |
| PAG1 | -0.9363573 | 1 | -0.97683 | 1 |
| ZXDB | -0.5320481 | 1 | -1.38014 | 1 |
| HSD3B1 | -1.7490548 | 1 | -0.16264 | 1 |

| | | | | |
|---|---|---|---|---|
| DLG5 | -1.3554563 | 1 | -0.55559 | 1 |
| MCTP1 | -1.7387712 | 1 | -0.17114 | 1 |
| RAB21 | -1.2375102 | 1 | -0.67162 | 1 |
| IFT52 | -1.3457703 | 1 | -0.56317 | 1 |
| WASL | -0.8702098 | 1 | -1.03601 | 1 |
| GTF2I | -1.8581032 | 1 | -0.04731 | 1 |
| CERT1 | -1.2053308 | 1 | -0.69795 | 1 |
| PSRC1 | -1.1941053 | 1 | -0.70695 | 1 |
| GRIA2 | -1.6935474 | 1 | -0.2058 | 1 |
| FCRL4 | -0.8379548 | 1 | -1.05729 | 1 |
| ZNF426 | -0.6141746 | 1 | -1.27657 | 1 |
| OR51G1 | -0.5821211 | 1 | -1.30509 | 1 |
| ZNF227 | -0.4202006 | 1 | -1.46533 | 1 |
| GATA3 | -0.1135316 | 1 | -1.77172 | 1 |
| SMAP2 | -0.8492992 | 1 | -1.03488 | 1 |
| SOCS1 | -0.6467412 | 1 | -1.23643 | 1 |
| SMAP1 | -0.5848718 | 1 | -1.29433 | 1 |
| USP33 | -0.981987 | 1 | -0.89721 | 1 |
| IL10 | -0.2102682 | 1 | -1.66869 | 1 |
| LPP | -0.865537 | 1 | -1.01203 | 1 |
| TRIM27 | -1.2845825 | 1 | -0.58899 | 1 |
| PDGFC | -0.9823915 | 1 | -0.88597 | 1 |
| SRGAP1 | -0.9990934 | 1 | -0.8685 | 1 |
| ZFR | -1.5048746 | 1 | -0.36243 | 1 |
| KCNH4 | -1.4451572 | 1 | -0.42209 | 1 |
| ZNF852 | -0.6210934 | 1 | -1.24384 | 1 |
| TBC1D16 | -0.28239 | 1 | -1.58143 | 1 |
| ATF2 | -1.4111793 | 1 | -0.45226 | 1 |
| TRIB1 | -0.2488868 | 1 | -1.61159 | 1 |
| GABRG2 | -0.4024588 | 1 | -1.45163 | 1 |
| MORC1 | -0.0668445 | 1 | -1.78446 | 1 |
| CNIH4 | -1.0226811 | 1 | -0.82674 | 1 |
| ZNF16 | -1.4817318 | 1 | -0.36683 | 1 |
| AHR | -0.334241 | 1 | -1.51147 | 1 |
| PDE8A | -1.3001619 | 1 | -0.54332 | 1 |
| MAGI1 | -1.4744885 | 1 | -0.36757 | 1 |
| MAPK14 | -0.7898171 | 1 | -1.04859 | 1 |
| INSIG2 | -1.0676179 | 1 | -0.77032 | 1 |
| TAPT1 | -0.5257319 | 1 | -1.31076 | 1 |
| SET | -0.8559294 | 1 | -0.9799 | 1 |
| PSD3 | -1.5439572 | 1 | -0.29122 | 1 |
| ZNF385D | -1.2021722 | 1 | -0.63023 | 1 |
| HBZ | -0.0247762 | 1 | -1.80753 | 1 |
| TNFRSF19 | -0.6188965 | 1 | -1.21145 | 1 |
| ZNF33B | -0.9001007 | 1 | -0.93016 | 1 |
| GMFB | 0.23459679 | 1 | -2.06277 | 1 |

| | | | | |
|---|---|---|---|---|
| OR5M1 | -1.6496491 | 1 | -0.17716 | 1 |
| MCM7 | -1.2531133 | 1 | -0.57273 | 1 |
| ERN1 | -0.4999596 | 1 | -1.32266 | 1 |
| ZNF737 | -0.7896375 | 1 | -1.03235 | 1 |
| OR5C1 | -1.3500429 | 1 | -0.46877 | 1 |
| NF1 | -1.2997291 | 1 | -0.51751 | 1 |
| ASCC1 | -0.7760829 | 1 | -1.0406 | 1 |
| DCX | -1.4320044 | 1 | -0.38308 | 1 |
| LGR4 | -0.4162614 | 1 | -1.39816 | 1 |
| CAND1 | -0.6518212 | 1 | -1.16103 | 1 |
| IFNAR1 | -1.3462521 | 1 | -0.46659 | 1 |
| SAV1 | -0.0177456 | 1 | -1.79369 | 1 |
| IRS2 | -0.6372891 | 1 | -1.17255 | 1 |
| TERF1 | -0.8534134 | 1 | -0.95323 | 1 |
| CDK5R1 | -1.5026725 | 1 | -0.30063 | 1 |
| GZMA | -0.7362749 | 1 | -1.06642 | 1 |
| NCSTN | -0.696226 | 1 | -1.10362 | 1 |
| STRN3 | -1.08568 | 1 | -0.71285 | 1 |
| ATF5 | -0.6549423 | 1 | -1.14158 | 1 |
| ZNF644 | -0.9641912 | 1 | -0.83159 | 1 |
| ZNF627 | -1.1711428 | 1 | -0.62304 | 1 |
| ZNF660 | -0.4952925 | 1 | -1.29799 | 1 |
| PTHLH | -0.8651223 | 1 | -0.92657 | 1 |
| ASCC3 | -1.4312252 | 1 | -0.35812 | 1 |
| PKN2 | -0.3746034 | 1 | -1.41406 | 1 |
| TSHZ1 | -1.2671703 | 1 | -0.51837 | 1 |
| TFB1M | -0.9174697 | 1 | -0.86702 | 1 |
| STC1 | -0.8738413 | 1 | -0.91054 | 1 |
| OR5B12 | -0.0734501 | 1 | -1.70875 | 1 |
| JAK2 | -0.3878075 | 1 | -1.3896 | 1 |
| CALCA | -0.3392872 | 1 | -1.43797 | 1 |
| AGFG1 | -0.4897259 | 1 | -1.28602 | 1 |
| SLC26A3 | -0.5896656 | 1 | -1.18522 | 1 |
| ZNF480 | -1.1488509 | 1 | -0.62505 | 1 |
| SMURF1 | -2.2776177 | 1 | 0.506267 | 1 |
| MAGI2 | -0.8242392 | 1 | -0.94604 | 1 |
| OR2J3 | -0.4371711 | 1 | -1.33291 | 1 |
| BCAR3 | 0.44346234 | 1 | -2.21329 | 1 |
| LPAR4 | -1.4924011 | 1 | -0.27493 | 1 |
| ADRA1B | -1.2673429 | 1 | -0.49957 | 1 |
| KDM6A | -0.8459053 | 1 | -0.91673 | 1 |
| OR6B3 | -1.2526012 | 1 | -0.50965 | 1 |
| TBK1 | -0.6561418 | 1 | -1.10608 | 1 |
| ZNF626 | -1.4061118 | 1 | -0.35526 | 1 |
| SP4 | -1.1534589 | 1 | -0.60758 | 1 |
| ITGA3 | -1.8507318 | 1 | 0.097949 | 1 |

| | | | | |
|---|---|---|---|---|
| AIMP1 | -0.1826351 | 1 | -1.56751 | 1 |
| GNAI3 | -1.0358338 | 1 | -0.71328 | 1 |
| CASP2 | -0.4360273 | 1 | -1.31121 | 1 |
| GRIA3 | -0.1969207 | 1 | -1.54788 | 1 |
| CCNE1 | -1.2977794 | 1 | -0.44535 | 1 |
| ZNF84 | -1.1929826 | 1 | -0.54944 | 1 |
| KLF12 | -1.4563642 | 1 | -0.28544 | 1 |
| RARB | -0.9748959 | 1 | -0.76424 | 1 |
| CNGB3 | -0.9179577 | 1 | -0.81981 | 1 |
| CTNNA1 | -1.0734265 | 1 | -0.6604 | 1 |
| CBX2 | -1.3448923 | 1 | -0.3865 | 1 |
| NFKB2 | -0.2199355 | 1 | -1.51104 | 1 |
| ZNF687 | -1.5343246 | 1 | -0.19635 | 1 |
| ANXA1 | -0.0663057 | 1 | -1.66409 | 1 |
| OR52K1 | -1.192256 | 1 | -0.53798 | 1 |
| NKX2.3 | -0.940427 | 1 | -0.78893 | 1 |
| TEAD4 | -0.374202 | 1 | -1.35501 | 1 |
| FGF14 | -1.0523302 | 1 | -0.67366 | 1 |
| BAIAP2 | -0.6695942 | 1 | -1.05415 | 1 |
| SMARCA5 | -1.2399888 | 1 | -0.48282 | 1 |
| DLX5 | -0.733708 | 1 | -0.98844 | 1 |
| ZNF266 | -1.1622463 | 1 | -0.55988 | 1 |
| OR5B2 | -0.5493374 | 1 | -1.17096 | 1 |
| DDIT3 | -1.0886125 | 1 | -0.63127 | 1 |
| ELK4 | -0.8399219 | 1 | -0.87933 | 1 |
| RAB3B | -0.3785022 | 1 | -1.34026 | 1 |
| ISL1 | -0.354225 | 1 | -1.36217 | 1 |
| ADAM9 | -0.4835885 | 1 | -1.23228 | 1 |
| KITLG | -0.4422268 | 1 | -1.27298 | 1 |
| OR52B6 | 0.08525813 | 1 | -1.80039 | 1 |
| OR8K1 | -0.1668243 | 1 | -1.54742 | 1 |
| H4C3 | -1.126774 | 1 | -0.58452 | 1 |
| IGF1R | -0.688299 | 1 | -1.02039 | 1 |
| H2BC4 | -1.5813829 | 1 | -0.12552 | 1 |
| CD59 | -1.7457579 | 1 | 0.039772 | 1 |
| ADNP | -0.4512767 | 1 | -1.25369 | 1 |
| GDNF | -1.3593479 | 1 | -0.34509 | 1 |
| NLK | -0.9490058 | 1 | -0.75348 | 1 |
| GHSR | -0.4263367 | 1 | -1.27466 | 1 |
| ZNF532 | -0.7223407 | 1 | -0.97774 | 1 |
| ATP2A2 | -1.5760669 | 1 | -0.12136 | 1 |
| ARHGEF10 | -0.1652384 | 1 | -1.52821 | 1 |
| MORC3 | -1.1860344 | 1 | -0.50663 | 1 |
| KCNH7 | -1.2496551 | 1 | -0.44052 | 1 |
| ZNF589 | -0.2747777 | 1 | -1.41483 | 1 |
| PARP1 | -1.2845844 | 1 | -0.40357 | 1 |

| | | | | |
|---|---|---|---|---|
| PPP1R2C | -1.2579367 | 1 | -0.43013 | 1 |
| CHD4 | -1.2988882 | 1 | -0.3891 | 1 |
| NR0B2 | 0.09269613 | 1 | -1.7804 | 1 |
| IL17A | -0.9306133 | 1 | -0.75537 | 1 |
| POT1 | -0.9560226 | 1 | -0.72697 | 1 |
| KIT | -0.7034554 | 1 | -0.97739 | 1 |
| PSMC3IP | -1.476822 | 1 | -0.19772 | 1 |
| ITGA2 | -1.0091405 | 1 | -0.66524 | 1 |
| SIN3A | -0.8807357 | 1 | -0.79264 | 1 |
| CNIH2 | -1.8846675 | 1 | 0.211814 | 1 |
| BCKDK | -1.5698088 | 1 | -0.10146 | 1 |
| CSNK1G3 | 0.245331 | 1 | -1.91502 | 1 |
| ZNF71 | -1.607169 | 1 | -0.05842 | 1 |
| DOK4 | -1.614169 | 1 | -0.04723 | 1 |
| PIK3C2G | -0.2023429 | 1 | -1.45848 | 1 |
| GPAT3 | -0.6265923 | 1 | -1.03257 | 1 |
| H1.2 | -1.3130626 | 1 | -0.34258 | 1 |
| PURA | -1.1558215 | 1 | -0.49738 | 1 |
| CCDC47 | -0.1884611 | 1 | -1.4635 | 1 |
| SCRT1 | -1.0028729 | 1 | -0.64879 | 1 |
| ROCK2 | -1.2133985 | 1 | -0.43335 | 1 |
| ETV3 | -0.5477629 | 1 | -1.09662 | 1 |
| SNX27 | -1.3669098 | 1 | -0.2764 | 1 |
| ISX | -1.0209817 | 1 | -0.62195 | 1 |
| ZNF605 | -0.2435957 | 1 | -1.39863 | 1 |
| ZNF442 | -1.1408931 | 1 | -0.50021 | 1 |
| APP | -2.5883433 | 1 | 0.948365 | 1 |
| ELK3 | -0.3168609 | 1 | -1.32287 | 1 |
| MCF2L2 | -1.6024593 | 1 | -0.03625 | 1 |
| TFB2M | -1.075183 | 1 | -0.56196 | 1 |
| MYC | -0.6996854 | 1 | -0.93684 | 1 |
| DNM1L | -1.1749536 | 1 | -0.46138 | 1 |
| TRAK1 | -0.0252734 | 1 | -1.606 | 1 |
| ARL4D | -1.3990656 | 1 | -0.23123 | 1 |
| CERS6 | -0.7531281 | 1 | -0.87459 | 1 |
| SGF29 | -0.6513408 | 1 | -0.97598 | 1 |
| MACC1 | -0.6620896 | 1 | -0.96462 | 1 |
| CEP57 | -1.1170052 | 1 | -0.50823 | 1 |
| CDC42SE2 | -1.2349227 | 1 | -0.39024 | 1 |
| THRAP3 | -1.2329298 | 1 | -0.39121 | 1 |
| GPR139 | -0.4498834 | 1 | -1.16967 | 1 |
| BCL10 | -0.5924293 | 1 | -1.02301 | 1 |
| CREBZF | -0.9525399 | 1 | -0.6624 | 1 |
| VEGFA | -0.2963483 | 1 | -1.31545 | 1 |
| RAB8B | -0.6497104 | 1 | -0.9616 | 1 |
| RXFP1 | -0.131202 | 1 | -1.48 | 1 |

| | | | | |
|---|---|---|---|---|
| FANCF | -0.4763627 | 1 | -1.13222 | 1 |
| POU3F4 | -0.8828733 | 1 | -0.72467 | 1 |
| KDM3A | -0.2450006 | 1 | -1.36156 | 1 |
| PDCL | -1.1707986 | 1 | -0.43343 | 1 |
| SMARCA4 | -1.0854853 | 1 | -0.51829 | 1 |
| PCGF5 | -1.3764551 | 1 | -0.22662 | 1 |
| ZBTB11 | -0.9287197 | 1 | -0.67345 | 1 |
| TAS2R39 | 0.00548597 | 1 | -1.60727 | 1 |
| ZNF521 | -0.9530232 | 1 | -0.64613 | 1 |
| TRHDE | -1.3567598 | 1 | -0.24218 | 1 |
| BCORL1 | -1.4099522 | 1 | -0.1872 | 1 |
| TP53BP2 | -1.3730806 | 1 | -0.22309 | 1 |
| ZNF585B | -0.1831022 | 1 | -1.41287 | 1 |
| ZNF22 | -1.1403202 | 1 | -0.45137 | 1 |
| ZNF711 | -0.3051748 | 1 | -1.28183 | 1 |
| BRWD1 | -1.1351827 | 1 | -0.44993 | 1 |
| MN1 | -0.3816668 | 1 | -1.20299 | 1 |
| RPS6KA6 | -0.2240995 | 1 | -1.35945 | 1 |
| CDKN1B | -1.1441899 | 1 | -0.4356 | 1 |
| STAG2 | -0.8304402 | 1 | -0.74884 | 1 |
| PPP1R15B | -0.4279154 | 1 | -1.15088 | 1 |
| RAB2B | -0.2217936 | 1 | -1.35558 | 1 |
| ARHGEF38 | -0.9603224 | 1 | -0.6169 | 1 |
| MSL3 | -1.3243237 | 1 | -0.25245 | 1 |
| JUN | -0.3051589 | 1 | -1.27029 | 1 |
| RGS16 | -1.0552309 | 1 | -0.51917 | 1 |
| ELL2 | -1.1507401 | 1 | -0.42119 | 1 |
| EPO | -0.9098934 | 1 | -0.65808 | 1 |
| PRAME | -0.0071844 | 1 | -1.55972 | 1 |
| ZNF714 | -1.0417318 | 1 | -0.52501 | 1 |
| ZNF385B | -1.1351631 | 1 | -0.43098 | 1 |
| PDE4D | -0.9667191 | 1 | -0.59751 | 1 |
| SSBP3 | -0.8378995 | 1 | -0.72601 | 1 |
| TFRC | -1.1393817 | 1 | -0.42416 | 1 |
| ZDHHC23 | -0.9299224 | 1 | -0.63357 | 1 |
| ITGA6 | -1.4705409 | 1 | -0.092 | 1 |
| OR4C12 | -0.1462205 | 1 | -1.41531 | 1 |
| TOB1 | -1.281252 | 1 | -0.27959 | 1 |
| MTSS1 | -2.2171197 | 1 | 0.658147 | 1 |
| ECE1 | -0.9764151 | 1 | -0.58243 | 1 |
| RRAD | -1.8129534 | 1 | 0.254684 | 1 |
| SIM1 | -0.1471716 | 1 | -1.41077 | 1 |
| NFATC1 | -0.7135462 | 1 | -0.84283 | 1 |
| NFYB | -1.4766789 | 1 | -0.07769 | 1 |
| ADA | -1.277209 | 1 | -0.27591 | 1 |
| OR4N5 | -0.103424 | 1 | -1.44886 | 1 |

| | | | | |
|---|---|---|---|---|
| ROCK1 | -0.7388419 | 1 | -0.81305 | 1 |
| RAD50 | -1.0380864 | 1 | -0.51295 | 1 |
| ADAM10 | -1.0120818 | 1 | -0.53875 | 1 |
| PSD4 | -0.5200159 | 1 | -1.02911 | 1 |
| MAPK8 | -1.3188452 | 1 | -0.229 | 1 |
| PIGU | -1.0311823 | 1 | -0.51658 | 1 |
| CHRNB2 | -0.9832744 | 1 | -0.56415 | 1 |
| NBN | -0.9197111 | 1 | -0.62649 | 1 |
| LRP6 | -1.1214661 | 1 | -0.4224 | 1 |
| OR2C3 | -0.0145055 | 1 | -1.52924 | 1 |
| GPSM2 | -0.5702864 | 1 | -0.97134 | 1 |
| HIVEP2 | -0.6652889 | 1 | -0.87475 | 1 |
| ZNF391 | -1.0981484 | 1 | -0.44149 | 1 |
| YWHAZ | -0.3949506 | 1 | -1.14382 | 1 |
| NELFE | -1.0141378 | 1 | -0.52427 | 1 |
| ZNF25 | -0.3956513 | 1 | -1.14264 | 1 |
| PKDREJ | -1.5461053 | 1 | 0.008647 | 1 |
| RGS7 | -1.0774962 | 1 | -0.45962 | 1 |
| PSIP1 | -0.3063823 | 1 | -1.23032 | 1 |
| TOP1 | -0.9742747 | 1 | -0.5612 | 1 |
| ROR1 | -1.2253285 | 1 | -0.30871 | 1 |
| POGK | -0.1930377 | 1 | -1.33984 | 1 |
| CCNH | -0.8393063 | 1 | -0.69355 | 1 |
| ZNF239 | -0.9129171 | 1 | -0.61969 | 1 |
| CASK | -0.3692749 | 1 | -1.16088 | 1 |
| VIPR1 | -1.0895948 | 1 | -0.44032 | 1 |
| USP9X | -1.2073223 | 1 | -0.31853 | 1 |
| OR4N2 | -1.3672111 | 1 | -0.15845 | 1 |
| NR2F2 | -1.1371241 | 1 | -0.38768 | 1 |
| PNRC2 | -1.1542034 | 1 | -0.37055 | 1 |
| RGMB | -0.4361911 | 1 | -1.088 | 1 |
| ZNF395 | -1.1221585 | 1 | -0.40133 | 1 |
| TDG | -0.6540344 | 1 | -0.86733 | 1 |
| RASGEF1B | -0.7225812 | 1 | -0.79749 | 1 |
| EPS15 | -1.2698616 | 1 | -0.25003 | 1 |
| TFDP2 | -0.3859243 | 1 | -1.13152 | 1 |
| BRCC3 | -0.7442351 | 1 | -0.77312 | 1 |
| ZNF672 | -1.5858544 | 1 | 0.071722 | 1 |
| MRE11 | -0.3848303 | 1 | -1.12786 | 1 |
| BMP2 | -1.2788922 | 1 | -0.23184 | 1 |
| OR56B1 | -2.6137308 | 1 | 1.109053 | 1 |
| ITGB8 | -0.4587369 | 1 | -1.04527 | 1 |
| PDE4B | -0.9983866 | 1 | -0.50445 | 1 |
| RASAL2 | -0.1878563 | 1 | -1.31418 | 1 |
| NEK11 | -0.1682647 | 1 | -1.3332 | 1 |
| GUCY1B1 | -0.2256557 | 1 | -1.27264 | 1 |

| | | | | |
|---|---|---|---|---|
| ARHGAP32 | -1.434582 | 1 | -0.06187 | 1 |
| PKNOX2 | -0.5417699 | 1 | -0.95058 | 1 |
| MBD3L3 | -1.1934555 | 1 | -0.29826 | 1 |
| GPR33 | -0.0857744 | 1 | -1.40446 | 1 |
| PLXNC1 | -0.9348817 | 1 | -0.55171 | 1 |
| VPS36 | -0.0950782 | 1 | -1.39125 | 1 |
| SMC1A | -0.5076316 | 1 | -0.97866 | 1 |
| RALGAPA1 | -0.5809909 | 1 | -0.90058 | 1 |
| IPO8 | -1.0883194 | 1 | -0.39144 | 1 |
| UBR5 | -0.7097866 | 1 | -0.76916 | 1 |
| CELSR1 | -0.9429612 | 1 | -0.53554 | 1 |
| ING3 | -1.0562346 | 1 | -0.42166 | 1 |
| ZNF350 | -0.9504986 | 1 | -0.52571 | 1 |
| MED4 | 0.26912205 | 1 | -1.74427 | 1 |
| CREB5 | -1.2333004 | 1 | -0.24174 | 1 |
| KLF4 | -0.96136 | 1 | -0.50822 | 1 |
| RHOT1 | -0.8529249 | 1 | -0.61577 | 1 |
| NRG3 | -1.6602499 | 1 | 0.191944 | 1 |
| OR4M1 | -0.2419459 | 1 | -1.22416 | 1 |
| UBE2L3 | 0.67511477 | 1 | -2.13954 | 1 |
| ZBED4 | -1.0610571 | 1 | -0.40113 | 1 |
| HGF | -0.6086056 | 1 | -0.85331 | 1 |
| ZNF664 | -0.4127546 | 1 | -1.03935 | 1 |
| ERO1A | -0.2361423 | 1 | -1.21243 | 1 |
| ZNF85 | -0.8516919 | 1 | -0.5918 | 1 |
| ZNF843 | -1.561745 | 1 | 0.120535 | 1 |
| MED30 | -1.1853305 | 1 | -0.25514 | 1 |
| RGS12 | -1.299731 | 1 | -0.13968 | 1 |
| KLF3 | -1.1867282 | 1 | -0.24796 | 1 |
| MXD1 | -1.033637 | 1 | -0.39975 | 1 |
| APLNR | -0.9445136 | 1 | -0.4878 | 1 |
| FOXO6 | -1.168861 | 1 | -0.26141 | 1 |
| ZNF669 | -0.3858158 | 1 | -1.04443 | 1 |
| EPHB3 | -0.5689897 | 1 | -0.85976 | 1 |
| ROBO1 | -0.0892633 | 1 | -1.33866 | 1 |
| KAT6A | -0.9278535 | 1 | -0.49951 | 1 |
| ELP3 | -0.4380811 | 1 | -0.98637 | 1 |
| ZMYM4 | -0.9504962 | 1 | -0.47261 | 1 |
| ZBTB39 | -1.2888766 | 1 | -0.13284 | 1 |
| FCGR2B | -0.3451961 | 1 | -1.07522 | 1 |
| MCHR1 | -1.2363375 | 1 | -0.18185 | 1 |
| PRKCA | -0.0587742 | 1 | -1.35662 | 1 |
| HOMER1 | -1.2259994 | 1 | -0.18839 | 1 |
| GATA4 | -0.902007 | 1 | -0.51224 | 1 |
| PAK3 | -0.8033345 | 1 | -0.60923 | 1 |
| GATA5 | -0.0129654 | 1 | -1.39882 | 1 |

| | | | | |
|---|---|---|---|---|
| TGFB1I1 | -1.0942802 | 1 | -0.31696 | 1 |
| ZFYVE9 | -0.5481584 | 1 | -0.85982 | 1 |
| ASH1L | -1.459426 | 1 | 0.052739 | 1 |
| ZNF184 | -0.9591203 | 1 | -0.44733 | 1 |
| HOXA3 | -0.2730284 | 1 | -1.13084 | 1 |
| ADRA1A | -0.2684404 | 1 | -1.13407 | 1 |
| ZEB1 | -1.0424981 | 1 | -0.35983 | 1 |
| INHBA | -0.7994478 | 1 | -0.60155 | 1 |
| ASAP3 | -0.1640329 | 1 | -1.23695 | 1 |
| ACKR4 | -0.6912985 | 1 | -0.70372 | 1 |
| HNF4A | -0.4748775 | 1 | -0.91944 | 1 |
| ARNTL2 | -1.0972599 | 1 | -0.29304 | 1 |
| TBC1D14 | -1.4371197 | 1 | 0.04712 | 1 |
| PPP3CB | -1.0729965 | 1 | -0.31455 | 1 |
| PDGFB | -0.477803 | 1 | -0.90799 | 1 |
| LBX1 | -0.0662746 | 1 | -1.31864 | 1 |
| ZNF326 | -1.0775194 | 1 | -0.30735 | 1 |
| MED7 | -1.1223533 | 1 | -0.2617 | 1 |
| ZNF117 | -0.5225385 | 1 | -0.86129 | 1 |
| PTGER4 | -0.030721 | 1 | -1.3528 | 1 |
| IL17RA | -0.0940161 | 1 | -1.28889 | 1 |
| RSF1 | -0.8224749 | 1 | -0.55953 | 1 |
| POU4F1 | -1.3436391 | 1 | -0.03536 | 1 |
| NKD1 | -0.4882752 | 1 | -0.89055 | 1 |
| HEYL | -0.8290834 | 1 | -0.54974 | 1 |
| INPP1 | -2.0606426 | 1 | 0.684401 | 1 |
| HLTF | -0.5042713 | 1 | -0.87194 | 1 |
| MAP3K1 | -0.97801 | 1 | -0.39723 | 1 |
| OPN1SW | -0.5495292 | 1 | -0.82143 | 1 |
| SOS1 | -0.4764387 | 1 | -0.89448 | 1 |
| OR2W1 | -0.0021879 | 1 | -1.36729 | 1 |
| PTPRM | -0.6784462 | 1 | -0.68828 | 1 |
| JAZF1 | -1.1275948 | 1 | -0.23896 | 1 |
| H4C13 | -0.9539136 | 1 | -0.40944 | 1 |
| OR10G6 | -0.6590605 | 1 | -0.70367 | 1 |
| SNAPC1 | -0.6489374 | 1 | -0.71115 | 1 |
| OR2G6 | -0.2525453 | 1 | -1.10456 | 1 |
| CCL24 | -0.7688217 | 1 | -0.58617 | 1 |
| BCOR | -0.843349 | 1 | -0.51103 | 1 |
| CHRM1 | -1.1928978 | 1 | -0.16082 | 1 |
| RHOBTB2 | -0.961592 | 1 | -0.38958 | 1 |
| ZNF93 | -0.7738594 | 1 | -0.57399 | 1 |
| DENND4A | -0.3815535 | 1 | -0.96468 | 1 |
| CPS1 | -0.2982512 | 1 | -1.04788 | 1 |
| SOX6 | -1.2793142 | 1 | -0.06514 | 1 |
| OR4C6 | 0.00993873 | 1 | -1.35404 | 1 |

| | | | | |
|---|---|---|---|---|
| STK38 | -1.1993585 | 1 | -0.14419 | 1 |
| ZNF518A | -1.0439033 | 1 | -0.29722 | 1 |
| GAPVD1 | -0.8471906 | 1 | -0.49366 | 1 |
| WDR12 | -0.7796356 | 1 | -0.554 | 1 |
| CA8 | -0.4605113 | 1 | -0.87299 | 1 |
| ZNF652 | -1.2481523 | 1 | -0.08299 | 1 |
| ARHGEF7 | -1.2976429 | 1 | -0.03233 | 1 |
| SF1 | -0.1130017 | 1 | -1.21473 | 1 |
| CBX8 | -0.1535462 | 1 | -1.17407 | 1 |
| MORF4L1 | -0.5993465 | 1 | -0.72814 | 1 |
| NR3C1 | -0.620579 | 1 | -0.70647 | 1 |
| PTGES3 | -0.6613629 | 1 | -0.66568 | 1 |
| CD69 | -0.2298979 | 1 | -1.09697 | 1 |
| ATR | -1.1559146 | 1 | -0.17049 | 1 |
| PTPRT | -0.3479022 | 1 | -0.97838 | 1 |
| DUSP16 | -0.2123926 | 1 | -1.11387 | 1 |
| UBE2D3 | 0.48006371 | 1 | -1.80442 | 1 |
| DIXDC1 | -2.4279324 | 1 | 1.103837 | 1 |
| H4C4 | -0.9823044 | 1 | -0.34104 | 1 |
| OR5J2 | -0.0656509 | 1 | -1.25667 | 1 |
| MAFF | -0.2857789 | 1 | -1.03547 | 1 |
| CNOT2 | -0.8727102 | 1 | -0.44633 | 1 |
| CNIH1 | -0.1289863 | 1 | -1.1899 | 1 |
| OR10J5 | -0.9339483 | 1 | -0.38159 | 1 |
| C1D | 0.0426632 | 1 | -1.35701 | 1 |
| ADORA2B | -0.0913608 | 1 | -1.22268 | 1 |
| H4C12 | -1.040476 | 1 | -0.27126 | 1 |
| PDE7A | -0.5801024 | 1 | -0.72952 | 1 |
| LMCD1 | -0.0900769 | 1 | -1.21163 | 1 |
| IRAK2 | -0.127612 | 1 | -1.17307 | 1 |
| GTF2H2C_2 | -0.9921491 | 1 | -0.30752 | 1 |
| PRKCQ | -1.7445642 | 1 | 0.445624 | 1 |
| BRMS1L | -0.1364508 | 1 | -1.16206 | 1 |
| USP8 | -0.9176189 | 1 | -0.37947 | 1 |
| OR9I1 | -0.3644256 | 1 | -0.93212 | 1 |
| BIRC2 | -0.7880264 | 1 | -0.50836 | 1 |
| TCF15 | -0.569741 | 1 | -0.72573 | 1 |
| TRIM63 | -0.4855466 | 1 | -0.8095 | 1 |
| ZKSCAN4 | -0.746593 | 1 | -0.54813 | 1 |
| ZNF234 | -0.5265004 | 1 | -0.76774 | 1 |
| RHOV | -0.0980101 | 1 | -1.19523 | 1 |
| ZBTB33 | -0.6144284 | 1 | -0.67494 | 1 |
| ZFP62 | -0.9867905 | 1 | -0.30229 | 1 |
| TRPS1 | 0.22724759 | 1 | -1.51482 | 1 |
| OR10A5 | -1.1687484 | 1 | -0.11872 | 1 |
| ZNF486 | -0.5286825 | 1 | -0.75846 | 1 |

| | | | | |
|---|---|---|---|---|
| DNER | -1.1520631 | 1 | -0.13389 | 1 |
| CYSLTR2 | -0.6243029 | 1 | -0.66147 | 1 |
| SOX9 | -0.0524675 | 1 | -1.23321 | 1 |
| ZNF781 | -0.7322624 | 1 | -0.55195 | 1 |
| ARHGEF3 | -0.8960143 | 1 | -0.38781 | 1 |
| PDE5A | -0.9291016 | 1 | -0.35419 | 1 |
| ZNF501 | -0.4242012 | 1 | -0.85713 | 1 |
| VEZF1 | -0.8903564 | 1 | -0.39026 | 1 |
| CBX3 | -0.3263962 | 1 | -0.95311 | 1 |
| OR2A42 | -0.8668387 | 1 | -0.40693 | 1 |
| ZFHX3 | -1.3339265 | 1 | 0.064098 | 1 |
| SMAD2 | 0.11765196 | 1 | -1.38613 | 1 |
| GABRA6 | -0.1467728 | 1 | -1.11795 | 1 |
| OR8I2 | -0.09255 | 1 | -1.16813 | 1 |
| RBAK | -0.4139952 | 1 | -0.84321 | 1 |
| MCL1 | -0.2144573 | 1 | -1.04266 | 1 |
| NAB1 | -0.350036 | 1 | -0.89915 | 1 |
| MAML1 | -0.9041201 | 1 | -0.3449 | 1 |
| STMN2 | -0.5665052 | 1 | -0.68224 | 1 |
| MAPK1 | 0.0469483 | 1 | -1.29354 | 1 |
| RAB33B | -0.2294123 | 1 | -1.01622 | 1 |
| ZBTB44 | -0.5540287 | 1 | -0.69148 | 1 |
| OR4C16 | -0.682525 | 1 | -0.5629 | 1 |
| POU5F1 | -0.2852381 | 1 | -0.95847 | 1 |
| ABI1 | -0.0246102 | 1 | -1.21787 | 1 |
| DUSP10 | -0.4135672 | 1 | -0.82788 | 1 |
| RNF141 | -1.6905066 | 1 | 0.450564 | 1 |
| TBX6 | -1.447864 | 1 | 0.208367 | 1 |
| ZNF26 | -1.3665539 | 1 | 0.129274 | 1 |
| AKT3 | -1.1136667 | 1 | -0.12309 | 1 |
| TSPYL2 | -1.2549038 | 1 | 0.018305 | 1 |
| TIAM2 | -0.5362089 | 1 | -0.70032 | 1 |
| HEY2 | -0.7830989 | 1 | -0.45341 | 1 |
| ZNF716 | 0.20418665 | 1 | -1.43988 | 1 |
| UHRF1 | -0.9308871 | 1 | -0.30269 | 1 |
| OR2AT4 | -0.847267 | 1 | -0.38516 | 1 |
| PRKAA1 | -0.3868289 | 1 | -0.84353 | 1 |
| ZNF804B | -1.0638148 | 1 | -0.16092 | 1 |
| CCT6B | -0.9802653 | 1 | -0.24429 | 1 |
| ITGB3 | 1.29089984 | 1 | -2.51507 | 1 |
| MAX | 0.13659886 | 1 | -1.36076 | 1 |
| LZTFL1 | -0.7736927 | 1 | -0.44818 | 1 |
| ADGRB1 | -1.2764996 | 1 | 0.055859 | 1 |
| ELN | -0.8964736 | 1 | -0.32329 | 1 |
| OR6B1 | -1.0254422 | 1 | -0.19333 | 1 |
| MKLN1 | -0.7680531 | 1 | -0.44947 | 1 |

| | | | | |
|---|---|---|---|---|
| TAS2R45 | -0.7811748 | 1 | -0.43551 | 1 |
| PRDM7 | -1.4978 | 1 | 0.28293 | 1 |
| KIR2DS2 | -1.1042129 | 1 | -0.10968 | 1 |
| CELSR3 | -0.4635935 | 1 | -0.74732 | 1 |
| TMX1 | -0.6895225 | 1 | -0.51971 | 1 |
| OR6F1 | -1.2319789 | 1 | 0.024118 | 1 |
| CD44 | -0.7708838 | 1 | -0.43653 | 1 |
| ADGRG6 | -1.142586 | 1 | -0.06294 | 1 |
| UBC | -0.9500702 | 1 | -0.25284 | 1 |
| ZNF443 | -0.0395153 | 1 | -1.16316 | 1 |
| PDGFRA | -1.193176 | 1 | -0.00738 | 1 |
| GABRA5 | -1.0979041 | 1 | -0.10221 | 1 |
| TAF1B | -0.0553815 | 1 | -1.14057 | 1 |
| ZNF417 | -0.208697 | 1 | -0.98707 | 1 |
| PAK2 | -0.4599215 | 1 | -0.73581 | 1 |
| NLE1 | -0.9319231 | 1 | -0.26378 | 1 |
| RXRG | -1.2507319 | 1 | 0.056019 | 1 |
| FZD2 | -1.0116549 | 1 | -0.18304 | 1 |
| EPHA8 | -0.0984971 | 1 | -1.09441 | 1 |
| OR6C1 | -0.5784123 | 1 | -0.61327 | 1 |
| DLL1 | -1.0866856 | 1 | -0.10493 | 1 |
| ZNF530 | -0.4570583 | 1 | -0.7344 | 1 |
| SCAI | -0.4704769 | 1 | -0.72077 | 1 |
| ZNF285 | -1.3304311 | 1 | 0.140167 | 1 |
| RHOA | -0.8135173 | 1 | -0.37514 | 1 |
| PCBD2 | -0.4118937 | 1 | -0.77553 | 1 |
| CRYM | -1.2117315 | 1 | 0.025154 | 1 |
| SERTAD2 | -0.5598265 | 1 | -0.62655 | 1 |
| GNA13 | -0.7289296 | 1 | -0.45567 | 1 |
| OR10K1 | 0.1010946 | 1 | -1.28567 | 1 |
| HMGN5 | -0.7319501 | 1 | -0.45235 | 1 |
| KHSRP | -0.950669 | 1 | -0.23103 | 1 |
| OR3A1 | -0.7224859 | 1 | -0.4592 | 1 |
| RPS6KC1 | -0.3074836 | 1 | -0.87373 | 1 |
| TBC1D8B | -0.9276305 | 1 | -0.25331 | 1 |
| OR52N4 | -0.7541662 | 1 | -0.42238 | 1 |
| TP73 | 0.01272047 | 1 | -1.18898 | 1 |
| AGTR1 | -0.3950251 | 1 | -0.78086 | 1 |
| ZNF563 | -0.630457 | 1 | -0.54512 | 1 |
| NEUROD6 | -0.2071922 | 1 | -0.96817 | 1 |
| ZNF614 | -0.265503 | 1 | -0.90976 | 1 |
| ZSCAN12 | -0.708252 | 1 | -0.46542 | 1 |
| ALX3 | -0.7764906 | 1 | -0.39455 | 1 |
| UBE2D1 | -0.8782619 | 1 | -0.29257 | 1 |
| NECTIN2 | -0.0613562 | 1 | -1.10756 | 1 |
| PALB2 | -0.5169659 | 1 | -0.65111 | 1 |

| | | | | |
|---|---|---|---|---|
| SLC30A9 | -0.5099534 | 1 | -0.65812 | 1 |
| TULP4 | -0.9376092 | 1 | -0.22975 | 1 |
| DAB2IP | -0.149764 | 1 | -1.01624 | 1 |
| CCNT1 | -0.1889061 | 1 | -0.97704 | 1 |
| DRD1 | -0.65508 | 1 | -0.51045 | 1 |
| PTPRO | -0.4505807 | 1 | -0.7137 | 1 |
| FABP4 | -1.1707287 | 1 | 0.006479 | 1 |
| F2RL3 | -0.0501838 | 1 | -1.11312 | 1 |
| CCL2 | -0.562499 | 1 | -0.59917 | 1 |
| ZNF609 | -0.8932434 | 1 | -0.26555 | 1 |
| ZNF682 | -0.9647055 | 1 | -0.19339 | 1 |
| MYO6 | -1.6387744 | 1 | 0.483058 | 1 |
| VAV2 | -0.6984724 | 1 | -0.45595 | 1 |
| OR11A1 | -0.1862842 | 1 | -0.96806 | 1 |
| RBBP7 | -0.348702 | 1 | -0.80549 | 1 |
| PRKAB1 | -0.069549 | 1 | -1.08416 | 1 |
| RALGAPA2 | -0.723243 | 1 | -0.43036 | 1 |
| ADCY3 | -0.9548504 | 1 | -0.19845 | 1 |
| CRTC3 | -1.29543 | 1 | 0.142276 | 1 |
| OR7G2 | -1.0487785 | 1 | -0.1009 | 1 |
| SPRED1 | -0.6811759 | 1 | -0.46848 | 1 |
| ARFGAP3 | -0.4150325 | 1 | -0.73426 | 1 |
| RAD21 | -0.3557775 | 1 | -0.79242 | 1 |
| DGKH | -0.5168237 | 1 | -0.63073 | 1 |
| NPY2R | -0.5174394 | 1 | -0.62948 | 1 |
| MARK2 | -1.2302349 | 1 | 0.084343 | 1 |
| ADGRL2 | -0.1655903 | 1 | -0.97671 | 1 |
| CHD7 | -0.9743745 | 1 | -0.16744 | 1 |
| GRIP1 | -0.8329014 | 1 | -0.30795 | 1 |
| LOXL2 | -0.846195 | 1 | -0.29394 | 1 |
| TMED4 | -0.7165345 | 1 | -0.4233 | 1 |
| BCLAF1 | -0.2243412 | 1 | -0.91521 | 1 |
| PURB | -0.6157154 | 1 | -0.52347 | 1 |
| SCX | -1.1734494 | 1 | 0.03548 | 1 |
| ZKSCAN8 | -0.5411443 | 1 | -0.59659 | 1 |
| HESX1 | -0.0417478 | 1 | -1.09599 | 1 |
| ZBTB43 | -0.2069661 | 1 | -0.93067 | 1 |
| IL1RL2 | -0.1563785 | 1 | -0.98107 | 1 |
| CALM2 | 0.08888079 | 1 | -1.22281 | 1 |
| TAOK1 | -0.5231266 | 1 | -0.6094 | 1 |
| CBL | -0.1128331 | 1 | -1.0186 | 1 |
| IRX6 | -0.9069471 | 1 | -0.22261 | 1 |
| HNRNPUL1 | -0.8232626 | 1 | -0.30621 | 1 |
| OXSR1 | -0.269129 | 1 | -0.85945 | 1 |
| RC3H1 | 0.09004039 | 1 | -1.21674 | 1 |
| TACR1 | -1.0088288 | 1 | -0.11704 | 1 |

| | | | | |
|---|---|---|---|---|
| FADD | -1.0185835 | 1 | -0.10706 | 1 |
| CDC42SE1 | -0.6963474 | 1 | -0.42881 | 1 |
| FLCN | -0.1293299 | 1 | -0.99563 | 1 |
| AFF1 | -0.1101876 | 1 | -1.01167 | 1 |
| OR51I2 | -0.281467 | 1 | -0.83884 | 1 |
| ZNF610 | -0.7680514 | 1 | -0.35041 | 1 |
| PTPN14 | -0.7707202 | 1 | -0.34737 | 1 |
| MAML2 | -2.2041621 | 1 | 1.086523 | 1 |
| LRP8 | -0.2506595 | 1 | -0.86534 | 1 |
| EGFR | -0.2010568 | 1 | -0.91429 | 1 |
| MARK1 | -0.7322898 | 1 | -0.38305 | 1 |
| ZKSCAN1 | -0.3845936 | 1 | -0.72927 | 1 |
| OR2T5 | -0.1012236 | 1 | -1.01255 | 1 |
| ZNF708 | -0.859442 | 1 | -0.25344 | 1 |
| RSC1A1 | -0.016981 | 1 | -1.09528 | 1 |
| IDH1 | -0.4598392 | 1 | -0.65009 | 1 |
| IRF2BP2 | -0.3055881 | 1 | -0.80188 | 1 |
| SIX3 | -1.3856138 | 1 | 0.279385 | 1 |
| CHN2 | -2.1741355 | 1 | 1.069334 | 1 |
| ZNF202 | -0.8082941 | 1 | -0.29576 | 1 |
| OR2C1 | -0.3776697 | 1 | -0.72558 | 1 |
| FGFBP1 | -0.7844275 | 1 | -0.31499 | 1 |
| TMED7.TICAM2 | -0.7820732 | 1 | -0.3167 | 1 |
| LEMD3 | -0.5865139 | 1 | -0.51189 | 1 |
| FOXK1 | -0.9565936 | 1 | -0.13968 | 1 |
| H2BC12 | -0.5421297 | 1 | -0.55187 | 1 |
| DNMT3A | -0.3371883 | 1 | -0.7567 | 1 |
| CEP290 | -0.8475934 | 1 | -0.24582 | 1 |
| ZNF101 | -0.4800114 | 1 | -0.61217 | 1 |
| SETD2 | -0.7624868 | 1 | -0.32879 | 1 |
| ACTL6B | -0.7549458 | 1 | -0.33389 | 1 |
| CACNB4 | -0.1562608 | 1 | -0.93249 | 1 |
| RPS6KA2 | -0.9656159 | 1 | -0.12207 | 1 |
| RINT1 | -0.2218231 | 1 | -0.86579 | 1 |
| PIK3CB | -0.2102439 | 1 | -0.87659 | 1 |
| HES7 | -1.317918 | 1 | 0.231292 | 1 |
| CBFA2T2 | 0.05678352 | 1 | -1.14328 | 1 |
| ARL3 | -0.1365798 | 1 | -0.94966 | 1 |
| NPFFR2 | -1.3687256 | 1 | 0.28273 | 1 |
| ZBTB14 | -1.0059771 | 1 | -0.07936 | 1 |
| HTR1B | -0.2914221 | 1 | -0.79176 | 1 |
| PLEKHG2 | -0.4524385 | 1 | -0.63012 | 1 |
| GPBP1 | -0.777183 | 1 | -0.30458 | 1 |
| ZNF468 | -0.7473976 | 1 | -0.33422 | 1 |
| GPHN | 0.2225834 | 1 | -1.30413 | 1 |
| L1CAM | -0.5255912 | 1 | -0.55417 | 1 |

| | | | | |
|---|---|---|---|---|
| OR4F4 | -0.246669 | 1 | -0.83205 | 1 |
| OR2M4 | -0.5537914 | 1 | -0.52479 | 1 |
| NFIL3 | -0.2317681 | 1 | -0.84471 | 1 |
| TNFRSF14 | -1.0744725 | 1 | -0.00171 | 1 |
| EBF2 | -0.1465246 | 1 | -0.92826 | 1 |
| ZNF148 | -0.326529 | 1 | -0.74697 | 1 |
| REL | -0.1507521 | 1 | -0.92111 | 1 |
| MSN | -0.2944308 | 1 | -0.7761 | 1 |
| ROS1 | -0.7668244 | 1 | -0.30257 | 1 |
| MDM2 | -0.5262434 | 1 | -0.54137 | 1 |
| NFIA | -0.1788093 | 1 | -0.88848 | 1 |
| OR4F17 | -0.2529996 | 1 | -0.81335 | 1 |
| ESR2 | 1.22783422 | 1 | -2.29367 | 1 |
| FRS2 | -0.330672 | 1 | -0.73354 | 1 |
| TPRX1 | 0.00902898 | 1 | -1.07296 | 1 |
| PLEKHG4B | -0.3572006 | 1 | -0.70362 | 1 |
| FOXN3 | -0.1805372 | 1 | -0.87894 | 1 |
| GTF2E2 | -0.0849716 | 1 | -0.97416 | 1 |
| LRRK2 | -0.970614 | 1 | -0.08788 | 1 |
| ZCCHC4 | -0.6679642 | 1 | -0.39036 | 1 |
| CNKSR2 | -0.6490495 | 1 | -0.40899 | 1 |
| MED6 | -0.2129415 | 1 | -0.84321 | 1 |
| GPR63 | -0.8939505 | 1 | -0.16079 | 1 |
| CAMKK2 | -2.0895621 | 1 | 1.035047 | 1 |
| PLCXD2 | -0.8035946 | 1 | -0.2508 | 1 |
| OR5M8 | -0.6691778 | 1 | -0.38514 | 1 |
| ANG | -0.2034133 | 1 | -0.84415 | 1 |
| ZNF677 | -0.1523631 | 1 | -0.89506 | 1 |
| RAB3C | -0.894846 | 1 | -0.15199 | 1 |
| RSPO1 | -0.0996514 | 1 | -0.94638 | 1 |
| ZBTB2 | -0.6516899 | 1 | -0.39406 | 1 |
| SPRY2 | -0.7561507 | 1 | -0.2889 | 1 |
| PEX11A | -0.678729 | 1 | -0.36582 | 1 |
| ARID2 | -0.5848892 | 1 | -0.45861 | 1 |
| ZNF277 | -0.8062263 | 1 | -0.2372 | 1 |
| IL4 | 0.05050617 | 1 | -1.09359 | 1 |
| SUPT3H | -0.2766862 | 1 | -0.7658 | 1 |
| ATG5 | 0.89757228 | 1 | -1.93922 | 1 |
| DUSP2 | -0.23082 | 1 | -0.81072 | 1 |
| RAP2B | -0.8000861 | 1 | -0.24011 | 1 |
| ZNF140 | -0.6668299 | 1 | -0.37326 | 1 |
| ATP2C1 | -0.5452385 | 1 | -0.49307 | 1 |
| MAP2K4 | -1.3142335 | 1 | 0.277047 | 1 |
| TBC1D19 | -0.4664659 | 1 | -0.56989 | 1 |
| IQSEC3 | -0.8380039 | 1 | -0.19835 | 1 |
| ZNF445 | -0.4084944 | 1 | -0.62764 | 1 |

| | | | | |
|---|---|---|---|---|
| VLDLR | -0.1112511 | 1 | -0.9229 | 1 |
| NRIP1 | -0.6690198 | 1 | -0.36441 | 1 |
| E2F5 | -0.5965127 | 1 | -0.43548 | 1 |
| KL | -0.5644829 | 1 | -0.46694 | 1 |
| FOXP1 | -0.8126039 | 1 | -0.21861 | 1 |
| GMCL1 | -0.1908867 | 1 | -0.84014 | 1 |
| APPL1 | -0.5390654 | 1 | -0.49186 | 1 |
| KMT2D | -1.7059125 | 1 | 0.675131 | 1 |
| BCL6B | -1.0057925 | 1 | -0.02477 | 1 |
| SFMBT1 | -0.4969386 | 1 | -0.53226 | 1 |
| ZNF474 | -0.0571286 | 1 | -0.96922 | 1 |
| CAMTA1 | -0.9514119 | 1 | -0.0718 | 1 |
| PTPRA | -0.8150466 | 1 | -0.20732 | 1 |
| HOXC8 | -0.0194362 | 1 | -1.00289 | 1 |
| HOXD10 | -1.0745112 | 1 | 0.0525 | 1 |
| OR2M3 | -0.6265193 | 1 | -0.39538 | 1 |
| KLRC3 | -0.9192135 | 1 | -0.10138 | 1 |
| BARD1 | -0.5843244 | 1 | -0.43575 | 1 |
| FANCC | -0.6471421 | 1 | -0.37237 | 1 |
| CBX4 | -0.8558815 | 1 | -0.16329 | 1 |
| MBTD1 | -0.2246572 | 1 | -0.79312 | 1 |
| PPM1L | -0.2790107 | 1 | -0.73737 | 1 |
| GEM | -0.8522584 | 1 | -0.16397 | 1 |
| ZNF557 | -0.7070671 | 1 | -0.30811 | 1 |
| ZNF860 | -0.1957334 | 1 | -0.81826 | 1 |
| FOXA1 | -0.3403281 | 1 | -0.67298 | 1 |
| MAPK8IP2 | -1.1310127 | 1 | 0.117769 | 1 |
| ZNF675 | -0.6544621 | 1 | -0.35787 | 1 |
| YEATS2 | -0.3602023 | 1 | -0.65205 | 1 |
| PITPNM1 | -1.0793664 | 1 | 0.068432 | 1 |
| PRDM5 | -0.3763622 | 1 | -0.63356 | 1 |
| IKZF2 | -0.3322917 | 1 | -0.67706 | 1 |
| OR10AG1 | -0.9232294 | 1 | -0.08589 | 1 |
| ZNF774 | -0.925834 | 1 | -0.07954 | 1 |
| TNFRSF1A | 1.01434655 | 1 | -2.01971 | 1 |
| OR4N4 | -1.0179971 | 1 | 0.014477 | 1 |
| RHPN2 | -0.8075789 | 1 | -0.19582 | 1 |
| NME1.NME2 | -0.2099909 | 1 | -0.79293 | 1 |
| GABRR3 | -0.6540321 | 1 | -0.34755 | 1 |
| ACVRL1 | -0.3950653 | 1 | -0.60436 | 1 |
| MAPK13 | -0.3251568 | 1 | -0.67346 | 1 |
| HIPK3 | 0.01354665 | 1 | -1.01162 | 1 |
| LRP5 | -0.1777701 | 1 | -0.81954 | 1 |
| APLP2 | -0.8443223 | 1 | -0.15275 | 1 |
| TRPC3 | -0.150149 | 1 | -0.84553 | 1 |
| ZNF620 | -0.347504 | 1 | -0.64798 | 1 |

| | | | | |
|---|---|---|---|---|
| NT5C2 | -0.2163563 | 1 | -0.77865 | 1 |
| SRPK2 | -0.2395607 | 1 | -0.75473 | 1 |
| FRYL | -0.9158309 | 1 | -0.07827 | 1 |
| ZNF81 | -0.9180352 | 1 | -0.07413 | 1 |
| NKD2 | -0.8490792 | 1 | -0.14299 | 1 |
| ATF7IP2 | -0.8231852 | 1 | -0.16836 | 1 |
| ZNF263 | -0.859215 | 1 | -0.13167 | 1 |
| H4C15 | -0.0646375 | 1 | -0.92484 | 1 |
| RAP2C | -0.3287486 | 1 | -0.66048 | 1 |
| TPD52L1 | 0.10960061 | 1 | -1.09855 | 1 |
| KLF13 | -1.5514684 | 1 | 0.563988 | 1 |
| TFDP3 | -0.7406503 | 1 | -0.2468 | 1 |
| DST | -0.8327177 | 1 | -0.1527 | 1 |
| RASSF1 | -0.2716079 | 1 | -0.71334 | 1 |
| ITGB1 | -0.0317432 | 1 | -0.95181 | 1 |
| DISP1 | -0.7324721 | 1 | -0.24691 | 1 |
| SF3B1 | -0.1645923 | 1 | -0.81316 | 1 |
| ITLN1 | -0.9202597 | 1 | -0.05683 | 1 |
| MYCBP2 | -0.4573243 | 1 | -0.51889 | 1 |
| OR52N5 | -0.3257764 | 1 | -0.65042 | 1 |
| SKIL | -0.6564493 | 1 | -0.3168 | 1 |
| GUCA1C | -0.0777823 | 1 | -0.89268 | 1 |
| MCF2 | -0.315812 | 1 | -0.65439 | 1 |
| TBC1D23 | -0.1470309 | 1 | -0.82211 | 1 |
| ZNF492 | -0.4896088 | 1 | -0.47924 | 1 |
| NPBWR1 | -0.1293343 | 1 | -0.83915 | 1 |
| TUT7 | -0.2384217 | 1 | -0.72993 | 1 |
| KDM4D | -0.6841687 | 1 | -0.28405 | 1 |
| ZRANB2 | -0.2858755 | 1 | -0.6813 | 1 |
| AKAP4 | -0.4207419 | 1 | -0.54543 | 1 |
| NGF | -0.2047116 | 1 | -0.75847 | 1 |
| MAPKAPK3 | -0.2217103 | 1 | -0.74111 | 1 |
| GABRG3 | -0.6995498 | 1 | -0.26283 | 1 |
| ASB15 | -0.0741363 | 1 | -0.88731 | 1 |
| DDX6 | -0.3628762 | 1 | -0.59833 | 1 |
| DTX1 | -0.9150773 | 1 | -0.04566 | 1 |
| RPH3AL | -1.4214568 | 1 | 0.461772 | 1 |
| CDKN2AIP | -0.3605992 | 1 | -0.59876 | 1 |
| ARHGAP19 | -0.0085831 | 1 | -0.95072 | 1 |
| CAV2 | -0.8417857 | 1 | -0.11524 | 1 |
| CHURC1 | -0.7297187 | 1 | -0.22594 | 1 |
| FERD3L | -0.5423032 | 1 | -0.41297 | 1 |
| MALT1 | -0.3067716 | 1 | -0.6468 | 1 |
| SCARB1 | -1.2375681 | 1 | 0.285944 | 1 |
| HLA.DOB | -0.3094286 | 1 | -0.64213 | 1 |
| SH3GL2 | -1.0509518 | 1 | 0.101941 | 1 |

| | | | | |
|---|---|---|---|---|
| ARHGAP29 | -0.491848 | 1 | -0.45714 | 1 |
| MATK | -1.6267647 | 1 | 0.679092 | 1 |
| CD79B | -0.1618724 | 1 | -0.78555 | 1 |
| AEN | -0.7371889 | 1 | -0.20931 | 1 |
| ACTL6A | -0.1803532 | 1 | -0.76597 | 1 |
| XCR1 | -0.7155835 | 1 | -0.22921 | 1 |
| RABGAP1 | -0.1200273 | 1 | -0.82428 | 1 |
| ZMYM2 | -0.3687555 | 1 | -0.57416 | 1 |
| ASB12 | -0.1055247 | 1 | -0.83683 | 1 |
| SUB1 | -0.3337639 | 1 | -0.6085 | 1 |
| ZNF141 | -0.3303458 | 1 | -0.61007 | 1 |
| MITF | -0.7104308 | 1 | -0.22988 | 1 |
| ZNF578 | -0.6629786 | 1 | -0.2763 | 1 |
| ADCY5 | -0.8858856 | 1 | -0.05231 | 1 |
| RAB32 | -0.9989796 | 1 | 0.062097 | 1 |
| RUNX1T1 | -0.2551623 | 1 | -0.68156 | 1 |
| IL17F | -0.2490238 | 1 | -0.68756 | 1 |
| NPAS3 | -0.9175071 | 1 | -0.0169 | 1 |
| IL1R1 | -0.2291443 | 1 | -0.70493 | 1 |
| ANGPTL4 | -0.2714657 | 1 | -0.65853 | 1 |
| MLH1 | -0.1204633 | 1 | -0.80921 | 1 |
| NCOA3 | -0.2413078 | 1 | -0.68782 | 1 |
| BDP1 | -0.235495 | 1 | -0.6927 | 1 |
| RHOJ | 0.05109379 | 1 | -0.97916 | 1 |
| GPR119 | -0.0669216 | 1 | -0.85872 | 1 |
| VHL | -0.1477485 | 1 | -0.77755 | 1 |
| RCOR3 | -0.5141896 | 1 | -0.41018 | 1 |
| KLF8 | -0.6355703 | 1 | -0.286 | 1 |
| DFFA | -0.2821564 | 1 | -0.63902 | 1 |
| BMP6 | -0.4302129 | 1 | -0.48997 | 1 |
| OR2B6 | 0.0239289 | 1 | -0.94403 | 1 |
| ZNF701 | -0.6620625 | 1 | -0.25606 | 1 |
| OR4C15 | -0.0217362 | 1 | -0.89396 | 1 |
| ARHGEF9 | -0.7749334 | 1 | -0.14074 | 1 |
| OR6B2 | -0.790657 | 1 | -0.12361 | 1 |
| SMC3 | -0.3232269 | 1 | -0.5902 | 1 |
| ZSCAN5C | -0.9138066 | 1 | 0.000482 | 1 |
| SGPL1 | -0.6621271 | 1 | -0.25107 | 1 |
| SEPTIN9 | -0.2333371 | 1 | -0.6791 | 1 |
| BAIAP2L1 | -1.6632084 | 1 | 0.752094 | 1 |
| TAF5 | -0.3455382 | 1 | -0.56506 | 1 |
| ZNF608 | -0.637021 | 1 | -0.27332 | 1 |
| STK17B | -0.5685435 | 1 | -0.34108 | 1 |
| GRIN1 | -0.5638538 | 1 | -0.34505 | 1 |
| ENY2 | -0.1066227 | 1 | -0.80178 | 1 |
| ZYX | -0.0640071 | 1 | -0.84228 | 1 |

| | | | | |
|---|---|---|---|---|
| ELAVL2 | -0.6956597 | 1 | -0.20842 | 1 |
| ZNF615 | -0.3051113 | 1 | -0.59858 | 1 |
| INSR | -0.181971 | 1 | -0.72066 | 1 |
| IRAK3 | -0.2842397 | 1 | -0.61778 | 1 |
| CARTPT | -0.8882119 | 1 | -0.01167 | 1 |
| GPR15 | -0.1753011 | 1 | -0.72451 | 1 |
| TAS2R16 | -0.8165048 | 1 | -0.08173 | 1 |
| RBSN | -0.7106027 | 1 | -0.18691 | 1 |
| ZNF831 | -0.0861593 | 1 | -0.811 | 1 |
| OR6K6 | -0.579397 | 1 | -0.31551 | 1 |
| FOS | -0.1606406 | 1 | -0.73394 | 1 |
| RND3 | 0.01393878 | 1 | -0.90836 | 1 |
| ARL8A | -0.9043317 | 1 | 0.011613 | 1 |
| STRADB | -0.7003143 | 1 | -0.19105 | 1 |
| HTR1E | -0.5032583 | 1 | -0.38761 | 1 |
| RALGPS1 | -0.2170382 | 1 | -0.6714 | 1 |
| GPR150 | -0.0576043 | 1 | -0.82796 | 1 |
| TBX1 | 0.11388881 | 1 | -0.99758 | 1 |
| ZNF275 | -0.2039284 | 1 | -0.6797 | 1 |
| ABCG1 | -0.4041388 | 1 | -0.47819 | 1 |
| PHTF2 | -0.2720066 | 1 | -0.6097 | 1 |
| FLT3LG | 0.13163199 | 1 | -1.01168 | 1 |
| RLF | -0.1536868 | 1 | -0.72607 | 1 |
| SERPINB9 | -0.0647533 | 1 | -0.81496 | 1 |
| PLXND1 | -0.1600672 | 1 | -0.71865 | 1 |
| CHRM3 | -0.5918362 | 1 | -0.28678 | 1 |
| GABPA | -0.3320086 | 1 | -0.54654 | 1 |
| MPP3 | -0.3106486 | 1 | -0.56633 | 1 |
| OR52E6 | -0.2155111 | 1 | -0.66069 | 1 |
| KSR1 | -0.9704998 | 1 | 0.096248 | 1 |
| BLZF1 | -0.2810544 | 1 | -0.5925 | 1 |
| PIR | -0.1439193 | 1 | -0.72911 | 1 |
| SCMH1 | -0.4834981 | 1 | -0.38724 | 1 |
| SEPTIN6 | -0.1376652 | 1 | -0.73274 | 1 |
| PGRMC2 | 0.10694854 | 1 | -0.97722 | 1 |
| ZFAT | -0.6561114 | 1 | -0.21414 | 1 |
| ZNF429 | -0.6032454 | 1 | -0.26669 | 1 |
| PEX11B | 0.14931859 | 1 | -1.0185 | 1 |
| MED24 | -0.9434415 | 1 | 0.076178 | 1 |
| ZNF292 | -0.3211752 | 1 | -0.54551 | 1 |
| TADA1 | -0.1769832 | 1 | -0.68927 | 1 |
| ZNF665 | -0.4234956 | 1 | -0.44046 | 1 |
| SOX21 | -0.8844258 | 1 | 0.02064 | 1 |
| ATM | -0.4546066 | 1 | -0.4085 | 1 |
| ZBTB20 | -0.6256689 | 1 | -0.23689 | 1 |
| HAND1 | -0.8553681 | 1 | -0.0066 | 1 |

| | | | | |
|---|---|---|---|---|
| HNF4G | -0.409482 | 1 | -0.45139 | 1 |
| OR6N2 | -0.8152648 | 1 | -0.04555 | 1 |
| ZCCHC10 | 0.06736185 | 1 | -0.92784 | 1 |
| TERF2IP | -1.665477 | 1 | 0.80529 | 1 |
| EDRF1 | -0.3686694 | 1 | -0.48884 | 1 |
| ZNF433 | -0.3288476 | 1 | -0.52774 | 1 |
| HRH4 | -0.8043944 | 1 | -0.05177 | 1 |
| EGR3 | -0.6993314 | 1 | -0.15479 | 1 |
| TSPAN12 | -0.0583876 | 1 | -0.79483 | 1 |
| RSU1 | 0.13438358 | 1 | -0.98748 | 1 |
| PBRM1 | -0.4085871 | 1 | -0.44421 | 1 |
| FGF10 | -0.80431 | 1 | -0.04766 | 1 |
| GNB4 | -0.4782699 | 1 | -0.37176 | 1 |
| ZNF573 | -0.1420054 | 1 | -0.70682 | 1 |
| PPP4R1 | -0.6197957 | 1 | -0.22885 | 1 |
| PEG3 | -0.6785473 | 1 | -0.16736 | 1 |
| ZNF260 | -0.2404171 | 1 | -0.60455 | 1 |
| EYA1 | -0.4383773 | 1 | -0.40631 | 1 |
| CDKN1A | -0.4916709 | 1 | -0.35123 | 1 |
| OR5A1 | -0.3685031 | 1 | -0.47112 | 1 |
| RUNX2 | -0.1059895 | 1 | -0.73272 | 1 |
| PPARGC1A | -0.6401552 | 1 | -0.19683 | 1 |
| PPP2R2A | -0.3066362 | 1 | -0.5303 | 1 |
| CD247 | -0.8408209 | 1 | 0.00443 | 1 |
| CAMK2A | -0.7223181 | 1 | -0.1137 | 1 |
| TEAD1 | -0.1205936 | 1 | -0.71397 | 1 |
| DPYSL3 | -0.0626035 | 1 | -0.76965 | 1 |
| CXXC5 | -1.2815977 | 1 | 0.450669 | 1 |
| ZNF431 | -0.414585 | 1 | -0.41618 | 1 |
| TAC1 | -1.0719989 | 1 | 0.241839 | 1 |
| ZNF8 | -0.8321075 | 1 | 0.002959 | 1 |
| OR5AP2 | -0.850866 | 1 | 0.022132 | 1 |
| PTPRZ1 | -0.5555545 | 1 | -0.27308 | 1 |
| ELF4 | -0.3161307 | 1 | -0.5098 | 1 |
| MLLT10 | -0.1989295 | 1 | -0.62664 | 1 |
| SPSB4 | -0.9909873 | 1 | 0.166802 | 1 |
| HCAR2 | -0.1472656 | 1 | -0.67605 | 1 |
| ARHGAP35 | -1.2940168 | 1 | 0.47162 | 1 |
| OR51B4 | -0.8844479 | 1 | 0.062345 | 1 |
| MED23 | -0.3098721 | 1 | -0.51137 | 1 |
| ZNF681 | -0.3451934 | 1 | -0.47581 | 1 |
| HIC1 | 0.12843064 | 1 | -0.94872 | 1 |
| TFE3 | -0.5127817 | 1 | -0.30564 | 1 |
| CD83 | -0.1258534 | 1 | -0.692 | 1 |
| MAPK15 | -0.2829058 | 1 | -0.53476 | 1 |
| KCNIP1 | -0.2682234 | 1 | -0.54917 | 1 |

| | | | | |
|---|---|---|---|---|
| KCNH5 | -0.2036269 | 1 | -0.61332 | 1 |
| TRAT1 | -1.8071404 | 1 | 0.990581 | 1 |
| UBE3A | -0.0281524 | 1 | -0.78786 | 1 |
| FOXI1 | -0.7339906 | 1 | -0.08085 | 1 |
| BTG1 | -0.6498532 | 1 | -0.16453 | 1 |
| EGR4 | -1.5192993 | 1 | 0.705536 | 1 |
| OR8B8 | -0.0797258 | 1 | -0.73342 | 1 |
| SIRT1 | -0.0563927 | 1 | -0.75662 | 1 |
| PDE1B | -0.668222 | 1 | -0.1443 | 1 |
| PTK2 | -0.1317658 | 1 | -0.68014 | 1 |
| CHRNA5 | -0.0710586 | 1 | -0.73982 | 1 |
| ACKR3 | -0.3544432 | 1 | -0.45586 | 1 |
| ASB13 | -0.9871868 | 1 | 0.176964 | 1 |
| OR2L2 | -0.2615104 | 1 | -0.54826 | 1 |
| OR6K3 | -0.0744605 | 1 | -0.73515 | 1 |
| CLOCK | -0.0971547 | 1 | -0.71183 | 1 |
| TIAL1 | -0.3884036 | 1 | -0.42015 | 1 |
| ZFAND1 | -0.2129334 | 1 | -0.59548 | 1 |
| PPP2R2B | -1.4449444 | 1 | 0.636862 | 1 |
| ZNF777 | -0.8384344 | 1 | 0.031373 | 1 |
| CYP26A1 | -0.1444989 | 1 | -0.66206 | 1 |
| OGT | -0.3448326 | 1 | -0.46161 | 1 |
| AJUBA | -1.0520394 | 1 | 0.246014 | 1 |
| PDE9A | -0.1356253 | 1 | -0.67023 | 1 |
| ZNF560 | -0.3992077 | 1 | -0.40635 | 1 |
| MYBL1 | -0.2953718 | 1 | -0.50836 | 1 |
| CRYAB | -0.2606191 | 1 | -0.54174 | 1 |
| KDM4C | -0.5621338 | 1 | -0.23982 | 1 |
| MYT1L | -0.7009978 | 1 | -0.1005 | 1 |
| PAK5 | -1.2759783 | 1 | 0.474943 | 1 |
| BCL2L13 | -0.0926183 | 1 | -0.70748 | 1 |
| SLC7A3 | -0.9347811 | 1 | 0.135005 | 1 |
| HTT | -0.4537473 | 1 | -0.34595 | 1 |
| HRH3 | -0.2459358 | 1 | -0.55265 | 1 |
| ARL5A | -0.2507611 | 1 | -0.54688 | 1 |
| GNAI1 | -1.0268746 | 1 | 0.22969 | 1 |
| CUL3 | 0.12467686 | 1 | -0.91817 | 1 |
| RASA2 | -0.1782016 | 1 | -0.61483 | 1 |
| MED13L | -0.4089244 | 1 | -0.38225 | 1 |
| TRIP4 | -0.3352157 | 1 | -0.45484 | 1 |
| KDSR | 0.11287647 | 1 | -0.90287 | 1 |
| MDFIC | -0.7623615 | 1 | -0.02615 | 1 |
| SORCS1 | -0.0292647 | 1 | -0.75885 | 1 |
| HPGD | -0.1060053 | 1 | -0.68147 | 1 |
| DEPDC7 | -0.2637951 | 1 | -0.5234 | 1 |
| TRAF1 | -0.3887794 | 1 | -0.3983 | 1 |

| | | | | |
|---|---|---|---|---|
| NKX6.1 | -0.0807589 | 1 | -0.70447 | 1 |
| PDZD8 | -0.4918184 | 1 | -0.29244 | 1 |
| RAPGEF2 | -0.8662459 | 1 | 0.085185 | 1 |
| UBP1 | -0.2128361 | 1 | -0.56735 | 1 |
| ID2 | -0.2333982 | 1 | -0.54677 | 1 |
| NR2C1 | -0.2446434 | 1 | -0.53535 | 1 |
| IL27RA | -0.854911 | 1 | 0.075152 | 1 |
| RAB37 | -0.9581117 | 1 | 0.178354 | 1 |
| NRGN | -0.8812838 | 1 | 0.101739 | 1 |
| MBIP | -0.4676943 | 1 | -0.31144 | 1 |
| GPBP1L1 | 0.04008549 | 1 | -0.81898 | 1 |
| NOBOX | -0.0731108 | 1 | -0.70549 | 1 |
| RANBP2 | -0.0975608 | 1 | -0.67907 | 1 |
| CSRNP3 | -0.6920937 | 1 | -0.08413 | 1 |
| OR5B3 | -0.147647 | 1 | -0.62646 | 1 |
| PDYN | -0.684122 | 1 | -0.08895 | 1 |
| HRH2 | -0.0611569 | 1 | -0.71166 | 1 |
| TACR2 | -0.1684117 | 1 | -0.60426 | 1 |
| UNC13B | -0.4684245 | 1 | -0.3039 | 1 |
| HUS1 | -0.3545453 | 1 | -0.41777 | 1 |
| G3BP1 | -0.1818773 | 1 | -0.58958 | 1 |
| ZBTB8A | -0.2697497 | 1 | -0.50153 | 1 |
| TNFRSF10B | -0.6971744 | 1 | -0.07204 | 1 |
| CNKSR3 | -0.0658387 | 1 | -0.70284 | 1 |
| STXBP4 | -0.4154642 | 1 | -0.35319 | 1 |
| NDFIP1 | -0.4189113 | 1 | -0.34857 | 1 |
| MSI2 | -0.3883278 | 1 | -0.37731 | 1 |
| ZNF823 | -0.1084104 | 1 | -0.65517 | 1 |
| PRDM6 | 0.07419357 | 1 | -0.83777 | 1 |
| SETD3 | -0.1410304 | 1 | -0.62242 | 1 |
| ZMAT3 | -0.7821674 | 1 | 0.018754 | 1 |
| CRABP1 | -0.5990602 | 1 | -0.16421 | 1 |
| CHD6 | -0.3412818 | 1 | -0.42037 | 1 |
| LITAF | -0.6045923 | 1 | -0.15666 | 1 |
| TIGD6 | -0.3428801 | 1 | -0.41836 | 1 |
| APC | -0.5499987 | 1 | -0.20728 | 1 |
| SSX4B | -0.710705 | 1 | -0.04531 | 1 |
| MAST1 | -0.402227 | 1 | -0.3532 | 1 |
| BAK1 | -0.5104293 | 1 | -0.24499 | 1 |
| SLK | -0.1314374 | 1 | -0.62398 | 1 |
| ZNF527 | -0.038304 | 1 | -0.7154 | 1 |
| RAB23 | -0.5205409 | 1 | -0.23309 | 1 |
| ERV3.1 | -0.1349928 | 1 | -0.61748 | 1 |
| ZFP69 | -0.4974902 | 1 | -0.25488 | 1 |
| RAF1 | -0.1176717 | 1 | -0.63437 | 1 |
| ING1 | -0.5551823 | 1 | -0.19684 | 1 |

| | | | | |
|---|---|---|---|---|
| TAF11 | 0.57366253 | 1 | -1.32524 | 1 |
| ZNF540 | 0.01629276 | 1 | -0.76584 | 1 |
| CNOT1 | -0.3410507 | 1 | -0.4069 | 1 |
| PRKAG2 | -0.7485969 | 1 | 0.002144 | 1 |
| PROX1 | -0.2282671 | 1 | -0.51755 | 1 |
| RPS6KA5 | -0.8185103 | 1 | 0.073348 | 1 |
| DNMT1 | -0.3182676 | 1 | -0.42639 | 1 |
| H4C8 | -0.6928875 | 1 | -0.0498 | 1 |
| LILRB3 | -0.152818 | 1 | -0.58963 | 1 |
| ZNF131 | -0.0598855 | 1 | -0.68255 | 1 |
| OR7A17 | -0.6322203 | 1 | -0.11018 | 1 |
| YBX2 | -0.7903313 | 1 | 0.048163 | 1 |
| PPARA | -1.1268271 | 1 | 0.386366 | 1 |
| RGS20 | 0.0411855 | 1 | -0.78149 | 1 |
| RGS17 | -0.1357198 | 1 | -0.60378 | 1 |
| KRIT1 | -0.2572674 | 1 | -0.48022 | 1 |
| OR51Q1 | -0.4045863 | 1 | -0.33267 | 1 |
| STAC | 0.01607593 | 1 | -0.75271 | 1 |
| CARF | 0.26479913 | 1 | -1.00108 | 1 |
| POU2F3 | 0.31075919 | 1 | -1.04593 | 1 |
| FOSB | 0.08983532 | 1 | -0.82496 | 1 |
| HOXB13 | 1.22596143 | 1 | -1.96026 | 1 |
| HTR5A | -0.3148784 | 1 | -0.41672 | 1 |
| TP53 | -0.7018911 | 1 | -0.02933 | 1 |
| BCL2L11 | -0.5243589 | 1 | -0.20629 | 1 |
| ZBTB21 | -0.2937227 | 1 | -0.43505 | 1 |
| GRAP2 | -0.6664972 | 1 | -0.05997 | 1 |
| GPRC5D | -0.1041457 | 1 | -0.62193 | 1 |
| PHLDA3 | -1.3287601 | 1 | 0.603187 | 1 |
| ADGRA1 | -2.1826805 | 1 | 1.458213 | 1 |
| RASL10B | -1.5588509 | 1 | 0.834996 | 1 |
| SDHA | -0.5799692 | 1 | -0.14336 | 1 |
| INPP4B | -0.4840473 | 1 | -0.23889 | 1 |
| ZNF813 | -0.2969517 | 1 | -0.42541 | 1 |
| ARHGAP42 | -0.3581539 | 1 | -0.36365 | 1 |
| FLRT3 | -0.3207833 | 1 | -0.40012 | 1 |
| ARHGAP28 | -0.0322152 | 1 | -0.68865 | 1 |
| CECR2 | -0.349063 | 1 | -0.37108 | 1 |
| MSX2 | -0.5354485 | 1 | -0.18402 | 1 |
| NR2E3 | -0.6059109 | 1 | -0.11305 | 1 |
| ARHGEF12 | -0.4966431 | 1 | -0.22135 | 1 |
| TAF7 | -0.3509453 | 1 | -0.36557 | 1 |
| SSBP2 | -0.5879428 | 1 | -0.12848 | 1 |
| RFC1 | -0.3025672 | 1 | -0.4136 | 1 |
| ZSWIM5 | -0.7094171 | 1 | -0.00641 | 1 |
| JAG2 | -0.2702845 | 1 | -0.44391 | 1 |

| | | | | |
|---|---|---|---|---|
| RAD54L2 | -0.4731934 | 1 | -0.2379 | 1 |
| OR2D3 | 0.03844442 | 1 | -0.74926 | 1 |
| RRAS2 | -0.6805329 | 1 | -0.03004 | 1 |
| ZXDA | -0.4224502 | 1 | -0.28812 | 1 |
| OR7A10 | -0.0991215 | 1 | -0.60594 | 1 |
| RGS21 | -0.9036328 | 1 | 0.199142 | 1 |
| MCTP2 | -0.349575 | 1 | -0.3549 | 1 |
| ARL6 | -0.063398 | 1 | -0.63999 | 1 |
| BATF3 | -0.463289 | 1 | -0.23872 | 1 |
| EIF2AK4 | -0.3993474 | 1 | -0.30238 | 1 |
| FMN2 | -0.1094606 | 1 | -0.59168 | 1 |
| NR6A1 | -0.203539 | 1 | -0.4968 | 1 |
| PTPN2 | -0.027408 | 1 | -0.67211 | 1 |
| PLAG1 | -0.2056419 | 1 | -0.4929 | 1 |
| PDE10A | -0.4584573 | 1 | -0.24007 | 1 |
| SFRP5 | -0.2671092 | 1 | -0.43103 | 1 |
| SCRIB | -0.0447832 | 1 | -0.65215 | 1 |
| RASEF | -0.6257981 | 1 | -0.07081 | 1 |
| PPM1A | -0.2896486 | 1 | -0.40692 | 1 |
| ELF2 | -0.1308172 | 1 | -0.56556 | 1 |
| JUP | -0.6327647 | 1 | -0.06335 | 1 |
| RABIF | 0.04628456 | 1 | -0.7399 | 1 |
| SPOCK1 | -0.7774664 | 1 | 0.084489 | 1 |
| EIF4EBP2 | -0.2603713 | 1 | -0.43228 | 1 |
| GADD45B | -0.0707273 | 1 | -0.62103 | 1 |
| ZNF175 | -0.744088 | 1 | 0.052706 | 1 |
| TCF4 | -0.1678178 | 1 | -0.52303 | 1 |
| TRAF6 | -0.3464663 | 1 | -0.34419 | 1 |
| NPY1R | -0.0380755 | 1 | -0.65162 | 1 |
| EFNA1 | -0.1693425 | 1 | -0.51887 | 1 |
| ARID4A | -0.4364395 | 1 | -0.25172 | 1 |
| CD101 | -0.0207168 | 1 | -0.6673 | 1 |
| ZNF250 | -0.3366185 | 1 | -0.35134 | 1 |
| SEL1L | -0.3174472 | 1 | -0.37037 | 1 |
| MRGPRE | -0.4426918 | 1 | -0.24341 | 1 |
| CORO2A | -0.777149 | 1 | 0.091942 | 1 |
| SDCBP | -0.1760792 | 1 | -0.50834 | 1 |
| OR4F5 | -0.7437989 | 1 | 0.060741 | 1 |
| BRPF1 | -0.3061582 | 1 | -0.37685 | 1 |
| TBC1D32 | -0.1917105 | 1 | -0.49007 | 1 |
| ZNF516 | -0.7051233 | 1 | 0.024648 | 1 |
| CYP27B1 | -0.7436157 | 1 | 0.063511 | 1 |
| AKAP11 | -0.5483471 | 1 | -0.13116 | 1 |
| GAB1 | -0.3892589 | 1 | -0.29018 | 1 |
| PDE8B | -0.5602016 | 1 | -0.1189 | 1 |
| AMER1 | -0.4346886 | 1 | -0.24434 | 1 |

| | | | | |
|---|---|---|---|---|
| GRIK1 | -1.0201884 | 1 | 0.341528 | 1 |
| PHIP | -0.2702574 | 1 | -0.40828 | 1 |
| PHF2 | -0.8882314 | 1 | 0.209858 | 1 |
| GNGT1 | -0.3417955 | 1 | -0.33657 | 1 |
| OCRL | -0.255786 | 1 | -0.42252 | 1 |
| STK24 | -0.2833156 | 1 | -0.3941 | 1 |
| CLCN6 | -0.6676059 | 1 | -0.00947 | 1 |
| YBX3 | -0.2160401 | 1 | -0.46 | 1 |
| OXTR | -1.2325246 | 1 | 0.557198 | 1 |
| OR4K17 | -0.2941023 | 1 | -0.38095 | 1 |
| ZBTB25 | -0.1251006 | 1 | -0.54903 | 1 |
| NSG1 | -0.4310691 | 1 | -0.24247 | 1 |
| CREB3L2 | -0.0290555 | 1 | -0.64427 | 1 |
| LIN54 | -0.2670862 | 1 | -0.40592 | 1 |
| CTNND1 | -0.4487351 | 1 | -0.22306 | 1 |
| INPP4A | -0.2473558 | 1 | -0.4228 | 1 |
| P2RY4 | 0.01265483 | 1 | -0.68169 | 1 |
| ACVR2A | -0.1062242 | 1 | -0.56273 | 1 |
| CCR7 | -0.5496905 | 1 | -0.11848 | 1 |
| PTH2R | -0.7674535 | 1 | 0.099701 | 1 |
| ZFP1 | -0.2446036 | 1 | -0.42284 | 1 |
| BTLA | -0.1428402 | 1 | -0.52429 | 1 |
| SAFB | -0.4695286 | 1 | -0.19735 | 1 |
| ZNF829 | -0.1083933 | 1 | -0.55839 | 1 |
| ILF3 | -0.2006725 | 1 | -0.46399 | 1 |
| DLC1 | -0.6878197 | 1 | 0.023921 | 1 |
| TGFBR2 | -0.3037463 | 1 | -0.35927 | 1 |
| RASSF5 | -0.2686497 | 1 | -0.39374 | 1 |
| GULP1 | -0.2921917 | 1 | -0.36992 | 1 |
| NCK1 | -0.1624394 | 1 | -0.49944 | 1 |
| PRDM10 | -0.2848956 | 1 | -0.37692 | 1 |
| PHOX2A | -0.0136355 | 1 | -0.64727 | 1 |
| PCM1 | -0.0418752 | 1 | -0.61894 | 1 |
| GNA12 | -0.7813238 | 1 | 0.120585 | 1 |
| RAPGEF4 | -0.5964793 | 1 | -0.063 | 1 |
| USP6NL | -0.2390586 | 1 | -0.41915 | 1 |
| HDAC3 | 0.08137199 | 1 | -0.73955 | 1 |
| GNB1 | -0.5176208 | 1 | -0.14054 | 1 |
| TNFSF15 | -0.3653026 | 1 | -0.29195 | 1 |
| SATB1 | -0.4316915 | 1 | -0.22445 | 1 |
| ZNF383 | -0.1521925 | 1 | -0.50363 | 1 |
| STRN | -0.2573158 | 1 | -0.39847 | 1 |
| KMT2C | -0.5094196 | 1 | -0.1455 | 1 |
| ELANE | -1.2496193 | 1 | 0.594797 | 1 |
| RAB31 | -0.3458621 | 1 | -0.3081 | 1 |
| TAS2R43 | -0.2468098 | 1 | -0.40564 | 1 |

| | | | | |
|---|---|---|---|---|
| NOTCH2NLA | -0.0103828 | 1 | -0.64075 | 1 |
| DBX1 | -1.1069783 | 1 | 0.456457 | 1 |
| CTNNB1 | -0.096936 | 1 | -0.55339 | 1 |
| ZC3H12C | -0.2695101 | 1 | -0.38066 | 1 |
| UBB | -0.2228472 | 1 | -0.42678 | 1 |
| ZDHHC15 | -0.4076158 | 1 | -0.24034 | 1 |
| OR9G1 | -0.1687321 | 1 | -0.47583 | 1 |
| AFDN | -0.0144705 | 1 | -0.62961 | 1 |
| MDM4 | 0.20060036 | 1 | -0.84409 | 1 |
| ASB11 | -0.0852257 | 1 | -0.55734 | 1 |
| FUBP1 | -0.2720333 | 1 | -0.3702 | 1 |
| HOXB3 | -0.1842161 | 1 | -0.45782 | 1 |
| TENT5C | -0.2608977 | 1 | -0.38112 | 1 |
| ZNF280B | -0.3507087 | 1 | -0.29109 | 1 |
| MAP4K3 | -0.2013028 | 1 | -0.44038 | 1 |
| ZFP82 | -0.2287551 | 1 | -0.41224 | 1 |
| ZNF699 | -0.2995513 | 1 | -0.34143 | 1 |
| HTR2A | -0.5773592 | 1 | -0.0634 | 1 |
| WNK3 | -0.2641048 | 1 | -0.37656 | 1 |
| ATF7IP | -0.492419 | 1 | -0.14668 | 1 |
| OR5AS1 | -0.3568796 | 1 | -0.28077 | 1 |
| HOXD13 | -0.3459557 | 1 | -0.28968 | 1 |
| CD27 | -0.5575822 | 1 | -0.07744 | 1 |
| CSNK1G1 | -0.3639309 | 1 | -0.27051 | 1 |
| TBX18 | -0.086885 | 1 | -0.54739 | 1 |
| ZCCHC8 | -0.1457931 | 1 | -0.48677 | 1 |
| ARHGDIG | -0.3299057 | 1 | -0.30218 | 1 |
| GTF2H3 | -0.1136813 | 1 | -0.51766 | 1 |
| DCAF6 | -0.5065316 | 1 | -0.12401 | 1 |
| ZNF543 | -0.1353514 | 1 | -0.49455 | 1 |
| DDX10 | -0.2695802 | 1 | -0.36016 | 1 |
| GCFC2 | -0.0900962 | 1 | -0.53869 | 1 |
| OR51E2 | 0.00386117 | 1 | -0.63068 | 1 |
| LEO1 | -0.1361109 | 1 | -0.49053 | 1 |
| ZNF10 | -0.0097404 | 1 | -0.61546 | 1 |
| BNIP3 | -0.3164037 | 1 | -0.308 | 1 |
| PRKD3 | -0.1752133 | 1 | -0.44911 | 1 |
| DVL3 | -0.6380475 | 1 | 0.014529 | 1 |
| CERS4 | -1.0106571 | 1 | 0.387359 | 1 |
| CBX5 | -0.329811 | 1 | -0.29334 | 1 |
| LECT2 | -0.1468584 | 1 | -0.47596 | 1 |
| NUFIP1 | -0.100125 | 1 | -0.52224 | 1 |
| PHF10 | -0.2277612 | 1 | -0.39445 | 1 |
| CHRNA3 | -0.5842312 | 1 | -0.03775 | 1 |
| OR13C5 | -0.13089 | 1 | -0.49104 | 1 |
| GLRX2 | 0.37646352 | 1 | -0.99797 | 1 |

| | | | | |
|---|---|---|---|---|
| MNAT1 | -0.2661424 | 1 | -0.35535 | 1 |
| RBM14 | -0.305556 | 1 | -0.31322 | 1 |
| EGR1 | -0.1236756 | 1 | -0.49408 | 1 |
| ZNF28 | -0.1519756 | 1 | -0.46537 | 1 |
| OR51T1 | -0.0519434 | 1 | -0.56517 | 1 |
| ZKSCAN5 | -0.1092466 | 1 | -0.50786 | 1 |
| GPR83 | -0.3796284 | 1 | -0.23625 | 1 |
| IFNB1 | -0.5502236 | 1 | -0.06467 | 1 |
| ZNF619 | -1.3361732 | 1 | 0.721314 | 1 |
| RALB | 0.58433092 | 1 | -1.19913 | 1 |
| CD164 | -0.0642079 | 1 | -0.54986 | 1 |
| OR6Y1 | -0.0969564 | 1 | -0.51647 | 1 |
| WWP1 | -0.2089002 | 1 | -0.40451 | 1 |
| TBX20 | -0.1147719 | 1 | -0.49841 | 1 |
| ITGA4 | -0.328467 | 1 | -0.28404 | 1 |
| EDARADD | 0.5074499 | 1 | -1.11962 | 1 |
| TFAM | 0.03535976 | 1 | -0.64559 | 1 |
| ZSCAN30 | -0.1853417 | 1 | -0.42395 | 1 |
| HDX | -0.3169202 | 1 | -0.29147 | 1 |
| RORA | -0.3233685 | 1 | -0.28394 | 1 |
| HOXC9 | 0.04592848 | 1 | -0.65262 | 1 |
| OR52R1 | 0.01581446 | 1 | -0.62155 | 1 |
| STK17A | -0.1205728 | 1 | -0.48453 | 1 |
| PPP1CB | 0.10529462 | 1 | -0.71021 | 1 |
| PDK1 | -0.1336018 | 1 | -0.47113 | 1 |
| RAB38 | -0.0594211 | 1 | -0.54483 | 1 |
| MED12L | -0.7980559 | 1 | 0.194733 | 1 |
| RBPJL | -0.6173825 | 1 | 0.014353 | 1 |
| PIK3C3 | -0.4749964 | 1 | -0.12739 | 1 |
| GRIK5 | -0.8831123 | 1 | 0.282206 | 1 |
| PRDM8 | -0.6044805 | 1 | 0.00361 | 1 |
| PLCE1 | -0.4417396 | 1 | -0.15782 | 1 |
| PCLO | -0.3952612 | 1 | -0.20355 | 1 |
| RHEB | -0.2825528 | 1 | -0.31583 | 1 |
| ALX4 | -0.5133556 | 1 | -0.08407 | 1 |
| YAP1 | -0.2279484 | 1 | -0.36835 | 1 |
| P2RY10 | 0.03192455 | 1 | -0.62768 | 1 |
| HOXA5 | -0.7222084 | 1 | 0.126489 | 1 |
| ZNF254 | -0.1241097 | 1 | -0.47158 | 1 |
| BLOC1S2 | 0.04485708 | 1 | -0.64049 | 1 |
| IL1RAP | -0.4015115 | 1 | -0.19381 | 1 |
| FAS | -0.3273244 | 1 | -0.26778 | 1 |
| APBB2 | -1.7760675 | 1 | 1.181376 | 1 |
| CELSR2 | -0.5217118 | 1 | -0.07296 | 1 |
| ZBTB26 | -0.0851012 | 1 | -0.50931 | 1 |
| ZNF805 | 0.01923425 | 1 | -0.61226 | 1 |

| | | | | |
|---|---|---|---|---|
| AGK | -0.1419317 | 1 | -0.45076 | 1 |
| STAMBP | -0.1554242 | 1 | -0.437 | 1 |
| ZNF407 | -1.2068494 | 1 | 0.615836 | 1 |
| OR4S2 | -0.6035427 | 1 | 0.0126 | 1 |
| ZBTB24 | 0.47728725 | 1 | -1.06708 | 1 |
| TRH | -0.5366511 | 1 | -0.05291 | 1 |
| OR6K2 | -0.0185913 | 1 | -0.57015 | 1 |
| DGKE | -0.2290103 | 1 | -0.35922 | 1 |
| OR2S2 | -0.5211194 | 1 | -0.06701 | 1 |
| NOLC1 | 0.1499042 | 1 | -0.73655 | 1 |
| RPS6KA1 | -0.7757092 | 1 | 0.189282 | 1 |
| ZNF595 | -0.1599244 | 1 | -0.42633 | 1 |
| PTGES2 | -0.6715678 | 1 | 0.085672 | 1 |
| HUNK | -0.3119555 | 1 | -0.27281 | 1 |
| DNMBP | -0.0719161 | 1 | -0.51283 | 1 |
| TAF1 | -0.266947 | 1 | -0.3168 | 1 |
| RASL11B | -0.3258481 | 1 | -0.25676 | 1 |
| HDAC9 | -0.4861013 | 1 | -0.09527 | 1 |
| ZCRB1 | -0.5956186 | 1 | 0.014558 | 1 |
| GALR3 | -1.3963527 | 1 | 0.815442 | 1 |
| ZNF32 | -0.6829781 | 1 | 0.102915 | 1 |
| ZNHIT3 | -0.0831606 | 1 | -0.49573 | 1 |
| BCL7A | -0.5043566 | 1 | -0.07428 | 1 |
| LGR5 | -0.0823481 | 1 | -0.49576 | 1 |
| ZBTB37 | -0.214286 | 1 | -0.36363 | 1 |
| GPR12 | -0.3073842 | 1 | -0.2702 | 1 |
| BRAP | -0.1515868 | 1 | -0.42583 | 1 |
| ZNF410 | 0.04289259 | 1 | -0.62003 | 1 |
| BBX | -0.3871137 | 1 | -0.18994 | 1 |
| GRM6 | -1.026359 | 1 | 0.449341 | 1 |
| TNFRSF13B | -0.4051872 | 1 | -0.17145 | 1 |
| RASL10A | -1.1840897 | 1 | 0.609143 | 1 |
| ZNF384 | -0.406753 | 1 | -0.16707 | 1 |
| PTH1R | -0.3943466 | 1 | -0.17788 | 1 |
| OR2L13 | -0.3478596 | 1 | -0.22431 | 1 |
| ZNF808 | 0.26528013 | 1 | -0.83727 | 1 |
| KLF6 | -0.072763 | 1 | -0.49915 | 1 |
| ZNF512 | -0.1854771 | 1 | -0.38596 | 1 |
| ZNF625 | -0.2354307 | 1 | -0.33565 | 1 |
| ERBIN | -0.2672293 | 1 | -0.30319 | 1 |
| MBD2 | -0.4761386 | 1 | -0.09402 | 1 |
| OR13F1 | -0.4834083 | 1 | -0.08459 | 1 |
| PRKG1 | -0.5070154 | 1 | -0.06041 | 1 |
| TUT4 | -0.09109 | 1 | -0.47624 | 1 |
| BIRC3 | -0.2901914 | 1 | -0.27708 | 1 |
| MAP3K14 | -0.701535 | 1 | 0.135746 | 1 |

| | | | | |
|---|---|---|---|---|
| RALGDS | -0.8715185 | 1 | 0.307148 | 1 |
| RGL4 | -0.2099427 | 1 | -0.35362 | 1 |
| ZMYM6 | -0.0995986 | 1 | -0.46381 | 1 |
| HCRTR2 | -0.4593942 | 1 | -0.10348 | 1 |
| STK4 | -0.1678247 | 1 | -0.39421 | 1 |
| DLL3 | -1.1490836 | 1 | 0.587143 | 1 |
| RASAL1 | -0.1026755 | 1 | -0.45854 | 1 |
| RALGPS2 | -0.18493 | 1 | -0.37514 | 1 |
| ZSCAN23 | -0.8770672 | 1 | 0.318434 | 1 |
| TAF4 | -0.2294894 | 1 | -0.32885 | 1 |
| APOA1 | -0.204506 | 1 | -0.35268 | 1 |
| PPP1R13L | -0.5789198 | 1 | 0.021768 | 1 |
| CHUK | 0.05996948 | 1 | -0.61709 | 1 |
| RPGRIP1L | -0.2855606 | 1 | -0.27084 | 1 |
| MAMLD1 | -0.2985564 | 1 | -0.25639 | 1 |
| ZNF264 | -0.3689315 | 1 | -0.1857 | 1 |
| ZNF280D | -0.28016 | 1 | -0.27371 | 1 |
| ABCA1 | 0.01786568 | 1 | -0.57114 | 1 |
| LPAR3 | -0.4614745 | 1 | -0.0909 | 1 |
| ONECUT1 | -0.5317167 | 1 | -0.02032 | 1 |
| GPR135 | -0.3372541 | 1 | -0.21418 | 1 |
| GALR1 | -0.2622429 | 1 | -0.28881 | 1 |
| ARHGAP12 | -0.17913 | 1 | -0.37158 | 1 |
| HES6 | -0.8658244 | 1 | 0.315219 | 1 |
| HBS1L | -0.0647914 | 1 | -0.48552 | 1 |
| SPRY3 | -0.3929709 | 1 | -0.15573 | 1 |
| GTPBP1 | -0.314929 | 1 | -0.23184 | 1 |
| TESC | -0.4532448 | 1 | -0.09162 | 1 |
| GRID2 | -0.1963133 | 1 | -0.34829 | 1 |
| NFXL1 | -0.132984 | 1 | -0.41034 | 1 |
| SETDB1 | -0.2698559 | 1 | -0.27339 | 1 |
| TRIO | -0.7499425 | 1 | 0.20731 | 1 |
| PDIA3 | -0.1830032 | 1 | -0.35769 | 1 |
| ANXA4 | -0.257997 | 1 | -0.2814 | 1 |
| FYB1 | -0.2512593 | 1 | -0.28808 | 1 |
| TENM2 | -0.2458938 | 1 | -0.29233 | 1 |
| ZNF98 | -0.2860567 | 1 | -0.25158 | 1 |
| DUSP8 | -0.3527117 | 1 | -0.18235 | 1 |
| RGS9BP | -0.8344517 | 1 | 0.299872 | 1 |
| ADAMTS1 | -0.1817494 | 1 | -0.35222 | 1 |
| GJA1 | -0.3500306 | 1 | -0.18267 | 1 |
| ASB2 | -0.1539104 | 1 | -0.37814 | 1 |
| ZNF273 | -0.2291407 | 1 | -0.30266 | 1 |
| TET2 | -0.3696498 | 1 | -0.16046 | 1 |
| CEP63 | -0.2986815 | 1 | -0.23126 | 1 |
| CARD8 | -0.2125963 | 1 | -0.31659 | 1 |

| | | | | |
|---|---|---|---|---|
| ZFP37 | -0.2355764 | 1 | -0.29316 | 1 |
| TNFRSF10C | -0.2715819 | 1 | -0.25572 | 1 |
| NPY5R | -0.1437507 | 1 | -0.38199 | 1 |
| LANCL1 | -0.2462284 | 1 | -0.27848 | 1 |
| FBXO8 | -0.1022666 | 1 | -0.42085 | 1 |
| PDE6H | -0.5662447 | 1 | 0.043401 | 1 |
| GPR151 | -0.2332022 | 1 | -0.28951 | 1 |
| AEBP2 | 0.00780262 | 1 | -0.53008 | 1 |
| PRKACA | 0.06078634 | 1 | -0.58112 | 1 |
| NFE2L3 | -0.27245 | 1 | -0.2475 | 1 |
| GPR45 | -0.3135891 | 1 | -0.20635 | 1 |
| OR2T6 | -0.3050773 | 1 | -0.21467 | 1 |
| PHF12 | -0.4025767 | 1 | -0.11661 | 1 |
| ITGA1 | -0.1553205 | 1 | -0.36292 | 1 |
| SLC7A1 | 0.02744904 | 1 | -0.54564 | 1 |
| ASCL3 | -0.0027622 | 1 | -0.5153 | 1 |
| ZNF721 | -0.0904918 | 1 | -0.4262 | 1 |
| ZC3HAV1L | -0.3375643 | 1 | -0.1791 | 1 |
| CAD | -0.3229417 | 1 | -0.19167 | 1 |
| GABRQ | -1.8207562 | 1 | 1.306296 | 1 |
| OR2T29 | -0.1614955 | 1 | -0.35251 | 1 |
| RORC | -0.4724444 | 1 | -0.04085 | 1 |
| ZNF17 | -0.0667308 | 1 | -0.44463 | 1 |
| EGR2 | -0.2565023 | 1 | -0.25461 | 1 |
| ZDHHC9 | -0.2589654 | 1 | -0.25209 | 1 |
| CLIC2 | 0.09653999 | 1 | -0.60726 | 1 |
| SSTR1 | -0.6015002 | 1 | 0.091977 | 1 |
| ZNF746 | -0.6689232 | 1 | 0.159438 | 1 |
| WSB1 | -0.2014663 | 1 | -0.30797 | 1 |
| MED17 | 0.05823537 | 1 | -0.56758 | 1 |
| EMSY | -0.2804062 | 1 | -0.22863 | 1 |
| DAPK3 | -1.3508888 | 1 | 0.842021 | 1 |
| XPR1 | -0.1373195 | 1 | -0.37088 | 1 |
| RGS1 | -0.3171836 | 1 | -0.191 | 1 |
| GFRA2 | -0.3795377 | 1 | -0.12857 | 1 |
| RRAGB | -0.4578464 | 1 | -0.04915 | 1 |
| ADGRL4 | 0.50147714 | 1 | -1.00844 | 1 |
| CIAO1 | 0.08514602 | 1 | -0.59209 | 1 |
| OR5T3 | -0.5511611 | 1 | 0.044439 | 1 |
| YWHAB | 0.63628516 | 1 | -1.14256 | 1 |
| HOXD4 | -0.4215867 | 1 | -0.08282 | 1 |
| GATA1 | -0.0284965 | 1 | -0.47521 | 1 |
| CRKL | -0.0933767 | 1 | -0.40987 | 1 |
| ZRANB1 | -0.3810827 | 1 | -0.12145 | 1 |
| GAL | -0.3265474 | 1 | -0.17584 | 1 |
| KIF13B | -1.4822881 | 1 | 0.980182 | 1 |

| | | | | |
|---|---|---|---|---|
| JMJD1C | -0.1488687 | 1 | -0.35312 | 1 |
| ADRA2A | -0.0235938 | 1 | -0.47774 | 1 |
| GNAZ | -0.4434877 | 1 | -0.05732 | 1 |
| NEO1 | -0.3182553 | 1 | -0.18252 | 1 |
| RARA | -0.3849003 | 1 | -0.11573 | 1 |
| NCOA2 | -0.0494595 | 1 | -0.45076 | 1 |
| ZNF791 | -0.118937 | 1 | -0.38104 | 1 |
| ETV4 | 0.50831878 | 1 | -1.00814 | 1 |
| MLLT3 | -0.0978306 | 1 | -0.40176 | 1 |
| JAK1 | -0.2885186 | 1 | -0.21071 | 1 |
| HSP90AB1 | 0.1586177 | 1 | -0.65707 | 1 |
| PRKN | -0.1817082 | 1 | -0.31635 | 1 |
| PLAGL1 | -0.1353572 | 1 | -0.36236 | 1 |
| WNT2 | -0.3956138 | 1 | -0.10204 | 1 |
| CDC42BPA | -0.2712805 | 1 | -0.2263 | 1 |
| ZBTB45 | -1.2790561 | 1 | 0.782403 | 1 |
| CD5 | -0.3046318 | 1 | -0.19151 | 1 |
| ASF1A | -0.0453803 | 1 | -0.45064 | 1 |
| VGLL4 | -0.2169104 | 1 | -0.27823 | 1 |
| TOM1L1 | -0.426889 | 1 | -0.06743 | 1 |
| NFYA | -0.2044716 | 1 | -0.28937 | 1 |
| BZW1 | 0.31891977 | 1 | -0.81264 | 1 |
| OR2A5 | -0.0385617 | 1 | -0.45486 | 1 |
| OR52H1 | -0.0258607 | 1 | -0.46713 | 1 |
| OR11H2 | -0.4365357 | 1 | -0.05385 | 1 |
| ZDHHC17 | -0.0901983 | 1 | -0.39994 | 1 |
| OR7G3 | -0.2581761 | 1 | -0.22934 | 1 |
| ZNF800 | -0.1418245 | 1 | -0.34341 | 1 |
| OR52I1 | -0.3429224 | 1 | -0.14143 | 1 |
| RABGAP1L | -0.1921061 | 1 | -0.29101 | 1 |
| OR10V1 | -0.4450431 | 1 | -0.03736 | 1 |
| GPR34 | -0.5490508 | 1 | 0.067549 | 1 |
| PARD3 | -0.219778 | 1 | -0.26116 | 1 |
| SSX7 | -0.6529319 | 1 | 0.172184 | 1 |
| RAP1GDS1 | 0.10492845 | 1 | -0.58541 | 1 |
| SAP30 | -0.4153888 | 1 | -0.06472 | 1 |
| ZNF558 | -0.268768 | 1 | -0.21106 | 1 |
| MTDH | -0.0710002 | 1 | -0.40864 | 1 |
| OVOL1 | -0.4590587 | 1 | -0.02021 | 1 |
| ZNF354C | -0.1889551 | 1 | -0.29014 | 1 |
| TEC | -0.2783995 | 1 | -0.20008 | 1 |
| GRPR | -0.2851209 | 1 | -0.19323 | 1 |
| ARF6 | 0.06576306 | 1 | -0.54381 | 1 |
| THOC1 | -0.0563272 | 1 | -0.42163 | 1 |
| ZNF2 | 0.17153875 | 1 | -0.64947 | 1 |
| FZD3 | -0.1289383 | 1 | -0.34879 | 1 |

| | | | | |
|---|---|---|---|---|
| IQSEC1 | -1.2854401 | 1 | 0.811753 | 1 |
| ADCY1 | -0.4323857 | 1 | -0.04104 | 1 |
| VGLL2 | -0.4986974 | 1 | 0.025992 | 1 |
| KLRK1 | -0.2170692 | 1 | -0.2553 | 1 |
| ZNF845 | -0.0842778 | 1 | -0.38798 | 1 |
| AKAP3 | -0.2876867 | 1 | -0.1844 | 1 |
| ZDHHC22 | -0.6592191 | 1 | 0.188308 | 1 |
| ZNF529 | 0.54970196 | 1 | -1.01987 | 1 |
| RAB27B | -0.2475254 | 1 | -0.22245 | 1 |
| EPHB1 | -1.0238003 | 1 | 0.554735 | 1 |
| PDE7B | -1.0655628 | 1 | 0.596756 | 1 |
| OR1C1 | -0.159826 | 1 | -0.30802 | 1 |
| ZNF713 | -0.3301741 | 1 | -0.13696 | 1 |
| EPHB4 | -0.2135378 | 1 | -0.2529 | 1 |
| PKNOX1 | -0.1943091 | 1 | -0.27136 | 1 |
| MEF2C | 0.22670919 | 1 | -0.69204 | 1 |
| MYSM1 | -0.0658636 | 1 | -0.39796 | 1 |
| ZMYM3 | -0.3311978 | 1 | -0.13202 | 1 |
| PAX6 | -1.6491845 | 1 | 1.186201 | 1 |
| HSF2 | -0.2052048 | 1 | -0.25692 | 1 |
| GRB14 | -0.1581337 | 1 | -0.30391 | 1 |
| ABR | -0.2469794 | 1 | -0.21267 | 1 |
| LATS1 | 0.02350998 | 1 | -0.48149 | 1 |
| IMPA1 | -0.1075653 | 1 | -0.34949 | 1 |
| SUPT5H | -0.4769765 | 1 | 0.021077 | 1 |
| ZNF79 | -0.0037289 | 1 | -0.45215 | 1 |
| SHB | -0.5500891 | 1 | 0.094258 | 1 |
| ZNF799 | -0.0590322 | 1 | -0.3963 | 1 |
| DEF8 | -0.7500898 | 1 | 0.295931 | 1 |
| FGF18 | -0.009987 | 1 | -0.44304 | 1 |
| AVPR1A | -0.3820847 | 1 | -0.07028 | 1 |
| LINGO1 | -0.5494912 | 1 | 0.097247 | 1 |
| CTCF | 0.47560568 | 1 | -0.92694 | 1 |
| ZNF827 | -0.3560893 | 1 | -0.095 | 1 |
| TACR3 | -0.2393843 | 1 | -0.21032 | 1 |
| ITGA8 | 0.58487405 | 1 | -1.03404 | 1 |
| GNG12 | 0.10597822 | 1 | -0.555 | 1 |
| CDK12 | -0.1620434 | 1 | -0.28677 | 1 |
| ZCCHC7 | 0.0057812 | 1 | -0.45396 | 1 |
| DIRAS3 | -0.2330375 | 1 | -0.21484 | 1 |
| CEBPB | -0.2654331 | 1 | -0.18223 | 1 |
| RGS6 | 0.00517425 | 1 | -0.45213 | 1 |
| GNG2 | -0.1822677 | 1 | -0.26457 | 1 |
| GPR182 | -0.2831752 | 1 | -0.16357 | 1 |
| FOXN2 | -0.2617488 | 1 | -0.18451 | 1 |
| HOXD12 | -0.0397363 | 1 | -0.40597 | 1 |

| | | | | |
|---|---|---|---|---|
| CDH1 | -0.1626965 | 1 | -0.28285 | 1 |
| ZNF160 | -0.2082776 | 1 | -0.23712 | 1 |
| DGKI | -0.2397465 | 1 | -0.20516 | 1 |
| CABP2 | -0.0189536 | 1 | -0.42432 | 1 |
| P2RX5 | -0.2076736 | 1 | -0.23557 | 1 |
| ARNT2 | -2.4032201 | 1 | 1.96027 | 1 |
| OR10T2 | -0.3921246 | 1 | -0.05072 | 1 |
| TBPL1 | 0.06173067 | 1 | -0.50447 | 1 |
| BCL6 | -0.2643802 | 1 | -0.1783 | 1 |
| TRIB2 | 0.13348769 | 1 | -0.57558 | 1 |
| MAPKAPK5 | -0.2017299 | 1 | -0.24015 | 1 |
| DKK1 | 0.30222042 | 1 | -0.74373 | 1 |
| CAMK2B | -0.8999135 | 1 | 0.459307 | 1 |
| RXFP2 | -1.001198 | 1 | 0.560756 | 1 |
| MAPK9 | -0.4365332 | 1 | -0.00289 | 1 |
| HEY1 | -0.0882793 | 1 | -0.35092 | 1 |
| BSX | -2.2739193 | 1 | 1.835172 | 1 |
| TCEANC2 | -0.2309474 | 1 | -0.20685 | 1 |
| PCSK9 | -0.0685231 | 1 | -0.36861 | 1 |
| DUOX1 | -0.3467383 | 1 | -0.09013 | 1 |
| PRKCB | -0.020871 | 1 | -0.41563 | 1 |
| FZD1 | -0.6548356 | 1 | 0.21847 | 1 |
| SPOCK3 | -0.779792 | 1 | 0.34371 | 1 |
| OR10A7 | -0.1156632 | 1 | -0.31972 | 1 |
| SP1 | -0.1858051 | 1 | -0.24914 | 1 |
| OR52I2 | -0.465355 | 1 | 0.031741 | 1 |
| NLRP6 | -0.1507061 | 1 | -0.2825 | 1 |
| HOXA9 | 0.05771909 | 1 | -0.48982 | 1 |
| GABRA4 | -0.2186784 | 1 | -0.21333 | 1 |
| HMGB1 | -0.1069778 | 1 | -0.32402 | 1 |
| ZFP28 | -0.1686962 | 1 | -0.2619 | 1 |
| PIM1 | -0.2300166 | 1 | -0.20021 | 1 |
| PTPRG | -0.06766 | 1 | -0.36171 | 1 |
| PTGER1 | -0.3961128 | 1 | -0.03276 | 1 |
| ZNF185 | -0.2397315 | 1 | -0.18894 | 1 |
| HDAC2 | -0.0410108 | 1 | -0.38587 | 1 |
| TCFL5 | -0.2454644 | 1 | -0.1813 | 1 |
| OR4A5 | -0.3700713 | 1 | -0.05597 | 1 |
| ADGRB2 | -0.4545837 | 1 | 0.029326 | 1 |
| GPR50 | 0.28052419 | 1 | -0.70475 | 1 |
| DLX2 | -0.2351271 | 1 | -0.18886 | 1 |
| ZNF510 | -0.192502 | 1 | -0.23132 | 1 |
| OR5V1 | -0.0492033 | 1 | -0.37342 | 1 |
| ZNF195 | -0.1904302 | 1 | -0.23189 | 1 |
| ZNF354B | 0.09010576 | 1 | -0.51088 | 1 |
| RGS7BP | -0.2979129 | 1 | -0.12174 | 1 |

| | | | | |
|---|---|---|---|---|
| GRHL1 | -0.1544573 | 1 | -0.26499 | 1 |
| NFAT5 | -0.1735314 | 1 | -0.24482 | 1 |
| LDB2 | -0.1799275 | 1 | -0.23837 | 1 |
| MRGPRX4 | -0.4269584 | 1 | 0.008966 | 1 |
| CSF1 | -1.3499591 | 1 | 0.932842 | 1 |
| ZDHHC6 | -0.0341226 | 1 | -0.38218 | 1 |
| OR51A7 | 0.10883787 | 1 | -0.5251 | 1 |
| GPR158 | -0.2607348 | 1 | -0.1555 | 1 |
| OR7C1 | -0.1288995 | 1 | -0.28716 | 1 |
| OR2AP1 | -0.1684709 | 1 | -0.24753 | 1 |
| KNDC1 | -0.5836372 | 1 | 0.168267 | 1 |
| CDKN2A | 0.78028886 | 1 | -1.19554 | 1 |
| TERF2 | -0.0689888 | 1 | -0.34562 | 1 |
| PGBD1 | -0.0426766 | 1 | -0.37071 | 1 |
| BAZ1B | -0.1947136 | 1 | -0.21843 | 1 |
| RASA1 | -0.0163504 | 1 | -0.39673 | 1 |
| SMARCA1 | -0.1602344 | 1 | -0.25226 | 1 |
| ZNF792 | -0.1931078 | 1 | -0.219 | 1 |
| P2RX7 | -0.7113871 | 1 | 0.299638 | 1 |
| LPIN2 | -0.2741415 | 1 | -0.13729 | 1 |
| RGS13 | -0.148785 | 1 | -0.26251 | 1 |
| NMRAL1 | -0.294753 | 1 | -0.11507 | 1 |
| MCTS1 | -0.0830796 | 1 | -0.3259 | 1 |
| RCAN2 | -0.6187058 | 1 | 0.211633 | 1 |
| ZNF304 | -0.0804117 | 1 | -0.32564 | 1 |
| FGF12 | -0.0711243 | 1 | -0.33425 | 1 |
| ZNF585A | -0.1327912 | 1 | -0.27241 | 1 |
| GLI2 | -0.1793768 | 1 | -0.22401 | 1 |
| DCC | -1.1908274 | 1 | 0.787831 | 1 |
| OR2T1 | -0.3276672 | 1 | -0.07525 | 1 |
| GRM5 | -0.6008764 | 1 | 0.198402 | 1 |
| H2AC6 | -0.200897 | 1 | -0.20113 | 1 |
| OXGR1 | -0.2744306 | 1 | -0.12655 | 1 |
| PAWR | -0.1991713 | 1 | -0.20176 | 1 |
| ZNF709 | -0.1677592 | 1 | -0.2329 | 1 |
| ARFGEF2 | -0.167831 | 1 | -0.23272 | 1 |
| H1.4 | -0.2157779 | 1 | -0.18466 | 1 |
| SMAD7 | -0.4227873 | 1 | 0.025335 | 1 |
| ZNF611 | -0.1611887 | 1 | -0.23624 | 1 |
| PTGFR | -0.1579395 | 1 | -0.23924 | 1 |
| ZFPM2 | -0.2324565 | 1 | -0.16459 | 1 |
| ZNF146 | -0.0155597 | 1 | -0.38138 | 1 |
| NDEL1 | -0.3744019 | 1 | -0.02254 | 1 |
| OR5H15 | -0.3149034 | 1 | -0.08152 | 1 |
| POLD1 | -0.2293369 | 1 | -0.16625 | 1 |
| ECEL1 | -0.1984546 | 1 | -0.1971 | 1 |

| | | | | |
|---|---|---|---|---|
| ITGB3BP | -0.1196035 | 1 | -0.27565 | 1 |
| GRIN3A | -0.2569976 | 1 | -0.13794 | 1 |
| ADGRE2 | -0.3036192 | 1 | -0.09093 | 1 |
| PPP2R5E | 1.07716799 | 1 | -1.47034 | 1 |
| CUX1 | -0.1923527 | 1 | -0.20073 | 1 |
| IRS1 | -0.209971 | 1 | -0.18153 | 1 |
| FANCL | -0.0279638 | 1 | -0.3633 | 1 |
| SCG2 | -0.1913319 | 1 | -0.19824 | 1 |
| MYCL | -2.3850106 | 1 | 1.995486 | 1 |
| GPR84 | 7.94E-05 | 1 | -0.38947 | 1 |
| FHL2 | -0.0522475 | 1 | -0.33508 | 1 |
| OR56A3 | -0.1735026 | 1 | -0.21376 | 1 |
| PKD2 | -0.1767912 | 1 | -0.21021 | 1 |
| NEUROG1 | -0.4132834 | 1 | 0.026742 | 1 |
| ZDHHC11B | -0.4848232 | 1 | 0.098336 | 1 |
| SOS2 | -0.2346826 | 1 | -0.1514 | 1 |
| PRKCG | -0.195093 | 1 | -0.18948 | 1 |
| ZNF157 | -0.1156462 | 1 | -0.26854 | 1 |
| MTA3 | -0.2469625 | 1 | -0.13715 | 1 |
| CHN1 | -0.0946953 | 1 | -0.28903 | 1 |
| PLRG1 | -0.1744828 | 1 | -0.20728 | 1 |
| ZFP36L1 | -0.1427026 | 1 | -0.23869 | 1 |
| F2RL1 | -0.0873568 | 1 | -0.29355 | 1 |
| FGF13 | -0.2088462 | 1 | -0.17181 | 1 |
| HELT | -0.7511663 | 1 | 0.37106 | 1 |
| SOCS6 | -0.1142382 | 1 | -0.26578 | 1 |
| KCNH6 | -0.3917626 | 1 | 0.012911 | 1 |
| ZNF566 | -0.0181548 | 1 | -0.36059 | 1 |
| ADIPOR1 | -0.2404218 | 1 | -0.13801 | 1 |
| ZNF318 | -0.3185273 | 1 | -0.05979 | 1 |
| IL1RAPL1 | -0.5860378 | 1 | 0.208393 | 1 |
| HBP1 | -0.0241983 | 1 | -0.35337 | 1 |
| MEAF6 | 0.06215626 | 1 | -0.43961 | 1 |
| RASSF10 | -1.1518051 | 1 | 0.774628 | 1 |
| OR4E2 | -0.2102357 | 1 | -0.16634 | 1 |
| NET1 | -0.0402561 | 1 | -0.33622 | 1 |
| ATIC | -0.1092524 | 1 | -0.2671 | 1 |
| ZZZ3 | -0.1219732 | 1 | -0.25433 | 1 |
| GPR18 | -0.2779655 | 1 | -0.09832 | 1 |
| BRS3 | -0.352058 | 1 | -0.02375 | 1 |
| UNC5A | -0.3154588 | 1 | -0.06026 | 1 |
| ZBTB18 | -0.2266478 | 1 | -0.14815 | 1 |
| CORO1C | -0.1738085 | 1 | -0.20066 | 1 |
| TRAF3 | -0.5002818 | 1 | 0.126033 | 1 |
| ZFP3 | 0.02327852 | 1 | -0.39707 | 1 |
| ETV1 | 0.09906929 | 1 | -0.47152 | 1 |

| | | | | |
|---|---|---|---|---|
| OPRD1 | -0.3668944 | 1 | -0.00542 | 1 |
| CBLB | -0.1523868 | 1 | -0.21962 | 1 |
| RAB40A | -0.1801242 | 1 | -0.18902 | 1 |
| CEP43 | -0.1394558 | 1 | -0.22938 | 1 |
| GNG4 | -0.2215612 | 1 | -0.14683 | 1 |
| KDM5A | 0.11079162 | 1 | -0.47892 | 1 |
| ARFGEF1 | 0.03080247 | 1 | -0.39889 | 1 |
| CCDC59 | 1.04429044 | 1 | -1.41127 | 1 |
| OR4P4 | 0.01758308 | 1 | -0.3844 | 1 |
| SHH | -0.1939215 | 1 | -0.17278 | 1 |
| ZNF236 | -0.2979322 | 1 | -0.06848 | 1 |
| NR0B1 | -0.2365405 | 1 | -0.12919 | 1 |
| OTP | -0.2793233 | 1 | -0.08484 | 1 |
| PRKACB | -0.0947579 | 1 | -0.26929 | 1 |
| S100A11 | -0.1078039 | 1 | -0.256 | 1 |
| ZNF397 | -0.3122908 | 1 | -0.05105 | 1 |
| DERL1 | 0.32454615 | 1 | -0.68739 | 1 |
| PPP2CA | 0.05495581 | 1 | -0.4168 | 1 |
| DNTTIP2 | 0.79200339 | 1 | -1.15376 | 1 |
| GFRA3 | -0.1946862 | 1 | -0.16577 | 1 |
| KLF7 | -0.1187647 | 1 | -0.24099 | 1 |
| ZNF565 | -0.1607865 | 1 | -0.19774 | 1 |
| ARHGAP5 | -0.2364457 | 1 | -0.12028 | 1 |
| BAG4 | 1.04095793 | 1 | -1.39583 | 1 |
| PPBP | -0.0521381 | 1 | -0.30167 | 1 |
| ARHGEF2 | -0.2063645 | 1 | -0.14743 | 1 |
| MAP3K13 | -0.07437 | 1 | -0.27864 | 1 |
| GPR22 | -0.162543 | 1 | -0.19019 | 1 |
| LEUTX | -0.1014327 | 1 | -0.25113 | 1 |
| CERK | -0.1566832 | 1 | -0.19533 | 1 |
| TFAP2B | 0.03646672 | 1 | -0.38839 | 1 |
| OR1Q1 | -0.2549986 | 1 | -0.09668 | 1 |
| NFATC3 | -0.1893372 | 1 | -0.16104 | 1 |
| ANK3 | -0.1295828 | 1 | -0.22012 | 1 |
| RIT1 | -0.0196577 | 1 | -0.32891 | 1 |
| ZNF623 | 0.28925162 | 1 | -0.6369 | 1 |
| FOXB1 | -0.0463898 | 1 | -0.30063 | 1 |
| HCAR3 | -0.2293226 | 1 | -0.11679 | 1 |
| ZNF776 | -0.1012658 | 1 | -0.24473 | 1 |
| ZBTB1 | 0.67304276 | 1 | -1.01871 | 1 |
| CERS3 | -0.1974063 | 1 | -0.1481 | 1 |
| CCNK | -0.0806231 | 1 | -0.26426 | 1 |
| GNRHR | -0.0176826 | 1 | -0.32614 | 1 |
| F2R | -0.0084499 | 1 | -0.33427 | 1 |
| ZSCAN1 | -0.7988465 | 1 | 0.456576 | 1 |
| ASAP2 | -0.087966 | 1 | -0.25417 | 1 |

| | | | | |
|---|---|---|---|---|
| OR14J1 | -0.1316204 | 1 | -0.21028 | 1 |
| OR6Q1 | -0.3007704 | 1 | -0.04085 | 1 |
| OR2H2 | -0.1561371 | 1 | -0.18511 | 1 |
| PLCZ1 | -0.3031138 | 1 | -0.03766 | 1 |
| DLX6 | -0.3278558 | 1 | -0.01277 | 1 |
| ZNF705A | -0.0561727 | 1 | -0.28372 | 1 |
| TP63 | -0.2486539 | 1 | -0.09083 | 1 |
| SPSB1 | -0.1861532 | 1 | -0.15222 | 1 |
| NPR3 | -0.1746797 | 1 | -0.16357 | 1 |
| TERT | -0.2170596 | 1 | -0.12115 | 1 |
| CHRM4 | -0.2416342 | 1 | -0.09619 | 1 |
| BNC1 | -0.1723391 | 1 | -0.16471 | 1 |
| FPR1 | -0.0821603 | 1 | -0.25407 | 1 |
| GLRB | 0.02987564 | 1 | -0.36559 | 1 |
| ZNF471 | -0.752422 | 1 | 0.417155 | 1 |
| ICAM1 | 0.00058667 | 1 | -0.3358 | 1 |
| REST | 0.09217335 | 1 | -0.42662 | 1 |
| IFNGR1 | 0.01056969 | 1 | -0.3449 | 1 |
| ZNF786 | -0.1317844 | 1 | -0.20251 | 1 |
| TLK2 | 0.03202136 | 1 | -0.36625 | 1 |
| OR1J4 | -0.1348805 | 1 | -0.19931 | 1 |
| KDR | -0.2183761 | 1 | -0.11528 | 1 |
| TBCK | -0.1682965 | 1 | -0.16531 | 1 |
| EIF4A2 | 0.05659609 | 1 | -0.3901 | 1 |
| ZNF404 | 0.31368152 | 1 | -0.64678 | 1 |
| NEK4 | -0.1044529 | 1 | -0.22843 | 1 |
| PKP1 | -0.2378335 | 1 | -0.09448 | 1 |
| NCOA4 | -0.0384817 | 1 | -0.2934 | 1 |
| GID4 | -0.1825694 | 1 | -0.14883 | 1 |
| SRGAP2 | -0.381545 | 1 | 0.052154 | 1 |
| CRX | -0.2620282 | 1 | -0.067 | 1 |
| OR6C65 | -0.4293299 | 1 | 0.100532 | 1 |
| POU6F2 | -0.1709701 | 1 | -0.15743 | 1 |
| ZDHHC24 | -0.5913849 | 1 | 0.263833 | 1 |
| TARDBP | 0.11072138 | 1 | -0.43824 | 1 |
| RAB5A | 0.07322354 | 1 | -0.40066 | 1 |
| DNASE1L3 | -0.1843771 | 1 | -0.14283 | 1 |
| GLP1R | -0.1341264 | 1 | -0.19287 | 1 |
| CHRNA7 | -0.2020629 | 1 | -0.12466 | 1 |
| IL13RA2 | -0.1299238 | 1 | -0.19518 | 1 |
| KIR2DS5 | 0.09324674 | 1 | -0.41829 | 1 |
| OR6C2 | -0.1009919 | 1 | -0.2237 | 1 |
| TNP1 | -0.1991885 | 1 | -0.12425 | 1 |
| ELOVL4 | -0.0019506 | 1 | -0.32057 | 1 |
| RASD2 | 0.36512998 | 1 | -0.68733 | 1 |
| BASP1 | -0.1146526 | 1 | -0.20748 | 1 |

| | | | | |
|---|---|---|---|---|
| ZNF562 | -0.0502995 | 1 | -0.27129 | 1 |
| PASK | -0.1349564 | 1 | -0.18591 | 1 |
| ZC3H11A | -0.1242113 | 1 | -0.19653 | 1 |
| TDP2 | 0.05631559 | 1 | -0.37501 | 1 |
| PRKG2 | -1.7621199 | 1 | 1.443516 | 1 |
| ZNF782 | -0.0712251 | 1 | -0.2458 | 1 |
| GHR | -0.3830244 | 1 | 0.066015 | 1 |
| GABPB2 | -0.4751017 | 1 | 0.158201 | 1 |
| CSRNP1 | -0.1140844 | 1 | -0.20194 | 1 |
| TAS2R30 | -0.1602211 | 1 | -0.15566 | 1 |
| IL5 | 0.31612382 | 1 | -0.63127 | 1 |
| SOCS7 | -0.2219893 | 1 | -0.09311 | 1 |
| GP1BA | -0.481173 | 1 | 0.16628 | 1 |
| RFXAP | -0.0814854 | 1 | -0.23258 | 1 |
| VIPR2 | -0.3920373 | 1 | 0.078277 | 1 |
| IRF6 | -0.6043704 | 1 | 0.290988 | 1 |
| SSTR2 | -0.1702016 | 1 | -0.1418 | 1 |
| SMAD1 | -0.092236 | 1 | -0.21925 | 1 |
| TAS2R10 | -0.1147842 | 1 | -0.19628 | 1 |
| NELFCD | -0.1158179 | 1 | -0.19508 | 1 |
| EPC1 | 0.32240844 | 1 | -0.63289 | 1 |
| MRTFA | -0.1711621 | 1 | -0.13902 | 1 |
| KLF10 | -0.1278165 | 1 | -0.18178 | 1 |
| NRF1 | -0.2191173 | 1 | -0.08971 | 1 |
| DCLK1 | -0.2226435 | 1 | -0.08578 | 1 |
| ITGB6 | 0.04756333 | 1 | -0.35556 | 1 |
| KIFAP3 | -0.1067 | 1 | -0.20115 | 1 |
| BAZ1A | 0.02766908 | 1 | -0.33552 | 1 |
| GHRH | -0.2114263 | 1 | -0.09625 | 1 |
| NKX1.1 | -0.0359742 | 1 | -0.27167 | 1 |
| CSNK2A1 | 0.32697052 | 1 | -0.6346 | 1 |
| SATB2 | -0.7708786 | 1 | 0.463923 | 1 |
| ROBO2 | -0.0533921 | 1 | -0.25352 | 1 |
| CSF2RB | -0.2394457 | 1 | -0.06731 | 1 |
| KDM5C | -0.0071449 | 1 | -0.29937 | 1 |
| ITGB5 | -0.1684221 | 1 | -0.13778 | 1 |
| ZNF559 | -0.0546185 | 1 | -0.25072 | 1 |
| RAB9A | -0.0771752 | 1 | -0.22707 | 1 |
| DMTF1 | -0.151265 | 1 | -0.15281 | 1 |
| MORC2 | -0.1658841 | 1 | -0.1379 | 1 |
| ZNHIT6 | 0.00457085 | 1 | -0.30815 | 1 |
| TRAFD1 | -1.0563877 | 1 | 0.75331 | 1 |
| ZDHHC5 | 0.00607705 | 1 | -0.30912 | 1 |
| ZNF717 | -0.1839514 | 1 | -0.11896 | 1 |
| AKT1 | -0.207656 | 1 | -0.09202 | 1 |
| OR10Q1 | 0.02260219 | 1 | -0.32225 | 1 |

| | | | | |
|---|---|---|---|---|
| SOCS4 | -0.0398691 | 1 | -0.25957 | 1 |
| PDE6A | -0.2267311 | 1 | -0.07177 | 1 |
| SRGAP3 | -2.5226371 | 1 | 2.224243 | 1 |
| STK26 | -0.2106049 | 1 | -0.08711 | 1 |
| TNFRSF4 | -0.1663321 | 1 | -0.13093 | 1 |
| GPR160 | 0.09541262 | 1 | -0.39252 | 1 |
| TSC22D1 | -0.1857732 | 1 | -0.11123 | 1 |
| NSD1 | -0.1603625 | 1 | -0.13649 | 1 |
| CAT | 0.84052235 | 1 | -1.13683 | 1 |
| DMRTC1 | -1.8612246 | 1 | 1.565289 | 1 |
| ZNF347 | -0.0932355 | 1 | -0.20126 | 1 |
| TNFRSF1B | -0.2311356 | 1 | -0.06331 | 1 |
| FOXS1 | -0.0244109 | 1 | -0.2699 | 1 |
| RPS6KB1 | 0.1383421 | 1 | -0.43225 | 1 |
| TRAPPC2 | -0.0999982 | 1 | -0.19387 | 1 |
| OR5AC2 | -0.3269109 | 1 | 0.033819 | 1 |
| MSS51 | -0.0560444 | 1 | -0.23686 | 1 |
| OR6C70 | -0.3094867 | 1 | 0.017387 | 1 |
| KRBOX4 | 0.15884984 | 1 | -0.45086 | 1 |
| PRPF6 | 0.03054679 | 1 | -0.32243 | 1 |
| LBX2 | -0.1733611 | 1 | -0.1183 | 1 |
| ZNF233 | -0.1409896 | 1 | -0.14973 | 1 |
| DGKK | -0.3229209 | 1 | 0.032432 | 1 |
| CAP2 | -0.0693212 | 1 | -0.22074 | 1 |
| AKIRIN2 | -0.0251659 | 1 | -0.26482 | 1 |
| CLU | -0.1432841 | 1 | -0.14614 | 1 |
| MYOG | -0.0764403 | 1 | -0.21254 | 1 |
| NPAS4 | -0.0607592 | 1 | -0.2282 | 1 |
| INVS | 0.06245052 | 1 | -0.35096 | 1 |
| POU3F2 | -0.1173676 | 1 | -0.17091 | 1 |
| C5AR2 | -0.193187 | 1 | -0.09493 | 1 |
| POGZ | -0.0018399 | 1 | -0.28622 | 1 |
| TENM3 | -0.4371509 | 1 | 0.149372 | 1 |
| ACSL6 | -0.213647 | 1 | -0.07368 | 1 |
| USP9Y | -0.2120982 | 1 | -0.07475 | 1 |
| DLX1 | -0.1449909 | 1 | -0.14047 | 1 |
| ZNF430 | -0.0229852 | 1 | -0.26242 | 1 |
| MS4A10 | -0.1336393 | 1 | -0.15136 | 1 |
| NSMCE3 | -0.1485151 | 1 | -0.13591 | 1 |
| CREBRF | -0.0868279 | 1 | -0.19722 | 1 |
| CDH13 | -0.1379711 | 1 | -0.14419 | 1 |
| ZNF329 | -0.2766824 | 1 | -0.00541 | 1 |
| ONECUT2 | -0.2018515 | 1 | -0.07975 | 1 |
| SPTBN1 | -0.2890328 | 1 | 0.007679 | 1 |
| TBC1D26 | -0.0638786 | 1 | -0.21712 | 1 |
| NTS | -0.3014823 | 1 | 0.020691 | 1 |

| | | | | |
|---|---|---|---|---|
| EPHA5 | -0.1868674 | 1 | -0.0923 | 1 |
| PDK3 | -0.1398709 | 1 | -0.13916 | 1 |
| CCND2 | -0.1206543 | 1 | -0.15814 | 1 |
| NDST1 | -0.3394296 | 1 | 0.060828 | 1 |
| CDYL | -0.1215241 | 1 | -0.15693 | 1 |
| ABCC9 | -0.1337098 | 1 | -0.14446 | 1 |
| ZNF382 | -0.027722 | 1 | -0.25031 | 1 |
| OPN3 | -0.053345 | 1 | -0.22467 | 1 |
| MAP3K21 | -0.1323571 | 1 | -0.1456 | 1 |
| CD86 | 0.1188146 | 1 | -0.39594 | 1 |
| VAPA | -0.0157325 | 1 | -0.26107 | 1 |
| ZNF180 | -0.1608771 | 1 | -0.11583 | 1 |
| NAA16 | 0.00721974 | 1 | -0.28305 | 1 |
| PLXNA3 | -0.2303314 | 1 | -0.04434 | 1 |
| RAB2A | 0.95140805 | 1 | -1.22595 | 1 |
| ZNF100 | 0.07059514 | 1 | -0.34458 | 1 |
| ZNF91 | -0.0342322 | 1 | -0.2394 | 1 |
| NEUROG2 | -0.0319104 | 1 | -0.24142 | 1 |
| IRX5 | -0.3285507 | 1 | 0.055387 | 1 |
| GUCY2F | -0.2607569 | 1 | -0.01226 | 1 |
| CHRNA1 | -0.0039756 | 1 | -0.26856 | 1 |
| OR2T12 | -0.1154016 | 1 | -0.15608 | 1 |
| EPS8 | -0.0079866 | 1 | -0.26337 | 1 |
| ZNF749 | -0.066048 | 1 | -0.20511 | 1 |
| RIC8B | -0.0908723 | 1 | -0.18011 | 1 |
| KMT2E | -0.1144231 | 1 | -0.15642 | 1 |
| ASH2L | 0.02999077 | 1 | -0.30045 | 1 |
| NR2C2 | -0.1432822 | 1 | -0.12689 | 1 |
| IRF2 | -0.087638 | 1 | -0.18253 | 1 |
| TCF7L2 | -0.2174336 | 1 | -0.05191 | 1 |
| ZNF14 | 0.03094868 | 1 | -0.29968 | 1 |
| HDAC6 | -0.4358333 | 1 | 0.167584 | 1 |
| RXRA | -0.4924231 | 1 | 0.224566 | 1 |
| OR8J3 | -0.1936619 | 1 | -0.07373 | 1 |
| CDKL2 | -0.1984121 | 1 | -0.06849 | 1 |
| SIX4 | -0.0453025 | 1 | -0.2214 | 1 |
| ZNF630 | -0.2077385 | 1 | -0.05881 | 1 |
| SSX5 | -0.1426596 | 1 | -0.12366 | 1 |
| MAP2K1 | 0.06674338 | 1 | -0.33295 | 1 |
| ATOH7 | -0.7105073 | 1 | 0.444583 | 1 |
| KLRB1 | 0.03183651 | 1 | -0.29718 | 1 |
| TIPIN | -0.0409737 | 1 | -0.22403 | 1 |
| MDC1 | -0.3245128 | 1 | 0.059676 | 1 |
| BHLHE41 | -0.1986172 | 1 | -0.06605 | 1 |
| FLNB | -0.1287817 | 1 | -0.13582 | 1 |
| OXT | -0.0563061 | 1 | -0.20814 | 1 |

| | | | | |
|---|---|---|---|---|
| HOXA7 | -0.3970014 | 1 | 0.132595 | 1 |
| ITPR1 | -0.1365283 | 1 | -0.12741 | 1 |
| DUSP19 | -0.0109495 | 1 | -0.25209 | 1 |
| ZNF780A | -0.1112463 | 1 | -0.15138 | 1 |
| SND1 | -0.0992532 | 1 | -0.16216 | 1 |
| AKT2 | -0.8493181 | 1 | 0.588305 | 1 |
| RAB33A | -0.4738345 | 1 | 0.213001 | 1 |
| IFT57 | -0.2518099 | 1 | -0.00799 | 1 |
| GKAP1 | -0.7225431 | 1 | 0.463051 | 1 |
| FOXA2 | -0.0530981 | 1 | -0.20599 | 1 |
| MS4A3 | -0.1425872 | 1 | -0.11542 | 1 |
| RTF1 | 0.12969616 | 1 | -0.38727 | 1 |
| ZNF718 | -0.060674 | 1 | -0.1959 | 1 |
| GUCY2C | -0.0267481 | 1 | -0.22944 | 1 |
| NIN | -0.0947228 | 1 | -0.16135 | 1 |
| OPCML | -0.1241522 | 1 | -0.1309 | 1 |
| RNF20 | -0.0649912 | 1 | -0.18991 | 1 |
| MYO1E | 0.60170559 | 1 | -0.85592 | 1 |
| RRH | -0.0128378 | 1 | -0.24077 | 1 |
| NPHP1 | -0.1186137 | 1 | -0.13441 | 1 |
| TRAF4 | -1.8991397 | 1 | 1.646572 | 1 |
| PEDS1.UBE2V1 | -0.0535543 | 1 | -0.19894 | 1 |
| ZNF182 | 0.07591393 | 1 | -0.32738 | 1 |
| YY2 | -0.0416379 | 1 | -0.20934 | 1 |
| DCDC1 | -0.1434418 | 1 | -0.10632 | 1 |
| SRFBP1 | 0.09316705 | 1 | -0.34258 | 1 |
| SH3BGRL | 0.22278893 | 1 | -0.47199 | 1 |
| GMNN | -0.0948278 | 1 | -0.15394 | 1 |
| SNIP1 | 0.09187713 | 1 | -0.34057 | 1 |
| PTGER2 | -0.137226 | 1 | -0.11099 | 1 |
| ZNF177 | -0.1062442 | 1 | -0.14091 | 1 |
| CHD5 | -0.2661637 | 1 | 0.019201 | 1 |
| FGF7 | -0.0573162 | 1 | -0.1882 | 1 |
| OR4D9 | -0.0241832 | 1 | -0.2204 | 1 |
| SLC44A2 | -0.1969354 | 1 | -0.04744 | 1 |
| YY1 | 1.18859728 | 1 | -1.43289 | 1 |
| SPATA13 | -0.1540747 | 1 | -0.08997 | 1 |
| ARID5B | -0.1510578 | 1 | -0.09293 | 1 |
| PIP5K1A | 0.25597907 | 1 | -0.49956 | 1 |
| OR52N2 | -0.3708828 | 1 | 0.127394 | 1 |
| ZNF692 | -0.3499584 | 1 | 0.10666 | 1 |
| ZDHHC4 | 0.37459857 | 1 | -0.61741 | 1 |
| WWTR1 | 0.01725834 | 1 | -0.25981 | 1 |
| CIT | -1.2452094 | 1 | 1.002892 | 1 |
| MAP3K9 | -0.2728551 | 1 | 0.03062 | 1 |
| ZNF324B | -0.2049284 | 1 | -0.03651 | 1 |

| | | | | |
|---|---|---|---|---|
| ADGRB3 | -0.4232794 | 1 | 0.182 | 1 |
| CYSLTR1 | 0.16107033 | 1 | -0.40214 | 1 |
| OR10X1 | -0.0092254 | 1 | -0.23173 | 1 |
| TLR9 | -0.0998502 | 1 | -0.14086 | 1 |
| ANTXR1 | -0.1204192 | 1 | -0.12018 | 1 |
| DDX17 | -0.2164223 | 1 | -0.02415 | 1 |
| NKIRAS1 | 1.23234074 | 1 | -1.47233 | 1 |
| ZIM2 | -0.3616635 | 1 | 0.121769 | 1 |
| PPP1R2 | 0.0216867 | 1 | -0.26129 | 1 |
| TAF6 | -0.1168635 | 1 | -0.12253 | 1 |
| POLR3F | 0.2256406 | 1 | -0.46502 | 1 |
| TRAF5 | 0.01262881 | 1 | -0.25103 | 1 |
| DCLK2 | -0.1024209 | 1 | -0.1356 | 1 |
| ARID3A | -0.0392712 | 1 | -0.19865 | 1 |
| OR2T4 | -0.1478181 | 1 | -0.09008 | 1 |
| PLEKHA1 | -0.1427168 | 1 | -0.09402 | 1 |
| PDCD11 | 0.06242071 | 1 | -0.29877 | 1 |
| MYCBP | 0.0516301 | 1 | -0.28778 | 1 |
| ZNF571 | -0.2019707 | 1 | -0.03388 | 1 |
| FGB | -0.1354054 | 1 | -0.09932 | 1 |
| ASB14 | 1.10564814 | 1 | -1.33934 | 1 |
| ARHGAP44 | -0.1700143 | 1 | -0.06351 | 1 |
| PAK6 | -0.1553431 | 1 | -0.07799 | 1 |
| GRM2 | -0.1766561 | 1 | -0.05648 | 1 |
| MTF1 | -0.0813255 | 1 | -0.15176 | 1 |
| MS4A5 | -0.1249482 | 1 | -0.10814 | 1 |
| OR8U8 | -0.1144208 | 1 | -0.11851 | 1 |
| FASLG | -0.0104493 | 1 | -0.22245 | 1 |
| TAS2R13 | 0.21590582 | 1 | -0.44856 | 1 |
| GSX2 | -0.1937011 | 1 | -0.03895 | 1 |
| OR6C75 | 0.00784162 | 1 | -0.24018 | 1 |
| OR52A5 | -0.0868627 | 1 | -0.14508 | 1 |
| ADGRD1 | -0.1139835 | 1 | -0.11774 | 1 |
| FUS | -0.0691256 | 1 | -0.16201 | 1 |
| PPP2R5A | -0.1525372 | 1 | -0.07793 | 1 |
| MTNR1B | -0.1489762 | 1 | -0.08111 | 1 |
| ZNF322 | 0.03176231 | 1 | -0.2616 | 1 |
| ZBTB34 | -0.0020561 | 1 | -0.22756 | 1 |
| TNFRSF18 | 1.07046194 | 1 | -1.29964 | 1 |
| ZMYM5 | 0.01058885 | 1 | -0.23903 | 1 |
| MYF6 | -0.1338247 | 1 | -0.09423 | 1 |
| OR52E4 | -0.1904478 | 1 | -0.03754 | 1 |
| ZDHHC2 | -0.1443015 | 1 | -0.08356 | 1 |
| ZNF114 | -0.2283121 | 1 | 0.001108 | 1 |
| RBPMS | 0.03984075 | 1 | -0.26703 | 1 |
| FOXJ2 | -0.0741869 | 1 | -0.15185 | 1 |

| | | | | |
|---|---|---|---|---|
| SIPA1L3 | -0.1234187 | 1 | -0.10261 | 1 |
| TLR8 | -0.1491678 | 1 | -0.076 | 1 |
| ZNF283 | -0.1186533 | 1 | -0.10649 | 1 |
| CCL16 | -0.1417726 | 1 | -0.08328 | 1 |
| SCG5 | -1.4148556 | 1 | 1.189936 | 1 |
| ELMO1 | -0.4934323 | 1 | 0.269056 | 1 |
| FOXO3 | -0.0111218 | 1 | -0.21238 | 1 |
| FCER1A | -0.1781402 | 1 | -0.04534 | 1 |
| BAZ2B | -0.2722103 | 1 | 0.048751 | 1 |
| AKAP17A | 0.0090725 | 1 | -0.23231 | 1 |
| PAX3 | 0.47703479 | 1 | -0.69988 | 1 |
| DAPP1 | -0.0898824 | 1 | -0.13264 | 1 |
| HIPK1 | -0.1485272 | 1 | -0.07276 | 1 |
| OR2A7 | 0.09247161 | 1 | -0.31316 | 1 |
| RBBP4 | 0.20432686 | 1 | -0.42496 | 1 |
| SHC3 | -0.1907657 | 1 | -0.027 | 1 |
| DEPDC4 | -0.0989424 | 1 | -0.11874 | 1 |
| RAB3A | -0.1546334 | 1 | -0.06263 | 1 |
| CLP1 | -0.0711634 | 1 | -0.14518 | 1 |
| ATRIP | -0.1273955 | 1 | -0.08872 | 1 |
| PROKR1 | -0.165944 | 1 | -0.05014 | 1 |
| ZNF816 | -0.0439192 | 1 | -0.17198 | 1 |
| TAS2R40 | -0.1363131 | 1 | -0.07951 | 1 |
| BMP3 | -0.1631569 | 1 | -0.0523 | 1 |
| MED12 | -1.2447566 | 1 | 1.029744 | 1 |
| NUDT4 | -0.0383726 | 1 | -0.17637 | 1 |
| ADH1A | -0.1404902 | 1 | -0.07416 | 1 |
| RELA | 0.08366635 | 1 | -0.29775 | 1 |
| OR8D1 | -0.0618538 | 1 | -0.15201 | 1 |
| GTF2E1 | 0.83982896 | 1 | -1.05367 | 1 |
| BCL2L2 | -0.7833389 | 1 | 0.569511 | 1 |
| SIPA1L2 | -0.1467165 | 1 | -0.06705 | 1 |
| ITGA9 | -0.1913953 | 1 | -0.02207 | 1 |
| F7 | -0.2138476 | 1 | 0.000387 | 1 |
| ZFP91 | 0.23029805 | 1 | -0.44357 | 1 |
| SIX6 | -0.082033 | 1 | -0.13107 | 1 |
| ZNF181 | -0.1937126 | 1 | -0.0186 | 1 |
| ECD | 0.01452377 | 1 | -0.22679 | 1 |
| ZNF607 | -0.0049027 | 1 | -0.20713 | 1 |
| CABYR | -0.2106329 | 1 | -0.00131 | 1 |
| PRRX1 | -0.2145671 | 1 | 0.003067 | 1 |
| MYRFL | -0.1460599 | 1 | -0.06534 | 1 |
| KLHL6 | -1.1502806 | 1 | 0.941097 | 1 |
| LYN | -0.0952634 | 1 | -0.1138 | 1 |
| ZNF697 | -0.2509646 | 1 | 0.041935 | 1 |
| LHX2 | -0.0332874 | 1 | -0.1747 | 1 |

| | | | | |
|---|---|---|---|---|
| MYNN | 0.20979438 | 1 | -0.41772 | 1 |
| CDON | -0.1307308 | 1 | -0.07692 | 1 |
| FHIT | -0.1620262 | 1 | -0.04558 | 1 |
| ZNF75D | 0.06492675 | 1 | -0.27232 | 1 |
| ASAH2B | 0.1291019 | 1 | -0.33633 | 1 |
| PDCD4 | -0.0590359 | 1 | -0.14793 | 1 |
| HLA.DRB3 | -0.0749636 | 1 | -0.13129 | 1 |
| TAS2R8 | -0.0957178 | 1 | -0.11051 | 1 |
| OR8G1 | -0.0525462 | 1 | -0.15347 | 1 |
| FCGR1A | -0.2659172 | 1 | 0.060171 | 1 |
| TIGD4 | -0.0702072 | 1 | -0.13418 | 1 |
| FLNA | -0.0919195 | 1 | -0.11204 | 1 |
| FBXW7 | -0.1127325 | 1 | -0.09104 | 1 |
| ZNF214 | 0.01432565 | 1 | -0.21804 | 1 |
| GDF6 | -0.2851957 | 1 | 0.082671 | 1 |
| ADGRF4 | -0.0299417 | 1 | -0.1723 | 1 |
| CAMK4 | -0.1136511 | 1 | -0.08843 | 1 |
| CD70 | -0.0997832 | 1 | -0.10198 | 1 |
| GRM4 | -0.1668607 | 1 | -0.03285 | 1 |
| ZNF226 | -0.0036352 | 1 | -0.19578 | 1 |
| PRKAA2 | 0.0493687 | 1 | -0.24851 | 1 |
| TBC1D25 | 0.96735839 | 1 | -1.16626 | 1 |
| OR6S1 | 0.02217924 | 1 | -0.22082 | 1 |
| ZNF281 | -0.0922232 | 1 | -0.10625 | 1 |
| ADCY10 | -0.0784295 | 1 | -0.12003 | 1 |
| NPBWR2 | -0.0660662 | 1 | -0.13237 | 1 |
| EPHA7 | 0.0023685 | 1 | -0.20022 | 1 |
| WASF2 | -0.145983 | 1 | -0.05147 | 1 |
| IFNK | -0.5775042 | 1 | 0.380074 | 1 |
| NF2 | -0.0098202 | 1 | -0.18583 | 1 |
| SRCAP | -0.1678747 | 1 | -0.0273 | 1 |
| OR4K1 | -0.0633437 | 1 | -0.1311 | 1 |
| ZNF154 | -0.07863 | 1 | -0.11567 | 1 |
| NR4A3 | -0.5842552 | 1 | 0.390011 | 1 |
| GAB2 | -1.0025397 | 1 | 0.808682 | 1 |
| NPAT | 0.17171651 | 1 | -0.36471 | 1 |
| MNDA | 0.02660967 | 1 | -0.21899 | 1 |
| GZMB | -0.1187669 | 1 | -0.0725 | 1 |
| CTBP2 | -0.0300778 | 1 | -0.16077 | 1 |
| LAMTOR3 | 0.20726986 | 1 | -0.39699 | 1 |
| RASA4 | -0.2861652 | 1 | 0.096457 | 1 |
| H4C5 | -0.1213707 | 1 | -0.06829 | 1 |
| MESD | -0.0975526 | 1 | -0.0912 | 1 |
| ZFP14 | -0.0316425 | 1 | -0.15654 | 1 |
| DMBX1 | -0.2421872 | 1 | 0.054845 | 1 |
| NEDD9 | -0.12508 | 1 | -0.06215 | 1 |

| | | | | |
|---|---|---|---|---|
| FOXO1 | -0.1566514 | 1 | -0.03025 | 1 |
| TFCP2L1 | -1.0595488 | 1 | 0.872951 | 1 |
| BARX2 | -0.058844 | 1 | -0.12769 | 1 |
| BMP15 | -0.0169352 | 1 | -0.16942 | 1 |
| RPE65 | -0.0734186 | 1 | -0.11273 | 1 |
| MIDEAS | 0.0076455 | 1 | -0.19336 | 1 |
| OPN4 | -0.7551493 | 1 | 0.569448 | 1 |
| EED | -0.0265556 | 1 | -0.1588 | 1 |
| EDAR | -0.1568308 | 1 | -0.0279 | 1 |
| ZNF416 | 0.14836773 | 1 | -0.33258 | 1 |
| MORC4 | -0.0511467 | 1 | -0.13294 | 1 |
| STAC2 | -0.1786791 | 1 | -0.0054 | 1 |
| LGR6 | -0.0446243 | 1 | -0.13921 | 1 |
| ASIP | -0.1002136 | 1 | -0.08356 | 1 |
| TAAR6 | -0.0516237 | 1 | -0.13198 | 1 |
| OBSCN | -0.1257846 | 1 | -0.0575 | 1 |
| NFRKB | -0.1144769 | 1 | -0.06733 | 1 |
| GPRC5C | -0.0897873 | 1 | -0.09148 | 1 |
| SLC20A1 | 0.80481404 | 1 | -0.98606 | 1 |
| SCML1 | -0.0168096 | 1 | -0.16438 | 1 |
| NEDD4 | -0.0161312 | 1 | -0.16504 | 1 |
| FLRT2 | -0.0358849 | 1 | -0.14524 | 1 |
| ASAP1 | -0.0510559 | 1 | -0.12943 | 1 |
| AMELX | 0.06282163 | 1 | -0.24296 | 1 |
| NOD2 | -0.3940603 | 1 | 0.215224 | 1 |
| MYH9 | 0.8142805 | 1 | -0.99298 | 1 |
| AVPR1B | -0.1328497 | 1 | -0.04575 | 1 |
| TAS2R19 | 0.02887278 | 1 | -0.20597 | 1 |
| SEPTIN5 | -0.1205819 | 1 | -0.0565 | 1 |
| PLXNA2 | -0.0622364 | 1 | -0.11442 | 1 |
| GPR155 | -0.0471157 | 1 | -0.12906 | 1 |
| ZNF420 | -0.058352 | 1 | -0.11751 | 1 |
| ZIC1 | -0.0882705 | 1 | -0.08719 | 1 |
| NPR1 | 0.18345676 | 1 | -0.35864 | 1 |
| ZNF317 | 0.04517311 | 1 | -0.22006 | 1 |
| FSHR | 0.01087877 | 1 | -0.18576 | 1 |
| ZMYND12 | -0.6756212 | 1 | 0.501101 | 1 |
| ZC3H12B | -0.3220508 | 1 | 0.148106 | 1 |
| RBM15 | 0.41015506 | 1 | -0.58393 | 1 |
| NFAM1 | -0.0855589 | 1 | -0.0882 | 1 |
| ERBB2 | -0.2069725 | 1 | 0.034394 | 1 |
| ZC3H15 | 0.38346623 | 1 | -0.55496 | 1 |
| GPR162 | -0.5248448 | 1 | 0.353922 | 1 |
| MSX1 | 0.04725544 | 1 | -0.21791 | 1 |
| CRABP2 | -0.2078781 | 1 | 0.037856 | 1 |
| TLK1 | 0.11323428 | 1 | -0.28311 | 1 |

| | | | | |
|---|---|---|---|---|
| GNG10 | 0.15502335 | 1 | -0.32321 | 1 |
| TMOD2 | -0.2200992 | 1 | 0.052226 | 1 |
| SREBF1 | -0.2819531 | 1 | 0.114999 | 1 |
| ZNF649 | -0.215821 | 1 | 0.049596 | 1 |
| CREG1 | -0.012288 | 1 | -0.1537 | 1 |
| SDCBP2 | -0.1234271 | 1 | -0.04255 | 1 |
| TBC1D9 | -0.0232042 | 1 | -0.14256 | 1 |
| CALCRL | 0.00664884 | 1 | -0.17211 | 1 |
| OR6C6 | -0.1105706 | 1 | -0.05396 | 1 |
| NMUR1 | -0.1556019 | 1 | -0.00892 | 1 |
| FGA | -0.0832154 | 1 | -0.08034 | 1 |
| NOD1 | -0.1250153 | 1 | -0.03788 | 1 |
| OR7A5 | -0.5069275 | 1 | 0.344609 | 1 |
| OR1L6 | -0.1316015 | 1 | -0.02956 | 1 |
| BCL2L10 | 0.0162969 | 1 | -0.17733 | 1 |
| CCL26 | 0.11848415 | 1 | -0.27945 | 1 |
| OR2W3 | -0.1204422 | 1 | -0.0396 | 1 |
| MYO9A | 0.07607383 | 1 | -0.23587 | 1 |
| OR1N2 | -0.0585929 | 1 | -0.1012 | 1 |
| ERN2 | -0.1513748 | 1 | -0.00811 | 1 |
| GABRB2 | -0.0150521 | 1 | -0.14422 | 1 |
| HHEX | -0.088601 | 1 | -0.07066 | 1 |
| OR10H5 | -0.1795164 | 1 | 0.020269 | 1 |
| ZNF790 | -0.0990611 | 1 | -0.06011 | 1 |
| SALL4 | -0.0795728 | 1 | -0.07947 | 1 |
| TRIM29 | -0.0765848 | 1 | -0.08237 | 1 |
| ZZEF1 | -0.017547 | 1 | -0.14141 | 1 |
| PPP1R1A | -0.1711668 | 1 | 0.012342 | 1 |
| PAX9 | -0.0845208 | 1 | -0.07399 | 1 |
| ELK1 | 0.92858447 | 1 | -1.08655 | 1 |
| CHIC2 | -0.1581963 | 1 | 0.000431 | 1 |
| OR5M10 | -0.0886504 | 1 | -0.06833 | 1 |
| STARD8 | -0.0978808 | 1 | -0.05877 | 1 |
| NOTCH1 | -0.3094535 | 1 | 0.15299 | 1 |
| LBP | -0.0715353 | 1 | -0.08397 | 1 |
| TXN | 0.10539479 | 1 | -0.2602 | 1 |
| RABL3 | 0.05814992 | 1 | -0.21257 | 1 |
| OR5AN1 | -0.0611051 | 1 | -0.09228 | 1 |
| TLR4 | -0.138929 | 1 | -0.01401 | 1 |
| TCEAL7 | 0.67894355 | 1 | -0.83183 | 1 |
| C5AR1 | 0.09301982 | 1 | -0.24514 | 1 |
| STK10 | -1.2049025 | 1 | 1.053415 | 1 |
| ELOA2 | -0.1171906 | 1 | -0.03405 | 1 |
| FRS3 | -0.3418024 | 1 | 0.191433 | 1 |
| DNAJA1 | 0.02264023 | 1 | -0.17281 | 1 |
| AIDA | 0.1178723 | 1 | -0.26739 | 1 |

| | | | | |
|---|---|---|---|---|
| FOXD4 | -0.0861537 | 1 | -0.06265 | 1 |
| TFEC | -0.0278672 | 1 | -0.12079 | 1 |
| KMT5B | 0.00355759 | 1 | -0.15152 | 1 |
| NANOG | -0.1844662 | 1 | 0.036965 | 1 |
| ZNF618 | -0.0787304 | 1 | -0.06824 | 1 |
| PPP1R12B | -0.2345473 | 1 | 0.087776 | 1 |
| HRH1 | 1.32490239 | 1 | -1.47148 | 1 |
| OR2G3 | -0.1113697 | 1 | -0.03495 | 1 |
| RAB41 | -0.0783014 | 1 | -0.06734 | 1 |
| DUOX2 | -0.0658828 | 1 | -0.07926 | 1 |
| MAS1L | -0.0374667 | 1 | -0.10764 | 1 |
| HOXB8 | -0.2340688 | 1 | 0.08949 | 1 |
| ITPR3 | 0.00142896 | 1 | -0.14573 | 1 |
| NFE2L1 | -0.2525006 | 1 | 0.108335 | 1 |
| NA. | -0.1413187 | 1 | -0.00284 | 1 |
| ZNF592 | -0.1316774 | 1 | -0.01242 | 1 |
| RTL4 | -0.1213306 | 1 | -0.02247 | 1 |
| ZNF112 | 0.03175892 | 1 | -0.17556 | 1 |
| SGSM1 | -0.2324227 | 1 | 0.088837 | 1 |
| TAS2R31 | -0.061256 | 1 | -0.08205 | 1 |
| GPR174 | -0.8870696 | 1 | 0.743794 | 1 |
| RAB36 | -0.0557105 | 1 | -0.08751 | 1 |
| TSC22D3 | -0.17177 | 1 | 0.0292 | 1 |
| ARHGEF28 | -0.1655352 | 1 | 0.023922 | 1 |
| ZEB2 | -0.1664119 | 1 | 0.025328 | 1 |
| TNIK | -0.1438315 | 1 | 0.002772 | 1 |
| ANK1 | -0.0840775 | 1 | -0.05626 | 1 |
| ZNF331 | -0.0627549 | 1 | -0.07749 | 1 |
| SOX12 | -0.0259125 | 1 | -0.1136 | 1 |
| ABCC8 | -0.3658897 | 1 | 0.226654 | 1 |
| RERGL | -0.0836983 | 1 | -0.05538 | 1 |
| UBN1 | -0.0893463 | 1 | -0.04973 | 1 |
| MEIS3 | -2.0841108 | 1 | 1.945318 | 1 |
| GRIN2B | -0.0770278 | 1 | -0.06085 | 1 |
| CX3CL1 | -0.2705466 | 1 | 0.133324 | 1 |
| NCOA7 | -0.0340659 | 1 | -0.1031 | 1 |
| EYA2 | 0.03409293 | 1 | -0.17065 | 1 |
| CHST11 | -0.7904304 | 1 | 0.653953 | 1 |
| PSPC1 | -0.064683 | 1 | -0.07119 | 1 |
| ZNF80 | -0.1865735 | 1 | 0.05089 | 1 |
| MED28 | -0.1390775 | 1 | 0.004538 | 1 |
| ABRAXAS1 | 0.02112077 | 1 | -0.15552 | 1 |
| NPY4R | -0.0391459 | 1 | -0.09503 | 1 |
| CANT1 | 0.85887804 | 1 | -0.99251 | 1 |
| ZNF732 | -0.114653 | 1 | -0.01872 | 1 |
| AGPAT2 | 0.00863343 | 1 | -0.14185 | 1 |

| | | | | |
|---|---|---|---|---|
| CASR | -0.0389777 | 1 | -0.09402 | 1 |
| TEX2 | -0.1344874 | 1 | 0.001677 | 1 |
| TRIM5 | -0.1419735 | 1 | 0.009274 | 1 |
| DUXA | -0.0495859 | 1 | -0.08273 | 1 |
| ZNF556 | -0.1018379 | 1 | -0.03026 | 1 |
| OR5D13 | -0.0991915 | 1 | -0.03224 | 1 |
| ZNF705B | -0.028208 | 1 | -0.10254 | 1 |
| OR2T7 | -0.0843233 | 1 | -0.0462 | 1 |
| ARL4A | -0.0580281 | 1 | -0.07236 | 1 |
| SH3BGR | -0.1153683 | 1 | -0.01424 | 1 |
| YBX1 | 0.25988674 | 1 | -0.38871 | 1 |
| MCHR2 | -0.0630171 | 1 | -0.06547 | 1 |
| FGF3 | -0.0537958 | 1 | -0.07403 | 1 |
| ZNF197 | -0.0309258 | 1 | -0.09686 | 1 |
| MLNR | -0.7570715 | 1 | 0.629399 | 1 |
| EPGN | -0.0721838 | 1 | -0.05524 | 1 |
| MLXIP | -0.0528295 | 1 | -0.07401 | 1 |
| ELP1 | 0.12846877 | 1 | -0.25531 | 1 |
| AGRN | -0.1749634 | 1 | 0.048181 | 1 |
| OR1L1 | -0.0647776 | 1 | -0.06192 | 1 |
| AXL | -0.0757838 | 1 | -0.05077 | 1 |
| RAB6C | -0.0569386 | 1 | -0.0693 | 1 |
| ZNF439 | -0.0671547 | 1 | -0.05885 | 1 |
| ZNF23 | -0.0294988 | 1 | -0.0963 | 1 |
| AKNA | -0.1084309 | 1 | -0.01695 | 1 |
| GPR171 | -0.1051688 | 1 | -0.01993 | 1 |
| HBEGF | -0.0930297 | 1 | -0.032 | 1 |
| SUPT20HL1 | -0.0768208 | 1 | -0.04798 | 1 |
| TAS1R1 | -0.0705909 | 1 | -0.05416 | 1 |
| OR1J2 | -0.0072658 | 1 | -0.11737 | 1 |
| IGHMBP2 | -0.2994077 | 1 | 0.175092 | 1 |
| SALL1 | -0.1486961 | 1 | 0.024536 | 1 |
| AGO1 | -0.1017645 | 1 | -0.02219 | 1 |
| HIF1A | 0.14282369 | 1 | -0.26677 | 1 |
| BATF | -0.027721 | 1 | -0.09567 | 1 |
| BACH2 | -0.1690879 | 1 | 0.045864 | 1 |
| EPHA6 | -0.1103508 | 1 | -0.01269 | 1 |
| ZNF45 | -0.0685657 | 1 | -0.05439 | 1 |
| GNB5 | -0.0707968 | 1 | -0.0518 | 1 |
| BLM | -0.0429189 | 1 | -0.07952 | 1 |
| NKX2.1 | -0.1101903 | 1 | -0.01215 | 1 |
| WFS1 | -0.2956816 | 1 | 0.173453 | 1 |
| UTF1 | -0.1253505 | 1 | 0.003395 | 1 |
| ATP2A1 | 0.01883866 | 1 | -0.1404 | 1 |
| SORCS3 | -0.1846793 | 1 | 0.063179 | 1 |
| ASIC1 | -1.6977621 | 1 | 1.576438 | 1 |

| | | | | |
|---|---|---|---|---|
| EDNRA | -0.0458212 | 1 | -0.07506 | 1 |
| USP21 | 0.01255272 | 1 | -0.13292 | 1 |
| SUDS3 | 0.34196174 | 1 | -0.46167 | 1 |
| SCARF1 | 0.01812992 | 1 | -0.13744 | 1 |
| RAC1 | 0.10867308 | 1 | -0.22792 | 1 |
| NFE2 | -0.0773443 | 1 | -0.041 | 1 |
| ZNF69 | -0.1331223 | 1 | 0.015398 | 1 |
| PRRX2 | -0.0570009 | 1 | -0.0604 | 1 |
| CHMP5 | 0.15373735 | 1 | -0.27112 | 1 |
| MED26 | 0.07628854 | 1 | -0.19253 | 1 |
| MICB | -0.1072036 | 1 | -0.00897 | 1 |
| PMAIP1 | -1.158039 | 1 | 1.041869 | 1 |
| ZNF418 | -0.0835651 | 1 | -0.03256 | 1 |
| RHEBL1 | -0.0683847 | 1 | -0.04727 | 1 |
| DLX4 | -0.0655737 | 1 | -0.04964 | 1 |
| CMKLR2 | -0.1389594 | 1 | 0.023799 | 1 |
| DRD4 | -0.1068883 | 1 | -0.0076 | 1 |
| TFAP2D | -0.0874471 | 1 | -0.02666 | 1 |
| OR1A1 | 0.01153095 | 1 | -0.12459 | 1 |
| ZNF454 | -0.1024824 | 1 | -0.01055 | 1 |
| ZNF212 | 0.02199719 | 1 | -0.13497 | 1 |
| CIDEA | -0.0668418 | 1 | -0.04583 | 1 |
| LIMK1 | 0.62582791 | 1 | -0.73821 | 1 |
| ZRSR2 | 0.02491687 | 1 | -0.13717 | 1 |
| OR8U1 | 0.01778942 | 1 | -0.12953 | 1 |
| CREBBP | -0.1371313 | 1 | 0.025632 | 1 |
| TRIM38 | -0.0496798 | 1 | -0.06122 | 1 |
| MAPK6 | 0.83060404 | 1 | -0.94094 | 1 |
| KHDRBS2 | -0.2921268 | 1 | 0.182109 | 1 |
| OR10G3 | -0.1059017 | 1 | -0.00372 | 1 |
| RGL1 | -1.2749233 | 1 | 1.165903 | 1 |
| UACA | -0.1003507 | 1 | -0.00854 | 1 |
| OR5W2 | -0.0648648 | 1 | -0.04349 | 1 |
| PHF5A | 0.23935164 | 1 | -0.34768 | 1 |
| OR2Y1 | -0.038595 | 1 | -0.06874 | 1 |
| OR4X2 | -0.0019972 | 1 | -0.10459 | 1 |
| OR14A16 | -0.1650129 | 1 | 0.058441 | 1 |
| S100A10 | 0.08979153 | 1 | -0.19603 | 1 |
| RIT2 | -0.0797953 | 1 | -0.02607 | 1 |
| ZNF766 | 0.05717197 | 1 | -0.16159 | 1 |
| OR4K5 | -0.0305294 | 1 | -0.07333 | 1 |
| ITPKA | -0.0068695 | 1 | -0.09697 | 1 |
| IMPA2 | -0.0407981 | 1 | -0.06253 | 1 |
| IKZF4 | -0.1330862 | 1 | 0.02988 | 1 |
| GTF2IRD1 | -0.0425508 | 1 | -0.06059 | 1 |
| ANKRD30A | -0.0463654 | 1 | -0.0567 | 1 |

| | | | | |
|---|---|---|---|---|
| ZNF684 | 0.17185634 | 1 | -0.27454 | 1 |
| AIRE | -0.1486635 | 1 | 0.046136 | 1 |
| CXCR4 | -0.1873883 | 1 | 0.085028 | 1 |
| TFDP1 | 0.06455453 | 1 | -0.16621 | 1 |
| PAX2 | -0.074259 | 1 | -0.02738 | 1 |
| EBP | -0.0292997 | 1 | -0.07229 | 1 |
| PPARG | 0.04516848 | 1 | -0.14618 | 1 |
| ARL5C | -0.0865707 | 1 | -0.01427 | 1 |
| SERTAD3 | 0.97576141 | 1 | -1.07658 | 1 |
| MAK | -0.0111426 | 1 | -0.08944 | 1 |
| DUSP22 | -0.1370736 | 1 | 0.036811 | 1 |
| FOXP2 | -0.0481598 | 1 | -0.05195 | 1 |
| GABRE | -0.073247 | 1 | -0.02646 | 1 |
| SORL1 | -0.2593868 | 1 | 0.161087 | 1 |
| ZFYVE16 | -0.0957473 | 1 | -0.00243 | 1 |
| ZMYND11 | -0.0307806 | 1 | -0.06733 | 1 |
| GRHL2 | -0.0628808 | 1 | -0.03473 | 1 |
| ERCC3 | -0.0650253 | 1 | -0.03257 | 1 |
| HMGA2 | -0.0244044 | 1 | -0.07318 | 1 |
| OR56B4 | 0.00513331 | 1 | -0.1027 | 1 |
| FOSL1 | 0.13017972 | 1 | -0.22667 | 1 |
| GMEB2 | -0.0739605 | 1 | -0.02248 | 1 |
| DMRTB1 | -0.0673408 | 1 | -0.02846 | 1 |
| HTR3A | -0.0625644 | 1 | -0.0324 | 1 |
| YPEL5 | 0.10762172 | 1 | -0.20243 | 1 |
| HOXC12 | -0.0325111 | 1 | -0.06187 | 1 |
| NCOR1 | -0.1051046 | 1 | 0.010722 | 1 |
| JDP2 | -0.1145128 | 1 | 0.02021 | 1 |
| ZNF705D | -0.0715482 | 1 | -0.02268 | 1 |
| ACAP1 | -0.1932614 | 1 | 0.100178 | 1 |
| IL15 | 0.00156076 | 1 | -0.09463 | 1 |
| ZNF280A | 0.01991385 | 1 | -0.11188 | 1 |
| SSTR4 | -0.1069446 | 1 | 0.015139 | 1 |
| CXXC1 | -0.3852968 | 1 | 0.294096 | 1 |
| TSSK4 | -0.1149164 | 1 | 0.024086 | 1 |
| TBX21 | -0.1000587 | 1 | 0.009577 | 1 |
| MESP2 | -0.0596346 | 1 | -0.03057 | 1 |
| RERG | -0.0460622 | 1 | -0.04345 | 1 |
| OR4D1 | 0.00981781 | 1 | -0.0991 | 1 |
| ZNF676 | -0.1730221 | 1 | 0.084463 | 1 |
| DNAJA3 | -0.0161165 | 1 | -0.07188 | 1 |
| CDK6 | -0.0832591 | 1 | -0.00452 | 1 |
| OR2L5 | -0.0326007 | 1 | -0.05501 | 1 |
| ZNF99 | -0.0990306 | 1 | 0.011583 | 1 |
| H4C9 | -0.0919102 | 1 | 0.004485 | 1 |
| ERRFI1 | 0.04444348 | 1 | -0.13173 | 1 |

| | | | | |
|---|---|---|---|---|
| CD38 | -0.2567376 | 1 | 0.169765 | 1 |
| ZNF48 | -0.0635867 | 1 | -0.02268 | 1 |
| PAFAH1B1 | 0.09274569 | 1 | -0.17877 | 1 |
| CD40 | -0.0441208 | 1 | -0.04184 | 1 |
| CHRND | -0.0280513 | 1 | -0.05789 | 1 |
| VGLL3 | -0.1608695 | 1 | 0.075832 | 1 |
| SKAP2 | -0.105608 | 1 | 0.020852 | 1 |
| ZNF587 | -0.0109422 | 1 | -0.07262 | 1 |
| ROPN1 | -0.1266171 | 1 | 0.0433 | 1 |
| ZC3H8 | -0.0264155 | 1 | -0.05675 | 1 |
| CNR2 | -0.0756326 | 1 | -0.00742 | 1 |
| NOTCH3 | -0.0611075 | 1 | -0.02183 | 1 |
| CCR1 | -0.0873448 | 1 | 0.00489 | 1 |
| ASZ1 | -0.0158895 | 1 | -0.06602 | 1 |
| SIGLEC9 | -0.046492 | 1 | -0.03512 | 1 |
| IL31RA | -0.0782845 | 1 | -0.0033 | 1 |
| ZNF584 | -0.0292629 | 1 | -0.05211 | 1 |
| CCR4 | -0.0847564 | 1 | 0.003931 | 1 |
| OR5F1 | 0.03491362 | 1 | -0.11559 | 1 |
| PLEKHG4 | -0.0585557 | 1 | -0.02173 | 1 |
| ZNF554 | 0.11046051 | 1 | -0.19009 | 1 |
| ADGRV1 | -0.0613879 | 1 | -0.01811 | 1 |
| ZNF343 | 0.10290829 | 1 | -0.1824 | 1 |
| CNKSR1 | -0.1299747 | 1 | 0.050657 | 1 |
| RAB19 | -0.1112168 | 1 | 0.032446 | 1 |
| MC3R | -0.0137608 | 1 | -0.06459 | 1 |
| PKN3 | -0.1150165 | 1 | 0.036855 | 1 |
| H4C16 | -0.1404333 | 1 | 0.06237 | 1 |
| OR10Z1 | -0.0873913 | 1 | 0.009619 | 1 |
| OR51B6 | -0.0198364 | 1 | -0.05722 | 1 |
| STK3 | -0.0143931 | 1 | -0.0626 | 1 |
| HIVEP3 | 1.71425284 | 1 | -1.79102 | 1 |
| ZC3H14 | -0.0700188 | 1 | -0.00661 | 1 |
| GRIK2 | -0.226679 | 1 | 0.150206 | 1 |
| DLL4 | -0.0240588 | 1 | -0.05225 | 1 |
| PLEK | 0.03170284 | 1 | -0.10763 | 1 |
| LASP1 | -0.0724918 | 1 | -0.00336 | 1 |
| SCRT2 | -0.1606667 | 1 | 0.085085 | 1 |
| FPR3 | -0.0007713 | 1 | -0.0746 | 1 |
| CDC42BPG | -0.1470183 | 1 | 0.071651 | 1 |
| LAX1 | -0.0681214 | 1 | -0.00707 | 1 |
| HTR1A | -0.031932 | 1 | -0.04271 | 1 |
| RAPGEF6 | 0.16362811 | 1 | -0.23762 | 1 |
| OR56A1 | -0.0628339 | 1 | -0.01 | 1 |
| MYLK2 | -0.0691578 | 1 | -0.00333 | 1 |
| IL3 | -0.0301069 | 1 | -0.04193 | 1 |

| | | | | |
|---|---|---|---|---|
| SORBS1 | -0.9784233 | 1 | 0.906405 | 1 |
| ZBTB40 | 0.02281735 | 1 | -0.09469 | 1 |
| OR51E1 | -0.0367362 | 1 | -0.03512 | 1 |
| TAS2R7 | -0.0103334 | 1 | -0.06115 | 1 |
| KIR3DL1 | -0.0148189 | 1 | -0.05596 | 1 |
| PTGES | -0.0762533 | 1 | 0.005777 | 1 |
| RAX2 | 0.67859315 | 1 | -0.74891 | 1 |
| ARAP3 | 0.00244359 | 1 | -0.07262 | 1 |
| NDN | -0.0406103 | 1 | -0.02869 | 1 |
| GRK6 | -0.1147685 | 1 | 0.045629 | 1 |
| FH | 0.09972022 | 1 | -0.16878 | 1 |
| ZNF586 | 0.02998819 | 1 | -0.09897 | 1 |
| ZNF483 | -0.0407924 | 1 | -0.02811 | 1 |
| HES3 | -0.0264135 | 1 | -0.0424 | 1 |
| ZNF836 | -0.0566687 | 1 | -0.01212 | 1 |
| RASGRP2 | -0.0860542 | 1 | 0.01748 | 1 |
| ZNF773 | 0.05916134 | 1 | -0.12724 | 1 |
| KDM2A | 0.04386662 | 1 | -0.11134 | 1 |
| IL22RA2 | -0.1767932 | 1 | 0.109375 | 1 |
| HLA.DRB4 | -0.1501438 | 1 | 0.082934 | 1 |
| UIMC1 | 0.1415764 | 1 | -0.20751 | 1 |
| PER2 | -0.1033264 | 1 | 0.037996 | 1 |
| CD34 | 0.08390119 | 1 | -0.14895 | 1 |
| AMBP | -0.0416581 | 1 | -0.02329 | 1 |
| TAS2R9 | 0.54884358 | 1 | -0.61298 | 1 |
| ZNF221 | -0.2763202 | 1 | 0.212243 | 1 |
| TDRD3 | 1.24458478 | 1 | -1.30835 | 1 |
| RASGEF1A | -0.119448 | 1 | 0.055742 | 1 |
| OR9G4 | 0.02585043 | 1 | -0.08898 | 1 |
| GUCA1A | 0.00792913 | 1 | -0.07056 | 1 |
| CNOT8 | 0.60010159 | 1 | -0.66272 | 1 |
| KLF14 | 0.13077685 | 1 | -0.1929 | 1 |
| TIFA | -0.0058073 | 1 | -0.05628 | 1 |
| SMAD3 | -0.0905372 | 1 | 0.028456 | 1 |
| ZNF469 | -1.4473997 | 1 | 1.385445 | 1 |
| HOXC11 | -0.0180582 | 1 | -0.04383 | 1 |
| ZNF704 | -0.0878335 | 1 | 0.02606 | 1 |
| OR2B3 | -0.0097998 | 1 | -0.05059 | 1 |
| POU3F3 | -0.054281 | 1 | -0.00611 | 1 |
| TAS2R60 | -0.0227848 | 1 | -0.03752 | 1 |
| PTGIS | -0.0714219 | 1 | 0.011486 | 1 |
| SMOC2 | -0.091422 | 1 | 0.032312 | 1 |
| PMS2 | -0.0004368 | 1 | -0.05855 | 1 |
| ZSCAN5A | 0.04835704 | 1 | -0.10716 | 1 |
| ARID1B | -0.7386538 | 1 | 0.679931 | 1 |
| PITPNC1 | -0.0531697 | 1 | -0.0055 | 1 |

| | | | | |
|---|---|---|---|---|
| ID4 | -0.0161881 | 1 | -0.04248 | 1 |
| WNT3A | -0.0517216 | 1 | -0.00604 | 1 |
| OR5K4 | -0.0875129 | 1 | 0.030199 | 1 |
| TFAP2A | -0.038517 | 1 | -0.01833 | 1 |
| POU4F2 | -0.1752859 | 1 | 0.118484 | 1 |
| IRX2 | -0.1752776 | 1 | 0.118563 | 1 |
| FOXD3 | 0.01550322 | 1 | -0.07168 | 1 |
| TOX2 | -0.0543108 | 1 | -0.00179 | 1 |
| ZNF470 | -0.0275683 | 1 | -0.02853 | 1 |
| ASB17 | 0.00309733 | 1 | -0.05916 | 1 |
| ZSCAN16 | -0.1542812 | 1 | 0.098519 | 1 |
| OR8D4 | -0.0802043 | 1 | 0.024559 | 1 |
| GTF2H4 | -0.0078265 | 1 | -0.04754 | 1 |
| P2RY8 | -0.069931 | 1 | 0.014797 | 1 |
| MYZAP | -0.1148242 | 1 | 0.059728 | 1 |
| TAL2 | -0.7513744 | 1 | 0.696397 | 1 |
| RHOXF2 | -0.0242015 | 1 | -0.03037 | 1 |
| ZNF506 | -0.0859826 | 1 | 0.031573 | 1 |
| MEIS2 | -0.0391346 | 1 | -0.01516 | 1 |
| CRK | 0.23093487 | 1 | -0.28468 | 1 |
| KLRD1 | -0.0501096 | 1 | -0.00338 | 1 |
| PFKFB1 | -0.8356058 | 1 | 0.78218 | 1 |
| FOXQ1 | -0.0700031 | 1 | 0.016624 | 1 |
| CCL28 | -0.0190985 | 1 | -0.03415 | 1 |
| CREBL2 | -0.0063359 | 1 | -0.04687 | 1 |
| LYL1 | -0.1087626 | 1 | 0.055999 | 1 |
| NLRP3 | 0.03543062 | 1 | -0.08779 | 1 |
| NSG2 | -0.1047881 | 1 | 0.05282 | 1 |
| CDC42 | 0.97760882 | 1 | -1.02742 | 1 |
| RAB27A | 0.10513784 | 1 | -0.15487 | 1 |
| ACTB | 0.15644094 | 1 | -0.20582 | 1 |
| FOXC2 | -0.0124893 | 1 | -0.0366 | 1 |
| SCML4 | -0.0551001 | 1 | 0.006638 | 1 |
| ARHGAP6 | -0.0626598 | 1 | 0.014213 | 1 |
| ADCY4 | -0.1680005 | 1 | 0.119624 | 1 |
| RAB29 | 0.03916815 | 1 | -0.08738 | 1 |
| MACF1 | -0.1164165 | 1 | 0.068218 | 1 |
| OR2T27 | -0.0038477 | 1 | -0.04413 | 1 |
| OR13C4 | -0.0449176 | 1 | -0.00284 | 1 |
| RHBDL3 | -0.2661226 | 1 | 0.218388 | 1 |
| ZNF750 | -0.0900693 | 1 | 0.042358 | 1 |
| BTG2 | -0.0503285 | 1 | 0.003103 | 1 |
| DUSP21 | -0.0535622 | 1 | 0.007266 | 1 |
| SH2D1A | 0.03013963 | 1 | -0.07628 | 1 |
| ANKRD1 | -0.0399946 | 1 | -0.00604 | 1 |
| ZCCHC13 | -0.0606161 | 1 | 0.015323 | 1 |

| | | | | |
|---|---|---|---|---|
| SH2D3A | 0.04628024 | 1 | -0.09049 | 1 |
| CD8B | -0.1316141 | 1 | 0.087673 | 1 |
| OR4X1 | -0.0648237 | 1 | 0.020918 | 1 |
| VAX1 | 0.04927956 | 1 | -0.09291 | 1 |
| NODAL | 0.04288688 | 1 | -0.08606 | 1 |
| KAT6B | -0.0661807 | 1 | 0.023302 | 1 |
| SSX3 | -0.1259622 | 1 | 0.083482 | 1 |
| OR2A14 | 0.00215605 | 1 | -0.0446 | 1 |
| RAN | 1.2728022 | 1 | -1.31494 | 1 |
| ZNF705G | -0.0081249 | 1 | -0.03393 | 1 |
| CSNK1A1L | -0.0428163 | 1 | 0.000805 | 1 |
| COL1A1 | -0.045595 | 1 | 0.003727 | 1 |
| SSX2B | 0.01730179 | 1 | -0.05887 | 1 |
| ZSCAN2 | -1.542155 | 1 | 1.500655 | 1 |
| FZD7 | 0.00563498 | 1 | -0.04602 | 1 |
| TSHZ3 | -0.1097058 | 1 | 0.069374 | 1 |
| SDHD | 0.14036383 | 1 | -0.18042 | 1 |
| SOHLH2 | 0.0459598 | 1 | -0.08602 | 1 |
| DUSP4 | 0.30897266 | 1 | -0.34901 | 1 |
| MC4R | -0.0601504 | 1 | 0.020262 | 1 |
| MYF5 | 0.00430722 | 1 | -0.04408 | 1 |
| IL4R | -0.0459352 | 1 | 0.006696 | 1 |
| WNT1 | -0.086728 | 1 | 0.047663 | 1 |
| TBC1D2 | -0.1034959 | 1 | 0.064873 | 1 |
| OR1J1 | -0.0453869 | 1 | 0.007092 | 1 |
| CEACAM6 | -0.0258034 | 1 | -0.0122 | 1 |
| ZNF132 | 0.02978722 | 1 | -0.06733 | 1 |
| CACNA1I | -0.1500231 | 1 | 0.112518 | 1 |
| OR4C11 | 0.0433124 | 1 | -0.08068 | 1 |
| OR8B4 | 0.0470044 | 1 | -0.08425 | 1 |
| GPR88 | 0.03692756 | 1 | -0.07414 | 1 |
| CASP10 | 0.00016806 | 1 | -0.03714 | 1 |
| RGS14 | 0.0346788 | 1 | -0.07146 | 1 |
| HTATIP2 | -0.4174332 | 1 | 0.381003 | 1 |
| OR5AK2 | -0.0459893 | 1 | 0.009635 | 1 |
| BCL9L | -0.1735554 | 1 | 0.137204 | 1 |
| PRDM12 | -0.0627974 | 1 | 0.027159 | 1 |
| DIDO1 | -0.0191838 | 1 | -0.0164 | 1 |
| ARL1 | 1.02701044 | 1 | -1.06257 | 1 |
| SIX2 | -0.0184912 | 1 | -0.01681 | 1 |
| CD4 | -0.0339142 | 1 | -0.00117 | 1 |
| EPHA3 | -0.0206857 | 1 | -0.01399 | 1 |
| IL2RG | -0.0692764 | 1 | 0.034806 | 1 |
| OR10P1 | -0.0515632 | 1 | 0.017233 | 1 |
| NDP | -0.4971807 | 1 | 0.462985 | 1 |
| RFX6 | -0.0031871 | 1 | -0.03097 | 1 |

| | | | | |
|---|---|---|---|---|
| SPSB2 | 0.0393382 | 1 | -0.07324 | 1 |
| ANKDD1B | -0.06083 | 1 | 0.027029 | 1 |
| OR10S1 | -0.0705084 | 1 | 0.037377 | 1 |
| MCF2L | -0.0961334 | 1 | 0.063313 | 1 |
| CBLL2 | 0.02638128 | 1 | -0.05813 | 1 |
| EID2 | 0.18118807 | 1 | -0.21294 | 1 |
| INTS6 | 0.16491854 | 1 | -0.19652 | 1 |
| OR2L8 | -0.0683747 | 1 | 0.03695 | 1 |
| OR1S1 | 1.31136279 | 1 | -1.34266 | 1 |
| CABP5 | 0.04500362 | 1 | -0.07581 | 1 |
| NKX3.2 | -0.0504024 | 1 | 0.019774 | 1 |
| ZNF224 | 0.01840239 | 1 | -0.04888 | 1 |
| TAAR8 | -0.0479605 | 1 | 0.017718 | 1 |
| ZNF365 | -0.0588405 | 1 | 0.028912 | 1 |
| EPC2 | 0.23808491 | 1 | -0.26761 | 1 |
| OR2M2 | -0.0066622 | 1 | -0.02207 | 1 |
| TSPAN6 | 0.12647444 | 1 | -0.15448 | 1 |
| ZNF225 | -0.053415 | 1 | 0.026566 | 1 |
| PI4KA | -0.1049599 | 1 | 0.078871 | 1 |
| RRAGC | 0.2268139 | 1 | -0.25284 | 1 |
| LAG3 | -0.0729731 | 1 | 0.047258 | 1 |
| GFRA1 | -0.0645963 | 1 | 0.038931 | 1 |
| HDAC1 | 0.92505106 | 1 | -0.95053 | 1 |
| LRP1 | -0.2279115 | 1 | 0.203236 | 1 |
| B2M | 0.06828673 | 1 | -0.0927 | 1 |
| ZNF334 | -0.0124955 | 1 | -0.01108 | 1 |
| OR13A1 | -0.0159881 | 1 | -0.00758 | 1 |
| GBX1 | 0.00169246 | 1 | -0.0248 | 1 |
| SLA2 | -0.0189637 | 1 | -0.00393 | 1 |
| PAK1IP1 | 0.50571158 | 1 | -0.52846 | 1 |
| ZNF502 | 0.18188048 | 1 | -0.20413 | 1 |
| PAK1 | -0.0213991 | 1 | -0.00012 | 1 |
| ARL14 | 0.03547068 | 1 | -0.05671 | 1 |
| KLF1 | -0.036554 | 1 | 0.015541 | 1 |
| CCKBR | -0.2125592 | 1 | 0.192058 | 1 |
| PRKAG3 | 0.00699668 | 1 | -0.02739 | 1 |
| TBC1D8 | 0.00126746 | 1 | -0.02131 | 1 |
| NKX2.6 | -0.0249622 | 1 | 0.006116 | 1 |
| CTNNBIP1 | -0.080551 | 1 | 0.062509 | 1 |
| ADGRG4 | 0.00216215 | 1 | -0.02003 | 1 |
| CNTNAP2 | -0.1306048 | 1 | 0.11333 | 1 |
| KLHL31 | -0.2609653 | 1 | 0.244225 | 1 |
| HMX3 | 0.01452289 | 1 | -0.0312 | 1 |
| LALBA | 0.01705912 | 1 | -0.0337 | 1 |
| LMX1B | 0.0191446 | 1 | -0.03574 | 1 |
| CALCR | 0.01924598 | 1 | -0.03537 | 1 |

| | | | | |
|---|---|---|---|---|
| GTF2IRD2 | 0.00591317 | 1 | -0.02198 | 1 |
| MS4A13 | 0.03888831 | 1 | -0.05452 | 1 |
| SOX17 | -0.0128541 | 1 | -0.00226 | 1 |
| PDE6G | -0.0661007 | 1 | 0.051615 | 1 |
| OR51F2 | -0.0352543 | 1 | 0.021516 | 1 |
| PRLH | -0.011451 | 1 | -0.00213 | 1 |
| OR2L3 | 0.05503593 | 1 | -0.06827 | 1 |
| LHX4 | -0.0830939 | 1 | 0.069955 | 1 |
| AZU1 | 0.01829524 | 1 | -0.03119 | 1 |
| MBD4 | 0.13609005 | 1 | -0.14866 | 1 |
| GBF1 | -0.0679575 | 1 | 0.055433 | 1 |
| IRS4 | 0.02150665 | 1 | -0.03376 | 1 |
| PHB2 | 0.86229067 | 1 | -0.87405 | 1 |
| IL5RA | -0.0868819 | 1 | 0.075282 | 1 |
| RASL11A | -0.0585902 | 1 | 0.047044 | 1 |
| ZNF503 | -0.0028152 | 1 | -0.0083 | 1 |
| ARHGEF17 | 0.01520719 | 1 | -0.02622 | 1 |
| OR3A3 | -0.0708803 | 1 | 0.060131 | 1 |
| ARX | -0.0305128 | 1 | 0.019794 | 1 |
| TIGD1 | -0.0317777 | 1 | 0.021066 | 1 |
| ZNF274 | -0.0399326 | 1 | 0.029379 | 1 |
| MS4A1 | 0.03683958 | 1 | -0.04729 | 1 |
| VN1R4 | 0.04628868 | 1 | -0.05621 | 1 |
| IL23R | -0.0204943 | 1 | 0.010789 | 1 |
| HOMER2 | -0.0318924 | 1 | 0.022612 | 1 |
| CCKAR | 0.03652606 | 1 | -0.04578 | 1 |
| RCAN3 | -0.0030055 | 1 | -0.00606 | 1 |
| ZFAND3 | 0.30599821 | 1 | -0.31427 | 1 |
| SPHK1 | 0.07602484 | 1 | -0.08393 | 1 |
| CCR6 | 0.03252623 | 1 | -0.04038 | 1 |
| RBM39 | 0.05127593 | 1 | -0.05892 | 1 |
| PROKR2 | -0.0613844 | 1 | 0.055074 | 1 |
| CRHR1 | -0.375614 | 1 | 0.369419 | 1 |
| ITGB1BP2 | -0.0407649 | 1 | 0.034795 | 1 |
| FGF19 | -0.0472313 | 1 | 0.041455 | 1 |
| ANKZF1 | -0.1976545 | 1 | 0.191913 | 1 |
| POU3F1 | 0.00012618 | 1 | -0.00515 | 1 |
| CYLD | -0.1103502 | 1 | 0.105402 | 1 |
| ADGRG2 | -0.0504009 | 1 | 0.045466 | 1 |
| TAS2R3 | -0.0469586 | 1 | 0.042057 | 1 |
| ETS2 | -0.0702987 | 1 | 0.065863 | 1 |
| FOXF2 | 0.03394395 | 1 | -0.03668 | 1 |
| PTGER3 | -0.0901023 | 1 | 0.087372 | 1 |
| FOXR1 | -0.0496909 | 1 | 0.047078 | 1 |
| ARAF | 0.14893391 | 1 | -0.15127 | 1 |
| AGAP1 | -0.0519862 | 1 | 0.04991 | 1 |

| | | | | |
|---|---|---|---|---|
| OR1A2 | -0.0329339 | 1 | 0.030933 | 1 |
| IFT88 | 0.14571194 | 1 | -0.14747 | 1 |
| FGD4 | -0.1798172 | 1 | 0.178114 | 1 |
| FAM13B | 0.06420134 | 1 | -0.06577 | 1 |
| RPS27L | 0.1400184 | 1 | -0.14113 | 1 |
| NKX6.3 | -0.1531463 | 1 | 0.152077 | 1 |
| HES4 | -0.002495 | 1 | 0.001722 | 1 |
| MAP3K4 | 0.04775878 | 1 | -0.04797 | 1 |
| SH2B3 | -0.0933851 | 1 | 0.093437 | 1 |
| WNK2 | -0.4525642 | 1 | 0.452666 | 1 |
| PYDC1 | -0.0297045 | 1 | 0.030854 | 1 |
| TAS2R38 | -0.0263161 | 1 | 0.027466 | 1 |
| EFNB3 | -0.1262667 | 1 | 0.127473 | 1 |
| TNFSF9 | -0.0309486 | 1 | 0.032473 | 1 |
| ZNF438 | 0.03595919 | 1 | -0.03359 | 1 |
| KIDINS220 | 0.14212435 | 1 | -0.13962 | 1 |
| NHLH1 | -0.0124512 | 1 | 0.015035 | 1 |
| BHLHE22 | -0.0325507 | 1 | 0.035536 | 1 |
| MBD3L1 | 0.01915723 | 1 | -0.01606 | 1 |
| OR2T11 | -0.0212817 | 1 | 0.024401 | 1 |
| PCSK6 | -0.2685607 | 1 | 0.27198 | 1 |
| ACOT11 | -0.0802622 | 1 | 0.08401 | 1 |
| ASCL4 | 0.00169533 | 1 | 0.002605 | 1 |
| PAXBP1 | 0.00036118 | 1 | 0.00403 | 1 |
| OR9Q1 | -0.0015308 | 1 | 0.005923 | 1 |
| OPN1MW | 0.13609306 | 1 | -0.13124 | 1 |
| AVP | -0.0350497 | 1 | 0.039961 | 1 |
| CDX2 | 0.02703711 | 1 | -0.02211 | 1 |
| AMHR2 | 0.01411609 | 1 | -0.00904 | 1 |
| GNA15 | -0.0078367 | 1 | 0.012952 | 1 |
| ECT2L | -0.1072422 | 1 | 0.112872 | 1 |
| MSGN1 | 0.01115106 | 1 | -0.00522 | 1 |
| NLRP12 | -0.0957004 | 1 | 0.101786 | 1 |
| FZD9 | -0.0246671 | 1 | 0.032328 | 1 |
| OR5I1 | -0.0165246 | 1 | 0.024283 | 1 |
| ZC3H12D | 0.0433676 | 1 | -0.03502 | 1 |
| PITX3 | 0.0169954 | 1 | -0.00855 | 1 |
| SOCS2 | 0.04539592 | 1 | -0.03685 | 1 |
| OR5AU1 | 0.04402948 | 1 | -0.03547 | 1 |
| RND2 | -0.0845453 | 1 | 0.093127 | 1 |
| NKX1.2 | 0.01864873 | 1 | -0.00976 | 1 |
| RAB24 | -0.0250318 | 1 | 0.033948 | 1 |
| HTR1F | -0.0385921 | 1 | 0.047512 | 1 |
| OR10G7 | 0.01810978 | 1 | -0.00887 | 1 |
| ARID3B | -0.0548087 | 1 | 0.064191 | 1 |
| PIAS3 | 0.00184351 | 1 | 0.007748 | 1 |

| | | | | |
|---|---|---|---|---|
| MAPKAPK2 | 0.10486273 | 1 | -0.09474 | 1 |
| SPIB | -0.1275911 | 1 | 0.137729 | 1 |
| MKNK1 | -0.0428384 | 1 | 0.053814 | 1 |
| TCEA2 | -0.0784469 | 1 | 0.089714 | 1 |
| IFNLR1 | -0.0394282 | 1 | 0.051356 | 1 |
| GPR26 | -0.0997709 | 1 | 0.112039 | 1 |
| CHRM2 | -0.6638731 | 1 | 0.676693 | 1 |
| MS4A7 | -0.0405604 | 1 | 0.05375 | 1 |
| SOX30 | 0.06349068 | 1 | -0.04968 | 1 |
| ZNF703 | -0.0193825 | 1 | 0.033364 | 1 |
| GRAP | 0.00313648 | 1 | 0.011839 | 1 |
| SPARCL1 | -0.000108 | 1 | 0.015147 | 1 |
| ZNF248 | 0.21680332 | 1 | -0.20162 | 1 |
| CHRNA4 | -0.4520542 | 1 | 0.467311 | 1 |
| MAP3K8 | -0.0191413 | 1 | 0.034545 | 1 |
| OR8U9 | -0.0101038 | 1 | 0.025673 | 1 |
| HOXC10 | 0.02350852 | 1 | -0.00769 | 1 |
| PRKCE | 0.22476271 | 1 | -0.20876 | 1 |
| HCRT | -0.005628 | 1 | 0.021919 | 1 |
| ZMAT2 | 0.77541826 | 1 | -0.75895 | 1 |
| HLF | -0.1506532 | 1 | 0.167537 | 1 |
| OR2T3 | 0.00871207 | 1 | 0.008561 | 1 |
| ZMYND8 | 0.04871021 | 1 | -0.03141 | 1 |
| TAF1D | 0.79787957 | 1 | -0.78027 | 1 |
| OR4A16 | -0.022152 | 1 | 0.039813 | 1 |
| NMBR | -0.4458051 | 1 | 0.463527 | 1 |
| RIN2 | -0.0325318 | 1 | 0.050293 | 1 |
| TBC1D7 | 0.73867719 | 1 | -0.72087 | 1 |
| STMN4 | -0.5932147 | 1 | 0.611075 | 1 |
| ZSCAN21 | 0.09396782 | 1 | -0.07554 | 1 |
| RP1 | 0.02076232 | 1 | -0.00218 | 1 |
| ZKSCAN3 | -0.0176126 | 1 | 0.036348 | 1 |
| ZNF662 | -0.0496833 | 1 | 0.068672 | 1 |
| ZNF583 | -0.1224999 | 1 | 0.141806 | 1 |
| OR5L2 | 0.02662113 | 1 | -0.00643 | 1 |
| ZSCAN4 | -0.0468945 | 1 | 0.06742 | 1 |
| TRPV4 | -0.0330287 | 1 | 0.053774 | 1 |
| CBFA2T3 | 0.46278012 | 1 | -0.44166 | 1 |
| GPR27 | -0.0841468 | 1 | 0.105339 | 1 |
| USP16 | 0.29792972 | 1 | -0.2765 | 1 |
| ZCCHC9 | 0.06033608 | 1 | -0.03825 | 1 |
| PIK3R1 | -0.0961838 | 1 | 0.118317 | 1 |
| GRIN2D | -0.0357309 | 1 | 0.058444 | 1 |
| PLEKHG7 | 0.00551846 | 1 | 0.017613 | 1 |
| ZMIZ1 | -0.0171663 | 1 | 0.040323 | 1 |
| EMX2 | 0.01442487 | 1 | 0.008741 | 1 |

| | | | | |
|---|---|---|---|---|
| SALL2 | -0.095202 | 1 | 0.118372 | 1 |
| MRGPRX2 | -0.0218547 | 1 | 0.045117 | 1 |
| OPRM1 | -0.0253274 | 1 | 0.049019 | 1 |
| IRAK1BP1 | 0.05668418 | 1 | -0.03292 | 1 |
| EYA3 | 0.10487789 | 1 | -0.08105 | 1 |
| PTPRF | -0.1292296 | 1 | 0.153723 | 1 |
| OR6T1 | 0.00365027 | 1 | 0.02088 | 1 |
| TFIP11 | 0.12841142 | 1 | -0.10385 | 1 |
| SHOX2 | 0.04052573 | 1 | -0.0159 | 1 |
| OR8G5 | -0.0132118 | 1 | 0.038484 | 1 |
| ZNF667 | 0.05606607 | 1 | -0.03052 | 1 |
| OPN1MW2 | 0.02710612 | 1 | -0.00155 | 1 |
| TRRAP | 0.07225843 | 1 | -0.04628 | 1 |
| PYGO2 | 0.05889361 | 1 | -0.0327 | 1 |
| PROP1 | -0.0011451 | 1 | 0.028092 | 1 |
| TEAD2 | -0.1401885 | 1 | 0.16804 | 1 |
| ARHGAP36 | -0.0346915 | 1 | 0.062778 | 1 |
| SIRPB1 | -0.0263018 | 1 | 0.054578 | 1 |
| OR4D2 | 0.04540817 | 1 | -0.01709 | 1 |
| SAP18 | 1.06739642 | 1 | -1.039 | 1 |
| OR13C9 | -0.0247995 | 1 | 0.053205 | 1 |
| MAVS | 0.01967893 | 1 | 0.008889 | 1 |
| TNFRSF10A | -0.0829994 | 1 | 0.111584 | 1 |
| EPHB6 | -1.1225668 | 1 | 1.15139 | 1 |
| C2CD3 | -0.1085778 | 1 | 0.137433 | 1 |
| OR4F16 | -0.0384027 | 1 | 0.067809 | 1 |
| OR4F29 | -0.0010843 | 1 | 0.0309 | 1 |
| EIF4EBP1 | 0.76413099 | 1 | -0.73431 | 1 |
| OR8H3 | 0.01974817 | 1 | 0.010386 | 1 |
| ZNF75A | 0.08836783 | 1 | -0.05763 | 1 |
| COPRS | -0.0432691 | 1 | 0.074034 | 1 |
| PSEN1 | 0.11345095 | 1 | -0.08258 | 1 |
| OR2T33 | 0.0101525 | 1 | 0.02225 | 1 |
| FLT1 | -0.0297827 | 1 | 0.062199 | 1 |
| KLF11 | 0.02265844 | 1 | 0.010074 | 1 |
| TEK | -0.0093594 | 1 | 0.042451 | 1 |
| GPR101 | -0.0049195 | 1 | 0.038613 | 1 |
| MAP3K5 | -0.0843309 | 1 | 0.118133 | 1 |
| KIR3DS1 | 0.05316872 | 1 | -0.01809 | 1 |
| SNCAIP | 0.12822707 | 1 | -0.09307 | 1 |
| TAAR5 | -0.0283544 | 1 | 0.064177 | 1 |
| ZNF200 | 0.1906188 | 1 | -0.15398 | 1 |
| ADGRE1 | -0.0459673 | 1 | 0.082783 | 1 |
| PPP1R1B | -0.1812589 | 1 | 0.218146 | 1 |
| PPP2R2D | 0.01137306 | 1 | 0.025861 | 1 |
| RTL3 | 0.13077546 | 1 | -0.09323 | 1 |

| | | | | |
|---|---|---|---|---|
| HMX2 | -0.0088813 | 1 | 0.047 | 1 |
| ZNF691 | -0.0954326 | 1 | 0.13358 | 1 |
| CD226 | 3.72E-05 | 1 | 0.038793 | 1 |
| TMPRSS6 | 0.03524867 | 1 | 0.00374 | 1 |
| CHRNB4 | 0.02455156 | 1 | 0.014673 | 1 |
| PKD2L1 | 0.03645051 | 1 | 0.00368 | 1 |
| DACH2 | 0.06505145 | 1 | -0.02474 | 1 |
| SCT | 0.0069813 | 1 | 0.03352 | 1 |
| IGSF6 | 0.03738433 | 1 | 0.003798 | 1 |
| SELE | 0.77475471 | 1 | -0.73344 | 1 |
| MRRF | 0.09096728 | 1 | -0.04966 | 1 |
| TAF15 | 0.04196226 | 1 | 0.000469 | 1 |
| OR9A4 | 0.0395698 | 1 | 0.003067 | 1 |
| FOXD1 | 0.01192716 | 1 | 0.030971 | 1 |
| MAP4K4 | -0.8987063 | 1 | 0.94176 | 1 |
| ZNF165 | 0.04561781 | 1 | -0.00247 | 1 |
| ADGRG3 | -0.0569675 | 1 | 0.100206 | 1 |
| MIXL1 | -0.0123582 | 1 | 0.055833 | 1 |
| OR5AR1 | 0.01424478 | 1 | 0.029242 | 1 |
| LMX1A | 0.02139113 | 1 | 0.022223 | 1 |
| BNC2 | 0.08930449 | 1 | -0.04563 | 1 |
| PRDM14 | 0.01394876 | 1 | 0.030067 | 1 |
| PRDM9 | 0.03738134 | 1 | 0.006814 | 1 |
| ZIC5 | -0.0041036 | 1 | 0.048358 | 1 |
| PCGF3 | 0.01305527 | 1 | 0.031255 | 1 |
| EVX1 | 0.04774961 | 1 | -0.0032 | 1 |
| ESR1 | -0.0940695 | 1 | 0.139322 | 1 |
| HMBOX1 | 0.02039898 | 1 | 0.025199 | 1 |
| FCRL2 | 0.04033245 | 1 | 0.005532 | 1 |
| HOXA4 | -0.0134551 | 1 | 0.059319 | 1 |
| OR1I1 | 0.01691508 | 1 | 0.029911 | 1 |
| REXO4 | 0.05045754 | 1 | -0.00333 | 1 |
| DMRT1 | 0.04513506 | 1 | 0.002468 | 1 |
| JARID2 | 0.07517303 | 1 | -0.02743 | 1 |
| FBXO31 | 0.01978454 | 1 | 0.028033 | 1 |
| ZNF606 | 0.01845651 | 1 | 0.029895 | 1 |
| GABRA1 | -0.1653686 | 1 | 0.21405 | 1 |
| NANOGNB | 0.02813343 | 1 | 0.02058 | 1 |
| MS4A15 | 0.03286031 | 1 | 0.017016 | 1 |
| GPR148 | -0.0351301 | 1 | 0.086376 | 1 |
| ZNF576 | 0.0034959 | 1 | 0.047757 | 1 |
| HAND2 | 0.03076899 | 1 | 0.020581 | 1 |
| TIRAP | -0.0462003 | 1 | 0.097734 | 1 |
| ARHGAP1 | 0.02481485 | 1 | 0.027452 | 1 |
| ASMTL | 0.01405663 | 1 | 0.038346 | 1 |
| NKIRAS2 | 0.3790374 | 1 | -0.32638 | 1 |

| | | | | |
|---|---|---|---|---|
| ZNF497 | -0.1093417 | 1 | 0.162333 | 1 |
| GLP2R | -0.000541 | 1 | 0.05386 | 1 |
| CASP9 | 0.0464355 | 1 | 0.007736 | 1 |
| ZFP2 | 0.03227648 | 1 | 0.022434 | 1 |
| TAGAP | -0.106833 | 1 | 0.161572 | 1 |
| GPR19 | 0.09950448 | 1 | -0.04455 | 1 |
| TSHB | 0.02744704 | 1 | 0.027626 | 1 |
| TPTEP2.CSNK1E | -0.0125966 | 1 | 0.06808 | 1 |
| PAX4 | 0.08749471 | 1 | -0.03186 | 1 |
| ARHGEF33 | -0.052044 | 1 | 0.108261 | 1 |
| KIR2DL1 | 0.00925259 | 1 | 0.047061 | 1 |
| OR2A2 | 0.01598429 | 1 | 0.040742 | 1 |
| RAB12 | 0.00998052 | 1 | 0.046773 | 1 |
| KLF15 | -1.8764832 | 1 | 1.933805 | 1 |
| SUPT20HL2 | -0.5671723 | 1 | 0.624579 | 1 |
| DNMT3L | -0.0425684 | 1 | 0.100021 | 1 |
| GPR52 | -0.4594684 | 1 | 0.517014 | 1 |
| CUZD1 | 0.03370557 | 1 | 0.024783 | 1 |
| NFATC4 | 0.88889262 | 1 | -0.8304 | 1 |
| HES1 | 0.0381955 | 1 | 0.020923 | 1 |
| GPR152 | 0.01453582 | 1 | 0.044628 | 1 |
| TBC1D12 | -2.5244829 | 1 | 2.583863 | 1 |
| PRDM15 | -0.0296317 | 1 | 0.089176 | 1 |
| OMP | 0.02363983 | 1 | 0.036102 | 1 |
| RASSF6 | 0.06345644 | 1 | -0.00356 | 1 |
| GNAT3 | 0.05403908 | 1 | 0.006024 | 1 |
| MEOX1 | 0.07027471 | 1 | -0.01014 | 1 |
| DGKD | 0.0374344 | 1 | 0.022793 | 1 |
| SPEN | -0.0309768 | 1 | 0.09134 | 1 |
| GML | 0.63067636 | 1 | -0.56987 | 1 |
| OR5P3 | 0.04791901 | 1 | 0.013018 | 1 |
| ZBED2 | 0.01091915 | 1 | 0.05037 | 1 |
| ZNF423 | -0.0469375 | 1 | 0.108939 | 1 |
| TNK1 | 0.0298954 | 1 | 0.032156 | 1 |
| OR4B1 | 0.01726416 | 1 | 0.04521 | 1 |
| OR7D4 | 0.03526948 | 1 | 0.028141 | 1 |
| EN1 | 0.00576438 | 1 | 0.057857 | 1 |
| AXIN2 | -0.1498207 | 1 | 0.213452 | 1 |
| SNAPC2 | 0.19412133 | 1 | -0.13037 | 1 |
| OR8H1 | -0.057284 | 1 | 0.121061 | 1 |
| ARR3 | 0.04785975 | 1 | 0.015925 | 1 |
| ZNF514 | 0.0343979 | 1 | 0.029673 | 1 |
| PLAGL2 | 0.10144931 | 1 | -0.03734 | 1 |
| ZNF490 | 0.12009851 | 1 | -0.05551 | 1 |
| GIPR | -0.1270831 | 1 | 0.192471 | 1 |
| ZNF568 | 0.13335381 | 1 | -0.0678 | 1 |

| | | | | |
|---|---|---|---|---|
| ZFX | 0.28517497 | 1 | -0.21952 | 1 |
| STK36 | 0.02091629 | 1 | 0.04514 | 1 |
| OR9K2 | 0.00452157 | 1 | 0.061636 | 1 |
| OR1D2 | -0.0017405 | 1 | 0.067963 | 1 |
| SIRT5 | 0.01002672 | 1 | 0.056312 | 1 |
| MSC | 0.02986253 | 1 | 0.036945 | 1 |
| PPP1R2B | 0.05137627 | 1 | 0.015724 | 1 |
| TNNI2 | 0.01549963 | 1 | 0.051649 | 1 |
| TGM2 | -0.0076589 | 1 | 0.074838 | 1 |
| OR51L1 | 0.02784966 | 1 | 0.039507 | 1 |
| SEC31A | 0.2778166 | 1 | -0.21011 | 1 |
| CCN4 | -0.0375876 | 1 | 0.105341 | 1 |
| ZNF398 | 0.08335824 | 1 | -0.01553 | 1 |
| SLTM | 0.05832588 | 1 | 0.010221 | 1 |
| BCL2 | 0.00910838 | 1 | 0.059724 | 1 |
| LHX3 | 0.02902917 | 1 | 0.040084 | 1 |
| GPR142 | -0.1033381 | 1 | 0.172687 | 1 |
| ZNF570 | 0.20887148 | 1 | -0.13929 | 1 |
| NIF3L1 | 0.21178162 | 1 | -0.14209 | 1 |
| CERKL | -0.0026785 | 1 | 0.072642 | 1 |
| ARHGAP9 | 0.03279979 | 1 | 0.037938 | 1 |
| PLEKHG3 | -0.0970985 | 1 | 0.168107 | 1 |
| OR10K2 | -0.0344517 | 1 | 0.107403 | 1 |
| ASB9 | 0.08418872 | 1 | -0.01075 | 1 |
| PRKD1 | -0.0274892 | 1 | 0.101602 | 1 |
| IRF3 | -0.035609 | 1 | 0.109915 | 1 |
| RANBP1 | 0.84701661 | 1 | -0.77252 | 1 |
| ANXA3 | -0.0414023 | 1 | 0.11624 | 1 |
| COPS5 | 0.14772355 | 1 | -0.07218 | 1 |
| ZAP70 | -0.0708243 | 1 | 0.146927 | 1 |
| CCL27 | 0.05154301 | 1 | 0.025444 | 1 |
| S1PR3 | -0.1315694 | 1 | 0.208832 | 1 |
| REN | 0.09659411 | 1 | -0.019 | 1 |
| ZNF142 | -0.0625652 | 1 | 0.140258 | 1 |
| KCNH3 | -0.0957595 | 1 | 0.173551 | 1 |
| GSC2 | 0.02879403 | 1 | 0.049061 | 1 |
| FOXR2 | -0.0222575 | 1 | 0.100132 | 1 |
| DLX3 | 0.06548746 | 1 | 0.012637 | 1 |
| SOCS5 | 0.65723912 | 1 | -0.57876 | 1 |
| TACSTD2 | -0.0502405 | 1 | 0.12957 | 1 |
| RHOXF2B | 0.05449355 | 1 | 0.025276 | 1 |
| ZSCAN26 | 0.10164792 | 1 | -0.02159 | 1 |
| TAS2R42 | 0.01713354 | 1 | 0.063138 | 1 |
| TAF3 | 0.10379803 | 1 | -0.0233 | 1 |
| OR10J1 | 0.05841642 | 1 | 0.022216 | 1 |
| MS4A2 | 0.09785619 | 1 | -0.01701 | 1 |

| | | | | |
|---|---|---|---|---|
| OR52W1 | 0.04955499 | 1 | 0.031682 | 1 |
| DERL3 | 0.00036788 | 1 | 0.080873 | 1 |
| SLC9A3R1 | -0.0711029 | 1 | 0.152612 | 1 |
| CD36 | 0.14144627 | 1 | -0.0599 | 1 |
| RASA4B | -0.0670108 | 1 | 0.148677 | 1 |
| CHKA | 0.06382972 | 1 | 0.018193 | 1 |
| PTPRC | 0.16433695 | 1 | -0.08201 | 1 |
| TRIM16 | 0.08006088 | 1 | 0.002832 | 1 |
| GMEB1 | 0.42613445 | 1 | -0.34291 | 1 |
| ZCWPW2 | -0.0081535 | 1 | 0.091772 | 1 |
| OR2A1 | 0.08594926 | 1 | -0.00214 | 1 |
| RAPGEF1 | -0.0934232 | 1 | 0.177328 | 1 |
| HCFC1 | 0.07635589 | 1 | 0.007796 | 1 |
| ADORA3 | -0.0729422 | 1 | 0.157139 | 1 |
| HTATSF1 | 0.11706252 | 1 | -0.03252 | 1 |
| OR10A3 | -0.0461592 | 1 | 0.130835 | 1 |
| ZMYND19 | 0.05173911 | 1 | 0.033198 | 1 |
| TAS2R1 | 0.12201438 | 1 | -0.03684 | 1 |
| TBPL2 | -0.0789945 | 1 | 0.164367 | 1 |
| FLT4 | 0.0330188 | 1 | 0.052519 | 1 |
| PHF20 | 0.16729935 | 1 | -0.08113 | 1 |
| GPR32 | 0.03175024 | 1 | 0.054534 | 1 |
| FOXF1 | -0.0685717 | 1 | 0.154867 | 1 |
| OR1B1 | 0.00588211 | 1 | 0.080856 | 1 |
| POU5F1B | -0.0077256 | 1 | 0.094748 | 1 |
| MS4A6A | 0.04101495 | 1 | 0.046158 | 1 |
| HOXA1 | -0.0208606 | 1 | 0.108155 | 1 |
| CLEC5A | 0.0689199 | 1 | 0.018543 | 1 |
| GPR21 | 0.05010204 | 1 | 0.037624 | 1 |
| OR51I1 | 0.02841813 | 1 | 0.059348 | 1 |
| OR2AK2 | 0.06107142 | 1 | 0.027525 | 1 |
| ASB4 | -0.0599429 | 1 | 0.148838 | 1 |
| NXN | 0.02529625 | 1 | 0.063782 | 1 |
| SPIC | 0.01921514 | 1 | 0.070123 | 1 |
| ULK2 | 0.03510284 | 1 | 0.054302 | 1 |
| NR2F1 | -0.0841384 | 1 | 0.173558 | 1 |
| TAAR2 | 0.02030241 | 1 | 0.069242 | 1 |
| OR13D1 | 0.02868039 | 1 | 0.061016 | 1 |
| LILRA2 | -0.0045495 | 1 | 0.09435 | 1 |
| PTAFR | -0.0484691 | 1 | 0.138679 | 1 |
| WWOX | 0.00915563 | 1 | 0.081181 | 1 |
| YWHAE | 0.14671535 | 1 | -0.05629 | 1 |
| ZSCAN29 | 0.12561568 | 1 | -0.03514 | 1 |
| TBXA2R | 0.03220046 | 1 | 0.058325 | 1 |
| KIR2DL4 | 0.02353317 | 1 | 0.067034 | 1 |
| OR11H1 | 0.08287976 | 1 | 0.008274 | 1 |

| | | | | |
|---|---|---|---|---|
| CNTNAP1 | -0.0944551 | 1 | 0.185643 | 1 |
| GTF2H2C | 0.0557899 | 1 | 0.035418 | 1 |
| OR4S1 | 0.05804638 | 1 | 0.03385 | 1 |
| ABRA | 0.01396835 | 1 | 0.077954 | 1 |
| NELFA | -0.122653 | 1 | 0.214929 | 1 |
| SNX29 | -0.0143147 | 1 | 0.106703 | 1 |
| RCVRN | 0.02562213 | 1 | 0.066922 | 1 |
| ZDHHC19 | -0.02683 | 1 | 0.119448 | 1 |
| SSX4 | 0.0437323 | 1 | 0.049224 | 1 |
| RASSF2 | -0.1863851 | 1 | 0.280527 | 1 |
| SP5 | 0.01095391 | 1 | 0.083367 | 1 |
| ZNF648 | 0.00797105 | 1 | 0.086623 | 1 |
| PAQR7 | 0.24197077 | 1 | -0.14729 | 1 |
| TCEAL2 | -0.0415745 | 1 | 0.13635 | 1 |
| VGLL1 | 0.0218828 | 1 | 0.073162 | 1 |
| TBL1X | 0.03365657 | 1 | 0.061691 | 1 |
| CNTFR | -0.0544653 | 1 | 0.150296 | 1 |
| APPL2 | -0.0492796 | 1 | 0.145176 | 1 |
| PDE2A | -0.0856329 | 1 | 0.181758 | 1 |
| TBC1D2B | -0.0452337 | 1 | 0.141872 | 1 |
| OR2M5 | 0.05859313 | 1 | 0.038292 | 1 |
| OR56A4 | -0.0955217 | 1 | 0.192835 | 1 |
| UNC119 | -0.0328867 | 1 | 0.130539 | 1 |
| OR6C4 | 0.07704734 | 1 | 0.020721 | 1 |
| DOK5 | 0.02366499 | 1 | 0.074155 | 1 |
| BTAF1 | 0.16544716 | 1 | -0.06762 | 1 |
| BAZ2A | -0.1403795 | 1 | 0.238428 | 1 |
| DPYSL2 | 0.08746537 | 1 | 0.010652 | 1 |
| HDAC4 | 0.03623712 | 1 | 0.062187 | 1 |
| ARHGEF5 | -1.3844134 | 1 | 1.482906 | 1 |
| TIGD7 | 0.14923544 | 1 | -0.05065 | 1 |
| ACVR1 | 0.96173959 | 1 | -0.86311 | 1 |
| ZDHHC16 | 0.13052665 | 1 | -0.03178 | 1 |
| CACNB3 | 0.07486175 | 1 | 0.024366 | 1 |
| HEXIM1 | 0.10629853 | 1 | -0.00561 | 1 |
| OVOL2 | 0.03429777 | 1 | 0.066552 | 1 |
| OR13C8 | 0.0246731 | 1 | 0.076222 | 1 |
| FOXD4L3 | -0.0983763 | 1 | 0.200634 | 1 |
| PLCL2 | 0.34823024 | 1 | -0.24544 | 1 |
| SS18 | 0.07956687 | 1 | 0.023532 | 1 |
| BMP10 | 0.07128638 | 1 | 0.031875 | 1 |
| ZFP92 | -0.1494548 | 1 | 0.253228 | 1 |
| FGF11 | 0.00920807 | 1 | 0.094573 | 1 |
| TRIAP1 | 0.42681334 | 1 | -0.32301 | 1 |
| ZMYND15 | -0.106502 | 1 | 0.210319 | 1 |
| OR6C76 | 0.11937578 | 1 | -0.01533 | 1 |

| | | | | |
|---|---|---|---|---|
| CSH2 | 0.09415563 | 1 | 0.009989 | 1 |
| GABRA2 | 0.02879419 | 1 | 0.075496 | 1 |
| TP53BP1 | 0.00926095 | 1 | 0.09585 | 1 |
| PARP15 | -0.0258133 | 1 | 0.131927 | 1 |
| LILRB4 | -0.0426277 | 1 | 0.148746 | 1 |
| XPA | 0.24473574 | 1 | -0.13841 | 1 |
| CXCR2 | -0.011161 | 1 | 0.11762 | 1 |
| ZDHHC11 | -0.0382965 | 1 | 0.146648 | 1 |
| ANHX | 0.0676165 | 1 | 0.041259 | 1 |
| BID | -0.2290581 | 1 | 0.338173 | 1 |
| KDM4A | 0.39377198 | 1 | -0.28383 | 1 |
| OR4A15 | 0.04407071 | 1 | 0.066008 | 1 |
| ELF1 | -0.2232343 | 1 | 0.333596 | 1 |
| APH1A | 0.28122701 | 1 | -0.17035 | 1 |
| ARRB2 | -0.0437643 | 1 | 0.154794 | 1 |
| PLCB3 | 0.09572902 | 1 | 0.015747 | 1 |
| SAG | 0.03983136 | 1 | 0.07188 | 1 |
| GAP43 | 1.12336236 | 1 | -1.0116 | 1 |
| MAS1 | -0.0144881 | 1 | 0.126492 | 1 |
| RAP1GAP2 | 0.12535369 | 1 | -0.01292 | 1 |
| REM1 | 0.06397221 | 1 | 0.048828 | 1 |
| MYBBP1A | 0.07112064 | 1 | 0.041814 | 1 |
| LCK | -0.0552394 | 1 | 0.168712 | 1 |
| SSX1 | 0.10856641 | 1 | 0.004981 | 1 |
| DPRX | 0.05742054 | 1 | 0.056333 | 1 |
| RBL2 | 0.08520074 | 1 | 0.02883 | 1 |
| RBM15B | 0.10195043 | 1 | 0.012608 | 1 |
| IRX1 | -0.0026369 | 1 | 0.117246 | 1 |
| OPTN | 0.04386304 | 1 | 0.071051 | 1 |
| OR10H1 | 0.0627393 | 1 | 0.053292 | 1 |
| ZNF780B | -0.0025619 | 1 | 0.119535 | 1 |
| LZTS2 | 0.03687697 | 1 | 0.080369 | 1 |
| ADGRF1 | 0.03148444 | 1 | 0.085772 | 1 |
| CREB3L3 | -0.0678291 | 1 | 0.185461 | 1 |
| OR5A2 | 0.06615507 | 1 | 0.051776 | 1 |
| OR5M11 | 0.05187628 | 1 | 0.066063 | 1 |
| PTPRU | -0.5397433 | 1 | 0.65802 | 1 |
| L3MBTL4 | -0.1055619 | 1 | 0.225064 | 1 |
| ZNF346 | -0.0041212 | 1 | 0.124101 | 1 |
| TBX15 | -0.0275277 | 1 | 0.147754 | 1 |
| ZPR1 | 0.19266736 | 1 | -0.07195 | 1 |
| MS4A12 | 0.0635269 | 1 | 0.057869 | 1 |
| MUSK | 0.06463209 | 1 | 0.056794 | 1 |
| ZC3H7B | -0.0624209 | 1 | 0.184914 | 1 |
| ZHX2 | 0.01276707 | 1 | 0.109836 | 1 |
| GRK4 | 0.03054969 | 1 | 0.092852 | 1 |

| | | | | |
|---|---|---|---|---|
| OR10W1 | -0.0372968 | 1 | 0.160713 | 1 |
| CDC42BPB | 0.03070586 | 1 | 0.092829 | 1 |
| ZNF37A | 0.09741721 | 1 | 0.02649 | 1 |
| OSM | 0.09503311 | 1 | 0.029354 | 1 |
| CYTH4 | 0.0330009 | 1 | 0.091559 | 1 |
| ADAMTS20 | 0.04824944 | 1 | 0.076845 | 1 |
| HSFY2 | 0.03007694 | 1 | 0.095043 | 1 |
| IL22 | 0.0692588 | 1 | 0.056056 | 1 |
| ZNF302 | 0.36946255 | 1 | -0.24405 | 1 |
| CHRNB3 | -0.0069518 | 1 | 0.132656 | 1 |
| TIE1 | -0.0749536 | 1 | 0.200883 | 1 |
| ZNF804A | 0.8074955 | 1 | -0.68134 | 1 |
| RASGEF1C | -0.2353145 | 1 | 0.361767 | 1 |
| ZFAND2A | 0.16070837 | 1 | -0.0323 | 1 |
| IL21 | 0.06257298 | 1 | 0.065944 | 1 |
| LILRA1 | 0.00181293 | 1 | 0.128586 | 1 |
| PXN | 0.05647881 | 1 | 0.074015 | 1 |
| OR10H3 | 0.04654155 | 1 | 0.084403 | 1 |
| ZSCAN10 | 0.04763917 | 1 | 0.083382 | 1 |
| ZCCHC3 | 0.45527668 | 1 | -0.32414 | 1 |
| IL2 | 0.11003035 | 1 | 0.021276 | 1 |
| GPR89B | 0.05843011 | 1 | 0.072934 | 1 |
| SHOX | 0.01730153 | 1 | 0.114957 | 1 |
| HIF1AN | -0.0721547 | 1 | 0.204955 | 1 |
| NUP98 | 0.29158997 | 1 | -0.15859 | 1 |
| OR2J2 | 0.05867234 | 1 | 0.074694 | 1 |
| CNOT9 | 0.48819511 | 1 | -0.35477 | 1 |
| OR2H1 | 0.06115334 | 1 | 0.072693 | 1 |
| WNK1 | -0.0548885 | 1 | 0.188983 | 1 |
| MS4A14 | 0.0390595 | 1 | 0.095214 | 1 |
| OR1G1 | 0.05617491 | 1 | 0.078839 | 1 |
| OR1E1 | 0.03994588 | 1 | 0.095704 | 1 |
| TFAP4 | 0.11768715 | 1 | 0.018141 | 1 |
| PRDX3 | 1.48722901 | 1 | -1.35131 | 1 |
| RHOH | 0.14992017 | 1 | -0.01398 | 1 |
| IRF4 | 0.10888219 | 1 | 0.027365 | 1 |
| GNA14 | 0.10541035 | 1 | 0.031596 | 1 |
| OR4K13 | 0.11489641 | 1 | 0.02217 | 1 |
| EDN1 | 0.0716535 | 1 | 0.065581 | 1 |
| PDZD3 | 0.12685013 | 1 | 0.010512 | 1 |
| ZNF394 | 0.07102779 | 1 | 0.06719 | 1 |
| DMRTC2 | 0.17346739 | 1 | -0.03456 | 1 |
| ZNF844 | 0.06740254 | 1 | 0.0718 | 1 |
| PDE6C | -0.0017437 | 1 | 0.14103 | 1 |
| PPP1R10 | -0.049496 | 1 | 0.188848 | 1 |
| SYK | 0.18889693 | 1 | -0.04917 | 1 |

| | | | | |
|---|---|---|---|---|
| TMEM9B | 0.07363101 | 1 | 0.066383 | 1 |
| TBC1D22A | -0.0209749 | 1 | 0.162189 | 1 |
| CDC42EP4 | 0.03857187 | 1 | 0.102877 | 1 |
| EPOR | 0.06575305 | 1 | 0.075858 | 1 |
| MAF1 | 0.18713538 | 1 | -0.0453 | 1 |
| IL18RAP | 0.03546004 | 1 | 0.106704 | 1 |
| RTN2 | -0.3536738 | 1 | 0.496145 | 1 |
| HTR4 | 0.05539474 | 1 | 0.08754 | 1 |
| NKRF | 0.57782253 | 1 | -0.43459 | 1 |
| EPHA1 | 0.07282869 | 1 | 0.070909 | 1 |
| OR2F1 | -0.0775749 | 1 | 0.221498 | 1 |
| MT1H | 0.12753598 | 1 | 0.016928 | 1 |
| ESRRG | -0.1177245 | 1 | 0.262909 | 1 |
| ZNF789 | 0.0738426 | 1 | 0.071618 | 1 |
| RAB17 | 0.08002554 | 1 | 0.06567 | 1 |
| RAB11B | -0.0209015 | 1 | 0.166721 | 1 |
| ZC3H13 | 0.13889322 | 1 | 0.007202 | 1 |
| OR5H1 | 0.0530933 | 1 | 0.093224 | 1 |
| CHRNB1 | 0.08900375 | 1 | 0.05766 | 1 |
| OR2F2 | 0.13297638 | 1 | 0.014408 | 1 |
| ARFGAP2 | 0.06287998 | 1 | 0.084867 | 1 |
| LIFR | -1.3075054 | 1 | 1.455503 | 1 |
| DOT1L | 0.12210816 | 1 | 0.025972 | 1 |
| ZNF638 | 0.18038669 | 1 | -0.03156 | 1 |
| LPAR1 | -0.280184 | 1 | 0.429407 | 1 |
| ZNF235 | -0.0095931 | 1 | 0.158898 | 1 |
| OR1K1 | 0.07578988 | 1 | 0.074043 | 1 |
| H4C2 | 0.0748083 | 1 | 0.075121 | 1 |
| DDX1 | 0.55842755 | 1 | -0.4081 | 1 |
| PASD1 | 0.06847618 | 1 | 0.081974 | 1 |
| ZNF414 | 0.14326902 | 1 | 0.00725 | 1 |
| FOXD4L5 | -0.0237854 | 1 | 0.174721 | 1 |
| ASCL2 | 0.07664807 | 1 | 0.074807 | 1 |
| CALY | -0.0433396 | 1 | 0.194803 | 1 |
| CR1 | -0.0059188 | 1 | 0.157453 | 1 |
| LSP1 | 0.07056895 | 1 | 0.081387 | 1 |
| ZBTB49 | 0.16252496 | 1 | -0.01055 | 1 |
| FOXA3 | -0.8527985 | 1 | 1.004799 | 1 |
| CD33 | -0.0309286 | 1 | 0.182978 | 1 |
| ARHGAP8 | -0.0654129 | 1 | 0.217563 | 1 |
| ARAP2 | 0.11832478 | 1 | 0.034371 | 1 |
| APBB1IP | 0.03437447 | 1 | 0.119 | 1 |
| SREBF2 | -0.0561008 | 1 | 0.209614 | 1 |
| POLR3G | 0.18540178 | 1 | -0.03182 | 1 |
| DRD2 | -0.0240374 | 1 | 0.177622 | 1 |
| TAAR1 | 0.10578367 | 1 | 0.048033 | 1 |

| | | | | |
|---|---|---|---|---|
| ZFP41 | 0.09437111 | 1 | 0.059652 | 1 |
| ADCYAP1 | 0.22796708 | 1 | -0.07374 | 1 |
| CIR1 | 0.40047008 | 1 | -0.24598 | 1 |
| ARID4B | 0.41108026 | 1 | -0.25573 | 1 |
| NAE1 | 0.19327123 | 1 | -0.03783 | 1 |
| CD22 | -0.0402623 | 1 | 0.196556 | 1 |
| TCEAL6 | -0.0768647 | 1 | 0.233534 | 1 |
| KLRC1 | 0.08524111 | 1 | 0.071819 | 1 |
| OR4F3 | 0.08855179 | 1 | 0.068782 | 1 |
| ZUP1 | 0.55020189 | 1 | -0.39232 | 1 |
| RAB43 | -0.0555765 | 1 | 0.213462 | 1 |
| TRHR | 0.34467104 | 1 | -0.18628 | 1 |
| CXCR1 | -0.0958594 | 1 | 0.254657 | 1 |
| IKBKB | 0.09454347 | 1 | 0.064468 | 1 |
| OR7G1 | 0.25630255 | 1 | -0.09667 | 1 |
| CHRNE | 0.02938573 | 1 | 0.130267 | 1 |
| RASGRP1 | -0.1000227 | 1 | 0.259766 | 1 |
| ZNF34 | 0.1206133 | 1 | 0.039464 | 1 |
| CXCL16 | -0.0036347 | 1 | 0.16415 | 1 |
| NR1D2 | 0.11610202 | 1 | 0.045822 | 1 |
| SFMBT2 | 0.01593714 | 1 | 0.147128 | 1 |
| ROPN1B | 0.04449822 | 1 | 0.118677 | 1 |
| ARAP1 | -0.0513978 | 1 | 0.21501 | 1 |
| ZNF7 | 0.17852591 | 1 | -0.01461 | 1 |
| PLEKHH3 | -0.0414804 | 1 | 0.205412 | 1 |
| OR8A1 | 0.07833544 | 1 | 0.085823 | 1 |
| ASB3 | 0.0064842 | 1 | 0.158344 | 1 |
| MERTK | 0.04181153 | 1 | 0.123246 | 1 |
| OR4D10 | 0.0963264 | 1 | 0.069383 | 1 |
| MAPRE2 | 0.04065562 | 1 | 0.12572 | 1 |
| ZNF311 | 0.08095441 | 1 | 0.086434 | 1 |
| PECAM1 | 0.12973399 | 1 | 0.037756 | 1 |
| MGST3 | 0.09511923 | 1 | 0.072431 | 1 |
| FZD8 | -0.0120509 | 1 | 0.179737 | 1 |
| CNTN6 | 0.097253 | 1 | 0.070496 | 1 |
| AHRR | -0.0363132 | 1 | 0.204656 | 1 |
| EVI5L | -1.0255693 | 1 | 1.194581 | 1 |
| U2AF1 | 0.10676558 | 1 | 0.062295 | 1 |
| ITGAD | -0.0256165 | 1 | 0.194742 | 1 |
| WNK4 | 0.0804159 | 1 | 0.089494 | 1 |
| SUPT7L | 0.21421494 | 1 | -0.04409 | 1 |
| GPR141 | 0.09038437 | 1 | 0.080815 | 1 |
| ARHGDIB | 0.11402812 | 1 | 0.058085 | 1 |
| GREM1 | 0.05167692 | 1 | 0.120953 | 1 |
| ZNF230 | 0.0210995 | 1 | 0.152271 | 1 |
| U2AF2 | 0.08514737 | 1 | 0.088829 | 1 |

| | | | | |
|---|---|---|---|---|
| CTCFL | 0.15525688 | 1 | 0.018884 | 1 |
| PPRC1 | 0.12316027 | 1 | 0.05156 | 1 |
| GRK7 | 0.06645945 | 1 | 0.108529 | 1 |
| DDAH1 | 0.19011119 | 1 | -0.01511 | 1 |
| CLEC2D | 0.06341833 | 1 | 0.111867 | 1 |
| KCTD1 | 0.0524786 | 1 | 0.122911 | 1 |
| MEN1 | 0.17027416 | 1 | 0.005422 | 1 |
| OR11H6 | 0.06837552 | 1 | 0.107456 | 1 |
| ZDHHC12 | 0.06784896 | 1 | 0.108075 | 1 |
| OR13J1 | -0.0239193 | 1 | 0.200162 | 1 |
| MS4A4A | 0.11034531 | 1 | 0.06596 | 1 |
| IL9R | 0.03630397 | 1 | 0.140492 | 1 |
| ETS1 | 1.17449326 | 1 | -0.99663 | 1 |
| HSFY1 | 0.09295642 | 1 | 0.085048 | 1 |
| GPR6 | -0.0868654 | 1 | 0.265183 | 1 |
| OR1E2 | 0.10564186 | 1 | 0.073171 | 1 |
| ZNF83 | 0.03995008 | 1 | 0.138891 | 1 |
| FGF4 | 0.13537581 | 1 | 0.043646 | 1 |
| ZNF396 | 0.00667728 | 1 | 0.172498 | 1 |
| CCR10 | -0.0956269 | 1 | 0.275234 | 1 |
| TOP2B | 0.5463834 | 1 | -0.36667 | 1 |
| REPS2 | -1.562086 | 1 | 1.741878 | 1 |
| INPP5B | 0.10093646 | 1 | 0.0789 | 1 |
| SIRT3 | -0.013281 | 1 | 0.193395 | 1 |
| TBP | 0.13890574 | 1 | 0.041604 | 1 |
| INS | 0.06586166 | 1 | 0.114706 | 1 |
| CIZ1 | -0.0009901 | 1 | 0.181595 | 1 |
| ZNHIT2 | 0.20967479 | 1 | -0.02867 | 1 |
| DZIP3 | 0.30935431 | 1 | -0.12796 | 1 |
| GPR68 | 0.09870934 | 1 | 0.082782 | 1 |
| ZNF577 | 0.08372892 | 1 | 0.09779 | 1 |
| JAK3 | -0.0302417 | 1 | 0.211999 | 1 |
| FLI1 | 0.0195784 | 1 | 0.162297 | 1 |
| PSD | -0.0299929 | 1 | 0.212192 | 1 |
| AGRP | -0.0145292 | 1 | 0.196738 | 1 |
| ZNF211 | 0.05610456 | 1 | 0.126188 | 1 |
| ARHGEF40 | -1.401014 | 1 | 1.583578 | 1 |
| ZNF841 | 0.08850732 | 1 | 0.094106 | 1 |
| NLRC4 | -0.158341 | 1 | 0.341156 | 1 |
| NHLH2 | 0.00206671 | 1 | 0.181206 | 1 |
| ALS2 | 0.16964434 | 1 | 0.013832 | 1 |
| ACIN1 | -0.0322408 | 1 | 0.216548 | 1 |
| ISL2 | 0.1652375 | 1 | 0.019129 | 1 |
| SDHC | 0.13170594 | 1 | 0.05285 | 1 |
| CERS2 | 0.05024822 | 1 | 0.134565 | 1 |
| ZNF189 | 0.12151146 | 1 | 0.063504 | 1 |

| | | | | |
|---|---|---|---|---|
| TSFM | 0.71567691 | 1 | -0.53046 | 1 |
| ZCCHC2 | 0.23124002 | 1 | -0.04513 | 1 |
| PNN | 0.31023766 | 1 | -0.12352 | 1 |
| ARHGEF6 | 0.17474249 | 1 | 0.012036 | 1 |
| GRIN2A | 0.06867964 | 1 | 0.118208 | 1 |
| OR8B2 | 0.11122842 | 1 | 0.075727 | 1 |
| ARHGAP21 | -0.1452909 | 1 | 0.332307 | 1 |
| ZNF582 | 0.09336536 | 1 | 0.093719 | 1 |
| TENM4 | 0.03464383 | 1 | 0.152787 | 1 |
| PLXNA4 | 0.07416394 | 1 | 0.113303 | 1 |
| DDAH2 | 0.14680872 | 1 | 0.04133 | 1 |
| ACKR1 | -0.0321105 | 1 | 0.220296 | 1 |
| OR10H4 | 0.0796618 | 1 | 0.108849 | 1 |
| PTK2B | 0.14570982 | 1 | 0.044026 | 1 |
| IRF8 | 0.10765562 | 1 | 0.082803 | 1 |
| OPN1LW | 0.03233687 | 1 | 0.158412 | 1 |
| HSF4 | -0.1036924 | 1 | 0.294889 | 1 |
| GNB2 | 1.18373493 | 1 | -0.99222 | 1 |
| RP1L1 | 0.1235488 | 1 | 0.068729 | 1 |
| TNFRSF8 | -0.1254692 | 1 | 0.317964 | 1 |
| POMC | 0.10964767 | 1 | 0.083014 | 1 |
| OR5M3 | 0.09844017 | 1 | 0.095182 | 1 |
| RHO | 0.10033338 | 1 | 0.093558 | 1 |
| CD3E | -0.0281843 | 1 | 0.223599 | 1 |
| DNAJB6 | 0.45778534 | 1 | -0.26175 | 1 |
| MPL | 0.0636494 | 1 | 0.134428 | 1 |
| IL1A | 0.00300131 | 1 | 0.195562 | 1 |
| TBC1D20 | 0.12944805 | 1 | 0.069318 | 1 |
| ZSWIM2 | 0.01580593 | 1 | 0.182979 | 1 |
| TLX1 | 0.11602687 | 1 | 0.083134 | 1 |
| VPS25 | 0.35739163 | 1 | -0.15821 | 1 |
| HNF1B | -0.0065408 | 1 | 0.206516 | 1 |
| OMD | 0.19046677 | 1 | 0.010176 | 1 |
| OR2A4 | 0.02747945 | 1 | 0.173388 | 1 |
| UTS2R | 0.08797998 | 1 | 0.113084 | 1 |
| BMX | 0.03654206 | 1 | 0.164569 | 1 |
| EFCAB6 | 0.07935774 | 1 | 0.122586 | 1 |
| OR4C13 | -0.057928 | 1 | 0.259923 | 1 |
| SPRED2 | 1.60714618 | 1 | -1.40515 | 1 |
| MAPK8IP1 | -0.0983638 | 1 | 0.300405 | 1 |
| ZNF222 | 0.17870602 | 1 | 0.024462 | 1 |
| EBF4 | 0.04868457 | 1 | 0.154684 | 1 |
| MGA | 0.22839676 | 1 | -0.02493 | 1 |
| ZNF169 | -0.0088623 | 1 | 0.212715 | 1 |
| XRCC6 | 0.67388656 | 1 | -0.46931 | 1 |
| RASSF4 | -0.0565923 | 1 | 0.261719 | 1 |

| | | | | |
|---|---|---|---|---|
| FEZF2 | -0.0225591 | 1 | 0.228033 | 1 |
| OR8B3 | 0.12403879 | 1 | 0.082709 | 1 |
| ARHGEF16 | 0.00884472 | 1 | 0.198078 | 1 |
| NR1I3 | -0.251523 | 1 | 0.458475 | 1 |
| RABEP1 | 0.83667026 | 1 | -0.62961 | 1 |
| ZNF282 | 0.22539967 | 1 | -0.01776 | 1 |
| ZNF461 | 0.22583558 | 1 | -0.01747 | 1 |
| OR4A47 | 0.09680582 | 1 | 0.111828 | 1 |
| CNBP | 0.86143568 | 1 | -0.65275 | 1 |
| OR51B5 | 0.0553654 | 1 | 0.153393 | 1 |
| CSNK1E | 0.13160258 | 1 | 0.077307 | 1 |
| ARHGEF15 | 0.03062091 | 1 | 0.178289 | 1 |
| BMI1 | 0.19917877 | 1 | 0.010352 | 1 |
| ZNF674 | 0.19349954 | 1 | 0.017195 | 1 |
| HOXB7 | 0.10893108 | 1 | 0.101915 | 1 |
| ZNF793 | 0.1285705 | 1 | 0.082478 | 1 |
| ARHGAP24 | 0.0640601 | 1 | 0.146995 | 1 |
| ADCY6 | 0.12307635 | 1 | 0.088153 | 1 |
| TBC1D3C | 0.11464566 | 1 | 0.096961 | 1 |
| CXCL3 | 0.05828772 | 1 | 0.153845 | 1 |
| PHF19 | 0.07123654 | 1 | 0.141525 | 1 |
| CLTCL1 | 0.09760331 | 1 | 0.115607 | 1 |
| PLEKHM3 | -0.7731225 | 1 | 0.986362 | 1 |
| SEBOX | 0.08536442 | 1 | 0.12796 | 1 |
| PTOV1 | 0.01871578 | 1 | 0.194657 | 1 |
| FGF6 | 0.0637618 | 1 | 0.150105 | 1 |
| ZNF546 | 0.05242042 | 1 | 0.161764 | 1 |
| RABL2B | 0.09672651 | 1 | 0.118276 | 1 |
| SIRPG | 0.09883328 | 1 | 0.116178 | 1 |
| ADIPOR2 | -0.0767304 | 1 | 0.291921 | 1 |
| HOXA13 | 0.18413213 | 1 | 0.032024 | 1 |
| VAV1 | 0.16623328 | 1 | 0.050002 | 1 |
| YY1AP1 | 0.21776909 | 1 | -0.00119 | 1 |
| CDK8 | 0.45310536 | 1 | -0.2363 | 1 |
| RAB25 | 0.10454449 | 1 | 0.112616 | 1 |
| CCNC | 1.49419403 | 1 | -1.27566 | 1 |
| RET | 0.07920776 | 1 | 0.139363 | 1 |
| ARGFX | 0.01528748 | 1 | 0.20358 | 1 |
| ID1 | 0.05421466 | 1 | 0.165009 | 1 |
| CHRNG | 0.26694582 | 1 | -0.04752 | 1 |
| CLEC1A | 0.08384857 | 1 | 0.135778 | 1 |
| KRBA2 | -0.0357677 | 1 | 0.25614 | 1 |
| OR8D2 | 0.09221072 | 1 | 0.12831 | 1 |
| CNTF | -0.1387727 | 1 | 0.359652 | 1 |
| DRD5 | 0.11813633 | 1 | 0.103155 | 1 |
| PKD1L3 | 0.0764593 | 1 | 0.14503 | 1 |

| | | | | |
|---|---|---|---|---|
| OR13G1 | 0.04373301 | 1 | 0.179033 | 1 |
| CAND2 | 0.00479766 | 1 | 0.21817 | 1 |
| MBD3L4 | 0.09355783 | 1 | 0.129479 | 1 |
| OR51F1 | 0.04268273 | 1 | 0.181032 | 1 |
| OR5D14 | 0.1350196 | 1 | 0.089376 | 1 |
| FGF23 | 0.05002689 | 1 | 0.176104 | 1 |
| GLMP | 0.02254411 | 1 | 0.204027 | 1 |
| CITED2 | 0.15700548 | 1 | 0.069738 | 1 |
| PAFAH1B2 | 1.86920147 | 1 | -1.64228 | 1 |
| RASSF8 | 0.17847131 | 1 | 0.048451 | 1 |
| IRF5 | 0.09423201 | 1 | 0.133798 | 1 |
| ADRB3 | 0.0077044 | 1 | 0.22046 | 1 |
| ZNF574 | 0.14061168 | 1 | 0.087597 | 1 |
| GRM8 | 0.31676936 | 1 | -0.08809 | 1 |
| EXT2 | 0.52906486 | 1 | -0.29978 | 1 |
| GRK2 | 0.07281992 | 1 | 0.157579 | 1 |
| CRHBP | -0.1258889 | 1 | 0.356419 | 1 |
| CARD19 | 0.00661154 | 1 | 0.224012 | 1 |
| SOD1 | 0.14792858 | 1 | 0.082697 | 1 |
| RIPK1 | 0.18252067 | 1 | 0.048264 | 1 |
| FZD6 | 0.16159119 | 1 | 0.069274 | 1 |
| RELN | 0.06981397 | 1 | 0.161212 | 1 |
| ARHGEF4 | 0.06347876 | 1 | 0.167859 | 1 |
| GNG8 | 0.07416913 | 1 | 0.157284 | 1 |
| OR2AG1 | 0.0550797 | 1 | 0.177036 | 1 |
| DFFB | 0.10040632 | 1 | 0.13186 | 1 |
| WDR90 | 0.05264093 | 1 | 0.180007 | 1 |
| CREB3 | 0.22934993 | 1 | 0.003399 | 1 |
| IL17C | -0.0284241 | 1 | 0.261944 | 1 |
| LTA | 0.05275036 | 1 | 0.18103 | 1 |
| OR5B21 | 0.0166032 | 1 | 0.218189 | 1 |
| GSX1 | 0.02430385 | 1 | 0.211197 | 1 |
| PDX1 | 0.09248745 | 1 | 0.143172 | 1 |
| NTSR2 | -0.025028 | 1 | 0.261123 | 1 |
| SOX13 | -0.2482978 | 1 | 0.484977 | 1 |
| TAS2R4 | 0.12033331 | 1 | 0.116356 | 1 |
| IL1RAPL2 | 0.10428603 | 1 | 0.132842 | 1 |
| CCR5 | -0.1410347 | 1 | 0.378825 | 1 |
| MSH3 | 0.56641176 | 1 | -0.32854 | 1 |
| LHX9 | 0.02890151 | 1 | 0.209245 | 1 |
| OR5M9 | 0.14659927 | 1 | 0.091719 | 1 |
| ZNF41 | 0.34945889 | 1 | -0.11061 | 1 |
| CSDC2 | 0.02020927 | 1 | 0.218834 | 1 |
| CHD2 | 0.34422016 | 1 | -0.10485 | 1 |
| KIF26A | -0.6913296 | 1 | 0.931306 | 1 |
| OR10A4 | 0.21301574 | 1 | 0.027519 | 1 |

| | | | | |
|---|---|---|---|---|
| SMPD2 | 0.07301757 | 1 | 0.168051 | 1 |
| TRAF3IP2 | 0.18543833 | 1 | 0.055726 | 1 |
| OR1N1 | 0.12604398 | 1 | 0.115497 | 1 |
| PTPRN2 | -0.0754083 | 1 | 0.317671 | 1 |
| TBC1D4 | -0.8846143 | 1 | 1.127142 | 1 |
| HOXD3 | 0.22551543 | 1 | 0.017454 | 1 |
| ZNF18 | 0.11356088 | 1 | 0.130387 | 1 |
| ZDHHC8 | 0.0621415 | 1 | 0.182117 | 1 |
| RGR | -0.7263198 | 1 | 0.97073 | 1 |
| BHLHA15 | 0.01350999 | 1 | 0.231479 | 1 |
| ORC2 | 0.20726777 | 1 | 0.038086 | 1 |
| ANK2 | 0.07717879 | 1 | 0.16836 | 1 |
| NCKIPSD | 0.08416677 | 1 | 0.161435 | 1 |
| ANGPTL3 | 0.28824128 | 1 | -0.04229 | 1 |
| GTF2IRD2B | -0.0580385 | 1 | 0.304243 | 1 |
| KCNH2 | 0.09960345 | 1 | 0.146875 | 1 |
| ZSWIM1 | 0.14815055 | 1 | 0.098662 | 1 |
| ARL11 | 0.15717712 | 1 | 0.090211 | 1 |
| ADCY7 | 0.04847878 | 1 | 0.199439 | 1 |
| KLF2 | -0.653941 | 1 | 0.902029 | 1 |
| PRLHR | 0.08070828 | 1 | 0.16769 | 1 |
| ZSWIM7 | -0.2083122 | 1 | 0.456828 | 1 |
| ZNF821 | 0.27401688 | 1 | -0.02484 | 1 |
| TBC1D5 | 0.10301742 | 1 | 0.146306 | 1 |
| ZBTB7C | 0.00556315 | 1 | 0.243804 | 1 |
| ARID3C | 0.11191671 | 1 | 0.137672 | 1 |
| OR5L1 | 0.11152949 | 1 | 0.138231 | 1 |
| CYTH1 | -0.0076786 | 1 | 0.257583 | 1 |
| PDE4A | 0.09925215 | 1 | 0.150744 | 1 |
| PADI4 | -0.025851 | 1 | 0.276031 | 1 |
| OR5H14 | 0.07328778 | 1 | 0.177055 | 1 |
| GPR156 | -0.1864148 | 1 | 0.436945 | 1 |
| RND1 | 0.01516199 | 1 | 0.235396 | 1 |
| LILRB2 | -0.0062991 | 1 | 0.257302 | 1 |
| PAX1 | -0.0308625 | 1 | 0.282449 | 1 |
| AKAP9 | 0.33889825 | 1 | -0.08655 | 1 |
| MED22 | 0.08012265 | 1 | 0.172378 | 1 |
| GRB7 | 0.09763187 | 1 | 0.155046 | 1 |
| PTH | 0.11799243 | 1 | 0.134764 | 1 |
| FCER1G | 0.16811176 | 1 | 0.084655 | 1 |
| CDX4 | 0.1666269 | 1 | 0.086353 | 1 |
| BCAM | 0.02688997 | 1 | 0.226335 | 1 |
| PMEPA1 | 1.31585047 | 1 | -1.06135 | 1 |
| ZBTB3 | 0.04955176 | 1 | 0.206228 | 1 |
| TADA2B | 0.13027621 | 1 | 0.125919 | 1 |
| MAEL | -0.1424853 | 1 | 0.398743 | 1 |

| | | | | |
|---|---|---|---|---|
| ZNF133 | 0.22412293 | 1 | 0.032141 | 1 |
| ZNF548 | 0.36808843 | 1 | -0.11175 | 1 |
| RHPN1 | -0.0792293 | 1 | 0.335746 | 1 |
| SOX1 | 0.20809593 | 1 | 0.049413 | 1 |
| ASB16 | 0.03777605 | 1 | 0.219812 | 1 |
| TBC1D10A | 0.13759194 | 1 | 0.121543 | 1 |
| GNAL | -0.0959681 | 1 | 0.355527 | 1 |
| ADCY2 | 0.04889999 | 1 | 0.210751 | 1 |
| CFLAR | 0.19681573 | 1 | 0.063329 | 1 |
| PTGDR | 0.09266999 | 1 | 0.167794 | 1 |
| ZNF768 | 0.15437301 | 1 | 0.106623 | 1 |
| IL6ST | 0.06829708 | 1 | 0.19307 | 1 |
| ZNF333 | 0.39858612 | 1 | -0.13702 | 1 |
| GHRHR | 0.03250294 | 1 | 0.229114 | 1 |
| PLEK2 | 0.17895219 | 1 | 0.082686 | 1 |
| HSPE1 | 0.37015795 | 1 | -0.10847 | 1 |
| RGS2 | 1.30698046 | 1 | -1.04491 | 1 |
| GNB1L | 0.14974948 | 1 | 0.112372 | 1 |
| DCDC2C | 0.10594738 | 1 | 0.156629 | 1 |
| RPTOR | 1.1057816 | 1 | -0.84308 | 1 |
| TXNDC12 | 1.18027209 | 1 | -0.91685 | 1 |
| VIP | 0.23357538 | 1 | 0.029912 | 1 |
| MC2R | 0.16313844 | 1 | 0.101142 | 1 |
| PRKCZ | -0.0027219 | 1 | 0.267455 | 1 |
| ARHGAP30 | 0.08557109 | 1 | 0.179409 | 1 |
| SCAP | 0.14823559 | 1 | 0.117038 | 1 |
| ARF3 | 0.13813208 | 1 | 0.127181 | 1 |
| ITGA7 | -0.7378024 | 1 | 1.00349 | 1 |
| TNFRSF17 | 0.12852266 | 1 | 0.137352 | 1 |
| TNFSF4 | 0.10160412 | 1 | 0.164803 | 1 |
| PTPN22 | 0.27972565 | 1 | -0.01317 | 1 |
| DAPK1 | 0.0565687 | 1 | 0.20999 | 1 |
| NPR2 | 0.0745919 | 1 | 0.192543 | 1 |
| CALCOCO1 | 0.01941434 | 1 | 0.248993 | 1 |
| OR52E5 | 0.011412 | 1 | 0.258013 | 1 |
| OR10R2 | 0.17852435 | 1 | 0.090917 | 1 |
| NTRK1 | -0.0289435 | 1 | 0.298944 | 1 |
| CGA | 0.10785182 | 1 | 0.162963 | 1 |
| ZCCHC14 | 0.17024036 | 1 | 0.100844 | 1 |
| OR51D1 | 0.02130092 | 1 | 0.249801 | 1 |
| FOXB2 | 0.20674479 | 1 | 0.065371 | 1 |
| RGS3 | 0.13040896 | 1 | 0.142692 | 1 |
| DMRTA1 | 0.21063251 | 1 | 0.062619 | 1 |
| CNTN1 | -1.6860162 | 1 | 1.959601 | 1 |
| ZNF134 | 0.5183522 | 1 | -0.2444 | 1 |
| NRL | 0.0380027 | 1 | 0.236081 | 1 |

| | | | | |
|---|---|---|---|---|
| BAIAP3 | 0.02402863 | 1 | 0.250123 | 1 |
| RIPPLY1 | 0.06944843 | 1 | 0.204791 | 1 |
| ZNF19 | 0.33338013 | 1 | -0.05825 | 1 |
| CEACAM1 | 0.05486207 | 1 | 0.220307 | 1 |
| IL12RB1 | 0.11691006 | 1 | 0.158312 | 1 |
| SIRT4 | 0.02049063 | 1 | 0.254768 | 1 |
| MAPK11 | 0.00137133 | 1 | 0.274279 | 1 |
| ZMYND10 | 0.08171779 | 1 | 0.194627 | 1 |
| ZIM3 | 0.0924327 | 1 | 0.183996 | 1 |
| OR6V1 | 0.11706674 | 1 | 0.159891 | 1 |
| TRIM28 | 0.12542546 | 1 | 0.151681 | 1 |
| OR2AG2 | -0.0645189 | 1 | 0.341881 | 1 |
| TLE3 | -0.776965 | 1 | 1.054547 | 1 |
| RASA3 | 0.01338364 | 1 | 0.264419 | 1 |
| PKD1 | 0.07132084 | 1 | 0.206884 | 1 |
| TPTE | 0.1162519 | 1 | 0.161968 | 1 |
| ID3 | 0.10858422 | 1 | 0.170065 | 1 |
| TBC1D24 | 0.20562416 | 1 | 0.073076 | 1 |
| GPS1 | 0.32866258 | 1 | -0.04983 | 1 |
| HTR2B | 0.26460847 | 1 | 0.014579 | 1 |
| MST1R | 0.02256726 | 1 | 0.25748 | 1 |
| ZNF621 | 0.07563109 | 1 | 0.204657 | 1 |
| ZNF567 | 1.28956723 | 1 | -1.00746 | 1 |
| ADGRF5 | -0.1261985 | 1 | 0.408684 | 1 |
| OR4F21 | 0.07066619 | 1 | 0.212524 | 1 |
| OR6C3 | 0.22983603 | 1 | 0.054197 | 1 |
| P2RX1 | 0.03305592 | 1 | 0.252581 | 1 |
| OR2V2 | 0.08667775 | 1 | 0.199149 | 1 |
| BTRC | -0.0213593 | 1 | 0.307913 | 1 |
| SNW1 | 0.54441304 | 1 | -0.25775 | 1 |
| SH3BP1 | 0.08843121 | 1 | 0.198238 | 1 |
| CIC | 0.02129891 | 1 | 0.26584 | 1 |
| OR52A1 | 0.00133002 | 1 | 0.285915 | 1 |
| DIRAS1 | -1.2177004 | 1 | 1.506685 | 1 |
| DBX2 | 0.39519201 | 1 | -0.10571 | 1 |
| BCL2L1 | 0.15713354 | 1 | 0.132677 | 1 |
| NOTCH2 | -0.4519772 | 1 | 0.742245 | 1 |
| PRKAR1B | -0.0906446 | 1 | 0.381093 | 1 |
| HHIP | 0.20205864 | 1 | 0.088783 | 1 |
| RASSF3 | 0.15928466 | 1 | 0.131625 | 1 |
| ZBBX | 0.11767607 | 1 | 0.173236 | 1 |
| ZFP64 | 0.42609312 | 1 | -0.13379 | 1 |
| PDE4C | 0.1051369 | 1 | 0.187537 | 1 |
| CGB3 | 0.93777805 | 1 | -0.6443 | 1 |
| PPIE | 0.3270916 | 1 | -0.03309 | 1 |
| SIM2 | 0.11636119 | 1 | 0.177815 | 1 |

| | | | | |
|---|---|---|---|---|
| ZBED9 | 0.00946418 | 1 | 0.284986 | 1 |
| SUFU | -0.0549364 | 1 | 0.349674 | 1 |
| KIR2DS1 | 0.17063869 | 1 | 0.124783 | 1 |
| ZNF613 | 0.09529519 | 1 | 0.200207 | 1 |
| CHD3 | 0.10488585 | 1 | 0.19128 | 1 |
| TSC1 | 0.10665722 | 1 | 0.190044 | 1 |
| TLR2 | 0.00307032 | 1 | 0.294528 | 1 |
| FEZF1 | -0.1835398 | 1 | 0.481709 | 1 |
| VN1R2 | 0.10404643 | 1 | 0.194449 | 1 |
| DOK3 | -0.0201017 | 1 | 0.319125 | 1 |
| OXER1 | 0.10797167 | 1 | 0.191108 | 1 |
| GABRR2 | 0.11779147 | 1 | 0.181652 | 1 |
| PGAP2 | 0.13217527 | 1 | 0.168485 | 1 |
| ABCG4 | -0.0713172 | 1 | 0.372248 | 1 |
| TSPO | 0.11972605 | 1 | 0.181294 | 1 |
| AZI2 | 0.47330519 | 1 | -0.17217 | 1 |
| CIAO3 | -0.0457413 | 1 | 0.347608 | 1 |
| PAX5 | 0.14890151 | 1 | 0.153379 | 1 |
| MED20 | 0.50743172 | 1 | -0.20497 | 1 |
| PTPN6 | 0.23829597 | 1 | 0.064765 | 1 |
| MAP4K1 | 0.02033176 | 1 | 0.282909 | 1 |
| OR2T8 | 0.29388303 | 1 | 0.00936 | 1 |
| BDKRB2 | 0.19080697 | 1 | 0.113365 | 1 |
| TPM3 | 0.53429331 | 1 | -0.22945 | 1 |
| DEDD | 0.52860723 | 1 | -0.22323 | 1 |
| ICOSLG | -0.0076612 | 1 | 0.313357 | 1 |
| SH3GL3 | -0.1031247 | 1 | 0.409443 | 1 |
| OR4D11 | 0.20377183 | 1 | 0.102563 | 1 |
| CACNA1A | 0.15364002 | 1 | 0.15347 | 1 |
| ARHGAP18 | 0.20551301 | 1 | 0.102517 | 1 |
| BLNK | 0.11699872 | 1 | 0.191281 | 1 |
| EREG | 0.18108855 | 1 | 0.127332 | 1 |
| TNFSF8 | 0.21372344 | 1 | 0.09474 | 1 |
| ARHGAP20 | 0.15628975 | 1 | 0.152463 | 1 |
| CFL1 | 0.34484421 | 1 | -0.03605 | 1 |
| INSM2 | -0.9556363 | 1 | 1.264441 | 1 |
| ZNF775 | 0.18774173 | 1 | 0.121302 | 1 |
| NDRG1 | 0.24495917 | 1 | 0.064431 | 1 |
| ZNF44 | 0.26703628 | 1 | 0.043458 | 1 |
| ATF7 | 0.17966975 | 1 | 0.131293 | 1 |
| GATAD2B | 0.46828717 | 1 | -0.15703 | 1 |
| P2RY6 | 0.68022782 | 1 | -0.36894 | 1 |
| TAAR9 | 0.90347018 | 1 | -0.59188 | 1 |
| DCDC2 | 0.06286996 | 1 | 0.248897 | 1 |
| ZSCAN9 | 0.50290382 | 1 | -0.18889 | 1 |
| TGFBR3 | 0.22920045 | 1 | 0.0854 | 1 |

| | | | | |
|---|---|---|---|---|
| SIGIRR | 0.11487725 | 1 | 0.200589 | 1 |
| LHX8 | 0.36676322 | 1 | -0.04995 | 1 |
| RABL2A | 0.19854764 | 1 | 0.118365 | 1 |
| GPR183 | 0.05930695 | 1 | 0.257741 | 1 |
| TAS2R46 | 0.197939 | 1 | 0.119271 | 1 |
| ITK | 0.2669974 | 1 | 0.050503 | 1 |
| CCR8 | 0.1547753 | 1 | 0.162797 | 1 |
| POLR1G | 0.19106599 | 1 | 0.126949 | 1 |
| BRDT | 0.06629046 | 1 | 0.253079 | 1 |
| AVPR2 | 0.15431998 | 1 | 0.165336 | 1 |
| RXFP3 | -1.1070214 | 1 | 1.42713 | 1 |
| OR52L1 | 0.10854968 | 1 | 0.21157 | 1 |
| TBC1D3H | 0.0474574 | 1 | 0.272986 | 1 |
| ZC3H10 | 0.2325858 | 1 | 0.088121 | 1 |
| IL2RB | 0.24183323 | 1 | 0.0794 | 1 |
| SSBP4 | 0.20668206 | 1 | 0.114736 | 1 |
| ZNF135 | 0.13711531 | 1 | 0.184317 | 1 |
| ZNF207 | 0.54087436 | 1 | -0.2184 | 1 |
| CDC42EP1 | 0.08366328 | 1 | 0.24041 | 1 |
| OR9A2 | -0.0353527 | 1 | 0.359546 | 1 |
| EBI3 | 0.13423359 | 1 | 0.190185 | 1 |
| EHMT1 | 0.05084685 | 1 | 0.273668 | 1 |
| PER1 | -0.0745632 | 1 | 0.39913 | 1 |
| PPP1R8 | 0.37159311 | 1 | -0.04591 | 1 |
| PLA2G4B | 0.06972796 | 1 | 0.256039 | 1 |
| OR52M1 | 0.0940487 | 1 | 0.232431 | 1 |
| BAIAP2L2 | 0.11680253 | 1 | 0.2104 | 1 |
| TAS2R41 | 0.11164821 | 1 | 0.215822 | 1 |
| ADGRL1 | -1.0608671 | 1 | 1.389101 | 1 |
| FGF8 | 0.18346754 | 1 | 0.145009 | 1 |
| ARHGAP45 | 0.13916344 | 1 | 0.189914 | 1 |
| VENTX | -0.027682 | 1 | 0.356764 | 1 |
| APOE | 0.15740463 | 1 | 0.172977 | 1 |
| ITGB2 | 0.19013293 | 1 | 0.141393 | 1 |
| PRDX2 | 0.30999488 | 1 | 0.02254 | 1 |
| MNT | 0.09749545 | 1 | 0.235048 | 1 |
| PTPRR | -0.2714441 | 1 | 0.605311 | 1 |
| OR52D1 | 0.16396256 | 1 | 0.170257 | 1 |
| ZNF223 | -0.4879509 | 1 | 0.822447 | 1 |
| HMX1 | 0.10634657 | 1 | 0.228766 | 1 |
| SAP30BP | 0.17087235 | 1 | 0.164759 | 1 |
| TLR6 | 0.1094276 | 1 | 0.226226 | 1 |
| BMP4 | 0.28982609 | 1 | 0.046169 | 1 |
| THRB | 0.13565024 | 1 | 0.201063 | 1 |
| FOXE3 | 0.3474377 | 1 | -0.01066 | 1 |
| IGF2R | 0.22262 | 1 | 0.114693 | 1 |

| | | | | |
|---|---|---|---|---|
| OR52B2 | 1.40613296 | 1 | -1.06866 | 1 |
| FGF20 | 0.15771386 | 1 | 0.180686 | 1 |
| OR10D3 | 0.22118571 | 1 | 0.117871 | 1 |
| CLDN3 | 0.1148445 | 1 | 0.2253 | 1 |
| JMJD7.PLA2G4B | -0.034275 | 1 | 0.374515 | 1 |
| ACTN2 | 0.24196722 | 1 | 0.098756 | 1 |
| DIABLO | 0.37299133 | 1 | -0.03107 | 1 |
| SKOR2 | 0.17953672 | 1 | 0.162899 | 1 |
| TGFB3 | 0.21040056 | 1 | 0.133052 | 1 |
| PAK4 | 0.04630766 | 1 | 0.297236 | 1 |
| CD79A | 0.25850673 | 1 | 0.08535 | 1 |
| NCOA1 | -0.472591 | 1 | 0.816779 | 1 |
| OR2T34 | 0.15613796 | 1 | 0.188068 | 1 |
| ADAP2 | 0.16021462 | 1 | 0.184004 | 1 |
| RGS5 | 0.17873407 | 1 | 0.165623 | 1 |
| HNF1A | 0.17461478 | 1 | 0.171325 | 1 |
| ZBTB8OS | 0.25253636 | 1 | 0.0948 | 1 |
| TGIF2 | 0.18421187 | 1 | 0.164473 | 1 |
| MBD1 | 0.13633046 | 1 | 0.212702 | 1 |
| PRKAB2 | 0.44297071 | 1 | -0.09365 | 1 |
| HSBP1 | 0.54130322 | 1 | -0.19068 | 1 |
| RAB22A | 0.41136319 | 1 | -0.06054 | 1 |
| CHRNA2 | 0.19006361 | 1 | 0.160948 | 1 |
| KMT5C | 0.16588688 | 1 | 0.18537 | 1 |
| TBR1 | 0.03941185 | 1 | 0.313611 | 1 |
| ZNF251 | 0.18295227 | 1 | 0.170472 | 1 |
| DEDD2 | 0.14300997 | 1 | 0.210483 | 1 |
| CTNNAL1 | 0.58475192 | 1 | -0.23112 | 1 |
| CARHSP1 | 0.2493278 | 1 | 0.10431 | 1 |
| HERPUD1 | 0.1117979 | 1 | 0.242111 | 1 |
| PITX2 | 0.05063496 | 1 | 0.303503 | 1 |
| PI4KB | 0.18334021 | 1 | 0.171163 | 1 |
| CDK11A | 0.17801476 | 1 | 0.176921 | 1 |
| CDIP1 | 0.02105009 | 1 | 0.33394 | 1 |
| OR10C1 | 0.18325119 | 1 | 0.172565 | 1 |
| ZFPM1 | 0.18532875 | 1 | 0.171025 | 1 |
| POU6F1 | 0.0873156 | 1 | 0.269411 | 1 |
| ZNF655 | 0.16521458 | 1 | 0.193019 | 1 |
| NR1H4 | 0.24850866 | 1 | 0.110319 | 1 |
| ESRRB | 0.14706652 | 1 | 0.211779 | 1 |
| GP6 | 0.06708617 | 1 | 0.292109 | 1 |
| OR6A2 | 0.31613916 | 1 | 0.043332 | 1 |
| SORCS2 | 0.09399631 | 1 | 0.265587 | 1 |
| BUD31 | 0.32289757 | 1 | 0.037739 | 1 |
| TG | 0.60706076 | 1 | -0.24579 | 1 |
| ZNF658 | 0.24978248 | 1 | 0.111755 | 1 |

| | | | | |
|---|---|---|---|---|
| NOTO | 0.19432152 | 1 | 0.16739 | 1 |
| FGFR4 | 0.12143292 | 1 | 0.240879 | 1 |
| USP20 | 0.07128133 | 1 | 0.291894 | 1 |
| IL17RC | -0.0086776 | 1 | 0.373432 | 1 |
| BRD4 | 0.28137849 | 1 | 0.083916 | 1 |
| PRKCH | 0.19887947 | 1 | 0.166615 | 1 |
| GARNL3 | 0.15923951 | 1 | 0.206428 | 1 |
| CAPN3 | 0.09277429 | 1 | 0.273799 | 1 |
| PSTPIP1 | 0.35525647 | 1 | 0.011465 | 1 |
| ZFC3H1 | 0.14636978 | 1 | 0.220628 | 1 |
| MOK | 0.05210319 | 1 | 0.315405 | 1 |
| ASB10 | 0.5625603 | 1 | -0.19407 | 1 |
| GPR143 | 0.18568095 | 1 | 0.182919 | 1 |
| MPP7 | 0.16221368 | 1 | 0.207842 | 1 |
| OR2G2 | 1.99970911 | 1 | -1.62898 | 1 |
| ZNF319 | 0.13819519 | 1 | 0.232735 | 1 |
| SLA | 0.19471083 | 1 | 0.176267 | 1 |
| TAS2R14 | 1.06592754 | 1 | -0.69491 | 1 |
| SLIRP | 0.36867685 | 1 | 0.003091 | 1 |
| RBM38 | 0.23563327 | 1 | 0.136346 | 1 |
| SOX15 | 0.08631366 | 1 | 0.285949 | 1 |
| NLRP1 | 0.04383089 | 1 | 0.328615 | 1 |
| NEURL2 | 0.0361601 | 1 | 0.336298 | 1 |
| CAMK1 | 0.24716338 | 1 | 0.126242 | 1 |
| PLCG1 | 0.22030372 | 1 | 0.153167 | 1 |
| CCN5 | 0.14767525 | 1 | 0.226722 | 1 |
| SOX7 | 0.02428108 | 1 | 0.350315 | 1 |
| IGF1 | 0.55663445 | 1 | -0.18173 | 1 |
| TAF12 | 0.86496867 | 1 | -0.49 | 1 |
| PSEN2 | 0.12643834 | 1 | 0.248839 | 1 |
| ZNF473 | 0.88158134 | 1 | -0.50618 | 1 |
| OR5H6 | 0.03806932 | 1 | 0.338118 | 1 |
| TFG | 1.01475403 | 1 | -0.63842 | 1 |
| GPER1 | 0.17375332 | 1 | 0.20291 | 1 |
| POU1F1 | 0.89402054 | 1 | -0.51722 | 1 |
| ZSCAN25 | 0.34559046 | 1 | 0.031706 | 1 |
| AGAP4 | 0.16218745 | 1 | 0.215235 | 1 |
| SBK2 | 0.15437152 | 1 | 0.223355 | 1 |
| PLEKHG6 | 0.12804059 | 1 | 0.250509 | 1 |
| ZSWIM3 | 0.12448583 | 1 | 0.254203 | 1 |
| RFX1 | 0.15264636 | 1 | 0.226244 | 1 |
| ZNF787 | 0.34684845 | 1 | 0.032228 | 1 |
| SCYL1 | 0.23161223 | 1 | 0.147518 | 1 |
| TLX2 | 0.09106615 | 1 | 0.288366 | 1 |
| ZFYVE26 | -0.525865 | 1 | 0.905683 | 1 |
| RASGRP4 | 0.07751677 | 1 | 0.302956 | 1 |

| | | | | |
|---|---|---|---|---|
| HOXA11 | 0.21576255 | 1 | 0.165022 | 1 |
| ADAP1 | 0.04830087 | 1 | 0.333268 | 1 |
| SMAD6 | 0.19825996 | 1 | 0.183469 | 1 |
| EHF | 0.34783357 | 1 | 0.034369 | 1 |
| MLXIPL | 0.16753999 | 1 | 0.215715 | 1 |
| OR5T1 | 0.00419158 | 1 | 0.379289 | 1 |
| ZFYVE28 | 0.07658824 | 1 | 0.306938 | 1 |
| SIPA1 | 0.09010878 | 1 | 0.294123 | 1 |
| ZNF208 | 0.05802824 | 1 | 0.326209 | 1 |
| GNAI2 | -0.0066235 | 1 | 0.390997 | 1 |
| EIF2AK2 | 0.22015284 | 1 | 0.164234 | 1 |
| PBX4 | 0.05935162 | 1 | 0.325094 | 1 |
| ITGAM | 0.10252316 | 1 | 0.282323 | 1 |
| AGAP3 | 0.15162098 | 1 | 0.233986 | 1 |
| MKX | -0.0511268 | 1 | 0.436956 | 1 |
| TFAP2C | -0.0543268 | 1 | 0.440958 | 1 |
| ASB8 | 0.2598602 | 1 | 0.126773 | 1 |
| ZNF3 | 0.49778303 | 1 | -0.11092 | 1 |
| ZNF449 | 0.52920242 | 1 | -0.14228 | 1 |
| ALX1 | -0.0214017 | 1 | 0.409031 | 1 |
| PLA2G1B | 0.34138526 | 1 | 0.04683 | 1 |
| CASP1 | 0.15741199 | 1 | 0.231008 | 1 |
| MED14 | 1.08710222 | 1 | -0.69702 | 1 |
| ASB18 | 0.257307 | 1 | 0.133397 | 1 |
| INPP5D | 0.12248101 | 1 | 0.26864 | 1 |
| CARD14 | 0.02043326 | 1 | 0.370933 | 1 |
| GTF2A1L | 0.2109221 | 1 | 0.180774 | 1 |
| CCND1 | 0.10767707 | 1 | 0.284138 | 1 |
| TYK2 | -0.0261486 | 1 | 0.418014 | 1 |
| ELOC | 0.61554405 | 1 | -0.22363 | 1 |
| CX3CR1 | 0.12542208 | 1 | 0.266956 | 1 |
| GAS7 | 0.14862869 | 1 | 0.24435 | 1 |
| PPP2R2C | -0.0290816 | 1 | 0.422257 | 1 |
| OR2A12 | 0.20268113 | 1 | 0.191687 | 1 |
| LCP1 | 0.29636133 | 1 | 0.099726 | 1 |
| TBX19 | 0.15174081 | 1 | 0.244568 | 1 |
| CIITA | 0.10203838 | 1 | 0.294744 | 1 |
| RIOX1 | -0.0278122 | 1 | 0.425044 | 1 |
| BAX | 0.17311193 | 1 | 0.2243 | 1 |
| PLXNB3 | -0.0335892 | 1 | 0.431131 | 1 |
| BARHL1 | 0.09248317 | 1 | 0.305202 | 1 |
| BTK | 0.20059981 | 1 | 0.197317 | 1 |
| ONECUT3 | 0.15438656 | 1 | 0.243823 | 1 |
| ERAS | 0.18413796 | 1 | 0.214588 | 1 |
| RING1 | 0.45606564 | 1 | -0.05703 | 1 |
| ZNF740 | 0.2790633 | 1 | 0.121771 | 1 |

| | | | | |
|---|---|---|---|---|
| ZNF385C | 0.19895717 | 1 | 0.202111 | 1 |
| ZNF76 | 0.12911403 | 1 | 0.272345 | 1 |
| NLRC3 | 0.10937199 | 1 | 0.292132 | 1 |
| DMD | 0.11826833 | 1 | 0.285111 | 1 |
| P2RX3 | 0.10804528 | 1 | 0.295972 | 1 |
| PDK2 | 0.20185719 | 1 | 0.203056 | 1 |
| CSF1R | 0.19645449 | 1 | 0.208644 | 1 |
| LRPPRC | 0.85648806 | 1 | -0.45113 | 1 |
| P2RX2 | -0.0250491 | 1 | 0.431072 | 1 |
| ARF1 | 0.56738208 | 1 | -0.16135 | 1 |
| TBC1D28 | 0.19941142 | 1 | 0.206923 | 1 |
| UNC5CL | 0.20138913 | 1 | 0.20501 | 1 |
| ZNF544 | 0.41192477 | 1 | -0.00503 | 1 |
| ZNF599 | 0.27126832 | 1 | 0.136084 | 1 |
| BAD | 0.3491926 | 1 | 0.058195 | 1 |
| CASZ1 | 0.74323441 | 1 | -0.33573 | 1 |
| PTCH2 | 0.05941078 | 1 | 0.348302 | 1 |
| IHH | 0.05340559 | 1 | 0.354395 | 1 |
| HTR2C | 0.56381384 | 1 | -0.15551 | 1 |
| AGPAT1 | 0.1421493 | 1 | 0.266525 | 1 |
| NPHP4 | 0.26383309 | 1 | 0.14502 | 1 |
| REM2 | 0.21433796 | 1 | 0.194756 | 1 |
| HLX | 0.05759843 | 1 | 0.351772 | 1 |
| HOXA2 | -0.9309945 | 1 | 1.340648 | 1 |
| ASCL1 | -0.0238244 | 1 | 0.433581 | 1 |
| GTF2A2 | 0.99235407 | 1 | -0.58242 | 1 |
| TLL2 | 0.04165564 | 1 | 0.369285 | 1 |
| SNCA | -0.1457228 | 1 | 0.556666 | 1 |
| MTERF1 | 0.59383615 | 1 | -0.18259 | 1 |
| TNFSF12 | 0.06927512 | 1 | 0.344486 | 1 |
| SIX5 | 0.08500179 | 1 | 0.329559 | 1 |
| ANGPTL6 | 0.05047072 | 1 | 0.366251 | 1 |
| FOXL2 | 0.24038138 | 1 | 0.17642 | 1 |
| E2F6 | 0.91228332 | 1 | -0.49537 | 1 |
| NEUROD1 | -0.1012727 | 1 | 0.51905 | 1 |
| LPXN | 0.15693786 | 1 | 0.260993 | 1 |
| ZNF419 | 0.17339853 | 1 | 0.24556 | 1 |
| TLX3 | 0.39581074 | 1 | 0.023259 | 1 |
| SH2D2A | -0.0584596 | 1 | 0.477685 | 1 |
| ZNF830 | 0.29420653 | 1 | 0.125797 | 1 |
| CD28 | -0.7082605 | 1 | 1.12886 | 1 |
| STAT4 | 0.33831052 | 1 | 0.082513 | 1 |
| AFF3 | -0.0786373 | 1 | 0.501015 | 1 |
| RREB1 | 0.1496158 | 1 | 0.272855 | 1 |
| RHOC | 0.22015407 | 1 | 0.202468 | 1 |
| ZNF707 | 0.3165526 | 1 | 0.106092 | 1 |

| | | | | |
|---|---|---|---|---|
| TRAF2 | 0.34566662 | 1 | 0.077427 | 1 |
| MAPK12 | 0.05941659 | 1 | 0.364299 | 1 |
| IGFBP3 | 0.400444 | 1 | 0.023494 | 1 |
| SETD1A | 0.14639839 | 1 | 0.2778 | 1 |
| EVL | 0.09458197 | 1 | 0.329709 | 1 |
| ADGRD2 | 0.15563914 | 1 | 0.270001 | 1 |
| ZNF880 | -0.5741858 | 1 | 1.000558 | 1 |
| PCGF1 | 0.26744219 | 1 | 0.158935 | 1 |
| ITGAE | 0.32073237 | 1 | 0.105851 | 1 |
| SETD1B | -0.1543244 | 1 | 0.582266 | 1 |
| DEPDC5 | 0.20699698 | 1 | 0.223041 | 1 |
| ARFRP1 | 0.07948685 | 1 | 0.350561 | 1 |
| MTERF2 | 0.22665488 | 1 | 0.203971 | 1 |
| PEBP1 | 0.24322541 | 1 | 0.18877 | 1 |
| CLIC3 | 0.22942977 | 1 | 0.202588 | 1 |
| CDC42EP5 | 0.23832944 | 1 | 0.193915 | 1 |
| HLA.DOA | 0.20323369 | 1 | 0.229693 | 1 |
| OR4C3 | 0.0429316 | 1 | 0.390009 | 1 |
| GCM2 | 0.11240106 | 1 | 0.320929 | 1 |
| CD3D | 0.19573788 | 1 | 0.23787 | 1 |
| ZNF561 | 0.71088055 | 1 | -0.27568 | 1 |
| NOSTRIN | 0.05841724 | 1 | 0.377512 | 1 |
| SLC11A1 | 0.15017955 | 1 | 0.286179 | 1 |
| SMARCAL1 | 0.31362417 | 1 | 0.12296 | 1 |
| ARHGAP10 | 0.01568809 | 1 | 0.423081 | 1 |
| ARHGAP23 | -0.8177125 | 1 | 1.257634 | 1 |
| GPR87 | 0.00322868 | 1 | 0.437414 | 1 |
| MUTYH | 0.2065669 | 1 | 0.234126 | 1 |
| ALDH1A2 | -0.0275741 | 1 | 0.468295 | 1 |
| AXIN1 | 0.07512399 | 1 | 0.366151 | 1 |
| GRM3 | -0.0374885 | 1 | 0.478847 | 1 |
| RNF14 | 0.60705336 | 1 | -0.16564 | 1 |
| IKBKE | 0.25019667 | 1 | 0.193127 | 1 |
| ARF4 | 0.86783798 | 1 | -0.42417 | 1 |
| VAX2 | 0.15775619 | 1 | 0.286131 | 1 |
| PRDM13 | 0.4146314 | 1 | 0.02933 | 1 |
| CDK9 | 0.13419683 | 1 | 0.310314 | 1 |
| AFP | 0.21378716 | 1 | 0.23076 | 1 |
| SOHLH1 | 0.18709418 | 1 | 0.258235 | 1 |
| ADORA2A | 0.19067579 | 1 | 0.255717 | 1 |
| ZNF534 | 0.02375705 | 1 | 0.423269 | 1 |
| ZCWPW1 | 0.21664433 | 1 | 0.230579 | 1 |
| OR2AE1 | 0.30995845 | 1 | 0.138051 | 1 |
| FADS1 | -1.0098599 | 1 | 1.459501 | 1 |
| GNA11 | 0.14730755 | 1 | 0.302688 | 1 |
| CABP4 | 0.07899897 | 1 | 0.372232 | 1 |

| | | | | |
|---|---|---|---|---|
| MC1R | 0.14895163 | 1 | 0.302748 | 1 |
| GFRAL | 0.3015731 | 1 | 0.151316 | 1 |
| ANXA9 | 0.1721612 | 1 | 0.28084 | 1 |
| ADORA1 | 0.10844144 | 1 | 0.344855 | 1 |
| STK25 | 0.24002414 | 1 | 0.21344 | 1 |
| ZSCAN20 | 0.32196453 | 1 | 0.132732 | 1 |
| IL10RB | 1.50185326 | 1 | -1.04548 | 1 |
| TDGF1 | 0.01383933 | 1 | 0.44263 | 1 |
| ERCC6 | -0.5968701 | 1 | 1.053343 | 1 |
| OR2K2 | 0.17844942 | 1 | 0.278279 | 1 |
| ZSCAN18 | 0.1673478 | 1 | 0.290741 | 1 |
| CERS5 | 0.40834009 | 1 | 0.05007 | 1 |
| LPAR5 | 0.18259618 | 1 | 0.275919 | 1 |
| CD2 | 0.14995771 | 1 | 0.309398 | 1 |
| TNFRSF25 | 0.13508216 | 1 | 0.324369 | 1 |
| GNB3 | 0.1286611 | 1 | 0.331398 | 1 |
| ASIC3 | 0.19535784 | 1 | 0.264835 | 1 |
| ITGB7 | 0.23192999 | 1 | 0.228472 | 1 |
| APBA3 | 0.21327226 | 1 | 0.247388 | 1 |
| CNOT6 | 0.81050807 | 1 | -0.34821 | 1 |
| HES2 | 0.1438579 | 1 | 0.319671 | 1 |
| WDTC1 | 0.08593218 | 1 | 0.377932 | 1 |
| SMARCC2 | -0.8127489 | 1 | 1.276615 | 1 |
| ESRRA | 0.2498874 | 1 | 0.214779 | 1 |
| SUPT6H | 0.38135654 | 1 | 0.084333 | 1 |
| PLPP3 | 0.17850379 | 1 | 0.287595 | 1 |
| GNG5 | 1.64882185 | 1 | -1.18233 | 1 |
| ADGRG5 | 0.11775634 | 1 | 0.349083 | 1 |
| ITGAX | 0.08415592 | 1 | 0.383118 | 1 |
| ZNF362 | 0.13215576 | 1 | 0.335172 | 1 |
| PLK2 | 1.34511627 | 1 | -0.87699 | 1 |
| LACRT | 0.90263989 | 1 | -0.43408 | 1 |
| ING4 | 0.34588259 | 1 | 0.123099 | 1 |
| NKX2.4 | -0.1365738 | 1 | 0.605846 | 1 |
| EEF1E1 | 0.67352993 | 1 | -0.20422 | 1 |
| C1QBP | 0.58769293 | 1 | -0.11796 | 1 |
| MED18 | 0.57119563 | 1 | -0.10112 | 1 |
| HOXA10 | 0.53879386 | 1 | -0.06745 | 1 |
| FSTL3 | 0.1818425 | 1 | 0.289785 | 1 |
| IKZF5 | 1.02492655 | 1 | -0.553 | 1 |
| KAT2B | 0.20562496 | 1 | 0.266627 | 1 |
| MLLT6 | 0.18514035 | 1 | 0.287498 | 1 |
| CNTNAP5 | 0.09491187 | 1 | 0.377862 | 1 |
| GFI1B | -0.0677241 | 1 | 0.543061 | 1 |
| EVI2A | 0.0668807 | 1 | 0.408704 | 1 |
| AKAP8 | 0.43685818 | 1 | 0.040951 | 1 |

| | | | | |
|---|---|---|---|---|
| TGIF2LY | -0.0539495 | 1 | 0.532009 | 1 |
| OR5K1 | 0.04601942 | 1 | 0.433698 | 1 |
| ZFPL1 | 0.26068719 | 1 | 0.219045 | 1 |
| AGAP5 | 0.15174098 | 1 | 0.328179 | 1 |
| IQGAP1 | 0.58455789 | 1 | -0.1045 | 1 |
| JMJD6 | 0.67910958 | 1 | -0.19868 | 1 |
| ADIPOQ | -0.0421075 | 1 | 0.523215 | 1 |
| ZNF232 | 0.28641834 | 1 | 0.194722 | 1 |
| RUNX3 | 0.37796682 | 1 | 0.103774 | 1 |
| OR4K15 | 0.49331673 | 1 | -0.01007 | 1 |
| PTPRB | 0.14029611 | 1 | 0.343149 | 1 |
| NFKB1 | 0.28195266 | 1 | 0.201886 | 1 |
| CCNL2 | 0.15374338 | 1 | 0.330496 | 1 |
| GIT1 | 0.15082854 | 1 | 0.335484 | 1 |
| OR6J1 | 0.32149156 | 1 | 0.16587 | 1 |
| MRAS | 0.10217974 | 1 | 0.385504 | 1 |
| ZNF600 | 0.04492632 | 1 | 0.444218 | 1 |
| MTNR1A | -0.4820055 | 1 | 0.971385 | 1 |
| ZNF30 | 0.17642578 | 1 | 0.313372 | 1 |
| COPS2 | 1.8095972 | 1 | -1.31814 | 1 |
| ZNF425 | 0.27287757 | 1 | 0.21866 | 1 |
| PPP2CB | 0.4774393 | 1 | 0.014189 | 1 |
| GPC3 | 0.43214207 | 1 | 0.059666 | 1 |
| IFI27 | 0.31782209 | 1 | 0.174308 | 1 |
| SIRT2 | -0.5429485 | 1 | 1.03536 | 1 |
| KLRG1 | -0.091941 | 1 | 0.584469 | 1 |
| CLDN4 | 0.17023461 | 1 | 0.32307 | 1 |
| TAL1 | 0.36129834 | 1 | 0.132549 | 1 |
| CARD9 | 0.15413533 | 1 | 0.339824 | 1 |
| ARHGAP4 | 0.22284461 | 1 | 0.272352 | 1 |
| GPSM3 | 0.18065786 | 1 | 0.315252 | 1 |
| SIX1 | 0.49751851 | 1 | -0.00144 | 1 |
| BAG1 | 0.46695226 | 1 | 0.029425 | 1 |
| SH2B2 | 0.51885082 | 1 | -0.02159 | 1 |
| OR10A6 | 0.01961805 | 1 | 0.478335 | 1 |
| OR10G8 | 0.15407225 | 1 | 0.344301 | 1 |
| PLCB2 | 0.15974945 | 1 | 0.338767 | 1 |
| GDF7 | 0.1104385 | 1 | 0.388303 | 1 |
| ADGRA2 | -0.0511975 | 1 | 0.549939 | 1 |
| EID3 | -0.0483462 | 1 | 0.547186 | 1 |
| ACKR2 | 0.21736152 | 1 | 0.281941 | 1 |
| CCNL1 | 0.20326509 | 1 | 0.296383 | 1 |
| IL7R | 0.3051938 | 1 | 0.194622 | 1 |
| P2RY11 | 0.21558585 | 1 | 0.284904 | 1 |
| H4C11 | 0.41542477 | 1 | 0.085367 | 1 |
| INSL4 | 0.78539617 | 1 | -0.28425 | 1 |

| | | | | |
|---|---|---|---|---|
| ZDHHC7 | 0.44737982 | 1 | 0.054442 | 1 |
| KLRF1 | 0.00300939 | 1 | 0.499964 | 1 |
| ARFIP2 | 0.33032918 | 1 | 0.172948 | 1 |
| HCRTR1 | 0.3953662 | 1 | 0.108792 | 1 |
| MLF1 | 0.23077513 | 1 | 0.274363 | 1 |
| TBC1D10C | 0.04117309 | 1 | 0.464794 | 1 |
| IRX4 | 0.17715437 | 1 | 0.329332 | 1 |
| OR52K2 | 0.23972479 | 1 | 0.267387 | 1 |
| ROPN1L | 0.06880803 | 1 | 0.438664 | 1 |
| THBS1 | 1.50616907 | 1 | -0.99833 | 1 |
| USF1 | 0.18303268 | 1 | 0.325567 | 1 |
| ARTN | 0.16512965 | 1 | 0.343885 | 1 |
| HTR1D | 0.45712451 | 1 | 0.052226 | 1 |
| CCL8 | 0.0885378 | 1 | 0.42097 | 1 |
| SECTM1 | -0.5701925 | 1 | 1.079946 | 1 |
| MYO10 | -0.8429032 | 1 | 1.352838 | 1 |
| SPOP | 1.08090253 | 1 | -0.57001 | 1 |
| LEPR | 1.08879052 | 1 | -0.57732 | 1 |
| MICAL1 | 0.1627713 | 1 | 0.34899 | 1 |
| NSMCE1 | 0.36311418 | 1 | 0.149137 | 1 |
| TFAP2E | 0.17244471 | 1 | 0.341298 | 1 |
| MYO18A | 0.05037103 | 1 | 0.463568 | 1 |
| HOXC6 | 0.15401322 | 1 | 0.360723 | 1 |
| TNIP2 | 0.20970066 | 1 | 0.306282 | 1 |
| HINFP | 0.22378433 | 1 | 0.292265 | 1 |
| ZBTB7B | 0.38084321 | 1 | 0.135262 | 1 |
| MTA1 | 0.4030055 | 1 | 0.113367 | 1 |
| NELFB | 0.10361685 | 1 | 0.413142 | 1 |
| ZFYVE27 | 0.28307751 | 1 | 0.234611 | 1 |
| CYP26C1 | 0.42626994 | 1 | 0.091866 | 1 |
| CHD1 | 1.90592841 | 1 | -1.38595 | 1 |
| CCL25 | 0.23547263 | 1 | 0.284832 | 1 |
| RGS9 | 0.22618823 | 1 | 0.294361 | 1 |
| LILRA3 | 0.29086667 | 1 | 0.230154 | 1 |
| MAGED1 | 0.2123456 | 1 | 0.308866 | 1 |
| DMAP1 | 0.31074104 | 1 | 0.210471 | 1 |
| UBE2V1 | 0.47408096 | 1 | 0.049198 | 1 |
| CD160 | 0.10551153 | 1 | 0.417996 | 1 |
| FOXC1 | -0.0634984 | 1 | 0.588092 | 1 |
| TCEAL5 | 0.0176175 | 1 | 0.508389 | 1 |
| PIK3CG | 0.1859356 | 1 | 0.340144 | 1 |
| LTBR | 0.19811155 | 1 | 0.328045 | 1 |
| VAC14 | 0.29978696 | 1 | 0.227871 | 1 |
| IL6R | 0.14638327 | 1 | 0.381603 | 1 |
| PRDM4 | 0.34819542 | 1 | 0.180044 | 1 |
| ZMIZ2 | -0.0833735 | 1 | 0.612163 | 1 |

| | | | | |
|---|---|---|---|---|
| GPR61 | 0.26157257 | 1 | 0.267847 | 1 |
| KEL | 0.14330999 | 1 | 0.386831 | 1 |
| PLCG2 | 0.05175481 | 1 | 0.478914 | 1 |
| PLXNB1 | 0.19625936 | 1 | 0.334612 | 1 |
| AGAP9 | 0.15324484 | 1 | 0.377974 | 1 |
| IL3RA | 0.20866159 | 1 | 0.323474 | 1 |
| ATRNL1 | 1.72567415 | 1 | -1.19328 | 1 |
| STAC3 | 0.1345223 | 1 | 0.397928 | 1 |
| GPR20 | 0.50639904 | 1 | 0.026057 | 1 |
| TRIM55 | -0.0026879 | 1 | 0.535302 | 1 |
| CAPN15 | 0.26665414 | 1 | 0.266667 | 1 |
| OR8U3 | 0.03653145 | 1 | 0.496821 | 1 |
| PPP2R5D | 0.97325223 | 1 | -0.43959 | 1 |
| GPR146 | 0.40258636 | 1 | 0.13115 | 1 |
| CREB3L4 | 0.13585324 | 1 | 0.398721 | 1 |
| LILRB1 | 0.38704422 | 1 | 0.147555 | 1 |
| AIFM2 | 0.34385754 | 1 | 0.19135 | 1 |
| SPA17 | 0.17197261 | 1 | 0.363517 | 1 |
| EZH1 | 0.10550566 | 1 | 0.430377 | 1 |
| NUTM2A | 0.05547808 | 1 | 0.480579 | 1 |
| LDB1 | 0.21745034 | 1 | 0.318701 | 1 |
| RBM10 | 0.30977266 | 1 | 0.227887 | 1 |
| ARHGAP27 | 0.1156131 | 1 | 0.422351 | 1 |
| SMPD1 | 0.29963167 | 1 | 0.238543 | 1 |
| DOK1 | 0.32245332 | 1 | 0.216442 | 1 |
| CLEC4M | 0.15388219 | 1 | 0.385819 | 1 |
| TAS2R20 | 0.00055693 | 1 | 0.539163 | 1 |
| SH3BP2 | 0.33567263 | 1 | 0.204793 | 1 |
| LAT2 | 0.24007133 | 1 | 0.300549 | 1 |
| RHOXF1 | 0.48907747 | 1 | 0.052005 | 1 |
| PRKAR2A | 1.24204345 | 1 | -0.7009 | 1 |
| RAB8A | 0.47692673 | 1 | 0.064257 | 1 |
| PNRC1 | 0.36619964 | 1 | 0.175176 | 1 |
| CD74 | 0.17066422 | 1 | 0.374439 | 1 |
| ARRB1 | -0.189626 | 1 | 0.736726 | 1 |
| ZNF696 | 0.31998488 | 1 | 0.227523 | 1 |
| GRHL3 | -0.1678895 | 1 | 0.71549 | 1 |
| ZNF547 | -0.0968048 | 1 | 0.644672 | 1 |
| RAX | 0.09399631 | 1 | 0.453967 | 1 |
| CBLC | 0.29462945 | 1 | 0.254037 | 1 |
| GPR153 | 0.42107478 | 1 | 0.128166 | 1 |
| RASD1 | 0.34723222 | 1 | 0.20202 | 1 |
| NKX3.1 | -0.487605 | 1 | 1.037438 | 1 |
| RNF10 | 0.46433838 | 1 | 0.087445 | 1 |
| GTF2H2 | 0.12563237 | 1 | 0.426168 | 1 |
| SGK1 | 0.0048558 | 1 | 0.547004 | 1 |

| | | | | |
|---|---|---|---|---|
| ASB1 | 0.13005004 | 1 | 0.421905 | 1 |
| ITGAL | 0.17609617 | 1 | 0.376035 | 1 |
| ZNF205 | 0.19404145 | 1 | 0.358469 | 1 |
| DVL1 | 0.4697014 | 1 | 0.082846 | 1 |
| AATF | 0.58930945 | 1 | -0.03676 | 1 |
| EP300 | 0.11492588 | 1 | 0.438205 | 1 |
| TIGD3 | 0.50397671 | 1 | 0.049642 | 1 |
| BAMBI | 0.20470404 | 1 | 0.348998 | 1 |
| GOLPH3 | 0.46026198 | 1 | 0.093683 | 1 |
| USP22 | -0.7715578 | 1 | 1.327269 | 1 |
| TGFBRAP1 | 0.49114399 | 1 | 0.064675 | 1 |
| HDAC11 | -0.8786601 | 1 | 1.434551 | 1 |
| SOX18 | 0.2995994 | 1 | 0.256405 | 1 |
| HIRA | 0.35047185 | 1 | 0.205822 | 1 |
| IL15RA | 0.30149965 | 1 | 0.255621 | 1 |
| DMRTA2 | 0.14139531 | 1 | 0.417811 | 1 |
| PRSS8 | 0.24011006 | 1 | 0.319899 | 1 |
| ASCC2 | 0.70995191 | 1 | -0.14796 | 1 |
| SDHB | 1.15706185 | 1 | -0.59467 | 1 |
| PARK7 | 0.72579923 | 1 | -0.16312 | 1 |
| GNGT2 | 0.14253654 | 1 | 0.421131 | 1 |
| NANOGP8 | 0.00345014 | 1 | 0.560244 | 1 |
| ZNF20 | 0.01830289 | 1 | 0.546422 | 1 |
| HSFX1 | 0.01911358 | 1 | 0.546053 | 1 |
| CHRM5 | -0.0210849 | 1 | 0.586287 | 1 |
| CXCR5 | 0.20823682 | 1 | 0.357768 | 1 |
| LILRB5 | 0.19527232 | 1 | 0.370922 | 1 |
| ZNF552 | 0.22732498 | 1 | 0.33895 | 1 |
| UNC13C | -0.0798391 | 1 | 0.646335 | 1 |
| OR12D3 | 0.00787005 | 1 | 0.559141 | 1 |
| BHLHA9 | -0.0441592 | 1 | 0.611308 | 1 |
| RASAL3 | 0.19825996 | 1 | 0.369025 | 1 |
| OR1S2 | 1.43620662 | 1 | -0.86792 | 1 |
| CALM1 | 0.51245945 | 1 | 0.057934 | 1 |
| HOXB4 | -0.017628 | 1 | 0.589095 | 1 |
| ZNF337 | 0.38785438 | 1 | 0.183621 | 1 |
| FIZ1 | 0.234307 | 1 | 0.33756 | 1 |
| ZNF296 | 0.2000392 | 1 | 0.373374 | 1 |
| TBL3 | 0.29304539 | 1 | 0.28101 | 1 |
| OR4K14 | 0.38250129 | 1 | 0.19242 | 1 |
| PREB | 0.50690609 | 1 | 0.068806 | 1 |
| RARG | 0.27720937 | 1 | 0.299128 | 1 |
| CAPN5 | 0.44704136 | 1 | 0.129521 | 1 |
| GPBAR1 | 0.28408163 | 1 | 0.294532 | 1 |
| HOXB6 | 0.75748231 | 1 | -0.1787 | 1 |
| CD81 | 0.31661248 | 1 | 0.26286 | 1 |

| | | | | |
|---|---|---|---|---|
| GPR39 | 0.05612577 | 1 | 0.523473 | 1 |
| PRKACG | 0.02588825 | 1 | 0.554133 | 1 |
| TNF | 0.21415818 | 1 | 0.368075 | 1 |
| MYCN | 0.14092082 | 1 | 0.441477 | 1 |
| S100A1 | 0.07595614 | 1 | 0.508153 | 1 |
| LPAR6 | 0.61878197 | 1 | -0.03334 | 1 |
| RAPGEFL1 | 0.1567394 | 1 | 0.429857 | 1 |
| POLR2L | 0.3914266 | 1 | 0.1955 | 1 |
| PLEKHG1 | 0.52250067 | 1 | 0.066333 | 1 |
| SHC2 | 0.06002126 | 1 | 0.529145 | 1 |
| MADCAM1 | 0.2109963 | 1 | 0.378382 | 1 |
| FCGRT | 0.31688187 | 1 | 0.273214 | 1 |
| RAD1 | 1.09019091 | 1 | -0.49984 | 1 |
| OR4F6 | 0.37122363 | 1 | 0.219711 | 1 |
| RAP1A | 0.80399461 | 1 | -0.21257 | 1 |
| SPTA1 | -0.0240092 | 1 | 0.615527 | 1 |
| ERF | 0.0498568 | 1 | 0.541912 | 1 |
| KEAP1 | 0.46958757 | 1 | 0.122265 | 1 |
| RHOG | 0.19216582 | 1 | 0.400145 | 1 |
| CYP26B1 | -0.212465 | 1 | 0.805244 | 1 |
| ZBED3 | 0.60594552 | 1 | -0.01175 | 1 |
| HOXD8 | 0.0079469 | 1 | 0.586403 | 1 |
| NUP62 | 0.21806478 | 1 | 0.376793 | 1 |
| ZNF324 | 0.14630751 | 1 | 0.448896 | 1 |
| RAB40C | 0.2152204 | 1 | 0.380245 | 1 |
| PHF21A | 0.85389462 | 1 | -0.25751 | 1 |
| TMED1 | 0.41092953 | 1 | 0.185836 | 1 |
| RGS19 | 0.33316074 | 1 | 0.263671 | 1 |
| CC2D1A | 0.26967911 | 1 | 0.327239 | 1 |
| TBX2 | 0.06799559 | 1 | 0.531093 | 1 |
| GABRD | -1.0840501 | 1 | 1.683296 | 1 |
| MADD | 0.17637259 | 1 | 0.423012 | 1 |
| BABAM1 | 0.41912494 | 1 | 0.180474 | 1 |
| RAPH1 | 0.59261115 | 1 | 0.00719 | 1 |
| NCR1 | 0.40339573 | 1 | 0.196734 | 1 |
| DAP | 0.18478223 | 1 | 0.415752 | 1 |
| GPR82 | 0.30083783 | 1 | 0.300291 | 1 |
| SLAMF1 | 0.02365423 | 1 | 0.577663 | 1 |
| FOXI3 | 0.18217501 | 1 | 0.419569 | 1 |
| OR5B17 | 0.54364349 | 1 | 0.059131 | 1 |
| ZC3H7A | 0.96252928 | 1 | -0.35871 | 1 |
| DVL2 | 0.24934015 | 1 | 0.356054 | 1 |
| PPP1R1C | 0.10570708 | 1 | 0.499901 | 1 |
| SLC2A4RG | 0.24861971 | 1 | 0.357091 | 1 |
| SNAI2 | -0.0386958 | 1 | 0.644688 | 1 |
| MED11 | 0.31134757 | 1 | 0.294709 | 1 |

| | | | | |
|---|---|---|---|---|
| HOXD1 | -0.0139974 | 1 | 0.622151 | 1 |
| ZNF837 | 0.28732157 | 1 | 0.321015 | 1 |
| ZNF646 | 0.2893383 | 1 | 0.3196 | 1 |
| GRIN2C | 0.26376689 | 1 | 0.345191 | 1 |
| PAX7 | -0.0394006 | 1 | 0.649092 | 1 |
| GABRA3 | -0.2895813 | 1 | 0.899581 | 1 |
| HES5 | -1.139727 | 1 | 1.750416 | 1 |
| ZNF500 | 0.27639114 | 1 | 0.334899 | 1 |
| TAB1 | 0.14896758 | 1 | 0.462483 | 1 |
| PDE1C | -0.065315 | 1 | 0.676824 | 1 |
| FRAT2 | 0.21198881 | 1 | 0.400645 | 1 |
| ZNF517 | 0.26527801 | 1 | 0.347704 | 1 |
| EID2B | 0.26692655 | 1 | 0.34615 | 1 |
| LIN28A | 0.28332425 | 1 | 0.329854 | 1 |
| ZNF596 | 0.63473899 | 1 | -0.02049 | 1 |
| PLCB4 | 1.15247208 | 1 | -0.53711 | 1 |
| RYK | 1.15094833 | 1 | -0.53501 | 1 |
| MINK1 | 0.15804893 | 1 | 0.458015 | 1 |
| TLR7 | 0.26339985 | 1 | 0.353295 | 1 |
| TRIB3 | 0.86773554 | 1 | -0.25076 | 1 |
| DDIT4 | 0.24263874 | 1 | 0.374498 | 1 |
| ZNF320 | 0.07913932 | 1 | 0.538545 | 1 |
| SKAP1 | 0.5035264 | 1 | 0.114176 | 1 |
| SPI1 | 0.1802538 | 1 | 0.439298 | 1 |
| WNT4 | -0.2688879 | 1 | 0.889204 | 1 |
| CCDC85B | 0.37539871 | 1 | 0.244932 | 1 |
| CD300C | 0.25421885 | 1 | 0.367016 | 1 |
| RCAN1 | 0.65273399 | 1 | -0.03077 | 1 |
| RAB7A | 0.73974675 | 1 | -0.11565 | 1 |
| PCBP1 | 1.08223871 | 1 | -0.45793 | 1 |
| KMT5A | 0.87549536 | 1 | -0.24877 | 1 |
| EDNRB | -0.0405111 | 1 | 0.667473 | 1 |
| OR6X1 | 0.262287 | 1 | 0.365761 | 1 |
| TWIST2 | 0.09239685 | 1 | 0.536717 | 1 |
| ZNF284 | 0.42465825 | 1 | 0.204633 | 1 |
| PTGIR | 0.04584431 | 1 | 0.584793 | 1 |
| CUX2 | -0.4009224 | 1 | 1.032787 | 1 |
| NCOA5 | 0.42774376 | 1 | 0.204883 | 1 |
| APBB1 | -0.1940214 | 1 | 0.828573 | 1 |
| UNC13A | -1.416851 | 1 | 2.051448 | 1 |
| OR4E1 | 0.03905579 | 1 | 0.595979 | 1 |
| IFT172 | 0.24531734 | 1 | 0.390068 | 1 |
| TBC1D3B | 0.25179771 | 1 | 0.384266 | 1 |
| ZNF155 | 0.48218436 | 1 | 0.15421 | 1 |
| CMKLR1 | 0.32035636 | 1 | 0.317402 | 1 |
| CXCR3 | 0.22774891 | 1 | 0.410344 | 1 |

| | | | | |
|---|---|---|---|---|
| CDK11B | 0.27522949 | 1 | 0.363549 | 1 |
| TYROBP | 0.35070355 | 1 | 0.289656 | 1 |
| RGN | 0.29773864 | 1 | 0.342839 | 1 |
| PWP2 | 0.6821358 | 1 | -0.04089 | 1 |
| ATXN3 | 0.22086481 | 1 | 0.422332 | 1 |
| FOXE1 | -0.0321244 | 1 | 0.675413 | 1 |
| MED9 | 0.41811948 | 1 | 0.225191 | 1 |
| CLN3 | 0.31682797 | 1 | 0.327062 | 1 |
| IL1RL1 | 0.60091694 | 1 | 0.043565 | 1 |
| CEBPA | 0.31820155 | 1 | 0.32942 | 1 |
| HSFX2 | -0.0072526 | 1 | 0.655221 | 1 |
| TCL1A | 0.02145649 | 1 | 0.626774 | 1 |
| PTPN11 | 1.00792453 | 1 | -0.35836 | 1 |
| RBCK1 | 0.34635992 | 1 | 0.304424 | 1 |
| NOC2L | 0.80145956 | 1 | -0.14866 | 1 |
| POFUT1 | 1.04144574 | 1 | -0.38754 | 1 |
| SFPQ | 1.7532228 | 1 | -1.09767 | 1 |
| SHC1 | 0.73587534 | 1 | -0.08014 | 1 |
| FFAR4 | 0.17779524 | 1 | 0.478129 | 1 |
| IL10RA | 0.31624082 | 1 | 0.339735 | 1 |
| BANP | 0.18458879 | 1 | 0.471464 | 1 |
| NOTCH4 | 0.29359504 | 1 | 0.364495 | 1 |
| SOX5 | 0.71557052 | 1 | -0.05666 | 1 |
| PRMT6 | 1.18350307 | 1 | -0.52456 | 1 |
| SMARCA2 | -0.4853909 | 1 | 1.144763 | 1 |
| PILRB | 0.12566464 | 1 | 0.53402 | 1 |
| CASP8 | 0.15909342 | 1 | 0.501144 | 1 |
| TBC1D3F | -0.0009573 | 1 | 0.661796 | 1 |
| RGS10 | 0.27419246 | 1 | 0.386726 | 1 |
| CSNK1G2 | 0.43523429 | 1 | 0.225804 | 1 |
| IKZF1 | 0.44414497 | 1 | 0.21757 | 1 |
| ATF6B | 0.39221707 | 1 | 0.270443 | 1 |
| NFIB | -0.129638 | 1 | 0.792442 | 1 |
| S1PR4 | 0.5552859 | 1 | 0.10767 | 1 |
| DGKG | 0.35834361 | 1 | 0.305051 | 1 |
| IFT27 | 0.16200917 | 1 | 0.501444 | 1 |
| GPSM1 | 0.26800591 | 1 | 0.395624 | 1 |
| KRTAP21.1 | 0.58126221 | 1 | 0.08249 | 1 |
| MS4A8 | 0.70576501 | 1 | -0.04166 | 1 |
| OR52B4 | 0.91443358 | 1 | -0.2492 | 1 |
| GPR4 | 0.48201234 | 1 | 0.183416 | 1 |
| MTA2 | 0.70104369 | 1 | -0.03494 | 1 |
| GATAD1 | 0.52137819 | 1 | 0.145309 | 1 |
| TSG101 | 0.83982986 | 1 | -0.17263 | 1 |
| PTK7 | 0.58993865 | 1 | 0.079568 | 1 |
| SOX14 | 0.1473607 | 1 | 0.522518 | 1 |

| | | | | |
|---|---|---|---|---|
| STMN1 | 3.35864886 | 1 | -2.68724 | 1 |
| CD53 | 0.28003302 | 1 | 0.391448 | 1 |
| KLHL12 | 0.78969864 | 1 | -0.11813 | 1 |
| AGER | 0.12426905 | 1 | 0.550508 | 1 |
| GDI1 | 0.97006421 | 1 | -0.29521 | 1 |
| ELF5 | 0.17792965 | 1 | 0.498657 | 1 |
| MRGPRX3 | 0.0607085 | 1 | 0.616214 | 1 |
| GALR2 | 0.2816271 | 1 | 0.395371 | 1 |
| FAM120B | 0.14123758 | 1 | 0.536283 | 1 |
| ARHGEF1 | 0.22989033 | 1 | 0.448298 | 1 |
| SP2 | 0.39084302 | 1 | 0.287695 | 1 |
| PEX14 | 0.68367721 | 1 | -0.00496 | 1 |
| ADRA2C | 0.79184794 | 1 | -0.11213 | 1 |
| MECP2 | 0.02896847 | 1 | 0.651724 | 1 |
| MSTN | 0.09436578 | 1 | 0.587168 | 1 |
| CD8A | -0.1575391 | 1 | 0.83913 | 1 |
| ZNF491 | 0.20802961 | 1 | 0.474129 | 1 |
| MAP2K7 | 0.25731684 | 1 | 0.426031 | 1 |
| CCL18 | 0.19867968 | 1 | 0.485764 | 1 |
| OR6N1 | -0.031497 | 1 | 0.717296 | 1 |
| DNLZ | 0.76730807 | 1 | -0.08151 | 1 |
| ING5 | 0.33242574 | 1 | 0.353419 | 1 |
| GPR89A | 0.4816494 | 1 | 0.205207 | 1 |
| OR5K2 | 0.72743977 | 1 | -0.04022 | 1 |
| ADGRA3 | -1.5730207 | 1 | 2.26109 | 1 |
| OR2V1 | 0.46185216 | 1 | 0.226286 | 1 |
| SPG21 | 1.8714818 | 1 | -1.183 | 1 |
| THAP1 | 0.86897719 | 1 | -0.17998 | 1 |
| ZNF354A | 1.57876145 | 1 | -0.88764 | 1 |
| FOXN1 | 0.63196278 | 1 | 0.059384 | 1 |
| ADCY9 | -0.4038851 | 1 | 1.095439 | 1 |
| VDR | 0.82552208 | 1 | -0.13393 | 1 |
| RFX2 | 0.1463866 | 1 | 0.545325 | 1 |
| LMO1 | -0.6024618 | 1 | 1.294606 | 1 |
| FOXP3 | -0.1278526 | 1 | 0.820931 | 1 |
| MAPK3 | 0.28391848 | 1 | 0.410878 | 1 |
| ELP5 | 0.43942575 | 1 | 0.255645 | 1 |
| LTB4R | 0.26480711 | 1 | 0.430374 | 1 |
| OR51A2 | 0.70134141 | 1 | -0.00605 | 1 |
| IL1B | -0.1020158 | 1 | 0.798036 | 1 |
| LRCH4 | 0.07103976 | 1 | 0.625058 | 1 |
| ZNF496 | 0.25969781 | 1 | 0.436842 | 1 |
| CGB5 | 0.17667248 | 1 | 0.52087 | 1 |
| SNAI3 | -0.1216401 | 1 | 0.82011 | 1 |
| ABLIM3 | -0.8307303 | 1 | 1.53128 | 1 |
| SRC | 0.23749184 | 1 | 0.46363 | 1 |

| | | | | |
|---|---|---|---|---|
| CTNND2 | -2.4513497 | 1 | 3.152559 | 1 |
| SYNGAP1 | 0.24388274 | 1 | 0.457505 | 1 |
| ADRA1D | 0.20700396 | 1 | 0.49456 | 1 |
| PSMC5 | 0.4951044 | 1 | 0.206589 | 1 |
| NOS3 | 0.32264914 | 1 | 0.379821 | 1 |
| TRIM25 | 0.26994154 | 1 | 0.434015 | 1 |
| ZIC3 | 0.49080835 | 1 | 0.214892 | 1 |
| PHOX2B | 0.68602788 | 1 | 0.02121 | 1 |
| EDA2R | 0.37544145 | 1 | 0.332089 | 1 |
| CSF3R | 0.2944509 | 1 | 0.413717 | 1 |
| PIP4K2B | 0.02903942 | 1 | 0.67969 | 1 |
| PAQR8 | -1.7563482 | 1 | 2.465836 | 1 |
| ZNF575 | 0.56042299 | 1 | 0.149103 | 1 |
| GIGYF1 | 0.18468886 | 1 | 0.52556 | 1 |
| OR1F1 | -0.9196371 | 1 | 1.63148 | 1 |
| OR1L4 | -0.0059847 | 1 | 0.718488 | 1 |
| FBXO11 | 0.9939036 | 1 | -0.28043 | 1 |
| ARL2 | 0.53172106 | 1 | 0.182446 | 1 |
| GIT2 | 0.75734077 | 1 | -0.04223 | 1 |
| TGIF2LX | 0.12304307 | 1 | 0.593435 | 1 |
| AGAP6 | -0.0170458 | 1 | 0.734516 | 1 |
| VAPB | 0.55446407 | 1 | 0.163283 | 1 |
| HIVEP1 | 1.24712644 | 1 | -0.52826 | 1 |
| RAB26 | 0.38226184 | 1 | 0.339877 | 1 |
| SNAPC5 | 0.48359788 | 1 | 0.239017 | 1 |
| DAP3 | 0.86626319 | 1 | -0.14346 | 1 |
| EID1 | 0.7138589 | 1 | 0.008958 | 1 |
| USP6 | 0.28409167 | 1 | 0.43954 | 1 |
| TBC1D1 | 0.6441123 | 1 | 0.081012 | 1 |
| KMT2B | 0.39631483 | 1 | 0.330669 | 1 |
| TCEAL8 | 0.88645094 | 1 | -0.15924 | 1 |
| OR2T2 | -0.0033488 | 1 | 0.731113 | 1 |
| ZGLP1 | 0.17100132 | 1 | 0.557742 | 1 |
| IL11 | 0.45590811 | 1 | 0.272864 | 1 |
| DTX2 | 0.2515933 | 1 | 0.47761 | 1 |
| ITSN1 | 0.26092536 | 1 | 0.468662 | 1 |
| SNAPC3 | 1.11781954 | 1 | -0.38726 | 1 |
| MBD3 | 0.32690908 | 1 | 0.403947 | 1 |
| ZNF493 | -0.33704 | 1 | 1.068597 | 1 |
| NPSR1 | 0.62853938 | 1 | 0.103214 | 1 |
| DYRK1B | 0.28142015 | 1 | 0.451709 | 1 |
| SUPT4H1 | 0.64088085 | 1 | 0.092354 | 1 |
| GSN | 0.13165706 | 1 | 0.602426 | 1 |
| ARFGAP1 | 0.58202315 | 1 | 0.153045 | 1 |
| RGL3 | -0.0455422 | 1 | 0.782056 | 1 |
| PRMT7 | 0.60080698 | 1 | 0.13613 | 1 |

| | | | | |
|---|---|---|---|---|
| MGST2 | 0.34453682 | 1 | 0.39249 | 1 |
| RASIP1 | 0.12659697 | 1 | 0.611896 | 1 |
| SCTR | 0.22692761 | 1 | 0.512693 | 1 |
| EVX2 | 0.24458785 | 1 | 0.495589 | 1 |
| GRIA1 | 0.0680548 | 1 | 0.672952 | 1 |
| CALML5 | 0.01223452 | 1 | 0.72916 | 1 |
| THOP1 | 0.49237444 | 1 | 0.249213 | 1 |
| ZGPAT | 0.34134507 | 1 | 0.400583 | 1 |
| PER3 | 0.26247925 | 1 | 0.480702 | 1 |
| PIAS1 | 0.99292507 | 1 | -0.24969 | 1 |
| KDM4B | -0.6470229 | 1 | 1.390631 | 1 |
| OTUD7B | 0.64605253 | 1 | 0.097829 | 1 |
| EDA | 0.13221415 | 1 | 0.611724 | 1 |
| TRIP10 | 0.63505923 | 1 | 0.110202 | 1 |
| SMARCD2 | 0.46466238 | 1 | 0.280895 | 1 |
| VSX1 | -0.0750106 | 1 | 0.820983 | 1 |
| NPAS2 | 0.88431942 | 1 | -0.13796 | 1 |
| RAB5B | 0.04107061 | 1 | 0.705399 | 1 |
| MED29 | 0.31494739 | 1 | 0.43221 | 1 |
| GUCA1B | 0.26404547 | 1 | 0.483831 | 1 |
| TRIM23 | 1.07831364 | 1 | -0.32948 | 1 |
| SYDE1 | 0.05914166 | 1 | 0.689796 | 1 |
| CDX1 | 0.92104404 | 1 | -0.17174 | 1 |
| GPR37L1 | 0.57957974 | 1 | 0.169967 | 1 |
| DRD3 | 0.0331122 | 1 | 0.716666 | 1 |
| OR5D18 | 0.85535075 | 1 | -0.10247 | 1 |
| OR2B11 | 0.0947616 | 1 | 0.658324 | 1 |
| HOXC5 | 0.01346541 | 1 | 0.740079 | 1 |
| LGALS3BP | 0.59175772 | 1 | 0.161898 | 1 |
| CCDC88C | -0.291303 | 1 | 1.045135 | 1 |
| PFDN1 | 0.63982328 | 1 | 0.114512 | 1 |
| ZNF853 | 0.68269463 | 1 | 0.072339 | 1 |
| RHOT2 | 0.4165393 | 1 | 0.338534 | 1 |
| TCEA3 | 0.68495141 | 1 | 0.071006 | 1 |
| CISH | 2.03634371 | 1 | -1.27857 | 1 |
| FBXO6 | -0.0681215 | 1 | 0.826057 | 1 |
| TNFSF14 | 0.38801611 | 1 | 0.372633 | 1 |
| RHOD | 0.06398864 | 1 | 0.69769 | 1 |
| L3MBTL2 | 0.42403469 | 1 | 0.337728 | 1 |
| ZBTB32 | 0.18874414 | 1 | 0.575343 | 1 |
| ZNF628 | 0.59190851 | 1 | 0.172661 | 1 |
| MAP3K7 | 1.32455598 | 1 | -0.55907 | 1 |
| RASSF7 | 0.55905034 | 1 | 0.206613 | 1 |
| STMN3 | -0.0377552 | 1 | 0.803452 | 1 |
| ZXDC | 0.3396484 | 1 | 0.427389 | 1 |
| GDF2 | 0.39580618 | 1 | 0.372121 | 1 |

| | | | | |
|---|---|---|---|---|
| SNAPC4 | 0.31833007 | 1 | 0.449846 | 1 |
| ZNF572 | 0.26531051 | 1 | 0.50355 | 1 |
| OR6C74 | 0.07369291 | 1 | 0.697254 | 1 |
| SP6 | 0.65876106 | 1 | 0.113536 | 1 |
| MYH11 | 0.70774336 | 1 | 0.064584 | 1 |
| RAD9A | 0.20830897 | 1 | 0.564161 | 1 |
| NR1H3 | 0.56797303 | 1 | 0.205043 | 1 |
| ARHGEF25 | 0.03445693 | 1 | 0.738695 | 1 |
| MRGBP | 0.86394735 | 1 | -0.09043 | 1 |
| LHX5 | 0.26508583 | 1 | 0.510688 | 1 |
| SRSF3 | 1.44899009 | 1 | -0.67306 | 1 |
| IL16 | 0.02349867 | 1 | 0.752618 | 1 |
| SMARCD3 | 0.26984874 | 1 | 0.506478 | 1 |
| FST | 0.00300244 | 1 | 0.7734 | 1 |
| PIAS2 | 0.86189873 | 1 | -0.08527 | 1 |
| OR8K3 | 0.72463525 | 1 | 0.052545 | 1 |
| DTX4 | -0.6227374 | 1 | 1.401115 | 1 |
| POU2AF1 | 0.2561391 | 1 | 0.522876 | 1 |
| OR3A2 | 0.2156838 | 1 | 0.563479 | 1 |
| BRD8 | 0.78048276 | 1 | -0.00034 | 1 |
| TSC22D4 | 0.41410418 | 1 | 0.367625 | 1 |
| BABAM2 | 0.71656074 | 1 | 0.065585 | 1 |
| ARHGAP39 | 0.33393176 | 1 | 0.448405 | 1 |
| HOXB5 | 0.53159034 | 1 | 0.251298 | 1 |
| ZNF688 | 0.31407134 | 1 | 0.469092 | 1 |
| NEUROG3 | -0.0395374 | 1 | 0.824193 | 1 |
| GNAT2 | 0.09334251 | 1 | 0.691535 | 1 |
| CAV1 | 0.79921355 | 1 | -0.01413 | 1 |
| CD80 | 0.00558458 | 1 | 0.780028 | 1 |
| ZMAT1 | 0.08603499 | 1 | 0.699594 | 1 |
| OPRK1 | -0.3421899 | 1 | 1.128239 | 1 |
| SNX6 | 1.39506557 | 1 | -0.60862 | 1 |
| MOV10 | 0.17715805 | 1 | 0.610123 | 1 |
| TLE1 | 0.00708884 | 1 | 0.782113 | 1 |
| OR1D5 | 0.21022645 | 1 | 0.580249 | 1 |
| RAB40AL | 0.03086231 | 1 | 0.762268 | 1 |
| MAP3K19 | 0.13670849 | 1 | 0.65799 | 1 |
| SLC22A17 | 0.3963114 | 1 | 0.398878 | 1 |
| GPR25 | 0.67543842 | 1 | 0.119817 | 1 |
| MIER2 | 0.2300091 | 1 | 0.565297 | 1 |
| KAT2A | 0.28405975 | 1 | 0.511445 | 1 |
| LRP1B | 0.82826808 | 1 | -0.03178 | 1 |
| PPP2R5C | 0.94522542 | 1 | -0.14675 | 1 |
| GLIS1 | -0.003021 | 1 | 0.801835 | 1 |
| ZNF174 | 0.63404147 | 1 | 0.164865 | 1 |
| CAMLG | 0.84298074 | 1 | -0.04289 | 1 |

| | | | | |
|---|---|---|---|---|
| FOXD4L6 | 0.59875624 | 1 | 0.202057 | 1 |
| SORT1 | -0.2554725 | 1 | 1.05633 | 1 |
| ZNF875 | 0.11763285 | 1 | 0.683897 | 1 |
| TBX10 | 0.54118765 | 1 | 0.261846 | 1 |
| ARHGEF26 | -0.0114585 | 1 | 0.814543 | 1 |
| OR6M1 | 0.03379154 | 1 | 0.77056 | 1 |
| GLRA1 | -0.0642824 | 1 | 0.86877 | 1 |
| MAP2K5 | 0.17499989 | 1 | 0.630162 | 1 |
| SP140 | 0.14532921 | 1 | 0.660331 | 1 |
| MAPRE3 | 0.71679219 | 1 | 0.089858 | 1 |
| SRY | 1.35987279 | 1 | -0.55218 | 1 |
| OR1M1 | 0.37815142 | 1 | 0.430706 | 1 |
| GPR65 | 0.25827099 | 1 | 0.551101 | 1 |
| LHB | 0.2337268 | 1 | 0.575776 | 1 |
| NSMAF | 0.89974066 | 1 | -0.09007 | 1 |
| TCF20 | 0.89711446 | 1 | -0.0868 | 1 |
| OR2T10 | 2.00524951 | 1 | -1.19413 | 1 |
| CARD11 | 0.23136889 | 1 | 0.580167 | 1 |
| ATP6AP2 | 1.18818209 | 1 | -0.37526 | 1 |
| POU4F3 | 0.38162564 | 1 | 0.433429 | 1 |
| CD19 | -0.0054519 | 1 | 0.820683 | 1 |
| TCF23 | 0.03933074 | 1 | 0.777271 | 1 |
| ETV2 | 0.77489655 | 1 | 0.042514 | 1 |
| PIK3CD | 0.04620064 | 1 | 0.7727 | 1 |
| LHX6 | 0.43841857 | 1 | 0.380684 | 1 |
| FOXP4 | 0.5349826 | 1 | 0.284202 | 1 |
| ARHGDIA | 1.19343804 | 1 | -0.37296 | 1 |
| ZNHIT1 | 0.62607299 | 1 | 0.194428 | 1 |
| BORCS8.MEF2B | 0.20538392 | 1 | 0.615707 | 1 |
| P2RX6 | 0.09291988 | 1 | 0.728911 | 1 |
| AIP | 0.67959796 | 1 | 0.142557 | 1 |
| SFSWAP | 0.31880053 | 1 | 0.503822 | 1 |
| SRSF2 | 0.74071619 | 1 | 0.083372 | 1 |
| FOXD4L1 | 0.60477414 | 1 | 0.222019 | 1 |
| RIN1 | 0.98432911 | 1 | -0.15748 | 1 |
| NRIP2 | 0.0524396 | 1 | 0.775108 | 1 |
| PDC | 0.04665003 | 1 | 0.780956 | 1 |
| GNG11 | 0.22313532 | 1 | 0.604929 | 1 |
| ECSIT | 0.09404946 | 1 | 0.735132 | 1 |
| LRRFIP2 | 0.52060341 | 1 | 0.308705 | 1 |
| SGSM3 | 0.19434007 | 1 | 0.636691 | 1 |
| APEX1 | 1.28648216 | 1 | -0.45505 | 1 |
| KNG1 | -0.017873 | 1 | 0.84931 | 1 |
| GCGR | 0.87932876 | 1 | -0.04721 | 1 |
| ZNF213 | 0.26528325 | 1 | 0.56692 | 1 |
| GADD45G | 0.67566378 | 1 | 0.156904 | 1 |

| | | | | |
|---|---|---|---|---|
| ULK1 | 0.30516497 | 1 | 0.528053 | 1 |
| OR14A2 | 0.00431818 | 1 | 0.829045 | 1 |
| RFX7 | 0.09667094 | 1 | 0.739833 | 1 |
| PRDM1 | 0.10101358 | 1 | 0.735634 | 1 |
| TRIM22 | 0.14052396 | 1 | 0.696864 | 1 |
| ZNF706 | 0.92462584 | 1 | -0.08629 | 1 |
| STAT3 | 0.00083031 | 1 | 0.837853 | 1 |
| CXCL14 | 0.57442808 | 1 | 0.264269 | 1 |
| ZC3H18 | 0.35947503 | 1 | 0.482271 | 1 |
| MED25 | 0.17868278 | 1 | 0.663881 | 1 |
| OR14C36 | 0.84258589 | 1 | 0.002685 | 1 |
| STK11 | 0.54428472 | 1 | 0.301661 | 1 |
| FOXH1 | 0.35248504 | 1 | 0.493488 | 1 |
| PRMT5 | 0.74494055 | 1 | 0.101976 | 1 |
| PDCD1LG2 | 0.48453865 | 1 | 0.362554 | 1 |
| ACAD8 | 0.35657702 | 1 | 0.491832 | 1 |
| SETD7 | 0.88243572 | 1 | -0.03396 | 1 |
| ZNF653 | 0.45008412 | 1 | 0.398703 | 1 |
| MAPK10 | 0.63638474 | 1 | 0.212712 | 1 |
| GRIK4 | -0.8759404 | 1 | 1.725976 | 1 |
| SKI | 0.15693789 | 1 | 0.693666 | 1 |
| CAV3 | 0.00702347 | 1 | 0.84508 | 1 |
| ZBTB9 | 1.06038326 | 1 | -0.20633 | 1 |
| RAP1GAP | 0.13791985 | 1 | 0.716273 | 1 |
| OR2A25 | 0.05576494 | 1 | 0.799248 | 1 |
| ARHGAP15 | 0.27407888 | 1 | 0.582602 | 1 |
| ARHGEF10L | 0.07491853 | 1 | 0.781809 | 1 |
| XRN2 | 1.29837688 | 1 | -0.44096 | 1 |
| ZNF253 | 1.27063507 | 1 | -0.41181 | 1 |
| PIN1 | 0.40560888 | 1 | 0.453552 | 1 |
| CDA | 0.2196986 | 1 | 0.640546 | 1 |
| OR1L8 | 0.43152035 | 1 | 0.429013 | 1 |
| OR4C5 | 0.89524388 | 1 | -0.03424 | 1 |
| TCF7 | 0.09263151 | 1 | 0.768745 | 1 |
| PRDM2 | 0.73853748 | 1 | 0.123729 | 1 |
| TBC1D17 | 0.33718779 | 1 | 0.527131 | 1 |
| NEUROD4 | 1.30608325 | 1 | -0.43959 | 1 |
| RAMP3 | 0.58944659 | 1 | 0.278046 | 1 |
| ZNF700 | -0.1159945 | 1 | 0.984232 | 1 |
| IL18R1 | -0.0709189 | 1 | 0.939467 | 1 |
| OR2D2 | 0.238223 | 1 | 0.6312 | 1 |
| MESP1 | 0.58064854 | 1 | 0.289037 | 1 |
| OR11H4 | 0.11442026 | 1 | 0.75554 | 1 |
| MRTFB | -0.292805 | 1 | 1.163018 | 1 |
| GMIP | -0.2495825 | 1 | 1.120253 | 1 |
| OR8K5 | -0.2999481 | 1 | 1.171394 | 1 |

| | | | | |
|---|---|---|---|---|
| HSF5 | -0.1078763 | 1 | 0.979554 | 1 |
| HMGA1 | 1.89756007 | 1 | -1.02312 | 1 |
| SERP2 | 0.65007571 | 1 | 0.224839 | 1 |
| DEPTOR | -0.4361483 | 1 | 1.311488 | 1 |
| CDC5L | 1.80123473 | 1 | -0.92565 | 1 |
| CEBPE | 0.16843216 | 1 | 0.707649 | 1 |
| TAS2R5 | 0.27297918 | 1 | 0.605651 | 1 |
| OPHN1 | 0.0912442 | 1 | 0.788816 | 1 |
| ZCCHC17 | 0.51296637 | 1 | 0.367222 | 1 |
| MLST8 | 0.4088409 | 1 | 0.471605 | 1 |
| TRADD | 0.75617198 | 1 | 0.124478 | 1 |
| RANGAP1 | 0.35756531 | 1 | 0.526122 | 1 |
| ZNF764 | 0.0085139 | 1 | 0.875938 | 1 |
| POLR2K | 1.4647502 | 1 | -0.58021 | 1 |
| AGFG2 | 0.06331027 | 1 | 0.821865 | 1 |
| POLR3K | 0.56829048 | 1 | 0.317697 | 1 |
| PRKD2 | 0.85619515 | 1 | 0.03129 | 1 |
| PICALM | 1.46096417 | 1 | -0.5725 | 1 |
| E2F4 | 0.53564857 | 1 | 0.353401 | 1 |
| GNAS | 1.90353341 | 1 | -1.01262 | 1 |
| OR10G2 | 0.13504649 | 1 | 0.756474 | 1 |
| MIB2 | 0.23072424 | 1 | 0.661001 | 1 |
| NGFR | 0.09775731 | 1 | 0.794411 | 1 |
| ZFP90 | 0.84183241 | 1 | 0.050376 | 1 |
| ZMAT5 | 0.64644108 | 1 | 0.246347 | 1 |
| SEC14L2 | 0.26055578 | 1 | 0.63289 | 1 |
| LRRD1 | 0.05274054 | 1 | 0.840989 | 1 |
| MYOCD | 0.75329722 | 1 | 0.142812 | 1 |
| ARHGEF18 | -0.0249025 | 1 | 0.921164 | 1 |
| SIK1 | 0.93333835 | 1 | -0.03591 | 1 |
| THEMIS | 0.61250674 | 1 | 0.286135 | 1 |
| ZNF551 | 0.31571179 | 1 | 0.583119 | 1 |
| CLNK | 0.76393576 | 1 | 0.135273 | 1 |
| PRDM16 | -0.2998859 | 1 | 1.199362 | 1 |
| OPRL1 | -0.1561653 | 1 | 1.056301 | 1 |
| MZF1 | 0.35276658 | 1 | 0.54778 | 1 |
| RASGRF1 | 0.41117187 | 1 | 0.490229 | 1 |
| SLC2A8 | 0.31378332 | 1 | 0.587874 | 1 |
| FOXJ1 | 0.0118036 | 1 | 0.890498 | 1 |
| TICAM1 | 0.87447657 | 1 | 0.027949 | 1 |
| OR51S1 | 0.9063064 | 1 | -0.00368 | 1 |
| CNOT3 | 0.37845188 | 1 | 0.524257 | 1 |
| UNC5C | 0.41799799 | 1 | 0.485533 | 1 |
| THAP11 | 0.65556933 | 1 | 0.248382 | 1 |
| BRSK1 | 0.33281197 | 1 | 0.571306 | 1 |
| CLEC1B | 0.13857936 | 1 | 0.765707 | 1 |

| | | | | |
|---|---|---|---|---|
| TAF8 | -0.0070953 | 1 | 0.911465 | 1 |
| RIN3 | 0.13124885 | 1 | 0.773129 | 1 |
| GHRL | 0.1935721 | 1 | 0.711174 | 1 |
| TXNIP | -0.063882 | 1 | 0.969494 | 1 |
| PSENEN | 0.90372878 | 1 | 0.002872 | 1 |
| LBH | 0.84008526 | 1 | 0.068736 | 1 |
| OR4K2 | 0.00871556 | 1 | 0.900168 | 1 |
| ITPKB | 0.51876034 | 1 | 0.390448 | 1 |
| SSTR3 | 0.69809863 | 1 | 0.21157 | 1 |
| AKAP10 | 0.77459882 | 1 | 0.135643 | 1 |
| TBX22 | 0.82320983 | 1 | 0.087199 | 1 |
| RXFP4 | -0.0201247 | 1 | 0.931003 | 1 |
| RAB3D | -1.1921367 | 1 | 2.10458 | 1 |
| DTHD1 | 0.16159296 | 1 | 0.752651 | 1 |
| OLIG2 | -0.2697215 | 1 | 1.184743 | 1 |
| ZNF444 | 0.25470664 | 1 | 0.663339 | 1 |
| ZFP42 | 0.07043419 | 1 | 0.847677 | 1 |
| TAS1R3 | 0.10407857 | 1 | 0.814346 | 1 |
| MIF | 0.90940628 | 1 | 0.00966 | 1 |
| STAT5B | 0.43199044 | 1 | 0.487814 | 1 |
| ANKDD1A | -0.0804844 | 1 | 1.000366 | 1 |
| CCR3 | 0.93232 | 1 | -0.01118 | 1 |
| LAT | 0.0928785 | 1 | 0.828486 | 1 |
| DR1 | 2.67625975 | 1 | -1.75462 | 1 |
| TIGD2 | 0.48257613 | 1 | 0.439148 | 1 |
| ZBTB17 | 0.44804307 | 1 | 0.476236 | 1 |
| YWHAQ | 1.93700743 | 1 | -1.01242 | 1 |
| TRIM54 | 0.81962663 | 1 | 0.108589 | 1 |
| CD7 | 0.25875499 | 1 | 0.67131 | 1 |
| ZNF785 | 0.29739648 | 1 | 0.63525 | 1 |
| TLE5 | 0.41388708 | 1 | 0.51967 | 1 |
| PRDX1 | 0.83092909 | 1 | 0.103972 | 1 |
| SPDEF | 0.14473863 | 1 | 0.790217 | 1 |
| HPGDS | 0.77766095 | 1 | 0.158356 | 1 |
| OR8B12 | -0.0183973 | 1 | 0.95676 | 1 |
| GSC | 0.53221939 | 1 | 0.406581 | 1 |
| TLE6 | -0.1022367 | 1 | 1.043158 | 1 |
| KLRC2 | 0.40173078 | 1 | 0.539961 | 1 |
| DOCK7 | -0.4297645 | 1 | 1.3716 | 1 |
| HOXB1 | 0.02059433 | 1 | 0.921483 | 1 |
| FBXW11 | 0.87689774 | 1 | 0.065224 | 1 |
| BBC3 | 0.91150276 | 1 | 0.031158 | 1 |
| GPR75 | 0.02428189 | 1 | 0.918902 | 1 |
| CXCR6 | 0.18016898 | 1 | 0.763686 | 1 |
| PHF1 | 0.40421511 | 1 | 0.539646 | 1 |
| CD14 | 0.7071343 | 1 | 0.240417 | 1 |

| | | | | |
|---|---|---|---|---|
| TRIP6 | 0.71112405 | 1 | 0.237477 | 1 |
| OR7D2 | -0.5798446 | 1 | 1.530469 | 1 |
| TMEM229A | 0.16231672 | 1 | 0.788895 | 1 |
| TCEAL3 | 0.2086954 | 1 | 0.742801 | 1 |
| TBX4 | 0.26940313 | 1 | 0.682879 | 1 |
| ZNF341 | 0.61319051 | 1 | 0.339794 | 1 |
| FASTK | 0.25526255 | 1 | 0.699306 | 1 |
| LGALS9 | 0.20815331 | 1 | 0.747167 | 1 |
| GUCY2D | 0.68751726 | 1 | 0.268914 | 1 |
| RAB35 | 0.57261962 | 1 | 0.384761 | 1 |
| HRAS | 0.81625882 | 1 | 0.141132 | 1 |
| GPR176 | -0.2683222 | 1 | 1.226324 | 1 |
| TXNDC17 | 0.60907026 | 1 | 0.353315 | 1 |
| SPRED3 | 1.17622332 | 1 | -0.21345 | 1 |
| LEF1 | 0.92148282 | 1 | 0.041387 | 1 |
| NDUFA13 | 0.84607245 | 1 | 0.11696 | 1 |
| PPP5C | 0.4993745 | 1 | 0.464407 | 1 |
| CLIC1 | 0.6228508 | 1 | 0.341241 | 1 |
| HTR3B | 0.48514473 | 1 | 0.480158 | 1 |
| PRKAG1 | 0.30022114 | 1 | 0.665804 | 1 |
| KAT5 | 0.76224627 | 1 | 0.206141 | 1 |
| RHOB | -0.78067 | 1 | 1.749525 | 1 |
| ZNF33A | 0.10887323 | 1 | 0.86443 | 1 |
| GGCT | 1.22589059 | 1 | -0.25131 | 1 |
| SMYD1 | 0.2599998 | 1 | 0.714594 | 1 |
| TNFRSF11A | 0.48843288 | 1 | 0.488613 | 1 |
| ZNF549 | -0.2679747 | 1 | 1.245066 | 1 |
| ADAM11 | -0.8859577 | 1 | 1.864141 | 1 |
| PRAM1 | 0.21108648 | 1 | 0.767174 | 1 |
| ZNF432 | -0.0068852 | 1 | 0.985153 | 1 |
| PLCL1 | -0.2399456 | 1 | 1.218359 | 1 |
| OR56A5 | 0.10298791 | 1 | 0.875961 | 1 |
| DPYSL5 | 0.6750798 | 1 | 0.303932 | 1 |
| CSNK1D | 0.62239693 | 1 | 0.358262 | 1 |
| CNTNAP3 | 0.11703923 | 1 | 0.865118 | 1 |
| RPS6KB2 | 0.41641646 | 1 | 0.57002 | 1 |
| CENPB | 0.84152177 | 1 | 0.145742 | 1 |
| HAT1 | 1.43565711 | 1 | -0.44751 | 1 |
| BLK | 0.91114611 | 1 | 0.077699 | 1 |
| OLIG1 | -1.0874872 | 1 | 2.079643 | 1 |
| TRIM13 | 1.24195836 | 1 | -0.24531 | 1 |
| SEMA4C | 0.32392783 | 1 | 0.674018 | 1 |
| TRPV1 | 0.57942522 | 1 | 0.418572 | 1 |
| ZNF770 | 1.54665889 | 1 | -0.5465 | 1 |
| SIK2 | 0.37156638 | 1 | 0.628874 | 1 |
| PDE4DIP | 0.32371332 | 1 | 0.679126 | 1 |

| | | | | |
|---|---|---|---|---|
| RAC3 | 0.51602539 | 1 | 0.487555 | 1 |
| DLG4 | 0.39912015 | 1 | 0.604861 | 1 |
| AGAP2 | 0.04329087 | 1 | 0.962867 | 1 |
| PITX1 | 0.14578511 | 1 | 0.862419 | 1 |
| NFATC2IP | -0.6936035 | 1 | 1.705189 | 1 |
| GLIS3 | -0.4267114 | 1 | 1.439627 | 1 |
| CAVIN1 | 0.10282367 | 1 | 0.910554 | 1 |
| PA2G4 | 1.49256945 | 1 | -0.47893 | 1 |
| ASB5 | 0.11051582 | 1 | 0.903273 | 1 |
| ADGRE3 | 0.26419121 | 1 | 0.750553 | 1 |
| IRX3 | 0.95590113 | 1 | 0.059453 | 1 |
| CRLF2 | 0.6237695 | 1 | 0.391823 | 1 |
| RCBTB1 | 0.18679985 | 1 | 0.82895 | 1 |
| MED19 | 0.6930111 | 1 | 0.327496 | 1 |
| RGL2 | 0.39813133 | 1 | 0.623409 | 1 |
| NUMA1 | 0.15519073 | 1 | 0.868528 | 1 |
| CASS4 | 0.26485752 | 1 | 0.758959 | 1 |
| FGF1 | -0.9704706 | 1 | 1.994834 | 1 |
| CCL5 | -0.0783141 | 1 | 1.103277 | 1 |
| ZDHHC3 | -0.1419136 | 1 | 1.168194 | 1 |
| ETV3L | 2.37132897 | 1 | -1.34455 | 1 |
| DKKL1 | -0.0682303 | 1 | 1.095661 | 1 |
| EIF2A | 1.50756937 | 1 | -0.47973 | 1 |
| HOXA6 | 0.70938723 | 1 | 0.318602 | 1 |
| ZNF594 | 0.79098328 | 1 | 0.238401 | 1 |
| MECOM | 0.66987241 | 1 | 0.360019 | 1 |
| TSC2 | 0.27237705 | 1 | 0.758091 | 1 |
| ZSCAN5B | 0.09273365 | 1 | 0.938077 | 1 |
| ZBTB4 | 0.359941 | 1 | 0.673537 | 1 |
| DTNA | 0.6690631 | 1 | 0.365183 | 1 |
| MNX1 | 0.73111201 | 1 | 0.303715 | 1 |
| LPAR2 | 0.82419175 | 1 | 0.210792 | 1 |
| CHD8 | 0.24410237 | 1 | 0.791217 | 1 |
| OSMR | 0.88501296 | 1 | 0.151713 | 1 |
| MED15 | 1.16817849 | 1 | -0.12865 | 1 |
| ZBTB48 | 0.46058202 | 1 | 0.580206 | 1 |
| IL13RA1 | 0.7447902 | 1 | 0.297174 | 1 |
| TARBP2 | 0.7720905 | 1 | 0.27041 | 1 |
| GLRA2 | -0.3179838 | 1 | 1.361966 | 1 |
| OR10G9 | 0.90984244 | 1 | 0.134203 | 1 |
| PPT1 | 0.82921815 | 1 | 0.215696 | 1 |
| SPRY4 | 0.84396333 | 1 | 0.201361 | 1 |
| OSTF1 | 0.14221371 | 1 | 0.903835 | 1 |
| ZNF771 | 0.49697996 | 1 | 0.549883 | 1 |
| MED27 | 1.91995101 | 1 | -0.8728 | 1 |
| SIGLEC8 | 0.29202317 | 1 | 0.755187 | 1 |

| | | | | |
|---|---|---|---|---|
| GPR78 | -0.0231663 | 1 | 1.071759 | 1 |
| PCSK1N | 0.20370522 | 1 | 0.846178 | 1 |
| ARHGAP17 | 1.03736482 | 1 | 0.017501 | 1 |
| HTR7 | -0.3658983 | 1 | 1.420947 | 1 |
| RAMP2 | 0.98606003 | 1 | 0.071281 | 1 |
| ZFYVE1 | 0.1946191 | 1 | 0.863155 | 1 |
| ZNF689 | 0.82955444 | 1 | 0.228899 | 1 |
| SQSTM1 | 0.27225875 | 1 | 0.787655 | 1 |
| RAB40B | 0.33612673 | 1 | 0.724314 | 1 |
| ZNF57 | 1.31356332 | 1 | -0.25015 | 1 |
| ADGRF3 | 0.47300418 | 1 | 0.591097 | 1 |
| TXNRD1 | 1.84867937 | 1 | -0.78319 | 1 |
| DMRT3 | 1.0304035 | 1 | 0.037466 | 1 |
| OR51M1 | -0.0250405 | 1 | 1.092952 | 1 |
| GABBR1 | -0.0871307 | 1 | 1.155798 | 1 |
| ZCCHC12 | -0.6054678 | 1 | 1.677318 | 1 |
| TARBP1 | 0.89057286 | 1 | 0.181629 | 1 |
| SCAND1 | 0.96608918 | 1 | 0.106697 | 1 |
| OR7E24 | 0.05059672 | 1 | 1.023599 | 1 |
| RNF4 | 1.34256001 | 1 | -0.26695 | 1 |
| MMS19 | 1.02048821 | 1 | 0.055917 | 1 |
| DRGX | 0.07611272 | 1 | 1.003326 | 1 |
| ZNF77 | 1.10046909 | 1 | -0.01853 | 1 |
| ZNF835 | 0.00534972 | 1 | 1.077015 | 1 |
| ZNF485 | 1.25188507 | 1 | -0.16685 | 1 |
| IL2RA | 0.0891763 | 1 | 0.997291 | 1 |
| SMOC1 | -1.1615718 | 1 | 2.249676 | 1 |
| FLT3 | -0.4135743 | 1 | 1.501968 | 1 |
| ZNF335 | 0.66177581 | 1 | 0.427467 | 1 |
| ELL3 | 0.84667056 | 1 | 0.243991 | 1 |
| DDX5 | 1.15999455 | 1 | -0.06911 | 1 |
| RPS6 | 1.17981635 | 1 | -0.08661 | 1 |
| IRF1 | -0.0099614 | 1 | 1.103237 | 1 |
| PPP1R9B | 1.18173171 | 1 | -0.08806 | 1 |
| TANK | 1.3865803 | 1 | -0.29246 | 1 |
| CCRL2 | 0.22682977 | 1 | 0.867358 | 1 |
| RIPK3 | 0.06269769 | 1 | 1.032159 | 1 |
| CCM2 | 0.65522758 | 1 | 0.439882 | 1 |
| ZNF778 | 0.13642292 | 1 | 0.960672 | 1 |
| TMEM101 | 0.84153575 | 1 | 0.256525 | 1 |
| VSX2 | -0.2632799 | 1 | 1.361776 | 1 |
| ZNF839 | 0.28171565 | 1 | 0.818074 | 1 |
| MLLT1 | 0.29179166 | 1 | 0.808019 | 1 |
| PYCARD | 0.28550405 | 1 | 0.814417 | 1 |
| RXRB | 0.83327699 | 1 | 0.268738 | 1 |
| GPR149 | 0.89891736 | 1 | 0.204227 | 1 |

| | | | | |
|---|---|---|---|---|
| PSMC3 | 0.58504848 | 1 | 0.520052 | 1 |
| ATF4 | 1.02141667 | 1 | 0.083896 | 1 |
| RASGRF2 | -0.1334653 | 1 | 1.239618 | 1 |
| PLCD4 | 0.18758898 | 1 | 0.918643 | 1 |
| MUC1 | 0.71973081 | 1 | 0.388082 | 1 |
| FGD6 | 0.4940021 | 1 | 0.614405 | 1 |
| OR2J1 | 1.06299772 | 1 | 0.045538 | 1 |
| PIDD1 | 0.25092994 | 1 | 0.858634 | 1 |
| PLCH2 | 0.18925554 | 1 | 0.921141 | 1 |
| LRRK1 | 0.36777762 | 1 | 0.743366 | 1 |
| MED8 | 2.14612129 | 1 | -1.0337 | 1 |
| PREX1 | 0.13450826 | 1 | 0.978027 | 1 |
| IFNGR2 | 0.25049335 | 1 | 0.866426 | 1 |
| PRDX4 | 1.73339691 | 1 | -0.61572 | 1 |
| TNFSF10 | 1.53030327 | 1 | -0.41234 | 1 |
| BCR | 0.05123134 | 1 | 1.067731 | 1 |
| RPS14 | 1.09935432 | 1 | 0.021575 | 1 |
| P2RY2 | 0.04551713 | 1 | 1.077064 | 1 |
| OR6C68 | 1.01728346 | 1 | 0.10606 | 1 |
| ZNF536 | -0.2694623 | 1 | 1.394043 | 1 |
| BEX3 | 0.10873943 | 1 | 1.015896 | 1 |
| SOX3 | 0.20024723 | 1 | 0.926226 | 1 |
| ZNF598 | 0.57699206 | 1 | 0.55303 | 1 |
| C3AR1 | 0.06012439 | 1 | 1.070316 | 1 |
| SIPA1L1 | 1.27629607 | 1 | -0.14457 | 1 |
| RAB1B | 0.92142195 | 1 | 0.211092 | 1 |
| GPR31 | 1.25873799 | 1 | -0.1238 | 1 |
| ZFR2 | 0.98875014 | 1 | 0.146932 | 1 |
| HOXC13 | 0.31543947 | 1 | 0.820307 | 1 |
| ZNF524 | 0.93802842 | 1 | 0.198355 | 1 |
| LZTR1 | 0.75911103 | 1 | 0.37752 | 1 |
| EMX1 | 0.2375183 | 1 | 0.900446 | 1 |
| CHRNA6 | 0.3261052 | 1 | 0.812662 | 1 |
| ACTR5 | 0.86700835 | 1 | 0.272034 | 1 |
| RGMA | 0.95774977 | 1 | 0.182554 | 1 |
| HDAC5 | 0.76500389 | 1 | 0.375805 | 1 |
| MAP3K6 | 0.50271121 | 1 | 0.641057 | 1 |
| FAF1 | 1.33144649 | 1 | -0.18708 | 1 |
| GRM7 | 0.90743139 | 1 | 0.237529 | 1 |
| DERL2 | 1.49631055 | 1 | -0.35078 | 1 |
| ITGA10 | 1.09160744 | 1 | 0.059971 | 1 |
| ZC3H4 | 0.79057481 | 1 | 0.361076 | 1 |
| ZNF512B | -1.1330058 | 1 | 2.286416 | 1 |
| CDK7 | 1.45248268 | 1 | -0.29897 | 1 |
| OR51B2 | 0.48283148 | 1 | 0.671356 | 1 |
| SMARCB1 | 0.59162231 | 1 | 0.562835 | 1 |

| | | | | |
|---|---|---|---|---|
| CHL1 | 1.23670149 | 1 | -0.08209 | 1 |
| CILK1 | -1.3740044 | 1 | 2.529877 | 1 |
| VOPP1 | 0.90390017 | 1 | 0.253155 | 1 |
| TREML1 | 0.1494245 | 1 | 1.007721 | 1 |
| DAXX | 1.44608789 | 1 | -0.28806 | 1 |
| OR4D5 | 1.21794049 | 1 | -0.05913 | 1 |
| OR13C2 | 0.30642899 | 1 | 0.852725 | 1 |
| RIIAD1 | -0.1322903 | 1 | 1.294135 | 1 |
| RPS6KA4 | 0.17361808 | 1 | 0.989843 | 1 |
| HSP90AA1 | 1.55770335 | 1 | -0.39285 | 1 |
| FKBP8 | 0.60531131 | 1 | 0.559767 | 1 |
| CYTH2 | 0.88470914 | 1 | 0.283964 | 1 |
| CCR2 | 0.30131958 | 1 | 0.868059 | 1 |
| BSG | 1.31564479 | 1 | -0.14504 | 1 |
| PLCD1 | 0.77872145 | 1 | 0.393018 | 1 |
| RNF43 | 0.02229286 | 1 | 1.149754 | 1 |
| NPAS1 | 1.05252627 | 1 | 0.11993 | 1 |
| SH2B1 | 0.09633915 | 1 | 1.076121 | 1 |
| PLP2 | 0.98717677 | 1 | 0.187387 | 1 |
| ZNF550 | 0.14477823 | 1 | 1.030202 | 1 |
| PCGF6 | 1.26372437 | 1 | -0.08822 | 1 |
| ERG | 0.12249524 | 1 | 1.054394 | 1 |
| GRTP1 | 0.16680654 | 1 | 1.011072 | 1 |
| SLC4A11 | 1.13048523 | 1 | 0.048308 | 1 |
| FRAT1 | 0.23502621 | 1 | 0.944086 | 1 |
| ITGB4 | -0.0251079 | 1 | 1.204311 | 1 |
| HSF1 | 1.07032396 | 1 | 0.11146 | 1 |
| PPP2R1A | 1.15735951 | 1 | 0.025105 | 1 |
| ERC1 | 0.95048556 | 1 | 0.23531 | 1 |
| EBF3 | 0.01142033 | 1 | 1.179284 | 1 |
| LHCGR | 0.84588452 | 1 | 0.347485 | 1 |
| GNL1 | 0.50225468 | 1 | 0.691728 | 1 |
| HACD3 | 1.430543 | 1 | -0.23411 | 1 |
| OR10AD1 | 1.06285996 | 1 | 0.134053 | 1 |
| MRGPRG | 1.15408888 | 1 | 0.043811 | 1 |
| NTF3 | 1.18511261 | 1 | 0.01302 | 1 |
| CHP1 | 1.32988514 | 1 | -0.13121 | 1 |
| PKN1 | 1.12839875 | 1 | 0.070383 | 1 |
| SLC9A1 | 1.34953512 | 1 | -0.15019 | 1 |
| NAT14 | 0.93939976 | 1 | 0.261263 | 1 |
| TAF1L | 0.05998697 | 1 | 1.141396 | 1 |
| CLEC4A | 0.50543317 | 1 | 0.69627 | 1 |
| ZSCAN32 | 0.74700262 | 1 | 0.455175 | 1 |
| NFIX | 0.0517404 | 1 | 1.151646 | 1 |
| TIAM1 | 0.8838425 | 1 | 0.319695 | 1 |
| HCLS1 | 0.43695785 | 1 | 0.766664 | 1 |

| | | | | |
|---|---|---|---|---|
| ZNF428 | 0.97052601 | 1 | 0.233671 | 1 |
| TLR5 | 0.56708567 | 1 | 0.638622 | 1 |
| WWC1 | 0.03150661 | 1 | 1.174414 | 1 |
| OR5H2 | 0.12751745 | 1 | 1.08064 | 1 |
| SH3BP5 | 0.62034563 | 1 | 0.588454 | 1 |
| CD24 | 0.05437915 | 1 | 1.158226 | 1 |
| PLXNB2 | 0.10672881 | 1 | 1.105994 | 1 |
| MAP2K6 | 0.98945333 | 1 | 0.223532 | 1 |
| IGBP1 | 1.30828851 | 1 | -0.09488 | 1 |
| NPFFR1 | 0.09531501 | 1 | 1.11981 | 1 |
| ZCCHC24 | 0.55919226 | 1 | 0.656409 | 1 |
| AKAP13 | -0.1194419 | 1 | 1.338571 | 1 |
| ARHGEF19 | 1.25904369 | 1 | -0.0394 | 1 |
| TPM4 | 2.24416897 | 1 | -1.02072 | 1 |
| GPRC5B | 1.04707798 | 1 | 0.177507 | 1 |
| F2RL2 | 1.077086 | 1 | 0.149616 | 1 |
| TBC1D13 | 0.89229531 | 1 | 0.337065 | 1 |
| STAT2 | 0.76147504 | 1 | 0.470082 | 1 |
| MAP3K11 | 0.69487239 | 1 | 0.536731 | 1 |
| PPP2R5B | 0.78184052 | 1 | 0.450968 | 1 |
| PREX2 | 0.82653227 | 1 | 0.407698 | 1 |
| OPN5 | 1.27697478 | 1 | -0.03919 | 1 |
| HGS | 0.6205276 | 1 | 0.617415 | 1 |
| DUSP1 | -0.8017005 | 1 | 2.040001 | 1 |
| TSPAN8 | 0.21345041 | 1 | 1.02561 | 1 |
| PLCXD3 | 1.15672994 | 1 | 0.084202 | 1 |
| FFAR3 | 0.74208191 | 1 | 0.501786 | 1 |
| FEV | 0.75435286 | 1 | 0.490327 | 1 |
| GTF2H5 | 1.09925781 | 1 | 0.147309 | 1 |
| STOML3 | 1.19522045 | 1 | 0.054467 | 1 |
| IGF2 | 1.52637485 | 1 | -0.27637 | 1 |
| CALM3 | -0.3032959 | 1 | 1.554838 | 1 |
| RABL6 | 0.88121272 | 1 | 0.372123 | 1 |
| SPSB3 | 0.34926273 | 1 | 0.907198 | 1 |
| KCNIP3 | -0.8461156 | 1 | 2.103027 | 1 |
| RFX5 | 0.42392818 | 1 | 0.833121 | 1 |
| IRAK1 | 0.28912675 | 1 | 0.969059 | 1 |
| AMFR | 0.47084622 | 1 | 0.793105 | 1 |
| FGFR3 | 0.0669874 | 1 | 1.197879 | 1 |
| DTX3 | 0.19344029 | 1 | 1.074545 | 1 |
| M6PR | 1.25768439 | 1 | 0.011954 | 1 |
| MAP4K2 | 0.36535493 | 1 | 0.9063 | 1 |
| KRT17 | 0.39966713 | 1 | 0.87266 | 1 |
| SGSM2 | 0.93131479 | 1 | 0.342502 | 1 |
| CD244 | 0.19012219 | 1 | 1.084228 | 1 |
| DGKA | 0.09298618 | 1 | 1.181766 | 1 |

| | | | | |
|---|---|---|---|---|
| FGD3 | 0.0246797 | 1 | 1.250571 | 1 |
| ZC3H12A | 1.04896812 | 1 | 0.226486 | 1 |
| CCL4L2 | 1.0389226 | 1 | 0.241949 | 1 |
| C18orf32 | 0.16401524 | 1 | 1.117046 | 1 |
| TBC1D22B | 1.01587171 | 1 | 0.265236 | 1 |
| STYXL1 | 0.33188547 | 1 | 0.949759 | 1 |
| NDUFS4 | 1.46278352 | 1 | -0.1806 | 1 |
| CARD10 | -0.19354 | 1 | 1.476077 | 1 |
| DCDC2B | 0.14380763 | 1 | 1.139977 | 1 |
| GDF3 | 1.19212955 | 1 | 0.09431 | 1 |
| PLCXD1 | -0.4149266 | 1 | 1.703116 | 1 |
| RFX4 | 0.21840754 | 1 | 1.070186 | 1 |
| TP53TG5 | 0.20007011 | 1 | 1.089189 | 1 |
| MRPL28 | 1.0535121 | 1 | 0.235856 | 1 |
| CPLANE2 | 0.86635439 | 1 | 0.426898 | 1 |
| PLD1 | -0.4529993 | 1 | 1.750025 | 1 |
| PDCD6 | 0.40006463 | 1 | 0.897274 | 1 |
| ATXN7L3 | 1.20040488 | 1 | 0.097606 | 1 |
| NR2F6 | 1.0232601 | 1 | 0.274803 | 1 |
| WNT5A | 1.20604642 | 1 | 0.094653 | 1 |
| CYTL1 | 1.27656263 | 1 | 0.024756 | 1 |
| FNTA | 0.79803286 | 1 | 0.503807 | 1 |
| EPAS1 | 1.08022882 | 1 | 0.22176 | 1 |
| AKT1S1 | 0.77746046 | 1 | 0.527133 | 1 |
| TIMM50 | 1.06438624 | 1 | 0.241324 | 1 |
| ZNF526 | 0.83353894 | 1 | 0.477173 | 1 |
| OR51V1 | -0.0908701 | 1 | 1.402735 | 1 |
| HDAC8 | 1.27785887 | 1 | 0.034091 | 1 |
| RUNDC3A | -0.1140086 | 1 | 1.427109 | 1 |
| CREB3L1 | 1.18448597 | 1 | 0.130687 | 1 |
| TNFSF13B | -0.1823664 | 1 | 1.498767 | 1 |
| PPARGC1B | 1.10032303 | 1 | 0.216412 | 1 |
| RPN1 | 1.65782832 | 1 | -0.34043 | 1 |
| TAMALIN | 0.5596573 | 1 | 0.761073 | 1 |
| SIRT6 | 0.7585246 | 1 | 0.563962 | 1 |
| MYO9B | 1.17356123 | 1 | 0.149074 | 1 |
| KALRN | -0.0342757 | 1 | 1.357954 | 1 |
| HDAC7 | 1.11918872 | 1 | 0.205467 | 1 |
| HABP4 | 0.17528102 | 1 | 1.149816 | 1 |
| RGS4 | 1.38569636 | 1 | -0.06008 | 1 |
| ERBB3 | -1.9949751 | 1 | 3.323119 | 1 |
| MED16 | 0.6217311 | 1 | 0.706456 | 1 |
| ZNF35 | 1.50057727 | 1 | -0.17047 | 1 |
| GABRB1 | 0.04189688 | 1 | 1.288571 | 1 |
| CDK4 | 1.44382519 | 1 | -0.11333 | 1 |
| RFX8 | 1.13617567 | 1 | 0.197851 | 1 |

| | | | | |
|---|---|---|---|---|
| FLYWCH1 | 0.27151378 | 1 | 1.063163 | 1 |
| MAMSTR | 0.13498479 | 1 | 1.200851 | 1 |
| TAF6L | 0.32244439 | 1 | 1.013518 | 1 |
| OSTN | 1.25426128 | 1 | 0.082888 | 1 |
| GPS2 | 1.13192605 | 1 | 0.205988 | 1 |
| ARL2BP | 1.4988662 | 1 | -0.16047 | 1 |
| ZFAND2B | 0.68172246 | 1 | 0.657512 | 1 |
| CCND3 | 1.14299427 | 1 | 0.196623 | 1 |
| CBX7 | 0.71719898 | 1 | 0.624658 | 1 |
| CASKIN1 | 0.98057766 | 1 | 0.364635 | 1 |
| UBASH3A | 1.04333259 | 1 | 0.303619 | 1 |
| FOXD4L4 | 0.1432828 | 1 | 1.206664 | 1 |
| CLEC7A | 0.18827066 | 1 | 1.163745 | 1 |
| NPM1 | 1.68417981 | 1 | -0.33185 | 1 |
| SARM1 | 0.11361177 | 1 | 1.240541 | 1 |
| PLAT | 0.11409149 | 1 | 1.240475 | 1 |
| ZNF467 | 0.3302965 | 1 | 1.02511 | 1 |
| ZNF862 | -0.0334994 | 1 | 1.38909 | 1 |
| HOPX | -0.4605968 | 1 | 1.81735 | 1 |
| NFATC2 | 0.13876211 | 1 | 1.219926 | 1 |
| ESX1 | 1.17179445 | 1 | 0.188074 | 1 |
| PBX2 | 0.25665373 | 1 | 1.104468 | 1 |
| UBE2N | 1.81444262 | 1 | -0.45187 | 1 |
| AFAP1L2 | -0.9595287 | 1 | 2.323596 | 1 |
| KCNIP2 | 0.30605367 | 1 | 1.058757 | 1 |
| CBFB | 0.11835474 | 1 | 1.24647 | 1 |
| ARNTL | -0.0062698 | 1 | 1.373102 | 1 |
| SARNP | 2.37269384 | 1 | -1.00531 | 1 |
| MAP3K12 | 0.27251304 | 1 | 1.096864 | 1 |
| MAPK4 | 0.28667247 | 1 | 1.084096 | 1 |
| ZNF668 | 0.5325713 | 1 | 0.839782 | 1 |
| OR4M2 | -0.0239884 | 1 | 1.397873 | 1 |
| TNFRSF10D | 0.08433751 | 1 | 1.289925 | 1 |
| ZNF710 | 0.14566004 | 1 | 1.228974 | 1 |
| CIDEB | 0.11098666 | 1 | 1.265028 | 1 |
| ZC3HC1 | 1.52621857 | 1 | -0.14905 | 1 |
| ZNF219 | 0.13094294 | 1 | 1.247091 | 1 |
| ZNF366 | -0.0593434 | 1 | 1.443879 | 1 |
| OR10G4 | 1.11923495 | 1 | 0.26629 | 1 |
| DNM2 | 0.26369667 | 1 | 1.12186 | 1 |
| HMOX1 | 0.13025164 | 1 | 1.257983 | 1 |
| NR1D1 | 1.18709517 | 1 | 0.202186 | 1 |
| GATAD2A | 1.53969632 | 1 | -0.14955 | 1 |
| GBX2 | 0.51824896 | 1 | 0.874325 | 1 |
| ELL | 0.2209092 | 1 | 1.173851 | 1 |
| PPP1R17 | 0.22954326 | 1 | 1.173414 | 1 |

| | | | | |
|---|---|---|---|---|
| MAPK7 | 0.52832914 | 1 | 0.87522 | 1 |
| USF2 | 0.98042326 | 1 | 0.423844 | 1 |
| TMEM30A | 1.04614772 | 1 | 0.358376 | 1 |
| WT1 | 0.7088155 | 1 | 0.696533 | 1 |
| C9 | 0.04739319 | 1 | 1.358457 | 1 |
| UTP11 | 2.16599614 | 1 | -0.75983 | 1 |
| DACT1 | 1.9034037 | 1 | -0.4963 | 1 |
| ZBTB16 | 0.90421798 | 1 | 0.504454 | 1 |
| MGLL | 1.23850566 | 1 | 0.17039 | 1 |
| GABRP | 0.47432728 | 1 | 0.935 | 1 |
| CTBP1 | 1.09229148 | 1 | 0.319306 | 1 |
| NCAM1 | -0.8334319 | 1 | 2.245247 | 1 |
| OR51A4 | 1.07834156 | 1 | 0.3355 | 1 |
| TTN | 0.10767345 | 1 | 1.306354 | 1 |
| MTOR | 0.61054387 | 1 | 0.805127 | 1 |
| THY1 | 0.67737327 | 1 | 0.740655 | 1 |
| HMG20B | 1.02511396 | 1 | 0.393149 | 1 |
| TGFB1 | 0.48029636 | 1 | 0.940221 | 1 |
| FGF22 | 0.1346493 | 1 | 1.288831 | 1 |
| IQSEC2 | 0.13320495 | 1 | 1.292119 | 1 |
| FZD10 | 1.51006572 | 1 | -0.08405 | 1 |
| ZNF593 | 1.44632106 | 1 | -0.01906 | 1 |
| CXCL2 | 0.45597383 | 1 | 0.973222 | 1 |
| RAB11A | 1.72137982 | 1 | -0.2895 | 1 |
| OR11H12 | 0.30688267 | 1 | 1.127969 | 1 |
| MAFK | 0.53633222 | 1 | 0.898753 | 1 |
| MEIS1 | 1.10664251 | 1 | 0.328875 | 1 |
| PBX1 | 2.05055018 | 1 | -0.61416 | 1 |
| ITGA2B | 1.1298646 | 1 | 0.307457 | 1 |
| ZFAND6 | 2.45253167 | 1 | -1.01395 | 1 |
| PLEKHG5 | 0.38412422 | 1 | 1.056409 | 1 |
| LHX1 | 1.30425858 | 1 | 0.138372 | 1 |
| DOK2 | 0.82247834 | 1 | 0.624568 | 1 |
| CCL13 | 0.26542501 | 1 | 1.182379 | 1 |
| ECM1 | 0.43430574 | 1 | 1.015478 | 1 |
| IL21R | 0.08831793 | 1 | 1.361741 | 1 |
| FIBP | 1.22548383 | 1 | 0.225341 | 1 |
| IDH2 | 0.60753824 | 1 | 0.843433 | 1 |
| PUF60 | 1.15891059 | 1 | 0.295772 | 1 |
| RFXANK | 0.9317514 | 1 | 0.523228 | 1 |
| OR52N1 | 0.24079405 | 1 | 1.215443 | 1 |
| NCLN | 1.38643023 | 1 | 0.070409 | 1 |
| OR4F15 | 0.57944576 | 1 | 0.878129 | 1 |
| NKX2.5 | 0.80957359 | 1 | 0.650214 | 1 |
| ZBTB46 | 0.85329244 | 1 | 0.610378 | 1 |
| CRADD | 1.3145987 | 1 | 0.149194 | 1 |

| | | | | |
|---|---|---|---|---|
| IFNG | 1.43987906 | 1 | 0.023983 | 1 |
| CCAR1 | 1.06506871 | 1 | 0.399742 | 1 |
| PDPN | 0.08736039 | 1 | 1.380109 | 1 |
| MC5R | -0.0297907 | 1 | 1.497622 | 1 |
| ZNF106 | 1.52410965 | 1 | -0.05561 | 1 |
| PDGFRB | 0.00840973 | 1 | 1.461773 | 1 |
| ZNF683 | -0.0541753 | 1 | 1.528895 | 1 |
| ZNF541 | 1.26031202 | 1 | 0.215133 | 1 |
| TAOK3 | 0.11606915 | 1 | 1.361057 | 1 |
| UNC5D | 0.4278101 | 1 | 1.051599 | 1 |
| PCP2 | 0.70011052 | 1 | 0.779644 | 1 |
| SIVA1 | 1.13272797 | 1 | 0.347122 | 1 |
| FLRT1 | -0.0491647 | 1 | 1.530891 | 1 |
| SNAI1 | -0.0167088 | 1 | 1.499222 | 1 |
| UNC5B | 0.52528253 | 1 | 0.9599 | 1 |
| CDS1 | 0.19452782 | 1 | 1.294094 | 1 |
| PPARD | 0.81015437 | 1 | 0.678718 | 1 |
| NCK2 | 0.44286581 | 1 | 1.046315 | 1 |
| PRMT1 | 1.25161588 | 1 | 0.244609 | 1 |
| ZNF513 | 1.41142454 | 1 | 0.08576 | 1 |
| OR7C2 | 1.37524974 | 1 | 0.12244 | 1 |
| OR10A2 | 0.58125477 | 1 | 0.918732 | 1 |
| ENDOG | 0.59010955 | 1 | 0.911838 | 1 |
| FKBP1A | 1.79951522 | 1 | -0.29539 | 1 |
| ARHGAP31 | -0.0317233 | 1 | 1.536669 | 1 |
| MRGPRD | 1.0527875 | 1 | 0.454915 | 1 |
| KREMEN2 | 1.46686199 | 1 | 0.042462 | 1 |
| ZNF671 | 0.19666441 | 1 | 1.313325 | 1 |
| NFKBIA | 0.1991209 | 1 | 1.31281 | 1 |
| MED10 | 2.1149834 | 1 | -0.59877 | 1 |
| SH3GL1 | 1.34765466 | 1 | 0.169155 | 1 |
| CCN6 | 0.5871744 | 1 | 0.930175 | 1 |
| OR2T35 | 0.03402199 | 1 | 1.484521 | 1 |
| PTPRN | 0.16498969 | 1 | 1.355854 | 1 |
| BAP1 | 1.30557439 | 1 | 0.220729 | 1 |
| TCF25 | 0.72859062 | 1 | 0.799993 | 1 |
| ACAP3 | 0.78786688 | 1 | 0.740777 | 1 |
| NGEF | 1.46054789 | 1 | 0.068795 | 1 |
| EAF2 | 0.11218883 | 1 | 1.418988 | 1 |
| PTPRE | 1.31761275 | 1 | 0.215036 | 1 |
| HIF3A | 0.11634654 | 1 | 1.417665 | 1 |
| HMGN1 | 1.19613876 | 1 | 0.338026 | 1 |
| GLI1 | 1.36667494 | 1 | 0.168743 | 1 |
| BRF2 | 1.07558859 | 1 | 0.460546 | 1 |
| COMMD7 | 1.70086871 | 1 | -0.16465 | 1 |
| FICD | 1.18971414 | 1 | 0.347013 | 1 |

| | | | | |
|---|---|---|---|---|
| E4F1 | 0.26459252 | 1 | 1.273846 | 1 |
| NUP93 | 0.74522899 | 1 | 0.794332 | 1 |
| ARHGEF11 | 0.18551598 | 1 | 1.356714 | 1 |
| ARHGAP22 | 0.27605923 | 1 | 1.268327 | 1 |
| P2RX4 | 1.42785573 | 1 | 0.116743 | 1 |
| ZNF415 | -0.3972063 | 1 | 1.943285 | 1 |
| MRGPRF | 0.09288144 | 1 | 1.453324 | 1 |
| ASGR2 | 1.41902119 | 1 | 0.12982 | 1 |
| CCL1 | -0.0076434 | 1 | 1.557173 | 1 |
| OR4C46 | 2.24244603 | 1 | -0.6928 | 1 |
| P2RY14 | 1.61184857 | 1 | -0.05951 | 1 |
| PLP1 | 0.40999173 | 1 | 1.144193 | 1 |
| ZNF287 | 1.39489864 | 1 | 0.16134 | 1 |
| SP8 | 0.6769904 | 1 | 0.880569 | 1 |
| DDR2 | 0.32698311 | 1 | 1.235665 | 1 |
| JUND | 0.12493146 | 1 | 1.438908 | 1 |
| OR2B2 | 0.29946117 | 1 | 1.268442 | 1 |
| ADCY8 | 1.63672071 | 1 | -0.06791 | 1 |
| RAB15 | 0.16949718 | 1 | 1.399594 | 1 |
| PFDN5 | 1.6596531 | 1 | -0.08604 | 1 |
| TADA3 | 1.75717184 | 1 | -0.1825 | 1 |
| DAPK2 | 1.11121824 | 1 | 0.46391 | 1 |
| LTB | 1.02457174 | 1 | 0.551027 | 1 |
| EPHA10 | -0.0920586 | 1 | 1.667733 | 1 |
| NR4A1 | 0.84678457 | 1 | 0.729682 | 1 |
| BIRC7 | -0.0052068 | 1 | 1.581722 | 1 |
| FURIN | 0.33827253 | 1 | 1.238406 | 1 |
| TMPRSS2 | 0.03660868 | 1 | 1.541293 | 1 |
| CXXC4 | 1.42987112 | 1 | 0.14947 | 1 |
| FGD2 | 0.08064225 | 1 | 1.505069 | 1 |
| TCEAL4 | 1.60072504 | 1 | -0.01106 | 1 |
| NCS1 | 1.60608168 | 1 | -0.01488 | 1 |
| HOXB2 | 1.39384394 | 1 | 0.200931 | 1 |
| OR13H1 | 0.95912885 | 1 | 0.639541 | 1 |
| L3MBTL1 | 0.21821231 | 1 | 1.381397 | 1 |
| VCP | 2.35550058 | 1 | -0.7552 | 1 |
| GPX1 | 1.23585141 | 1 | 0.365534 | 1 |
| HDAC10 | 0.34806029 | 1 | 1.254895 | 1 |
| ZNF528 | 0.12721652 | 1 | 1.479928 | 1 |
| DRAP1 | 1.50556149 | 1 | 0.102501 | 1 |
| POU2F2 | 0.56811432 | 1 | 1.041463 | 1 |
| ZDHHC18 | 1.87229619 | 1 | -0.26207 | 1 |
| LANCL2 | 0.84470112 | 1 | 0.76594 | 1 |
| ERCC2 | 1.35497004 | 1 | 0.255695 | 1 |
| CNIH3 | 0.24478644 | 1 | 1.366668 | 1 |
| EFS | 0.03131408 | 1 | 1.584659 | 1 |

| | | | | |
|---|---|---|---|---|
| F10 | 0.22840301 | 1 | 1.388296 | 1 |
| IRF2BP1 | 1.2851577 | 1 | 0.332968 | 1 |
| ADRA2B | 0.10420894 | 1 | 1.521211 | 1 |
| FFAR2 | 0.23726143 | 1 | 1.38869 | 1 |
| ILK | 2.74897334 | 1 | -1.12176 | 1 |
| CABIN1 | 0.68340783 | 1 | 0.946285 | 1 |
| PTPRH | 1.12589023 | 1 | 0.507178 | 1 |
| FNBP1 | 1.26305991 | 1 | 0.370224 | 1 |
| OR5P2 | 1.36968706 | 1 | 0.264237 | 1 |
| PORCN | 1.07372932 | 1 | 0.574488 | 1 |
| GP1BB | 1.2930917 | 1 | 0.356042 | 1 |
| FOXD2 | 0.23215974 | 1 | 1.418438 | 1 |
| EXOSC6 | 1.57072243 | 1 | 0.083029 | 1 |
| GNG3 | 0.31199063 | 1 | 1.342924 | 1 |
| ITPR2 | 1.27811225 | 1 | 0.377194 | 1 |
| HNRNPAB | 2.16172308 | 1 | -0.50585 | 1 |
| TBC1D9B | 1.09803063 | 1 | 0.558869 | 1 |
| ZNF580 | 1.32511034 | 1 | 0.334429 | 1 |
| LY6E | 0.31210907 | 1 | 1.348767 | 1 |
| SENP2 | 1.45006855 | 1 | 0.21369 | 1 |
| TBC1D21 | 1.0649407 | 1 | 0.604159 | 1 |
| HINT1 | 1.83756695 | 1 | -0.16713 | 1 |
| DPF2 | 0.60817559 | 1 | 1.062611 | 1 |
| NOS2 | 1.22591865 | 1 | 0.447869 | 1 |
| SHISA5 | 0.30114968 | 1 | 1.373845 | 1 |
| HIC2 | -1.2862714 | 1 | 2.96469 | 1 |
| IL22RA1 | 1.51618056 | 1 | 0.162876 | 1 |
| ZNF622 | 1.56615039 | 1 | 0.115406 | 1 |
| GNG7 | 1.28835846 | 1 | 0.40956 | 1 |
| MAFA | 1.17216296 | 1 | 0.528216 | 1 |
| SPHK2 | 1.08116368 | 1 | 0.62133 | 1 |
| TFPT | 1.00892918 | 1 | 0.695374 | 1 |
| CDC37 | 1.41103597 | 1 | 0.293354 | 1 |
| PILRA | 0.62284623 | 1 | 1.081674 | 1 |
| TNXB | 1.02733139 | 1 | 0.67731 | 1 |
| PPFIA1 | 0.29210897 | 1 | 1.41311 | 1 |
| TCF21 | 1.62897114 | 1 | 0.076559 | 1 |
| ABT1 | 1.54652824 | 1 | 0.160037 | 1 |
| DAB1 | 1.36472507 | 1 | 0.346584 | 1 |
| EBF1 | 0.80559562 | 1 | 0.906415 | 1 |
| SIN3B | 1.49736831 | 1 | 0.215105 | 1 |
| RHBDL1 | 0.41166666 | 1 | 1.304608 | 1 |
| LTK | 0.0775571 | 1 | 1.639912 | 1 |
| MAZ | 1.41869013 | 1 | 0.300208 | 1 |
| ZFHX2 | 0.33371922 | 1 | 1.3891 | 1 |
| GNAO1 | 1.32908403 | 1 | 0.394447 | 1 |

| | | | | |
|---|---|---|---|---|
| CHMP1A | 1.36243883 | 1 | 0.362919 | 1 |
| SMO | 0.16323827 | 1 | 1.568238 | 1 |
| STAT6 | 0.24821 | 1 | 1.485127 | 1 |
| S100A6 | 0.1182157 | 1 | 1.619877 | 1 |
| CD209 | 0.37810345 | 1 | 1.361551 | 1 |
| SHARPIN | 1.49550671 | 1 | 0.247498 | 1 |
| TBC1D10B | 0.50821267 | 1 | 1.236034 | 1 |
| SUMO1 | 2.06773954 | 1 | -0.32334 | 1 |
| TWIST1 | 0.32271419 | 1 | 1.42381 | 1 |
| TBXT | -0.0459488 | 1 | 1.802499 | 1 |
| BDKRB1 | 0.56427884 | 1 | 1.194273 | 1 |
| CNOT4 | 0.6585255 | 1 | 1.10203 | 1 |
| DACH1 | 0.78203351 | 1 | 0.981139 | 1 |
| MKNK2 | 0.77035588 | 1 | 0.994503 | 1 |
| GON4L | 1.24301689 | 1 | 0.528635 | 1 |
| ZNF763 | 0.46170695 | 1 | 1.311382 | 1 |
| SAFB2 | 0.23684005 | 1 | 1.538661 | 1 |
| SNF8 | 1.52034613 | 1 | 0.258528 | 1 |
| ZNF358 | 0.44287237 | 1 | 1.336341 | 1 |
| PICK1 | 0.36765712 | 1 | 1.412988 | 1 |
| TLE2 | 0.07709843 | 1 | 1.708685 | 1 |
| ZFHX4 | 0.14014509 | 1 | 1.646289 | 1 |
| NFYC | 1.54201762 | 1 | 0.252945 | 1 |
| XPC | 0.97001713 | 1 | 0.82499 | 1 |
| NPHS1 | 0.17883453 | 1 | 1.616532 | 1 |
| TRAF7 | 1.3703772 | 1 | 0.426931 | 1 |
| NEDD8 | 1.69540973 | 1 | 0.104644 | 1 |
| TCF7L1 | 0.95297577 | 1 | 0.847594 | 1 |
| SSX2 | 0.74014321 | 1 | 1.060858 | 1 |
| TPRA1 | 1.17031691 | 1 | 0.63121 | 1 |
| UBE2I | 1.07290708 | 1 | 0.731839 | 1 |
| MTSS2 | 0.60702879 | 1 | 1.198929 | 1 |
| FZD4 | 0.76030167 | 1 | 1.048588 | 1 |
| HEXIM2 | 0.0681938 | 1 | 1.748114 | 1 |
| NEUROD2 | 0.14459574 | 1 | 1.674819 | 1 |
| ADAM17 | 2.30291716 | 1 | -0.482 | 1 |
| RASGRP3 | 0.75964393 | 1 | 1.061914 | 1 |
| EEF1D | 1.67604803 | 1 | 0.147305 | 1 |
| TFEB | 0.19976654 | 1 | 1.625221 | 1 |
| SAP30L | 0.25310369 | 1 | 1.572691 | 1 |
| RTN1 | 0.95536333 | 1 | 0.871073 | 1 |
| ENG | 0.74179605 | 1 | 1.084746 | 1 |
| FOXI2 | 1.96839322 | 1 | -0.13865 | 1 |
| RNF213 | -0.1019558 | 1 | 1.941647 | 1 |
| TXK | 0.00893808 | 1 | 1.831246 | 1 |
| ZNF385A | 0.9905527 | 1 | 0.852394 | 1 |

| | | | | |
|---|---|---|---|---|
| NFIC | 0.84654295 | 1 | 0.998466 | 1 |
| BMP7 | 1.26940179 | 1 | 0.577913 | 1 |
| OTX2 | 1.06912531 | 1 | 0.778672 | 1 |
| MYD88 | 1.75481194 | 1 | 0.097867 | 1 |
| KCNH8 | 0.59970536 | 1 | 1.25486 | 1 |
| PIAS4 | 1.5653097 | 1 | 0.297766 | 1 |
| ZBTB22 | 1.54294934 | 1 | 0.322293 | 1 |
| KAT8 | 1.30990406 | 1 | 0.559439 | 1 |
| MAP3K10 | 1.55985097 | 1 | 0.311709 | 1 |
| UNC50 | 2.28177197 | 1 | -0.40277 | 1 |
| GPR132 | 0.09321264 | 1 | 1.792166 | 1 |
| RELB | 0.89790884 | 1 | 0.989782 | 1 |
| FOXK2 | 1.40048892 | 1 | 0.487795 | 1 |
| NMI | 0.22878711 | 1 | 1.659947 | 1 |
| DBNL | 0.44149448 | 1 | 1.45218 | 1 |
| KDM3B | 0.80539987 | 1 | 1.09317 | 1 |
| FHL5 | 0.89914218 | 1 | 1.001393 | 1 |
| PRMT2 | 0.95684268 | 1 | 0.948654 | 1 |
| GTF2F1 | 1.84961163 | 1 | 0.056075 | 1 |
| CCN2 | 1.12174498 | 1 | 0.789518 | 1 |
| NR1I2 | 0.134443 | 1 | 1.777189 | 1 |
| NACC1 | 1.33175579 | 1 | 0.581596 | 1 |
| DRG2 | 1.21902942 | 1 | 0.697846 | 1 |
| PDCD1 | 0.33320473 | 1 | 1.5894 | 1 |
| NACC2 | 0.89990639 | 1 | 1.024794 | 1 |
| HOXB9 | 0.86077739 | 1 | 1.064456 | 1 |
| NISCH | 1.16080343 | 1 | 0.765038 | 1 |
| OR11G2 | 1.14803154 | 1 | 0.780095 | 1 |
| RAC2 | 0.45109431 | 1 | 1.480142 | 1 |
| ZFP57 | 1.09141813 | 1 | 0.840099 | 1 |
| ELF3 | 0.1913924 | 1 | 1.744437 | 1 |
| GRN | 1.21181108 | 1 | 0.726899 | 1 |
| RIPK2 | 1.90233547 | 1 | 0.036441 | 1 |
| ATOH8 | 0.51169926 | 1 | 1.430422 | 1 |
| FYN | 0.24613191 | 1 | 1.698796 | 1 |
| MEF2D | 1.04351312 | 1 | 0.903181 | 1 |
| GATA2 | 0.87515595 | 1 | 1.080141 | 1 |
| NTSR1 | 0.520035 | 1 | 1.442062 | 1 |
| FBXW4 | 1.51251601 | 1 | 0.456165 | 1 |
| IL11RA | 0.73817031 | 1 | 1.231285 | 1 |
| MAP2K2 | 1.29210531 | 1 | 0.677846 | 1 |
| EOMES | 1.45756916 | 1 | 0.514344 | 1 |
| YOD1 | 2.06641214 | 1 | -0.09336 | 1 |
| NCR2 | 1.05718261 | 1 | 0.919001 | 1 |
| CYP24A1 | 0.06542571 | 1 | 1.913541 | 1 |
| NME2 | 2.09134178 | 1 | -0.11023 | 1 |

| | | | | |
|---|---|---|---|---|
| TRAK2 | 1.15552499 | 1 | 0.827477 | 1 |
| LCP2 | 0.03735434 | 1 | 1.946653 | 1 |
| ZDHHC13 | 2.31518561 | 1 | -0.33004 | 1 |
| HOXC4 | -0.006324 | 1 | 1.993686 | 1 |
| P2RY13 | 0.78124166 | 1 | 1.213132 | 1 |
| ARRDC2 | 0.36148716 | 1 | 1.641056 | 1 |
| S1PR1 | 0.90072254 | 1 | 1.103952 | 1 |
| ZNF784 | 1.64846939 | 1 | 0.35852 | 1 |
| ZNF581 | 1.63293095 | 1 | 0.374377 | 1 |
| BCAR1 | 0.91270865 | 1 | 1.094851 | 1 |
| AGT | 0.54697596 | 1 | 1.461961 | 1 |
| RAB13 | 0.5567523 | 1 | 1.45305 | 1 |
| UNCX | 0.98453139 | 1 | 1.027763 | 1 |
| AEBP1 | 1.31285086 | 1 | 0.704142 | 1 |
| HCAR1 | 1.6805544 | 1 | 0.344567 | 1 |
| SKOR1 | 0.61142108 | 1 | 1.416794 | 1 |
| NKX2.8 | 0.75119211 | 1 | 1.280423 | 1 |
| PQBP1 | 1.9718067 | 1 | 0.062379 | 1 |
| ETV7 | 0.24073788 | 1 | 1.79522 | 1 |
| FLII | 1.28103896 | 1 | 0.756473 | 1 |
| MSRB2 | 1.9516497 | 1 | 0.089584 | 1 |
| TNFSF13 | 0.14718579 | 1 | 1.894112 | 1 |
| KCNK10 | 0.54751113 | 1 | 1.495029 | 1 |
| ZFYVE19 | 1.39717849 | 1 | 0.646834 | 1 |
| ZNFX1 | 1.0495736 | 1 | 0.999789 | 1 |
| QRFPR | 0.1219844 | 1 | 1.929589 | 1 |
| IFNL1 | 1.3024283 | 1 | 0.749641 | 1 |
| FFAR1 | 0.14802761 | 1 | 1.906573 | 1 |
| ZSCAN22 | 1.18268308 | 1 | 0.87586 | 1 |
| NR1H2 | 1.06085702 | 1 | 1.000537 | 1 |
| ZSCAN31 | 0.82393148 | 1 | 1.239068 | 1 |
| CBY1 | 0.92012516 | 1 | 1.146957 | 1 |
| OR52J3 | 0.31184669 | 1 | 1.756161 | 1 |
| GREM2 | 0.04064674 | 1 | 2.027622 | 1 |
| ZNF641 | 0.57613407 | 1 | 1.505021 | 1 |
| PC | 1.13422222 | 1 | 0.948949 | 1 |
| EGLN2 | 1.23104489 | 1 | 0.85372 | 1 |
| KLF9 | 1.46400266 | 1 | 0.623118 | 1 |
| DEAF1 | 0.89859653 | 1 | 1.188658 | 1 |
| PLPP1 | 1.12402238 | 1 | 0.964056 | 1 |
| NR2E1 | 1.26013526 | 1 | 0.832152 | 1 |
| RAB5C | 1.92299159 | 1 | 0.170927 | 1 |
| UBA52 | 2.38318037 | 1 | -0.28832 | 1 |
| ADRB2 | 1.42605848 | 1 | 0.670623 | 1 |
| VPS72 | 1.61645233 | 1 | 0.48143 | 1 |
| SPN | 0.21268569 | 1 | 1.887715 | 1 |

| | | | | |
|---|---|---|---|---|
| PTGDR2 | 0.71925144 | 1 | 1.383789 | 1 |
| TAX1BP3 | 1.86538618 | 1 | 0.239528 | 1 |
| PDE11A | 0.33083489 | 1 | 1.781792 | 1 |
| GRK1 | 0.79021645 | 1 | 1.323188 | 1 |
| MLX | 2.17251318 | 1 | -0.05612 | 1 |
| ITPK1 | 0.95479662 | 1 | 1.164901 | 1 |
| RUVBL2 | 1.90450556 | 1 | 0.217278 | 1 |
| ELOB | 1.80439201 | 1 | 0.318189 | 1 |
| RAPGEF3 | 0.53861437 | 1 | 1.590067 | 1 |
| RADIL | 0.21131565 | 1 | 1.921275 | 1 |
| CSNK2B | 2.19026444 | 1 | -0.05729 | 1 |
| BHLHE23 | 0.91082813 | 1 | 1.225943 | 1 |
| TAOK2 | 1.11524383 | 1 | 1.028797 | 1 |
| MRPL12 | 2.05882922 | 1 | 0.08693 | 1 |
| CPE | 1.21440995 | 1 | 0.932777 | 1 |
| NOX1 | 1.78874369 | 1 | 0.360467 | 1 |
| LRRN2 | -0.0835798 | 1 | 2.234067 | 1 |
| CRY2 | 0.82341776 | 1 | 1.33318 | 1 |
| CALR | 1.27951253 | 1 | 0.882753 | 1 |
| NCOR2 | 1.57425602 | 1 | 0.589877 | 1 |
| PML | 0.94238787 | 1 | 1.224694 | 1 |
| OR4L1 | 1.19402435 | 1 | 0.973606 | 1 |
| RAB7B | 1.12390371 | 1 | 1.045831 | 1 |
| ZHX3 | 0.19889532 | 1 | 1.972144 | 1 |
| FARP2 | 1.02978287 | 1 | 1.151702 | 1 |
| GPR62 | 0.57476208 | 1 | 1.611871 | 1 |
| MAP3K3 | 0.1238082 | 1 | 2.062997 | 1 |
| SCGB1A1 | 0.88764079 | 1 | 1.301025 | 1 |
| TNK2 | -0.0772781 | 1 | 2.270325 | 1 |
| SRA1 | 1.68685256 | 1 | 0.507327 | 1 |
| NR5A2 | 0.31441379 | 1 | 1.883117 | 1 |
| RGS11 | 0.2603653 | 1 | 1.938187 | 1 |
| CD72 | 1.3740678 | 1 | 0.830279 | 1 |
| SPRY1 | 1.41559167 | 1 | 0.795344 | 1 |
| PSD2 | 0.59922203 | 1 | 1.617327 | 1 |
| FUT8 | 1.07144067 | 1 | 1.160552 | 1 |
| MPP2 | 1.71758461 | 1 | 0.5196 | 1 |
| HOXD11 | 1.1636445 | 1 | 1.074695 | 1 |
| ARHGAP26 | 1.38934716 | 1 | 0.859943 | 1 |
| AKAP7 | 0.77881337 | 1 | 1.476414 | 1 |
| GUCY1A1 | 1.21178288 | 1 | 1.047747 | 1 |
| IKBKG | 1.63938628 | 1 | 0.624642 | 1 |
| MAML3 | 0.9404888 | 1 | 1.331297 | 1 |
| OVOL3 | 0.27444077 | 1 | 2.001355 | 1 |
| ITGB1BP1 | 1.50721767 | 1 | 0.771328 | 1 |
| GRIA4 | 0.20067401 | 1 | 2.081765 | 1 |

| | | | | |
|---|---|---|---|---|
| MED31 | 0.38980135 | 1 | 1.901417 | 1 |
| LGALS1 | 0.93122682 | 1 | 1.36002 | 1 |
| EN2 | 1.20455061 | 1 | 1.08777 | 1 |
| PBXIP1 | -0.0485961 | 1 | 2.35716 | 1 |
| CD3G | 0.98898191 | 1 | 1.328243 | 1 |
| CDK5 | 1.83386169 | 1 | 0.484435 | 1 |
| APOL3 | 0.21298446 | 1 | 2.107821 | 1 |
| SERGEF | 1.1557282 | 1 | 1.166016 | 1 |
| CDKN1C | 0.5321629 | 1 | 1.794597 | 1 |
| SORBS3 | 0.18770033 | 1 | 2.144774 | 1 |
| FEZ2 | 1.24894363 | 1 | 1.085366 | 1 |
| MXD4 | 1.68751198 | 1 | 0.656204 | 1 |
| ZNF446 | 1.47286756 | 1 | 0.873375 | 1 |
| PDE6B | 0.28037656 | 1 | 2.075359 | 1 |
| RPS27A | 2.30077781 | 1 | 0.056572 | 1 |
| ZNF12 | 0.06905616 | 1 | 2.2968 | 1 |
| OR2M7 | 1.76485396 | 1 | 0.602709 | 1 |
| PRKCSH | 1.69975766 | 1 | 0.668205 | 1 |
| CTSH | 1.48181419 | 1 | 0.886979 | 1 |
| GLIS2 | 0.99494952 | 1 | 1.374095 | 1 |
| STUB1 | 2.14566508 | 1 | 0.225521 | 1 |
| SP100 | 0.50372625 | 1 | 1.876273 | 1 |
| ADRB1 | 0.62669767 | 1 | 1.756629 | 1 |
| TMEM102 | 0.75738333 | 1 | 1.627199 | 1 |
| CAMTA2 | 1.39908979 | 1 | 0.986005 | 1 |
| EVI5 | 1.1489853 | 1 | 1.23917 | 1 |
| MFN2 | 0.81636322 | 1 | 1.574871 | 1 |
| FOXL1 | 0.20129924 | 1 | 2.201032 | 1 |
| CCR9 | 1.41823672 | 1 | 0.984473 | 1 |
| AIFM3 | 1.29595213 | 1 | 1.115139 | 1 |
| POLR2I | 2.25089006 | 1 | 0.16047 | 1 |
| PCSK7 | 0.69860952 | 1 | 1.720784 | 1 |
| IGSF1 | 1.5906772 | 1 | 0.838049 | 1 |
| PAF1 | 0.26502959 | 1 | 2.165229 | 1 |
| PPP4C | 1.88417311 | 1 | 0.553555 | 1 |
| FRY | -0.0486394 | 1 | 2.500347 | 1 |
| PDAP1 | 2.09965787 | 1 | 0.352754 | 1 |
| EDF1 | 2.21587048 | 1 | 0.241086 | 1 |
| TCF3 | 2.50380125 | 1 | -0.04248 | 1 |
| MPP1 | 1.15225225 | 1 | 1.318383 | 1 |
| ZNF488 | -1.4614055 | 1 | 3.93927 | 0.5016578 |
| PIK3C2B | 0.81904245 | 1 | 1.658827 | 1 |
| GPR35 | 0.2880994 | 1 | 2.195288 | 1 |
| RPL7L1 | 1.35370689 | 1 | 1.133275 | 1 |
| PDE1A | 1.69653358 | 1 | 0.800602 | 1 |
| NR5A1 | 1.43468112 | 1 | 1.062789 | 1 |

| | | | | |
|---|---|---|---|---|
| PIM2 | 0.84965723 | 1 | 1.649393 | 1 |
| TBX3 | 1.1229707 | 1 | 1.378327 | 1 |
| VN1R1 | 0.18986089 | 1 | 2.312247 | 1 |
| ZNF408 | 2.38465009 | 1 | 0.126137 | 1 |
| THAP7 | 1.93686529 | 1 | 0.574618 | 1 |
| PHTF1 | 1.27540393 | 1 | 1.243173 | 1 |
| CSK | 1.75847895 | 1 | 0.763279 | 1 |
| RAMP1 | 0.99927219 | 1 | 1.527656 | 1 |
| ARHGAP40 | 0.95725953 | 1 | 1.577973 | 1 |
| CDK18 | 0.11110753 | 1 | 2.424999 | 1 |
| RAB34 | 2.35701279 | 1 | 0.181622 | 1 |
| DDX54 | 2.51600288 | 1 | 0.022888 | 1 |
| HOMEZ | 2.26986681 | 1 | 0.269352 | 1 |
| GSK3A | 1.59022134 | 1 | 0.954685 | 1 |
| ASXL2 | 2.09659763 | 1 | 0.450876 | 1 |
| EYA4 | 0.65167391 | 1 | 1.908771 | 1 |
| PCBD1 | 1.49077836 | 1 | 1.070224 | 1 |
| PDE3B | 1.47901555 | 1 | 1.085666 | 1 |
| KREMEN1 | 0.73202658 | 1 | 1.834551 | 1 |
| PSMD9 | 2.31783612 | 1 | 0.262208 | 1 |
| NFKBIB | 2.41758853 | 1 | 0.164613 | 1 |
| ARF5 | 2.43374771 | 1 | 0.151148 | 1 |
| SPOCK2 | 1.95801227 | 1 | 0.630151 | 1 |
| MYPOP | 1.78248491 | 1 | 0.824316 | 1 |
| FGFRL1 | 1.88476066 | 1 | 0.72829 | 1 |
| SP7 | 0.2183327 | 1 | 2.394728 | 1 |
| NACA | 2.65499712 | 1 | -0.03684 | 1 |
| ZBTB12 | 1.34696707 | 1 | 1.281346 | 1 |
| NRG2 | 0.59016585 | 1 | 2.050679 | 1 |
| ZBTB47 | 1.00041295 | 1 | 1.643979 | 1 |
| ADCYAP1R1 | 1.31987171 | 1 | 1.324551 | 1 |
| ZNF747 | 1.83310831 | 1 | 0.811947 | 1 |
| GATA6 | 1.23155358 | 1 | 1.42408 | 1 |
| PRKCD | 0.37272579 | 1 | 2.283963 | 1 |
| SERTAD1 | 1.43757485 | 1 | 1.235026 | 1 |
| STARD13 | 1.26887065 | 1 | 1.421867 | 1 |
| ZDHHC14 | 1.77617412 | 1 | 0.922125 | 1 |
| NDRG2 | 1.54257074 | 1 | 1.165456 | 1 |
| PLD2 | 0.16439687 | 1 | 2.544753 | 1 |
| TADA2A | 1.40824846 | 1 | 1.307045 | 1 |
| SPZ1 | 1.06804723 | 1 | 1.653019 | 1 |
| PCGF2 | 1.71750345 | 1 | 1.009653 | 1 |
| TYRO3 | 1.35871466 | 1 | 1.368598 | 1 |
| NTRK3 | 0.70100088 | 1 | 2.039644 | 1 |
| ELOF1 | 2.58230279 | 1 | 0.1668 | 1 |
| SIRT7 | 1.05184427 | 1 | 1.698974 | 1 |

| | | | | |
|---|---|---|---|---|
| RHBDL2 | 1.23200337 | 1 | 1.518875 | 1 |
| SLC35B2 | 1.78460187 | 1 | 0.97826 | 1 |
| DGKQ | 0.44047508 | 1 | 2.328161 | 1 |
| INSRR | 0.77344663 | 1 | 2.012088 | 1 |
| IGFBP6 | 1.69751867 | 1 | 1.094078 | 1 |
| STAT5A | 1.18003738 | 1 | 1.612963 | 1 |
| NR3C2 | 1.55402306 | 1 | 1.238987 | 1 |
| ENPP2 | 1.05982797 | 1 | 1.735203 | 1 |
| TNFAIP6 | 1.50061799 | 1 | 1.296648 | 1 |
| ZBTB7A | 2.30406249 | 1 | 0.49849 | 1 |
| RAB4B | 1.4499328 | 1 | 1.366391 | 1 |
| CAPS | 0.88397229 | 1 | 1.933458 | 1 |
| TSHR | 0.72789811 | 1 | 2.100127 | 1 |
| TAF10 | 1.72990206 | 1 | 1.099555 | 1 |
| PTCH1 | 0.96489821 | 1 | 1.865426 | 1 |
| PPP1R16B | 1.44133336 | 1 | 1.401394 | 1 |
| PINK1 | 0.24780699 | 1 | 2.598575 | 1 |
| RASL12 | 1.20910244 | 1 | 1.642876 | 1 |
| ARHGEF37 | 1.01894491 | 1 | 1.837806 | 1 |
| IKZF3 | 0.37622667 | 1 | 2.480651 | 1 |
| RALBP1 | 1.48675885 | 1 | 1.37685 | 1 |
| WSB2 | 1.00173106 | 1 | 1.862063 | 1 |
| P2RY12 | 2.089894 | 1 | 0.776392 | 1 |
| GIPC1 | 1.62752182 | 1 | 1.239101 | 1 |
| PTPRD | 1.63822881 | 1 | 1.237893 | 1 |
| PLA2G4C | 1.46202317 | 1 | 1.414677 | 1 |
| TNS2 | 0.4804552 | 1 | 2.420924 | 1 |
| GABRR1 | 0.89628991 | 1 | 2.00523 | 1 |
| OR5T2 | 0.79861775 | 1 | 2.113409 | 1 |
| TBX5 | 0.86757998 | 1 | 2.061646 | 1 |
| LTB4R2 | 2.03123093 | 1 | 0.904177 | 1 |
| GPR179 | 0.97052984 | 1 | 1.971947 | 1 |
| BTF3 | 2.98656425 | 1 | -0.04209 | 1 |
| POLR1H | 2.89159796 | 1 | 0.072007 | 1 |
| OR10H2 | 0.88702116 | 1 | 2.076674 | 1 |
| MTCH1 | 1.69378669 | 1 | 1.273254 | 1 |
| GABRB3 | 1.53973567 | 1 | 1.429481 | 1 |
| ERBB4 | 1.06360304 | 1 | 1.910456 | 1 |
| CD274 | 0.21278765 | 1 | 2.763844 | 1 |
| GABRG1 | 2.07281894 | 1 | 0.904437 | 1 |
| KLF16 | 2.2894819 | 1 | 0.698601 | 1 |
| SOX10 | 1.64581588 | 1 | 1.372376 | 1 |
| ZNF316 | 2.11593816 | 1 | 0.905776 | 1 |
| RHOF | 1.26188013 | 1 | 1.774108 | 1 |
| ZNF579 | 2.59121384 | 1 | 0.450651 | 1 |
| OR13C3 | 0.956866 | 1 | 2.085578 | 1 |

| | | | | |
|---|---|---|---|---|
| RRAS | 1.66070066 | 1 | 1.382396 | 1 |
| CASP7 | 1.55013097 | 1 | 1.493524 | 1 |
| CCL15 | 1.08718608 | 1 | 1.966223 | 1 |
| ITGBL1 | 0.76926785 | 1 | 2.295742 | 1 |
| OR9G9 | 1.80918395 | 1 | 1.261811 | 1 |
| DMRT2 | 1.12039142 | 1 | 1.958438 | 1 |
| RAB6B | 1.31071772 | 1 | 1.769489 | 1 |
| ZNF629 | 1.20799553 | 1 | 1.88778 | 1 |
| TOLLIP | 1.6167191 | 1 | 1.48717 | 1 |
| PDK4 | 1.57615738 | 1 | 1.52816 | 1 |
| ZC3H3 | 1.85301468 | 1 | 1.272859 | 1 |
| ZNF276 | 1.17472336 | 1 | 1.964864 | 1 |
| RPL22 | 3.30141614 | 1 | -0.15869 | 1 |
| RAB28 | 2.11866195 | 1 | 1.027217 | 1 |
| BARHL2 | 2.61665241 | 1 | 0.550722 | 1 |
| MYRF | 0.9300307 | 1 | 2.256667 | 1 |
| MAP2K3 | 1.92195227 | 1 | 1.277257 | 1 |
| ATN1 | 2.05421956 | 1 | 1.16386 | 1 |
| ZFY | 1.73854058 | 1 | 1.483972 | 1 |
| CNTNAP4 | 1.54111779 | 1 | 1.692183 | 1 |
| SGK2 | 1.62757341 | 1 | 1.605772 | 1 |
| ZNF511 | 2.39087307 | 1 | 0.854577 | 1 |
| RTKN | 0.82193473 | 1 | 2.441405 | 1 |
| SSTR5 | 1.40483245 | 1 | 1.859809 | 1 |
| RAB4A | 2.21902955 | 1 | 1.045955 | 1 |
| PLCD3 | 1.56902762 | 1 | 1.715627 | 1 |
| CSPG4 | 1.1978781 | 1 | 2.091625 | 1 |
| PFN1 | 2.70137118 | 1 | 0.592108 | 1 |
| SRF | 2.58895645 | 1 | 0.716837 | 1 |
| NAB2 | 1.62541046 | 1 | 1.684956 | 1 |
| S1PR5 | 0.72518869 | 1 | 2.597855 | 1 |
| TLR1 | 1.08053937 | 1 | 2.25603 | 1 |
| SLC1A2 | 1.6898592 | 1 | 1.654948 | 1 |
| BARX1 | 1.58227292 | 1 | 1.777527 | 1 |
| ATOH1 | 1.82926742 | 1 | 1.536271 | 1 |
| RRBP1 | 1.84021403 | 1 | 1.532424 | 1 |
| CITED4 | 1.60316737 | 1 | 1.77832 | 1 |
| MYT1 | 1.82731338 | 1 | 1.556379 | 1 |
| PTF1A | 0.91757127 | 1 | 2.474661 | 1 |
| PTPN1 | 2.15201882 | 1 | 1.245574 | 1 |
| OR52E2 | 1.37795714 | 1 | 2.025597 | 1 |
| ZSWIM4 | 1.41863869 | 1 | 1.994608 | 1 |
| RPL6 | 3.40550319 | 1 | 0.016756 | 1 |
| DIRAS2 | 1.68441496 | 1 | 1.750706 | 1 |
| PLEKHB1 | 1.1119157 | 1 | 2.329188 | 1 |
| HELZ2 | 1.85097208 | 1 | 1.60006 | 1 |

| | | | | |
|---|---|---|---|---|
| GRIN3B | 1.63492689 | 1 | 1.860956 | 1 |
| ZNF761 | 0.95407484 | 1 | 2.549422 | 1 |
| MUL1 | 2.13246756 | 1 | 1.404192 | 1 |
| KLK6 | 0.42072289 | 1 | 3.117412 | 1 |
| ATP8B1 | 1.79942332 | 1 | 1.75125 | 1 |
| CXCL10 | 0.65294359 | 1 | 2.905597 | 1 |
| SIT1 | 1.58108874 | 1 | 1.993949 | 1 |
| CD48 | 1.61585192 | 1 | 1.965385 | 1 |
| RAPGEF5 | 1.73483565 | 1 | 1.932009 | 1 |
| HOMER3 | 2.08816797 | 1 | 1.596551 | 1 |
| RPS3 | 3.86594691 | 0.679224994 | -0.17092 | 1 |
| PHB1 | 3.56559945 | 1 | 0.139583 | 1 |
| MEOX2 | 1.72856641 | 1 | 1.978747 | 1 |
| CRTC2 | 2.6784165 | 1 | 1.057385 | 1 |
| MPZL1 | 1.96546103 | 1 | 1.801553 | 1 |
| ADGRG1 | 2.8193098 | 1 | 0.991247 | 1 |
| PGR | 0.81630811 | 1 | 3.011652 | 1 |
| MAPK8IP3 | 1.66730899 | 1 | 2.219299 | 1 |
| HIPK2 | 1.31489126 | 1 | 2.574909 | 1 |
| OR5K3 | 1.92186365 | 1 | 2.007273 | 1 |
| TLR3 | 1.97929607 | 1 | 1.952265 | 1 |
| ZFYVE21 | 1.78317945 | 1 | 2.213644 | 1 |
| GPR55 | 0.47213095 | 1 | 3.525402 | 1 |
| RPL7 | 4.27009878 | 0.119928182 | -0.26046 | 1 |
| GPR37 | 2.54952416 | 1 | 1.477229 | 1 |
| ASB6 | 2.76438032 | 1 | 1.282084 | 1 |
| SBNO2 | 2.35188617 | 1 | 1.700299 | 1 |
| SNX17 | 3.72147212 | 1 | 0.355048 | 1 |
| ZDHHC1 | 2.22333435 | 1 | 1.860092 | 1 |
| TEF | 1.98455675 | 1 | 2.15278 | 1 |
| IRF9 | 2.13797239 | 1 | 2.025695 | 1 |
| LY96 | 1.32384482 | 1 | 2.859909 | 1 |
| NMUR2 | 1.0998332 | 1 | 3.087048 | 1 |
| STAP1 | 0.68599524 | 1 | 3.50528 | 1 |
| PKIG | 2.61103696 | 1 | 1.586545 | 1 |
| PDPK1 | 1.59854835 | 1 | 2.6246 | 1 |
| PMF1 | 2.15691814 | 1 | 2.096092 | 1 |
| ST18 | 0.84291695 | 1 | 3.418806 | 1 |
| ZNF846 | 1.01252163 | 1 | 3.294622 | 1 |
| SP110 | 1.3417391 | 1 | 3.001843 | 1 |
| SOX8 | 2.04650867 | 1 | 2.375357 | 1 |
| GFI1 | 1.12528134 | 1 | 3.361592 | 1 |
| FOXO4 | 2.65275012 | 1 | 1.860818 | 1 |
| HR | 2.08918887 | 1 | 2.482904 | 1 |
| LIF | 2.11548594 | 1 | 2.655689 | 1 |
| NKX6.2 | 1.86656668 | 1 | 2.98296 | 1 |

| | | | | |
|---|---|---|---|---|
| CRTC1 | 2.27364328 | 1 | 2.656115 | 1 |
| NRK | 2.48703311 | 1 | 2.607239 | 1 |
| RGS18 | 2.21020776 | 1 | 2.975704 | 1 |
| IFI16 | 2.0600983 | 1 | 3.164621 | 1 |
| APC2 | 2.66153499 | 1 | 2.610587 | 1 |
| PLEKHM1 | 3.27926139 | 1 | 2.005229 | 1 |
| TGFA | 2.69456931 | 1 | 2.648249 | 1 |
| MDFI | 2.03857298 | 1 | 3.390769 | 1 |
| THRA | 2.34614158 | 1 | 3.367955 | 1 |
| LDLRAP1 | 2.79884733 | 1 | 3.049315 | 1 |
| BST2 | 2.36467519 | 1 | 3.730621 | 1 |
| DBP | 2.65410479 | 1 | 3.445058 | 1 |
| ZC3HAV1 | 2.59260865 | 1 | 3.962187 | 0.455847 |
| PARP14 | 2.45070314 | 1 | 4.158152 | 0.1969274 |
| GPR17 | 2.42442391 | 1 | 4.21767 | 0.1515105 |
| IFI6 | 2.56205271 | 1 | 4.591296 | 0.027038 |
| IFITM1 | 2.1001939 | 1 | 5.136392 | 0.001719 |
| IRF7 | 2.75398931 | 1 | 4.683366 | 0.0173216 |
| STAT1 | 3.14537691 | 1 | 5.863447 | 2.78E-05 |
| BATF2 | 4.28513933 | 0.112095038 | 5.032557 | 0.0029707 |
| MX1 | 3.32835863 | 1 | 6.037775 | 9.59E-06 |

| | log2FoldChange_A-367 | log2FoldChange_KAPP |
|---|---|---|
| CEP55 | -2.670625487 | -1.763125359 |
| ASF1B | -3.099804772 | -1.29747228 |
| PRC1 | -2.648466861 | -1.575531374 |
| ASPM | -2.704685379 | -1.182380438 |
| MAD2L1 | -2.081044854 | -1.772675198 |
| CKAP2L | -2.258642608 | -1.389082608 |
| FANCI | -2.575532109 | -1.030873646 |
| DEPDC1 | -2.130404245 | -1.373093362 |
| BIRC5 | -2.115392941 | -1.372222137 |
| CENPE | -2.092576063 | -1.302567484 |
| CENPF | -2.224599983 | -1.15875039 |
| KIF18A | -1.879849926 | -1.480819768 |
| NCAPG | -2.159753954 | -1.193829485 |
| MKI67 | -2.152398949 | -1.109815448 |
| CDCA3 | -2.165360673 | -1.078356335 |
| DTL | -2.266125966 | -0.967011225 |
| RAD51AP1 | -2.21360722 | -0.992914878 |
| HMMR | -1.925258215 | -1.248233857 |
| TPX2 | -1.897774048 | -1.200668778 |
| KIF4A | -2.117863946 | -0.900177373 |
| NDC80 | -1.970785858 | -1.032549617 |
| DLGAP5 | -1.871725258 | -1.097384519 |
| TOP2A | -2.131435696 | -0.835512451 |
| FBXO5 | -1.994433289 | -0.959328104 |
| KIF15 | -2.080781784 | -0.85800709 |
| KIF11 | -2.177317077 | -0.750139317 |
| CENPK | -1.854626454 | -1.044115481 |
| FANCD2 | -2.204725398 | -0.682203267 |
| CYP2C18 | -1.859465552 | -1.025027642 |
| PBK | -2.087117928 | -0.770900981 |
| RRM2 | -2.195558991 | -0.605971636 |
| ECT2 | -1.774575105 | -1.012380202 |
| NEK2 | -1.923202602 | -0.843294627 |
| TROAP | -2.322695939 | -0.424614079 |
| CCNA2 | -1.843459403 | -0.90205445 |
| CCNB2 | -1.879903695 | -0.791970372 |
| BUB1 | -1.947397152 | -0.712577317 |
| KIF23 | -1.805466352 | -0.837175692 |
| ESCO2 | -1.930640452 | -0.711585415 |
| DSCC1 | -1.900130045 | -0.716682095 |
| SMC4 | -1.48004534 | -1.113970496 |
| KIF2C | -1.905846651 | -0.682783832 |
| AURKA | -1.776360341 | -0.790433095 |
| CDC25C | -1.958932771 | -0.575714932 |
| BRIP1 | -1.707751657 | -0.814484384 |

| | | |
|---|---|---|
| STIL | -1.957258251 | -0.528951982 |
| ARHGEF39 | -2.082445991 | -0.402978945 |
| CKAP2 | -1.69940814 | -0.742986375 |
| TTK | -1.648895072 | -0.775244072 |
| BUB1B | -1.959168704 | -0.45909119 |
| CDCA5 | -2.10078432 | -0.297967859 |
| CCNB1 | -1.767599658 | -0.62904895 |
| PARPBP | -1.664014169 | -0.584269589 |
| RACGAP1 | -1.667991131 | -0.571860589 |
| GTSE1 | -1.814745718 | -0.423708679 |
| PHF6 | -1.397361644 | -0.806899534 |
| EME1 | -1.529147245 | -0.65702082 |
| SPAG5 | -1.998788565 | -0.1268282 |
| CENPA | -1.554183973 | -0.541172779 |
| NCAPH | -1.605983622 | -0.474840026 |
| CDK1 | -1.606000176 | -0.465618165 |
| ERCC6L | -1.617184186 | -0.403714421 |
| TMPO | -1.256566315 | -0.758145906 |
| EXO1 | -1.560760495 | -0.426586986 |
| TRAIP | -1.506391639 | -0.463925708 |
| POC1A | -1.11884259 | -0.829640821 |
| MCM8 | -1.256032279 | -0.643534548 |
| NCAPD2 | -1.516101181 | -0.375029062 |
| TCF19 | -1.767591717 | -0.112723817 |
| CEP70 | -1.175321745 | -0.65311004 |
| SPC24 | -1.296411309 | -0.479970997 |
| HJURP | -1.2526701 | -0.379056158 |
| LMNB1 | -1.063491495 | -0.475514875 |
| ESPL1 | -1.535034109 | 0.033192055 |
| CDCA7 | -1.01747649 | -0.459861065 |
| PTTG1 | -1.779569625 | 0.305732773 |
| CENPM | -1.474655601 | 0.049405789 |
| GINS1 | -0.664299302 | -0.689523043 |
| MCM5 | -1.364252595 | 0.028320744 |
| EZH2 | -0.409739178 | -0.874108509 |
| ZNF280C | -0.386470481 | -0.894908676 |
| ZWINT | -0.285313296 | -0.798029634 |
| TOPBP1 | -0.641588784 | -0.34346359 |
| RFC4 | -0.746520161 | -0.115341045 |
| POLQ | -0.559790251 | -0.20936041 |
| CDT1 | -0.374315809 | 0.012619675 |
| CDC7 | 0.092013613 | 0.055893589 |
| E2F2 | -0.395789341 | 0.880617115 |

| | Antigen | Clone | Reactivity | Fluorophore | Company | Catalog number | Dilution |
|---|---|---|---|---|---|---|---|
| | CD155 | TX56 | Mouse | APC | Ebioscience | 17-1551-82 | |
| | cd96 | 3.3 | Mouse | APC | Biolegend | 131711 | |
| | TIGIT | IG9 | Mouse | APC | Proteintech | APC-65079 | |
| | DNAM1 | 10E5- | Mouse | PE-Cy7 | Biolegend | 128811 | |
| | B7-H3 | RTAA15 | Mouse | PE | Biolegend | 124507 | |
| | CD45 | 30-F11 | Mouse | Pacific Blue | Biolegend | 103126 | |
| | CD8a | 53-6.7 | Mouse | APC | Ebioscience | 17-0081-82 | |
| | CD25 | PC61 | Mouse | BV421 | Biolegend | 102043 | |
| | CD44 | IM7 | Mouse | BUV496 | BD | 741057 | |
| | CD69 | H1.2F3 | Mouse | PE-CY7 | Biolegend | 104511 | |
| | CD112 | 829038 | Mouse | BUV650 | BD | 748048 | |
| | Interferon gamma | XMG1.2 | Mouse | APC | Biolegend | 505809 | |
| | Granzyme B | QA16A02 | Mouse/Human | PE | Biolegend | 372207 | |
| | CD45 | 2D1 | Human | FITC | BD | 345808 | |
| | HLA-ABC | W6/32 | Human | FITC | Biolegend | 311404 | |
| | CD155 | SKII.4 | Human | APC | Biolegend | 337618 | |
| | B7-H3 | DCN.70 | Human | PE | Biolegend | 331605 | |
| | CD155 | SKII.4 | Human | | Biolegend | 337602 | |
| | B7-H3 | DCN.70 | Human | - | Biolegend | 331602 | |
| | 4-1BB/CD137L | 5F4 | Human | - | Biolegend | 311502 | |
| | CD86 | IT2.2 | Human | - | Biolegend | 305401 | |
| | CD66a/c/e | ASL-32 | Human | - | Biolegend | 342302 | |
| | CD252/OX40 | RM134L | Human | - | Biolegend | 108802 | |
| | Galectin 9 | 9M1-3 | Human | - | Biolegend | 348902 | |
| | PD-L2 | 24F.10C12 | Human | - | Biolegend | 329602 | |
| secondary antibody | goat-anti-mouse | Poly4053 | mouse | PE | Biolegend | 405307 | |
| | Annexin V | - | - | APC | Biolegend | 640920 | |

| mouse | TRC Clone ID/Plasmid ID | Company | target sequence |
|---|---|---|---|
| CD155 1 | TRCN0000112545 | Millipore/Sigma | CCCAGACACTATTCTGACTTA |
| CD155 2 | TRCN0000112548 | Millipore/Sigma | CGTCCAGTATTCATCTGTGAA |
| FOXM1 1 | TRCN0000304361 | Millipore/Sigma | ACTTCCTATTCAGTCCATTAA |
| FOXM1 2 | TRCN0000304362 | Millipore/Sigma | ACTTAGAGAGGCCTATCAAAG |
| human | | | |
| CD155 1 | TRCN0000062909 | Millipore/Sigma | CTAATGGGCATGTCTCCTATT |
| CD155 2 | TRCN0000062912 | Millipore/Sigma | CGGCAAGAATGTGACCTGCAA |
| negative controls | | | |
| scramble shRNA | Addgene 1864 | Addgene | CCTAAGGTTAAGTCGCCCTCGCTCGAGCGAGGGCGACTTAACCTTAGG |
| non-target control | SHC002 | Millipore/Sigma | CCGGCAACAAGATGAAGAGCACCAACTCGAGTTGGTGCTCTTCATCTTGTTGTTTTT |

Primers qRT-PCR

| Gene | Species | Forward | Reverse |
|---|---|---|---|
| CD155 | mouse | CGGGTGGGGATATACGTGTG | ACTACAGTGCAAGGTGGTCG |
| FOXM1 | mouse | CCAAAAGTCAAGCCTGAGCC | ACTGCTCAGGACAACCTGAAA |
| ROCK2 | mouse | AACTGTGATCCCAAGGGAAGG | ACTGACAGCAGCAGTATGCC |
| EGFR | mouse | AAGTCATCATGCAGCAGTGG | AGCATCAAGCAGGCATTCTGT |
| SH2D3C | mouse | CAGCCACATGCTAGCAAACC | TTCAAAGGTGTCAGGCTCCG |
| ITGA4 | mouse | CCAGGCATTCATGCGGAAAG | ATGCCCAAGGTGGTATGTGG |
| beta Actin | mouse | CCGAGCGTGGCTACAGCTTC | ACCTGGCCGTCAGGCAGCTC |
| HPRT | mouse | AGTCCCAGCGTCGTGATTAG | TTTCCAAATCCTCGGCATAATGA |

Western Blot Antibodies

| Antigen | Host species | Reactivity | Clone | Dilution | Company |
|---|---|---|---|---|---|
| Integrin alpha 4 | rabbit | mouse | 2DE1 | 1::2000 | Cell Signaling |
| FOXM1 | rabbit | mouse | EPR17379 | 1::1000 | Abcam |
| Phospho-p44/42 MAPK (Erk1/2) (Thr202/Tyr204) | mouse | mouse | E10 | 1::2000 | Cell Signaling |
| p44/42 MAPK (Erk1/2) | rabbit | mouse | polyclonal | 1::2000 | Cell Signaling |
| beta-Actin | mouse | mouse | 8H10D10 | 1::2000 | Cell Signaling |
| CD155 | rabbit | mouse | polyclonal | 1::750 | Abcam |

**Supplementary Table 1:**

Patient characteristics of our brain tumor patient cohort (median age and range; number of patients and percentages for further data; n.k.= not known).

**Supplementary Table 2:**

Flow cytometry data of brain tumor patients (percentage of positive cells, as well as CD45-negative and CD45-positive cells).

**Supplementary Table 3:**

Z score of upstream regulators following IPA analysis of gene expression data from CD155-negative and control A-367 and KAPP cells.

**Supplementary Table 4:**

List of differentially regulated proteins / master regulators in CD155-deficient versus control A-367 and KAPP cells (sh CD155 / sh CTR); Shown is the normalized expression score (NES) and the adjusted p value per cell line.

**Supplementary Table 5:**

FOXM1-related genes differentially expressed in CD155-deficient versus control A-367 and KAPP cells (sh CD155 / sh CTR)

**Supplementary Table 6:**

Antibodies used in flow cytometry experiments

**Supplementary Table 7:**

shRNAs used in lentiviral transduction experiments

**Supplementary Table 8:**

Primers used in qRT-PCR experiments

**Supplementary Table 9:**

Antibodies used in Western Blot experiments
